# Phenotypic high-throughput screening identifies modulators of gut microbial choline metabolism

**DOI:** 10.1101/2024.11.08.621386

**Authors:** Amelia Y. M. Woo, Walter J. Sandoval–Espinola, Maud Bollenbach, Alison Wong, Mariko Sakanaka–Yokoyama, Qijun Zhang, Vincent Nieto, Federico E. Rey, Emily P. Balskus

## Abstract

Anaerobic metabolism of dietary choline to trimethylamine (TMA) by the human gut microbiome is a disease-associated pathway. The host’s impaired ability to oxidize TMA to trimethylamine-*N*-oxide (TMAO) results in trimethylaminuria (TMAU), while elevated serum TMAO levels have been positively correlated with cardiometabolic disease. Small molecule inhibition of gut bacterial choline metabolism attenuates the development of disease in mice, highlighting the therapeutic potential of modulating this metabolism. Inhibitors previously developed to target this pathway are often designed to mimic choline, the substrate of the key TMA-generating enzyme choline trimethylamine-lyase (CutC). Here, we use a growth-based phenotypic high-throughput screen and medicinal chemistry to identify distinct chemical scaffolds that can modulate anaerobic microbial choline metabolism and lower TMAO levels *in vivo*. These results illustrate the potential of using phenotypic screening to rapidly discover new inhibitors of gut microbial metabolic activities.

## Introduction

The human gut microbiome has been increasingly associated with host physiology, and this has prompted the development and application of various approaches to manipulate the composition and functions of this microbial community^1^. These strategies include fecal microbiome transplant (FMT), use of antibiotics, prebiotics, wild-type or genetically modified probiotics, and phage therapy^2,3^. Small molecule inhibitors that confer precise control over specific metabolic activities broadly across multiple gut microbial species, without exhibiting broad-spectrum antimicrobial activity, are ideal not just for therapeutic applications, but also for basic studies of the gut microbiome^4,5^. Changes in host phenotypes induced by treatment with gut microbiome-targeted small molecule inhibitors in animal models could be used to infer the causality of host–microbiome associations and could provide critical support for microbiome-targeted therapeutic interventions^6^.

Efforts to develop small molecule inhibitors as tools to study the gut microbiome have accelerated over the last decade, beginning with the seminal example of bacterial β-glucuronidase inhibitors^7–11^. Other key examples include inhibitors of choline trimethylamine-lyase^12–15^, amino acid decarboxylases^16,17^, bile salt hydrolases^18,19^, and tryptophanases^20^. However, inhibitor discovery in this space has primarily relied on having a biochemically well-characterized target for inhibitor development. This sharply contrasts with antibiotic^21^, anti-virulence^22–24^, and anti-infective^25–27^ drug discovery, in which target-agnostic, phenotypic screening campaigns are common.

While underutilized in the context of the gut microbiome, phenotypic screening offers several advantages over target-based inhibitor discovery strategies^28^. First, the requirement for compounds to access their targets in a cellular setting is built into the screen. This may be especially important when targeting metabolic activities that are widespread across bacterial phyla, as differences in cell membrane architectures can significantly influence the permeability of small molecules^29^. Secondly, such screens can be performed with more complex microbial communities and gut microbiome samples^30^, which may provide results that are more physiologically relevant, such as in the case of phenotypic screens in organoids^31^. Lastly, phenotypic screens are target-agnostic and therefore provide an opportunity to both discover new biologically relevant targets and shed light on previously unappreciated aspects of microbial function.

One disease-associated gut metabolic activity of interest is the anaerobic catabolism of choline to trimethylamine (TMA)^32–34^. TMA is an exclusively microbial metabolite in the body, with choline as its major precursor^32,35^. TMA generated by the gut microbiome is typically oxidized in the liver by flavin monooxygenase 3 (FMO3) enzymes to form trimethylamine-*N*-oxide (TMAO), which is then excreted^36^ (Figure 1A). However, individuals who have an impaired ability to oxidize TMA excrete this malodorous metabolite and suffer from a socially debilitating condition called trimethylaminuria (TMAU)^37^. Multiple studies have also found that high levels of circulating serum TMAO in humans strongly correlate with the incidence of cardiometabolic diseases such as cardiovascular disease^38–40^, chronic kidney disease^41,42^, type II diabetes^43^ and non-alcoholic fatty liver disease^44^. Mice that have been fed with high choline or high TMAO diets develop symptoms associated with cardiovascular disease and have more severe heart failure^45^, and this pathway has also been shown to play a causal role in atherosclerosis development in ApoE knockout mice^38^. Moreover, the metabolism of choline by gut microbes can also reduce availability of this essential nutrient for the host^46^. Due to the prevalence of diseases linked to TMA, there is strong interest in developing inhibitors of gut microbial choline metabolism as potential therapeutics. Besides the opportunity for drug development, small molecules that can selectively inhibit anaerobic choline metabolism in complex gut microbiomes would be useful tools for understanding this metabolic activity, as there remains questions regarding how dietary choline impacts chronic plasma TMAO levels^47^.

**Figure 1.**
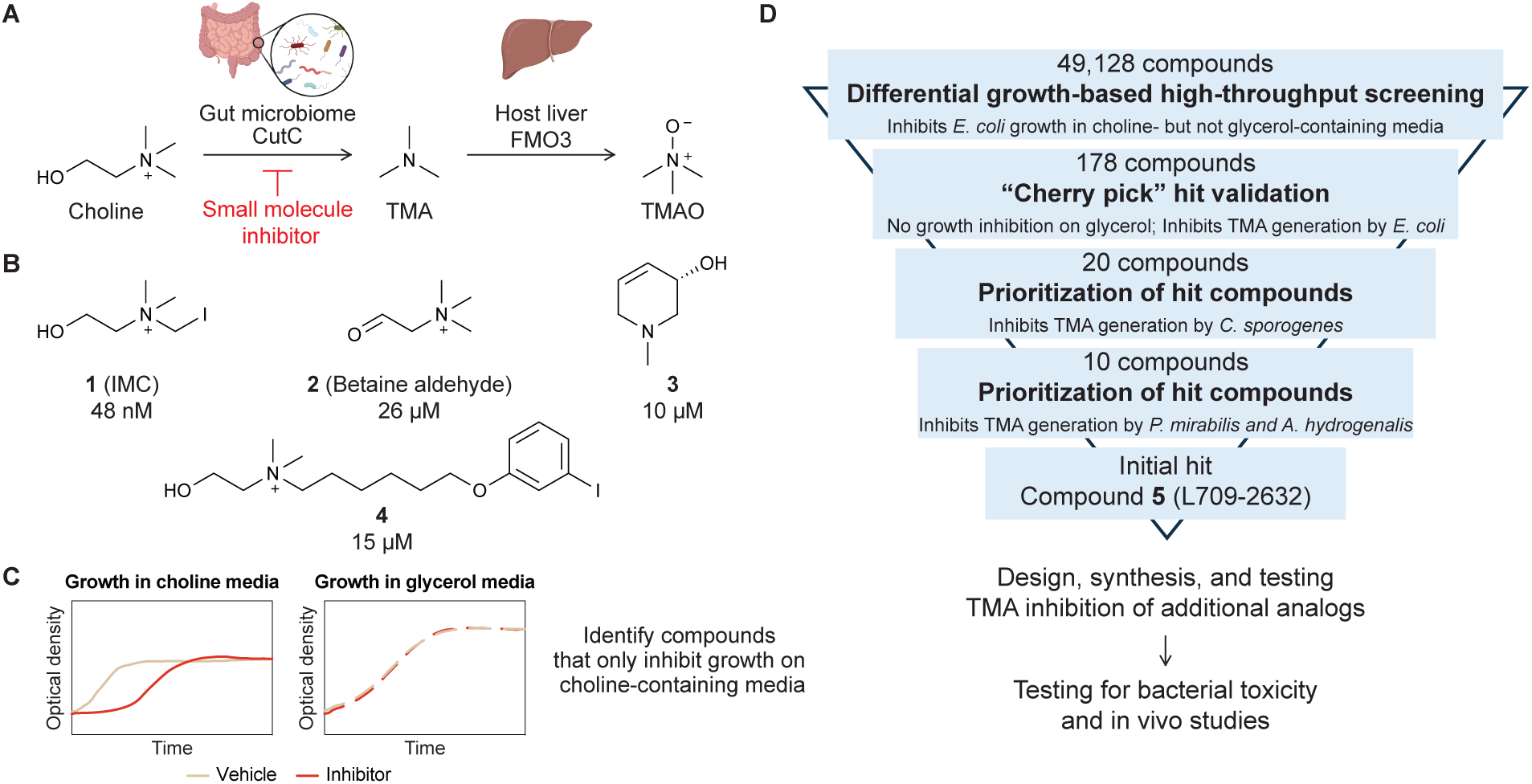
Small molecule inhibitors of microbial choline metabolism can be identified through phenotypic screening. (A) Dietary choline is metabolized by the gut microbial enzyme CutC into TMA, which is further oxidized to TMAO by host FMO3 enzymes in the liver. Both TMA and TMAO are disease-associated metabolites, and generation of these metabolites can potentially be reduced by targeting gut microbial choline metabolism with small molecule inhibitors. (B) Chemical structures of reported inhibitors of anaerobic choline metabolism and their IC_50_ values against bacterial cultures. (C) Small molecule inhibitors of choline metabolism can be identified by comparing bacterial growth on minimal media supplemented with choline or an alternate carbon and energy source. (D) Overall inhibitor development workflow began with identifying inhibitors using high-throughput phenotypic screening for growth inhibition of *E. coli* MS 200-1 followed by validation and prioritization of compounds based on their EC_50_ for inhibition of TMA production from choline across multiple choline-metabolizing bacteria. New analogs were designed, synthesized, and iteratively tested for their TMA inhibitory activity, and promising compounds were evaluated for their bacterial toxicity and *in vivo* activity.

Anaerobic choline metabolism is widespread across bacterial phyla, and this activity has been observed in both gram-positive and gram-negative gut bacteria^48^. This metabolism requires enzymes encoded within the choline utilization (*cut*) gene cluster^49^, including the glycyl radical enzyme choline trimethylamine-lyase (CutC), which performs the chemically challenging C–N bond cleavage of choline to generate TMA and acetaldehyde^50^. Bacteria that encode and express the *cut* gene cluster metabolize the acetaldehyde byproduct into ethanol and acetyl-CoA, enabling energy production, while TMA is excreted. As such, CutC has been the canonical target for the development of inhibitors of anaerobic choline metabolism.

Structural mimics of choline such as halomethylcholines^13^, betaine aldehyde (**2**)^14^, and cyclic choline analogs (**3**)^15^ have been identified based on the structure of CutC and its proposed mechanism^50,51^ and shown to be potent inhibitors of anaerobic choline metabolism both *in vitro* and *in vivo*^13,52–54^ (Figure 1B). Iodomethylcholine (**1**, IMC), one of the most potent CutC inhibitors, has a half-maximal inhibitory concentration (IC_50_) of 1.5 nM against *Proteus mirabilis* lysates^13^. In animal models of disease, IMC has also been shown to reduce renal tubulointerstitial fibrosis^52^, reduce markers of renal injury^53^ and improve cardiac function^54^. While IMC is found at around 6 mM in mice cecum and colon samples and is thus largely retained in the gut, the inhibitor and its metabolites can be detected at low micromolar levels in serum^13^. IMC has previously been shown to be a potent inhibitor of choline transport in rat synaptosomes with an IC_50_ of 0.49 µM and is thought to bind irreversibly to the high affinity choline transporter (HAChT)^55^, which raises concerns of potential neurotoxicity and motivates the discovery of additional inhibitors.

To our knowledge, there has only been one prior attempt to identify inhibitors of anaerobic choline metabolism using phenotypic screening. A matrix-assisted laser desorption/ionization time-of-flight (MALDI–TOF) mass spectrometry-based phenotypic screen measured derivatized TMA in bacterial cultures and identified an inhibitor of a human choline transporter (**4**) as a modulator of anaerobic choline metabolism with an IC_50_ of 15 µM in *Hungatella hathewayi* cultures^56^. This compound inhibits choline transport in human cells that overexpress choline transporters with an IC_50_ of 44 nM^57^.

Given the structural similarity of reported inhibitors to choline and their potential ability to interact with host choline receptors and transporters, we sought to identify inhibitors of microbial choline metabolism with more drug-like scaffolds. Here, we identify novel, broad-spectrum inhibitors of anaerobic microbial choline metabolism using a target-agnostic differential growth-based high-throughput screen (HTS). Our inhibitor optimization efforts reveal that a cyclic amine is essential for activity. These compounds are structurally distinct from previously reported CutC inhibitors. They are non-lethal to both gram-positive and gram-negative commensal bacteria and have improved broad-spectrum activity compared to our initial hit. Finally, we demonstrate that these inhibitors can lower TMAO levels in gnotobiotic and conventional mouse models. We anticipate that this type of growth-based HTS screening workflow can be readily adapted to identify compounds that modulate additional gut bacterial metabolic activities, including those with no known associated enzyme targets.

## Results

### Differential growth as a phenotype for high-throughput screening

To identify inhibitors of choline metabolism in a target-agnostic manner, we envisioned using a phenotypic HTS with choline-metabolizing bacteria. The previous phenotypic screen for choline metabolism inhibitors used high-throughput quantitation of derivatized TMA by MALDI–TOF mass spectrometry to screen a library of 10,229 compounds^56^. While this method successfully identified compounds that inhibit the transformation of choline to TMA, this approach has several disadvantages, including a requirement for specialized instrumentation, the need for chemical derivatization, and analysis time that scales considerably with library size. Microbial growth is a more straightforward readout for screening, as optical density of cultures can be measured quickly with a plate reader in a high-throughput fashion. This type of screen is routinely used for antibiotic discovery but is rarely used to identify inhibitors of specific microbial metabolic activities because many functions of interest are not readily linked to bacterial growth^58,59^.

Inspired by our previous data showing that a rationally designed CutC inhibitor could inhibit bacterial growth on choline^14^, we conceived of an alternative phenotypic screening approach. Anaerobic choline metabolism is a non-essential pathway when alternate carbon and energy sources are available. However, it can become essential when choline is the sole carbon and energy source during anaerobic growth^46^. Thus, choline metabolism can be linked directly to a growth phenotype to enable a HTS. To identify and remove broad-spectrum bacterial growth inhibitors from consideration, such as antibiotics and antimicrobials, we envisioned performing a counter-screen using the same minimal medium containing an alternative carbon and energy source. In this format, compounds that specifically interfere with anaerobic choline metabolism should inhibit bacterial growth in choline-containing medium but not in the alternate medium (Figure 1C). This differential growth-based screening strategy is straightforward to implement, making it easily scalable to screen large compound libraries and readily extendable to other organisms and metabolic activities.

We chose to use the choline-metabolizing strain *Escherichia coli* MS 200-1 for the HTS as it is a human gut isolate and can be grown anaerobically at 37 °C in No Carbon Essential (NCE) medium supplemented with choline^46^. This medium contains fumarate as the terminal electron acceptor and can be supplemented with other carbon sources. Using kanamycin and DMSO as the positive and negative controls, respectively, we optimized the assay in a 384-well plate format to have Z’ values > 0.5, making it suitable for the HTS^60^. We chose to measure optical densities of the cultures at 7 h and 20 h after inoculation to identify partial (decreased growth rate) or total growth inhibitors in choline-containing media. A simultaneous counter-screen with glycerol-containing NCE medium would also be performed. Thus, compounds that inhibited *E. coli*’s growth on choline but not glycerol NCE media, would be considered potential hits.

We then planned to validate the activity of potential hits toward *E. coli* MS 200-1 by testing both their growth-inhibitory effects in NCE glycerol medium and their TMA-inhibitory activity. Half-maximal effective concentration (EC_50_) values of TMA inhibition were calculated after measurement using a liquid chromatography–mass spectrometry (LC–MS/MS)-based assay. We planned to further prioritize compounds to test against additional choline-metabolizing gut isolates based on their EC_50_ values. Direct measurement of TMA inhibition would be a key step in our HTS efforts because the growth-based screen was not designed to differentiate compounds that specifically inhibit TMA generation from compounds that may inhibit the downstream metabolism of acetaldehyde, which is also linked to growth. Scaffolds identified through this screening would be further optimized for their potency for TMA inhibition using medicinal chemistry. Finally, the most potent analogs would be evaluated for their toxicity against a panel of gut bacterial isolates and their ability to lower TMAO levels *in vivo* (Figure 1D).

### Growth-based phenotypic HTS identifies new inhibitor scaffolds

The primary screen was performed in duplicate using the ChemDiv7 commercial library from the Institute of Chemistry and Cell Biology–Longwood (ICCB-L) Screening Facility, consisting of 49,128 compounds (Figure 1D). Each compound was tested at a concentration of 20 µg/mL. As described above, compounds that differentially inhibited *E. coli* MS 200-1 growth on choline (Z-score <–2) but not glycerol (Z-score ≥0) media at either the 7 h or 20 h time points were considered potential hits (Figure 2A).

**Figure 2.**
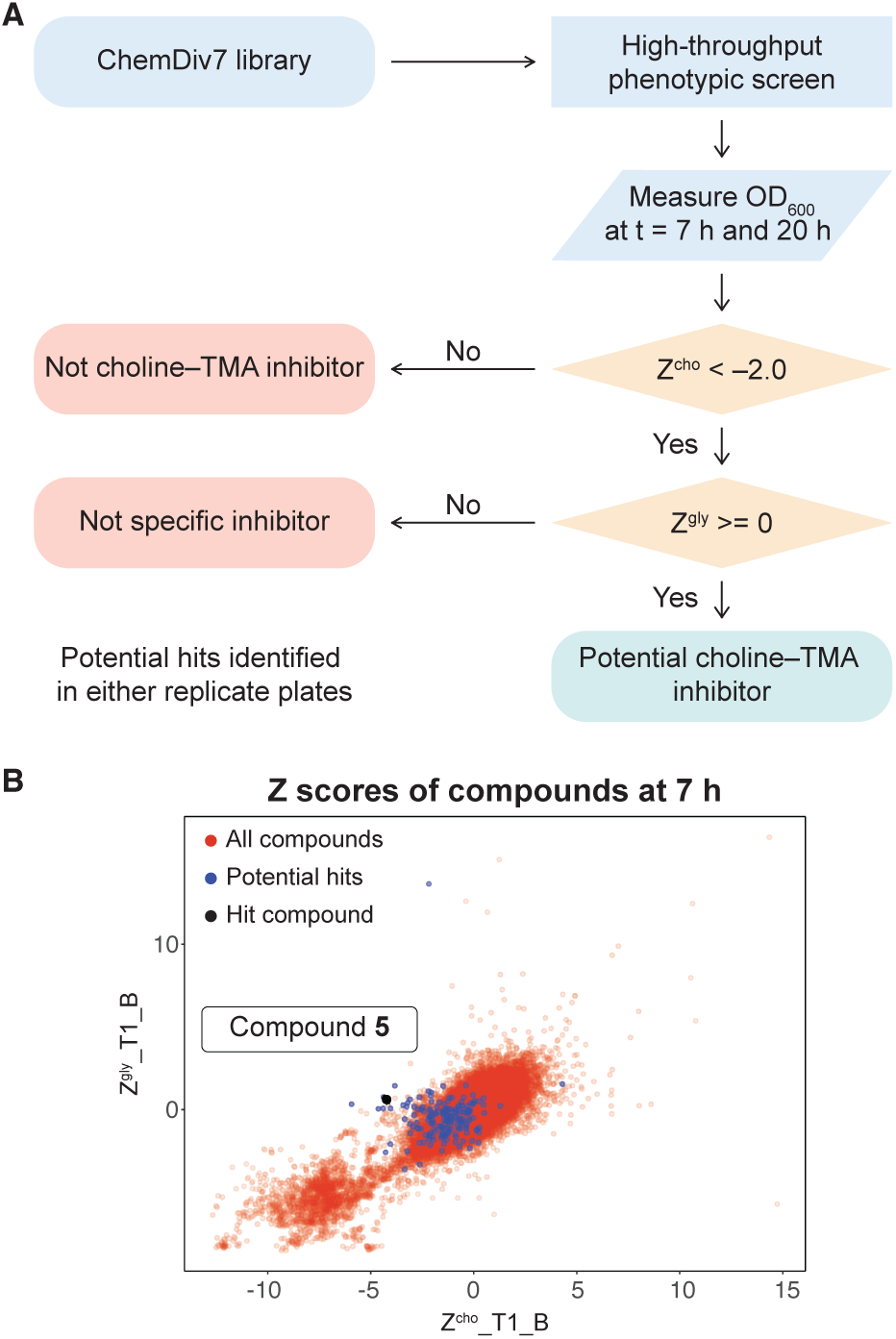
Differential growth-based phenotypic screening identified inhibitors of anaerobic gut bacterial choline metabolism. (A) Flowchart of data analysis pipeline to identify potential hit compounds. (B) Representative scatter plot of one replicate plate comparing Z-scores of compounds in choline medium versus glycerol medium at 7 h. Compounds that met the criteria on either replicate plate and chosen for retesting are colored in blue. Hit compound **5** is labeled and colored in black. Positive and negative control wells with kanamycin and DMSO vehicle were not given Z-scores.

The HTS identified 60 compounds that differentially inhibited growth on choline at 7 h and 122 compounds that differentially inhibited growth at 20 h on either replicate plates. Of these, 4 compounds differentially inhibited growth at both 7 h and 20 h. We next performed a “cherry pick” screen (i.e. hit validation) with 178 compounds, or 0.362% of the ChemDiv7 library (Figure 2B). Comparatively, when screening a smaller library of 10,229 compounds, the MALDI–TOF method had a higher hit rate of 2.5%^56^, suggesting that this growth-based screening strategy is not only easier to scale but also may afford fewer false positives.

To validate the potential hits from the HTS, we tested the 178 compounds for inhibition of growth of *E. coli* MS 200-1 in glycerol minimal media at 30 µg/mL, following up with those without activity. As the number of compounds re-evaluated was significantly fewer than the primary screen, we defined growth inhibition as cultures that had an optical density below 5% standard deviation from the mean optical density of the plate, rather than using Z-scores. We then determined EC_50_ values for inhibition of TMA production from choline by bacterial cultures incubated with [trimethyl-*d*_9_]choline in phosphate-buffered saline. Of the 178 compounds tested in both growth and TMA inhibition assays, 20 inhibited TMA generation by *E. coli* MS 200-1 without growth inhibitory effects in glycerol minimal medium.

We then used secondary assays to prioritize scaffolds that had broad-spectrum TMA inhibitory activity against additional choline-metabolizing gut bacteria. For this, we used previously identified choline-metabolizing human gut isolates *Clostridium sporogenes* ATCC 15579, *Anaerococcus hydrogenalis* DSM 7454 and *Proteus mirabilis* ATCC 29906^61^. This group of strains included both gram-positive and gram-negative bacteria. Gram-negative bacteria have an additional outer membrane layer and associated efflux pumps^29^, which typically decreases small molecule permeability as compared to gram-positive bacteria. It also included bacteria with *cut* gene cluster architectures that differ in the number and types of microcompartment proteins encoded and the size of the CutC enzyme^48^. We hypothesized that testing the inhibitors against multiple choline-utilizing gut bacteria would increase the likelihood of identifying compounds with broad-spectrum activity.

This series of secondary assays identified compound **5** as a promising lead for medicinal chemistry efforts (Figure 2B). This compound displayed the best potency across three of the four choline-metabolizers, with EC_50_ values for TMA inhibition ranging from 12 to 33 µM, and it had no impact on growth of *E. coli* MS 200-1 in glycerol medium (Figure 3D). Compound **5** has a pyrrole-3-carboxamide as its core scaffold, with a phenyl group at C5 of the pyrrole and an amine linked through the carboxamide. **5** was an attractive starting point for our medicinal chemistry efforts as it was more drug-like than existing inhibitors and analogs were synthetically accessible. Testing commercially available analogs of **5** identified inhibitor **6**, which was active across all four choline-metabolizers tested (Figure 3B–D). Compound **6** had an EC_50_ of 9 µM toward against *E. coli* in our LC–MS/MS-based TMA inhibition assay and was subsequently used as our reference compound when comparing inhibition of TMA generation in downstream EC_50_ assays across additional analogs.

**Figure 3.**
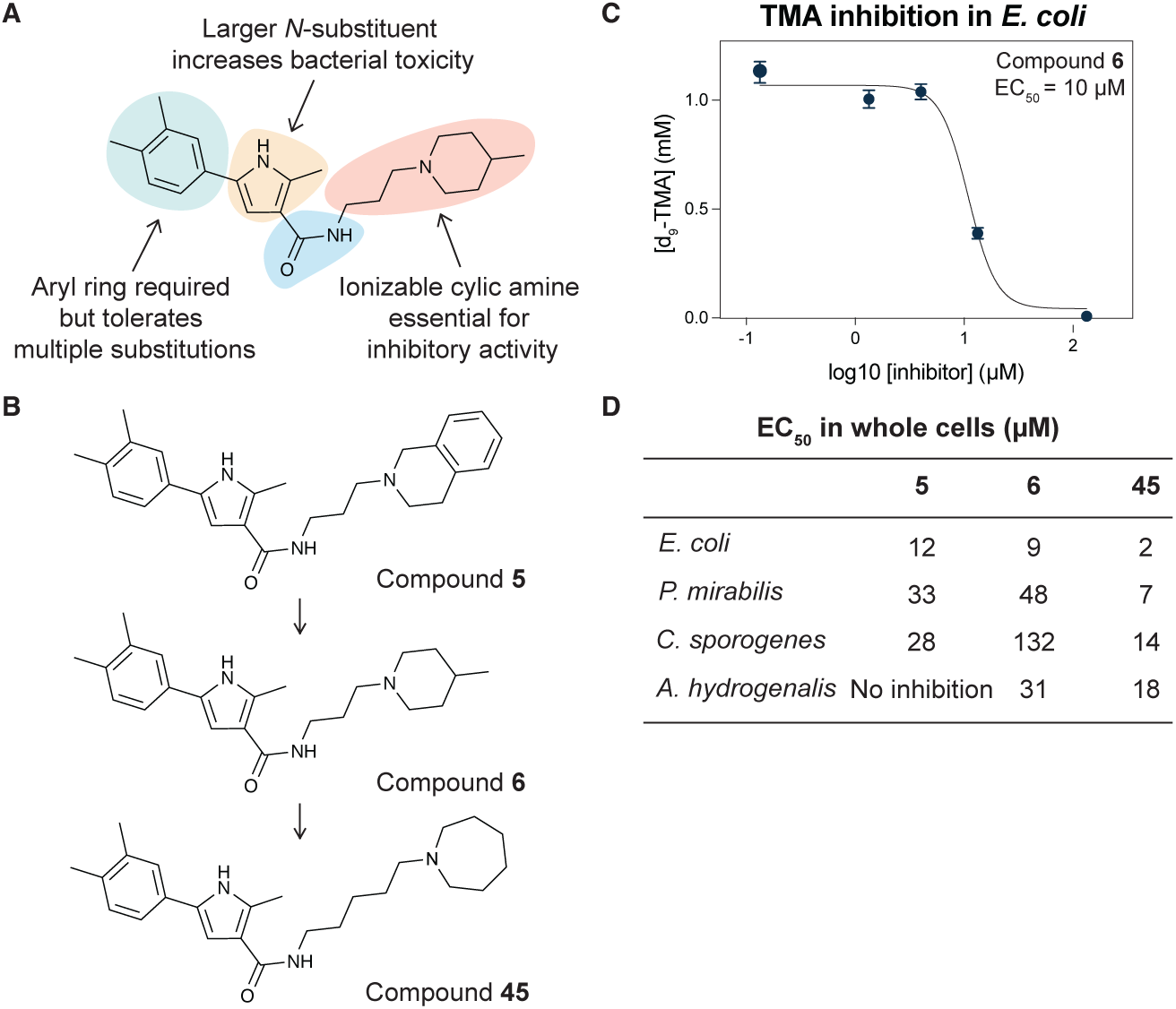
Iterative design, synthesis, and testing of analogs resulted in improved inhibitor **45**. (A) Summary of structure–activity relationships around each component of compound **6**. (B) Chemical structures of initial hit compound **5**, subsequent analog **6**, and most potent lead compound **45**. (C) TMA inhibitory activity (EC_50_ in µM) of compound **6** against *E. coli* MS 200-1. (D) TMA inhibitory activity (EC_50_ in µM) of compounds **5**, **6**, and **45** against the panel of four choline-metabolizing gut bacteria.

### Medicinal chemistry effort improves inhibitor potency and broad-spectrum activity

We next sought to improve the potency of inhibitor **6** in reducing TMA generation from choline across various choline-metabolizing gut bacteria, while minimizing its effects on the growth of other gut bacteria. Thus, we generated and evaluated analogs of **6** for their TMA inhibitory activity and gut bacterial growth inhibition. To guide these efforts, we tested analogs’ ability to inhibit the generation of TMA from choline by the four choline-metabolizing gut bacteria described above. We prioritized compounds that maintained activity across at least one gram-positive and one gram-negative strain for further testing. We next tested the effects of prioritized compounds on the anaerobic growth of nine gut commensal bacteria in complex media, including the four choline-metabolizers and five non-choline-metabolizing organisms (*Bacteroides caccae* ATCC 43185, *Bacteroides ovatus* ATCC 8483, *Bacteroides thetaiotaomicron* VPI-5482, *Collinsella aerofaciens* ATCC 25986 and *Eubacterium rectale* ATCC 33656). These additional species are prevalent human gut bacteria and have been used in prior gnotobiotic mouse studies of choline metabolism^61^. We prioritized compounds that had minimum inhibitory concentrations (MICs) of ≥200 µM across all strains, i.e., 10-fold higher than their EC_50_s for inhibition of TMA generation by choline-metabolizing bacteria.

We first investigated the importance of the heterocyclic core of compound **6** (see Table S1). Removing the C2 methyl group on the pyrrole (**7**) reduced potency by about 2-fold across all four choline-metabolizing bacteria, while moving the methyl group to the C4 position (**8**) increased the EC_50_ in *E. coli* by more than 10-fold. Changing the C2 methyl substituent to an ethyl group (**9**) improved activity slightly in *C. sporogenes* but reduced activity in *E. coli*. We then changed the relative positions of the aryl and carboxamide on the pyrrole. Keeping the aryl substituent at C5 and moving the carboxamide from C3 (**7**) to C2 (**10**) largely decreased activity in *E. coli* and *C. sporogenes* while slightly improving activity in *P. mirabilis*. Keeping the carboxamide on C3 and moving the aryl substituent from C5 (**7**) to C4 (**11**) similarly decreased activity in *C. sporogenes* while improving activity in *P. mirabilis*. Replacing the pyrrole core (**6**) with a furan (**12**) decreased activity across bacteria except for *C. sporogenes*. Comparatively, replacing the pyrrole core (**7**) with thiophene (**13**) maintained activity in *E. coli* and *P. mirabilis*, and improved in activity in *C. sporogenes*, while a thiazole (**14**) scaffold decreased activity across all bacteria tested.

Keeping the pyrrole scaffold, we then introduced larger substituents on the pyrrole nitrogen (see Table S2). Methylating the pyrrole nitrogen (**15**) improved activity in *C. sporogenes* compared to the non-methylated counterpart (**7**) but reduced activity across the rest of the bacteria panel. Analogs with larger *N*–substituents (**16**–**19**) were found to be at least 10-fold more active against *C. sporogenes* but were also less active towards *E. coli*. We next explored modifying the aryl group at the C5 position (see Table S3). We found this substituent was critical, as removal of this group (**20**) greatly reduced activity across all bacteria. Replacement of the aryl group with cyclohexane (**21**) improved potency in *C. sporogenes* but reduced potency in *E. coli*. Removal of the 3,4-dimethyl substituents (**22**) or the 4-methyl substituent (**23**) on the aryl group decreased activity against *P. mirabilis* while generally retaining activity across other bacteria, while removal of the 3-methyl substituent (**24**) greatly reduced activity in *E. coli*. Moving the 4- methyl substituent to the 5-position (**25**) retained activity across all bacteria. However, replacing the aryl group with a 2-methylpyridin-4-yl substituent (**26**) abolished activity. We then examined the necessity of the carboxamide linkage at the C3 position and found that the methylated amide (**27**) and ester (**28**) derivatives were less active in *E. coli* than the original hit (see Table S4).

Lastly, we studied the importance of the cyclic amine linked to the carboxamide (Table 1). Varying the linker length between 2 and 5 carbons (**29**–**31**) showed that the longest linker length improved potency in *C. sporogenes* and retained activity in *E. coli* and *P. mirabilis*, and *A. hydrogenalis*, while shortening the linker was detrimental in all organisms except *E. coli*. Compared to compound **29**, replacement of the cyclic amine with a cyclohexene (**32**) or ether (**33**) abolished activity. When the cyclic amine in **29** is replaced with a primary (**34**), secondary (**35**), or acyclic tertiary amine (**36**), activity across bacteria is greatly reduced. Replacing the 4- methylpiperidine of **6** with a morpholine (**37**) or 4-methyl-3-oxopiperazine (**39**) largely decreased activity while the 4-methylpiperzaine (**38**) analog lost activity in *A. hydrogenalis*. Varying the ring size of the cyclic amine of **6** (**40**–**43**) identified eight-membered ring analog **43** with generally improved activity across bacteria. Combining an increase in the ring size from piperidine to azepane with an increased linker length (**44**, **45**) improved potencies across bacteria and led to the most promising derivative (**45**) which is 2–10-fold more potent across the panel of bacteria than **6**.

**Table 1.** EC_50_ values of compounds 6, 29–45 with varying amine substituents.

| EC<sub>50</sub> in whole cells (μM) |  |  |  |  |
| --- | --- | --- | --- | --- |
| Compound | *E. coli* | *P. mirabilis* | *C. sporogenes* | *A. hydrogenalis* |
| **6** | 9 | 48 | 132 | 31 |
| **29** | 7 | 64 | 213 | 176 |
| **30** | 5 | 35 | 138 | 27 |
| **31** | 11 | 66 | 84 | 33 |
| **32** | >100 | N/A | No inhibition at 100 | N/A |
| **33** | No inhibition at 100 | N/A | No inhibition at 100 | N/A |
| **34** | 140 | No inhibition at 312 | No inhibition at 312 | N/A |
| **35** | 60 | >312 | >312 | N/A |
| **36** | 200 | >312 | >312 | N/A |
| **37** | >125 | No inhibition at 312 | No inhibition at 312 | N/A |
| **38** | 14 | 58 | 92 | No inhibition at 312 |
| **39** | >125 | No inhibition at 312 | >312 | N/A |
| **40** | 11 | 104 | 78 | 83 |
| **41** | 10 | 193 | 267 | 244 |
| **42** | 2 | 40 | 104 | 43 |
| **43** | 3 | 28 | 40 | 28 |
| **44** | 8 | 33 | 68 | 7 |
| **45** | 2 | 7 | 14 | 18 |

\*N/A indicates that compound was not tested against bacterial strain

To ensure that the TMA inhibition observed was not due to antibacterial activity, we tested select compounds for their growth inhibition on commensal bacteria (see Table S5). While most analogs did not display antibacterial activity up to 200 µM, an analog with a benzyl substituent on the pyrrole nitrogen (**19**) inhibited growth (MIC < 200 µM) in 5 out of 9 bacterial strains tested. While the potent TMA inhibitory activity of this compound against *C. sporogenes* was not likely due to antibacterial activity, this highlighted the need to validate potent compounds in the MIC assay to ensure that any reduction in choline metabolism was not due to growth inhibition and broad toxicity.

Overall, through our medicinal chemistry efforts, we found that the cyclic amine was crucial for TMA inhibitory activity. Larger *N*–substituents on the pyrrole scaffold increased bacterial toxicity while the aryl ring generally tolerated various substitutions. Interestingly, we observed that the activity of the inhibitors did not correlate with whether the target bacteria were gram-positive or negative. In general, the compounds tested were most active against *E. coli* and *A. hydrogenalis* and less active against *C. sporogenes* and *P. mirabilis*. Most compounds tested were non-toxic to commensal bacteria (MICs >200 µM across all strains) with the notable exception of analogs with larger *N*-substitutions on the pyrrole. Compared to compound **6**, the most active compound, **45**, contains an azepane instead of the 4-methylpiperidine and has a linker length of five carbons. Compound **45** has improved EC_50_ values against all bacteria tested as compared to **6**, with a marked improvement in activity in *C. sporogenes*. Our medicinal chemistry efforts highlight the importance of testing both compound efficacy as well as antibacterial activity across multiple bacteria.

### Inhibitor 45 lowers serum TMAO in gnotobiotic and conventional mice

We then evaluated a promising inhibitor in a gnotobiotic mouse model. We ranked the compounds based on their potency in *E. coli* whole cells and lack of toxicity against the gut commensal panel. We chose to test inhibitor **45** as it was one of the most potent compounds across the choline-metabolizing bacteria (EC_50_ values between 2 and 18 µM) while remaining non-toxic to other commensals (MICs >200 µM). In our experiment, we used germ-free C57BL/6 mice colonized with a previously described defined microbial community consisting of five non-choline metabolizing strains from diverse phylogenetic groups and *E. coli* MS 200-1 as the sole choline metabolizing strain. This gnotobiotic model has been used to investigate the impact of gut microbial choline metabolism on host physiology^46^. Since this *E. coli* strain is susceptible to our inhibitor *in vitro* and is the only choline-metabolizing strain in this community, any reduction in TMA generation *in vivo* can be attributed to the activity of inhibitor **45** *in vivo*.

Mice were kept on a 1% choline diet and given inhibitor **45** at a dose of either 3 mg/kg or 30 mg/kg daily through oral gavage over a period of 3 days. Serum samples were collected 24 h after the last inhibitor dose and serum concentrations of TMAO were measured (see Figure S1A). We observed reduced serum TMAO levels in the animals administered the higher inhibitor dose (Figure 4A). This experiment was also replicated in conventional C57BL/6 mice with similar results (Figure 4B). Fecal samples from the gnotobiotic mice were also collected prior to and after inhibitor treatment and the community profiled as previously described^46^. This analysis showed that inhibitor treatment did not significantly alter community composition (Figure 4C, S1B). In both gnotobiotic and conventional mice, inhibitor treatment at the higher concentration lowered TMAO levels to about 60% of vehicle treatment, with p-values of 0.054 and 0.027 respectively (Figure 4A, 4B). While the reduction in serum TMAO levels was moderate, these results suggest the inhibitor is active *in vivo*.

**Figure 4.**
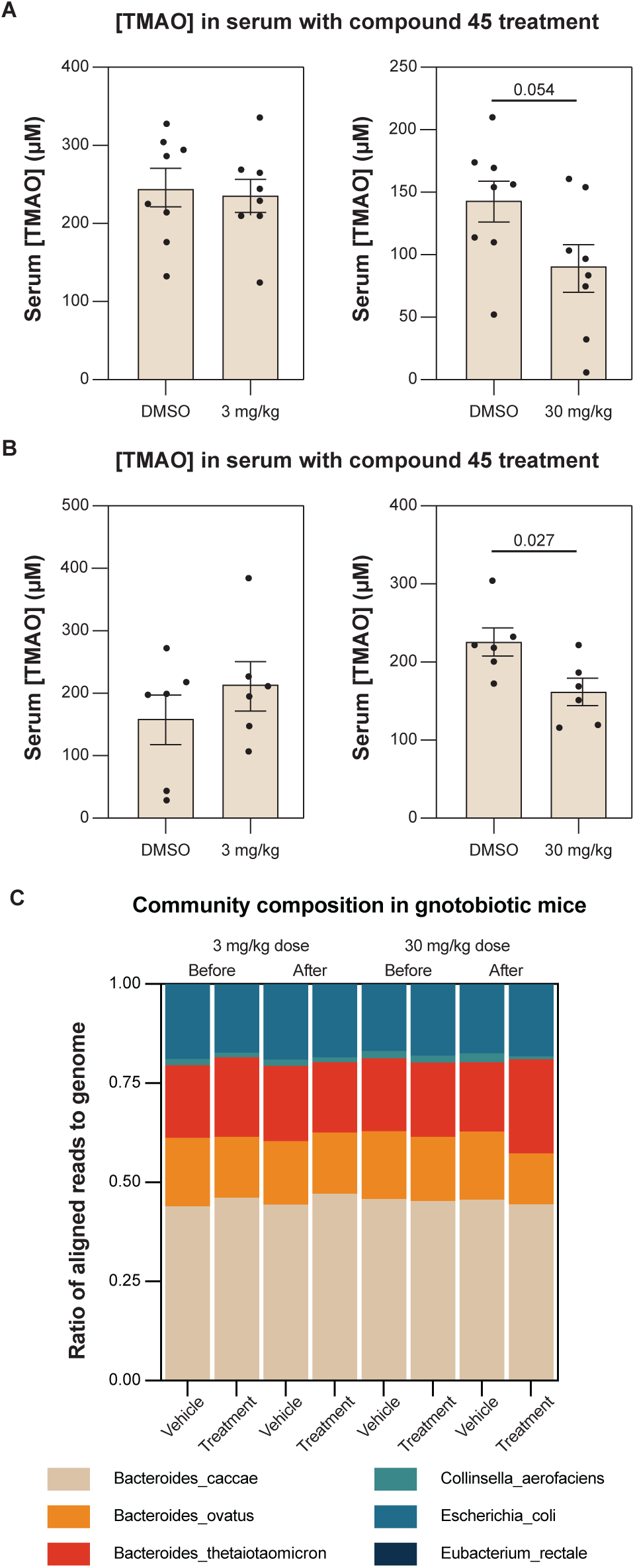
Treatment with inhibitor **45** decreased serum TMAO in gnotobiotic and conventional mice. (A) Serum TMAO levels in gnotobiotic mice after inhibitor **45** treatment. TMAO levels decreased by ∼40% when dosed at 30 mg/kg. Statistical significance calculated by unpaired t-test, n = 8, p = 0.054. (B) Serum TMAO levels in conventional mice after inhibitor **45** treatment. TMAO levels decreased by ∼40% when dosed at 30 mg/kg. Statistical significance calculated by unpaired t-test, n = 6, p = 0.027. (C) Abundance ratio of bacterial species as determined by COPRO-seq shows no significant differences in community composition before and after inhibitor **45** treatment at both low and high doses (See also Figure S1B).

## Discussion

We have designed a phenotypic HTS strategy to discover modulators of anaerobic gut microbial choline metabolism. This screen allowed for the rapid testing of a large chemical library and the identification of a novel scaffold that is less structurally similar to choline compared to existing inhibitors. The importance of the cyclic amine for activity suggests that it could potentially be involved in target engagement by mimicking choline or aid the compound in entering cells. These inhibitors also did not display antibacterial activity at 200 µM, a concentration 100-fold higher than the EC_50_ of 2 µM required to half-maximally inhibit TMA generation in cultures. This may minimize potential unwanted shifts in gut microbiome composition due to inhibitor treatment. When tested in the gnotobiotic mouse model, inhibitor **45** did not significantly change the composition of the defined bacterial community, suggesting that inhibitor treatment at this concentration may not apply substantial selective pressure on the gut microbiome. In contrast, when choline metabolism is eliminated in this defined microbial community by replacing the wild-type *E. coli* strain with the Δ*cutC* strain, the abundance ratio of *C. aerofaciens* increases at the expense of *E. coli*^46^. In conventional mice, inhibition of choline metabolism by IMC treatment induced a significant shift in the gut microbial community, with a notable increase in proportion of *Akkermansia*^13^. Further studies on how inhibitor treatment may impact a more complex gut microbial community over time will be helpful in illuminating possible secondary effects beyond direct inhibition of TMA generation.

We also observed that while inhibitor **45** reduced the conversion of choline to TMA by bacteria efficiently *in vitro* (>80%), it had moderate efficacy *in vivo*, decreasing TMAO levels by approximately 40% compared to vehicle treatment. In contrast, the halomethylcholine inhibitors previously reported completely inhibited TMAO production *in vivo* when dosed at 100 mg/kg, a concentration 3.3-fold higher than that used in our experiments, with an EC_50_ of 3.4 mg/kg and 45 mg/kg for fluoromethylcholine (FMC) and IMC respectively^13^. Further optimization of inhibitor dosage and an understanding of its pharmacokinetics may be required to improve *in vivo* efficacy.

A key consideration in developing small molecule inhibitors to study gut microbial metabolism is the potential metabolic crosstalk between microbiome and host. Many metabolites, dietary compounds, and xenobiotics can be metabolized by both gut microbial enzymes and human enzymes. As such, designing inhibitors that are specific for the microbial enzyme is critical for the compound to be a useful tool for disentangling host effects. Phenotypic HTS may be advantageous in this respect as it may be possible to identify novel chemical scaffolds that are distinct from the endogenous enzyme substrates. Although IMC and FMC are more potent than compound **45** *in vivo*, their structural similarity to choline may make them more likely to interact with host choline receptors, as evidenced by their potential neurotoxicity and the detection of related inhibitor metabolites in mice serum^13,55^.

While there is the possibility to discover new biological targets from phenotypic screening, another challenge associated with inhibitors discovered from this approach is the need for target identification. Because the gut microbiome harbors diverse bacterial taxa with different metabolic capabilities, it may be necessary to identify molecular targets and validate their functions in multiple organisms. The targets of these inhibitors remain unknown, and further characterization of their cellular targets will be required to better understand their mechanisms of action and potential off-target effects.

Although target-agnostic phenotypic screening is commonly used in antibiotic drug discovery, it has not yet been widely employed to discover non-lethal inhibitors of gut microbial metabolism. Major challenges associated with this screening approach include choosing an appropriate metabolic activity of interest and detection method. Methods that measure changes in levels of a specific metabolite of interest are often more resource and time intensive than measuring bacterial growth. However, it is not always straightforward to link a specific metabolic transformation of interest to growth. As such, previously used phenotypic screens focused gut bacterial metabolism have tended to rely on MS-based detection methods^56^. One notable exception is a recent report that screened extracts containing dietary phytochemicals for inhibition of the aerobic microbial transformation of L-carnitine to TMA by measuring growth of *Klebsiella pneumoniae* on media containing either L-carnitine or glucose as the sole carbon source and measuring resazurin reduction as a readout for metabolism^59^. This proof-of-concept study screened 39 extracts and found 2 that could reduce TMA generation from L-carnitine *in vitro* by 84.6% and 61.3%. However, a major limitation of measuring resazurin reduction as a proxy for metabolic activity rather than optical density for growth is that anaerobes likely do not reduce resazurin^62^. As such, optical density may be a more appropriate measurement when studying strictly anaerobic bacteria or metabolic processes. Our work similarly leveraged the knowledge that anaerobic choline metabolism is essential for growth only under certain conditions. A better understanding of how various bacterial enzymatic activities may be relevant to primary metabolism could allow for accelerated discovery of inhibitors through growth-based HTS.

The discovery of new inhibitors of anaerobic microbial choline metabolism in this work will inform future efforts to characterize biological targets involved in choline metabolism and to investigate how specifically modulating choline metabolism impacts host physiology. These inhibitors or additional analogs may be candidate tool compounds to study the link between TMAO generated through the anaerobic choline metabolism pathway and cardiometabolic diseases or developed as potential therapeutics for TMAU, a condition for which there are limited treatment options besides antibiotics^63^. Because these inhibitors are structurally distinct from previously established inhibitors of anaerobic choline metabolism, their activity could not have been predicted or identified based on structure-based rational design. Our success in identifying novel inhibitors of anaerobic choline metabolism using this differential growth-based phenotypic HTS not only demonstrates the utility of this approach but also suggests it may be applied more broadly to identify inhibitors of other gut bacterial metabolic activities of interest.

### Limitations of the study

The molecular target of these compounds and the mechanism by which they exert their TMA- inhibitory activity is still currently unknown. It is conceivable that the inhibitors may have complex modes of action including through modulating the gut microbiome composition *in vivo*. While the compounds have been shown to delay bacterial growth on choline when it is the sole carbon and energy source and inhibit TMA generation both *in vitro* and *in vivo*, less is known about how the gut microbiome would respond to long-term inhibitor treatment, such as developing resistance.

### Significance

Gut microbial metabolic activities play important roles in human health with many gut microbiome metabolites correlating with disease. This has prompted interest in the discovery of small molecule inhibitors both to study gut bacterial metabolism *in vivo* and as potential therapeutics. Anaerobic choline metabolism by the gut microbiome generates disease-associated metabolites TMA and its downstream metabolite TMAO, which are correlated with the incidence of TMAU and other cardiometabolic diseases. Current strategies for the development of gut microbiome-targeted inhibitors primarily rely on having a known microbial enzyme target, and in the context of anaerobic choline metabolism this has largely resulted in CutC inhibitors that resemble the choline substrate. We show that a growth-based phenotypic HTS can identify inhibitors of anaerobic choline metabolism with chemical scaffolds that are structurally distinct from choline and existing inhibitors. These compounds can be optimized further and can lower serum TMAO in gnotobiotic mice without significantly perturbing the composition of the gut microbiome. This work highlights the potential of using target-agnostic phenotypic screening to rapidly discover additional inhibitors of microbial metabolic activities, which would accelerate mechanistic studies of the gut microbiome and deepen our understanding of disease biology from correlation to causation.

## Acknowledgements

This work was supported by The Blavatnik Biomedical Accelerator at Harvard University and Servier Pharmaceuticals under the award A37535 Optimizing Inhibitors of Microbial Choline Metabolism for the Treatment of Type 2 Diabetes and Non-Alcoholic Fatty Liver Disease. E.P.B. is a Howard Hughes Medical Institute Investigator. We acknowledge the support of ICCB– Longwood Screening Facility at Harvard Medical School and Paraza Pharma Inc. for providing access to the compound libraries and chemical synthesis, respectively. We also extend our appreciation to Eugeneio I. Vivas, M.S., Director of the Gnotobiotic Core at UW–Madison for providing support for the animal experiments, Greg-Barrett-Wilt, Ph.D., Director of the Mass Spectrometry Facility Biotechnology Center at UW–Madison for LC–MS/MS measurements of TMAO, and Lindsay Kalan, Ph.D., Assistant Professor at UW–Madison for access to the iSeq100. We note that this article is subject to HHMI’s Open Access to Publications policy. HHMI laboratory heads have previously granted a nonexclusive CC BY 4.0 license to the public and a sublicensable license to HHMI in their research articles. Pursuant to those licenses, the author-accepted manuscript of this article can be made freely available under a CC BY 4.0 license immediately upon publication.

## Author contributions

Conceptualization, A.Y.M.W., M.B., W.J.S-E. and E.P.B.; Investigation, A.Y.M.W., W.J.S-E., M.B., A.W., M.S-Y. and Q.Z.; Writing — Original Draft, A.Y.M.W. and E.P.B.; Writing — Review & Editing, A.Y.M.W., W.J.S-E., M.B., M.S-Y., F.E.R. and E.P.B.; Project Administration, V.N.; Supervision, V.N., F.E.R. and E.P.B.; Funding Acquisition, E.P.B.

## Declaration of interests

E.P.B., A.Y.M.W., W.J.S-E. and M.B. are inventors on an unpublished U.S. provisional patent application submitted by President and Fellows of Harvard College that covers choline–TMA inhibitors. W.J.S-E. is currently affiliated with Facultad de Ciencias Exactas y Naturales, Departamento de Biotecnología, Laboratorio de Biotecnología Microbiana, Universidad Nacional de Asunción, San Lorenzo, Paraguay, Departamento de Análisis de Microbiomas, GeneBiome E.A.S, Luque, Paraguay, and Departamento de Investigación Aplicada de Microbiomas, MicroBios S.A, Montevideo, Uruguay. M.B. is an employee of Bayer SAS, France and A.W. is an employee of Takeda Pharmaceuticals USA, Inc. All work for this manuscript was performed when the authors were at the institutions listed in the author list.

## Supplemental information

Document S1. Tables S1–S5 and Figure S1

Document S2. Synthetic procedures for Compounds **5**–**45**

Document S3. NMR spectra of Compounds **5**–**45**

Table S6. Excel file containing HTS data, related to Figure 2

## STAR methods

### KEY RESOURCES TABLE

STAR methods KEY RESOURCES TABLE
| REAGENT or RESOURCE | SOURCE | IDENTIFIER |
| --- | --- | --- |
| Bacterial and virus strains |  |  |
| *Escherichia coli* | BEI | MS 200-1 |
| *Clostridium sporogenes* | ATCC | 15579 |
| *Proteus mirabilis* | ATCC | 29906 |
| *Anaerococcus hydrogenalis* | DSMZ | DSM 7454 |
| *Bacteroides thetaiotaomicron* VPI-5482 | ATCC | 29148 |
| *Bacteroides ovatus* | ATCC | 8483 |
| *Bacteroides caccae* | ATCC | 43185 |
| *Collinsella aerofaciens* | ATCC | 25986 |
| *Eubacterium rectale* | ATCC | 33656 |
| Chemicals, peptides, and recombinant proteins |  |  |
| (trimethyl-d<sub>9</sub>)-choline chloride | Cambridge Isotope Laboratories | DLM-549-PK |
| d<sub>9</sub>-trimethylamine hydrochloride | Sigma Aldrich | 613843 |
| Deposited data |  |  |
| High-throughput screening raw data | PubChem | AID 1963822 |
| Experimental models: Organisms/strains |  |  |
| *Mus musculus* C57BL/6 | Germ-free mice raised in the gnotobiotic facility at the University of Wisconsin – Madison | N/A |
| *Mus musculus* C57BL/6 | Conventional mice raised in the gnotobiotic facility at the University of Wisconsin – Madison | N/A |
| Software and algorithms |  |  |
| MassHunter Workstation Qualitative Analysis | Agilent Technologies |  |
| GraphPad Prism 10 | GraphPad Software | graphpad.com |

## CONTACT FOR REAGENT AND RESOURCE SHARING

Further information and requests for resources and reagents should be directed to the lead contact, Emily Balskus.

## EXPERIMENTAL MODEL AND SUBJECT DETAILS

### *In vivo* Animal Studies

#### Preparation of Core Gut Microbial Community

A core community of 5 gut bacterial species: *Bacteroides caccae* ATCC 43185*, Bacteroides ovatus* ATCC 8483*, Bacteroides thetaiotaomicron* VPI-5482*, Colinsella aerofaciens* ATCC 25986, and *Eubacterium rectale* ATCC 33656, as well as *Escherichia coli* MS 200-1 obtained from the Balskus lab, were cultured in CMM medium at 37 °C in an anaerobic environment. 1 mL of culture from each of the 6 individually grown strains was added to create this mixed culture.

### Preparation of Inhibitor

In a biosafety cabinet, a 300 mg/mL stock of sterile inhibitor in filter sterilized DMSO was prepared. After this preparation, a working solution of 3 mg/mL of the inhibitor stock in sterile 1% hydroxyethyl cellulose / 1% Polysorbate 80 was prepared within 4 days of gavaging the mice and stored at room temperature. A low dose of the inhibitor was also prepared, with a working solution of 0.3 mg/mL of the inhibitor stock in sterile 1% hydroxyethyl cellulose / 1% Polysorbate 80. The vehicle was prepared in the same way with DMSO without inhibitor.

### Inhibitor Treatment and Sample Collection

All mice were handled as outlined in animal use protocols approved by the University of Wisconsin-Madison Animal Care and Use Committee. Female C57BL/6 germ-free mice (n=8 per inhibitor group) at 10–12 weeks of age in an isolator were colonized with 200 µL of mix containing the five bacterial species mentioned above ∼10^8^ bacterial cells/mL. Mice were fed a 1% choline diet (Envigo) *ad libitum*. Two weeks after colonization, mice were administrated by gavage with 3 mg/kg of the inhibitor or vehicle between 6 and 8 a.m. once a day for 3 consecutive days. After a 10-day washout period, mice were orally administered 30 mg/kg of the inhibitor once a day (same time as above) for 3 consecutive days and blood was collected 24 hours after the last gavage.

Eight weeks old conventionally raised female C57BL/6 mice (n=6 per inhibitor group) were fed a 1% choline diet (Envigo) *ad libitum* for 10–13 days. All mice were administered by gavage with 3 mg/kg or 30 mg/kg of the inhibitor once a day (between 6 and 8 a.m.) for 3 consecutive days.

A fecal sample was collected before the first gavage. At sacrifice, tissues collected were blood, cecal contents, a fecal pellet, proximal colon, distal small intestine, liver, and kidney. Mice were anesthetized with isoflurane and underwent cardiac puncture for a terminal blood draw, followed by cervical dislocation. Tissue was snap frozen in liquid nitrogen. Blood was spun down at 8,000 × *g* for 8 min at room temperature; the resulting serum layer was removed and snap frozen. All snap frozen tissues were stored at −80 °C until needed.

## METHOD DETAILS

### Bacterial Culture Conditions and Media Preparation

Bacteria were grown overnight in an anaerobic chamber (Coy Laboratories) at 37 °C under an atmosphere of either 95% N_2_ and 5% H_2_ for the HTS and TMA inhibition EC_50_ assays or 75% N_2_, 20% CO_2_ and 5% H_2_ for MIC assays. For TMA inhibition assays, *E. coli* and *P. mirabilis* were grown in NCE medium supplemented with 20 mM choline chloride, *A. hydrogenalis* was grown in MEGA medium supplemented with 3 mM choline chloride, and *C. sporogenes* was grown in BHI medium supplemented with 20 mM of choline chloride to induce expression of the *cut* gene cluster. For MIC assays, all bacteria strains were grown in MEGA medium.

NCE media was adapted based on the “no carbon E” media as previously described^14^, and contains 29 mM potassium phosphate monobasic, 29 mM potassium phosphate dibasic, 17 mM ammonium sodium phosphate dibasic, 40 mM sodium fumarate dibasic, 0.1% (wt/v) casamino acids, and either 20 mM choline chloride or 3 g/L glycerol. This mixture was supplemented with 1 mM magnesium sulfate, 1% ATCC vitamin supplement, and 1% ATCC trace mineral supplement. MEGA medium was prepared as previously described^61^ and CMM medium was prepared by supplementing MEGA medium with 0.9 g of D-maltose, 0.86 g of D-cellobiose and 0.43 g of D-fructose per L of medium.

### Growth-Based Phenotypic High-Throughput Screening

Compounds from the Institute of Chemistry and Cell Biology-Longwood Screening Facility (ICCB-L) commercial ChemDiv7 library were transferred to 384-well plates at 5 mg/mL in DMSO.

*E. coli* was grown anaerobically in LB media containing 1 mM choline chloride overnight at 37 °C and then resuspended (2 log-fold dilution) in either NCE media supplemented with 20 mM choline or 3 g/L glycerol for screening. The *E. coli* cultures were then added to the compound plates and incubated at 37 °C anaerobically. The final concentration of compounds in culture was 20 µg/mL. OD_600_ was measured at 7 h and 20 h in duplicate compound plates. For control, 60 µg/mL kanamycin was used. Z-scores of each compound were calculated by measuring the number of standard deviations of each well’s OD_600_ with respect to the plate’s mean OD_600_. Compounds with a Z-score of less than –2 on choline but more than or equal 0 on glycerol, in at least one replicate plate, were selected for further validation. Z’ was calculated using the control columns, as an internal quality control.

### Determination of EC_50_ for TMA Inhibition in Bacterial Whole Cells

Compounds were stored as 10 mM DMSO stocks at –20 °C and diluted in DMSO to prepare varying concentrations of inhibitors for testing. Bacteria were grown anaerobically overnight in media supplemented with choline as described above. Cells were pelleted and resuspended in PBS, and d_9_-choline chloride was added to a final concentration of 1 mM. The final OD_600_ of the PBS cell resuspensions are as follows: 0.4 (*E. coli*), 0.15–0.2 (*P. mirabilis*), 0.2 (*C. sporogenes*), and 0.4 (*A. hydrogenalis*). The resuspended cells were added to 96-well plates containing inhibitors (final concentration of 1% DMSO, with final inhibitor concentrations ranging between 0 and 325 µM). The plates were sealed and incubated for 75 min (or 90 min for *P. mirabilis*) at 37 °C. Following incubation, the plates were removed from the anaerobic chamber and analyzed immediately or kept at –20 °C until LC–MS/MS analysis. Each inhibitor concentration was tested in triplicate. EC_50_ values were calculated with GraphPad Prism by fitting the log(inhibitor) vs. response—Variable slope (four parameters) model for non-linear regression.

### Determination of MIC Against Bacterial Panel

Compounds were diluted in DMSO to prepare a series of concentrations ranging from 0 to 10 mM in a 384-well plate. From this compound plate, 1 µL of each compound concentration was transferred into three new 384-well plates. Overnight cultures of bacteria in MEGA media were resuspended in fresh MEGA media (1% inoculum). 49 µL of these bacterial cultures were added to each well containing compounds. The final concentration of compounds in culture ranged between 0 and 200 µM. DMSO was used as the growth control and media was used as the sterile control. Plates were incubated anaerobically at 37 °C for 24 h, and OD_600_ was measured after incubation. Percent growth of wells was calculated with the following formula: (Sample OD_600_ – Sterile control OD_600_) / (DMSO control OD_600_ – Sterile control OD_600_) x 100%. Wells with less 50% growth were determined as no growth. Starter cultures of *B. caccae*, *C. aerofaciens* and *E. rectale* were grown for 2 days rather than overnight before testing as they grow slowly.

### Quantification of d_9_-TMA and d_9_-Choline Using LC–MS/MS

Samples were kept frozen until analysis and kept cold on ice while handling due to the volatility of TMA. Bacterial cell resuspensions in PBS were diluted 100-fold in LCMS-grade acetonitrile with 0.1% formic acid to lyse cells, precipitate proteins and extract metabolites. Samples were centrifuged at 4000 rpm for 10 min at 4 °C to pellet proteins. The supernatants were then further diluted 20-fold in LCMS-grade acetonitrile with 0.1% formic acid, and the resulting solution was analyzed by LC–MS/MS.

LC–MS/MS analysis was performed on an Agilent 1290/6470 Triple Quadrupole LC–MS instrument (Agilent Technologies) using electrospray ionization. The mass spectrometer was operated in multiple reaction monitoring (MRM) mode with positive ionization monitoring. The precursor–product ion pairs used in MRM mode were: m/z 69.1m/z → 49.1 (d_9_-TMA) and m/z 113.2 m/z → 69.1 (d_9_-choline). The capillary voltage was set to 4.0 kV and the fragmentor voltage to 30 V (d_9_-TMA) and 115 V (d_9_-choline). The collision energy for the precursor–product ion pairs was set to 29 V (d_9_-TMA) and 21 V (d_9_-choline). MS^1^ resolution was set to wide and MS^2^ resolution to unit. The drying gas temperature was maintained at 250 °C, with a flow rate of 11 L/min and a nebulizer pressure of 45 psi. The time filter width used was 0.07 min. The injection volume of all samples was 1 μL. Samples were injected onto InfinityLab Poroshell 120 (Agilent) HILIC columns (2.7 μm, 100 mm × 2.1 mm, 100 Å) preceded by InfinityLab Poroshell 120 HILIC guard columns (2.1 mm, 1.9µm UHPLC guard). The LC conditions on the binary pump were: a gradient of 5–35% A increasing over 1.95 min (1.2 mL/min), a gradient of 35–5% A decreasing over 0.05 min (1.6 mL/min) and held at 5% A for 0.5 min (2 mL/min). The columns were regenerated on the quaternary pump with the following conditions: 60% A for 1.25 min (1.2 mL/min), a decreasing gradient of 60–5% A over 0.05 min (1.2 mL/min) and held at 5% A for 0.5 min (1.5 mL/min). Solvent A = 10 mM ammonium formate with 0.1% formic acid; solvent B = acetonitrile with 0.1% formic acid. Data analysis was performed with Mass Hunter Workstation Data Acquisition and Qualitative Analysis software (Agilent Technologies).

### Quantification of TMAO in Mouse Serum Samples Using LC–MS/MS

One volume of plasma and 4 volumes of ice-cold extraction solution (HPLC-grade methanol with 2.5 µM d_9_-TMAO internal standard) were combined and spun at 21,100 x *g* for 3 min at 4 °C. One volume of the resulting supernatant was then mixed with 1 volume of HPLC-grade water. The resulting samples were injected onto a Waters ACQUITY C18 UPLC column (1.7 µm, 2.1 mm x 100 mm) that was coupled to a Thermo Fisher Q-Exactive mass spectrometer at a flow rate of 0.2 mL/min. Elution of samples occurred over a 7 min isocratic gradient of 25% water, 5 mM ammonium acetate, 0.05% acetic acid and 75% methanol. TMAO was quantified in the positive mode using Parallel Reaction Monitoring (PRM) using an inclusion list of 76.076 and 85.132 m/z (for TMAO and d_9_-TMAO respectively). El-MAVEN was used for peak quantification, and internal standards were utilized as a comparison to calculate plasma concentrations. Statistical significance was calculated using unpaired t-test.

### Sample Preparation and Sequencing of Mice Samples

DNA was extracted from fecal samples according to published bead-beating procedures^64,65^ and pellets were dried with ethanol and resuspended in 10 mM Tris-HCl, pH 8.5. NucleoSpin Gel and PCR Clean-up Kit (Macherey-Nagel) was used to remove contaminants. Isolated DNA was stored at −80°C until downstream processing. Libraries were prepared according to the manufacturer’s protocol (15031942 v05) for Illumina Nextera XT DNA Library Preparation Kit (Illumina). Briefly, 1.0 ng of each input DNA was enzymatically fragmented and tagged by fragmentation. The cleaved DNA was subjected to limited-cycle PCR for indexing with i5 and i7 index adaptors. The PCR products were purified using Agencourt AMPure XP beads (Beckman Coulter) and normalized using the Nextera XT Library Normalization Beads (Illumina). The normalized libraries were diluted and pooled at final loading concentration of 150 pM according to the manufacturer’s protocol (1000000036024 v07). The pooled library was spiked with 2% non-denatured TailorMix Dual Indexed PhiX Control Library (SeqMatic) and sequenced with Illumina iSeq 100 (v3.0.0.359, 2×150 cycles, paired-end). Quality control monitoring of the library preparation process was performed using agarose gel electrophoresis. Results were processed using the software pipeline detailed by McNulty et al.^66^.

## QUANTIFICATION AND STATISTICAL ANALYSIS

All data fitting and statistical analysis were performed with GraphPad Prism version 10, GraphPad Software, La Jolla, California, USA, www.graphpad.com. Statistical values and statistical significance were also reported in the related figure or figure legends.

## DATA AND SOFTWARE AVAILABILITY

### Data Resources

The HTS screening data reported in the paper is deposited in PubChem under the PubChem AID 1963822.

### Software

All software used in this study was listed in the Key Resources Table and describes in the detailed methods where applicable.

## References

1. Durack, J., and Lynch, S.V. (2019). The gut microbiome: Relationships with disease and opportunities for therapy. Journal of Experimental Medicine 216, 20–40. 10.1084/jem.20180448.

2. Fan, Y., and Pedersen, O. (2021). Gut microbiota in human metabolic health and disease. Nat Rev Microbiol 19, 55–71. 10.1038/s41579-020-0433-9.

3. Jiménez-Avalos, J.A., Arrevillaga-Boni, G., González-López, L., García-Carvajal, Z.Y., and González-Avila, M. (2021). Classical methods and perspectives for manipulating the human gut microbial ecosystem. Critical Reviews in Food Science and Nutrition 61, 234–258. 10.1080/10408398.2020.1724075.

4. Woo, A.Y.M., Aguilar Ramos, M.A., Narayan, R., Richards-Corke, K.C., Wang, M.L., Sandoval-Espinola, W.J., and Balskus, E.P. (2023). Targeting the human gut microbiome with small-molecule inhibitors. Nat Rev Chem 7, 319–339. 10.1038/s41570-023-00471-4.

5. Visconti, A., Le Roy, C.I., Rosa, F., Rossi, N., Martin, T.C., Mohney, R.P., Li, W., De Rinaldis, E., Bell, J.T., Venter, J.C., et al. (2019). Interplay between the human gut microbiome and host metabolism. Nat Commun 10, 4505. 10.1038/s41467-019-12476-z.

6. Chaudhari, S.N., McCurry, M.D., and Devlin, A.S. (2021). Chains of evidence from correlations to causal molecules in microbiome-linked diseases. Nature Chemical Biology 17, 1046–1056. 10.1038/s41589-021-00861-z.

7. Wallace, B.D., Wang, H., Lane, K.T., Scott, J.E., Orans, J., Koo, J.S., Venkatesh, M., Jobin, C., Yeh, L., Mani, S., et al. (2010). Alleviating Cancer Drug Toxicity by Inhibiting a Bacterial Enzyme. Science 330, 831–835. 10.1126/science.1191175.

8. Wallace, B.D., Roberts, A.B., Pollet, R.M., Ingle, J.D., Biernat, K.A., Pellock, S.J., Venkatesh, M.K., Guthrie, L., O’Neal, S.K., Robinson, S.J., et al. (2015). Structure and Inhibition of Microbiome β-Glucuronidases Essential to the Alleviation of Cancer Drug Toxicity. Chemistry and Biology 22, 1238–1249. 10.1016/j.chembiol.2015.08.005.

9. Pellock, S.J., Creekmore, B.C., Walton, W.G., Mehta, N., Biernat, K.A., Cesmat, A.P., Ariyarathna, Y., Dunn, Z.D., Li, B., Jin, J., et al. (2018). Gut Microbial β-Glucuronidase Inhibition via Catalytic Cycle Interception. ACS Cent. Sci. 4, 868–879. 10.1021/acscentsci.8b00239.

10. Ervin, S.M., Hanley, R.P., Lim, L., Walton, W.G., Pearce, K.H., Bhatt, A.P., James, L.I., and Redinbo, M.R. (2019). Targeting Regorafenib-Induced Toxicity through Inhibition of Gut Microbial β - Glucuronidases. ACS Chemical Biology 14. 10.1021/acschembio.9b00663.

11. Letertre, M.P.M., Bhatt, A.P., Harvey, M., Nicholson, J.K., Wilson, I.D., Redinbo, M.R., and Swann, J.R. (2022). Characterizing the metabolic effects of the selective inhibition of gut microbial β-glucuronidases in mice. Sci Rep 12, 17435. 10.1038/s41598-022-21518-4.

12. Wang, Z., Roberts, A.B., Buffa, J.A., Levison, B.S., Zhu, W., Org, E., Gu, X., Huang, Y., Zamanian-Daryoush, M., Culley, M.K., et al. (2015). Non-lethal Inhibition of Gut Microbial Trimethylamine Production for the Treatment of Atherosclerosis. Cell 163, 1585–1595. 10.1016/j.cell.2015.11.055.

13. Roberts, A.B., Gu, X., Buffa, J.A., Hurd, A.G., Wang, Z., Zhu, W., Gupta, N., Skye, S.M., Cody, D.B., Levison, B.S., et al. (2018). Development of a gut microbe–targeted nonlethal therapeutic to inhibit thrombosis potential. Nat Med 24, 1407–1417. 10.1038/s41591-018-0128-1.

14. Orman, M., Bodea, S., Funk, M.A., Campo, A.M., Drennan, C.L., Balskus, E.P., Martínez-Del Campo, A., Bollenbach, M., Drennan, C.L., and Balskus, E.P. (2018). Structure-Guided Identification of a Small Molecule That Inhibits Anaerobic Choline Metabolism by Human Gut Bacteria. Journal of the American Chemical Society 17, 34. 10.1021/jacs.8b04883.

15. Bollenbach, M., Ortega, M., Orman, M., Drennan, C.L., and Balskus, E.P. (2020). Discovery of a Cyclic Choline Analog That Inhibits Anaerobic Choline Metabolism by Human Gut Bacteria. ACS Medicinal Chemistry Letters, 0–5. 10.1021/acsmedchemlett.0c00005.

16. Williams, B.B., Van Benschoten, A.H., Cimermancic, P., Donia, M.S., Zimmermann, M., Taketani, M., Ishihara, A., Kashyap, P.C., Fraser, J.S., and Fischbach, M.A. (2014). Discovery and Characterization of Gut Microbiota Decarboxylases that Can Produce the Neurotransmitter Tryptamine. Cell Host & Microbe 16, 495–503. 10.1016/j.chom.2014.09.001.

17. Rekdal, V.M., Bess, E.N., Bisanz, J.E., Turnbaugh, P.J., and Balskus, E.P. (2019). Discovery and inhibition of an interspecies gut bacterial pathway for Levodopa metabolism. Science 364. 10.1126/science.aau6323.

18. Adhikari, A.A., Seegar, T.C.M., Ficarro, S.B., McCurry, M.D., Ramachandran, D., Yao, L., Chaudhari, S.N., Ndousse-Fetter, S., Banks, A.S., Marto, J.A., et al. (2020). Development of a covalent inhibitor of gut bacterial bile salt hydrolases. Nature Chemical Biology 16, 318–326. 10.1038/s41589-020-0467-3.

19. Adhikari, A.A., Ramachandran, D., Chaudhari, S.N., Powell, C.E., Banks, A.S., and Sloan Devlin, A. A gut-restricted lithocholic acid analog as an 1 inhibitor of gut bacterial bile salt hydrolases. ACS Chemical Biology. 10.1101/2021.03.15.435552.

20. Graboski, A.L., Kowalewski, M.E., Simpson, J.B., Cao, X., Ha, M., Zhang, J., Walton, W.G., Flaherty, D.P., and Redinbo, M.R. (2023). Mechanism-based inhibition of gut microbial tryptophanases reduces serum indoxyl sulfate. Cell Chemical Biology, S2451945623002441. 10.1016/j.chembiol.2023.07.015.

21. Zahir, T., Camacho, R., Vitale, R., Ruckebusch, C., Hofkens, J., Fauvart, M., and Michiels, J. (2019). High-throughput time-resolved morphology screening in bacteria reveals phenotypic responses to antibiotics. Commun Biol 2, 269. 10.1038/s42003-019-0480-9.

22. Kim, S.M., Park, J., Kim, M.S., Song, H., Jo, A., Park, H., Kim, T.S., Choi, S.H., and Park, S.B. (2020). Phenotypic Discovery of an Antivirulence Agent against Vibrio vulnificus via Modulation of Quorum-Sensing Regulator SmcR. ACS Infect. Dis. 6, 3076–3082. 10.1021/acsinfecdis.0c00587.

23. Garcia, C., Burgain, A., Chaillot, J., Pic, É., Khemiri, I., and Sellam, A. (2018). A phenotypic small-molecule screen identifies halogenated salicylanilides as inhibitors of fungal morphogenesis, biofilm formation and host cell invasion. Sci Rep 8, 11559. 10.1038/s41598-018-29973-8.

24. Kirienko, D.R., Revtovich, A.V., and Kirienko, N.V. (2016). A High-Content, Phenotypic Screen Identifies Fluorouridine as an Inhibitor of Pyoverdine Biosynthesis and Pseudomonas aeruginosa Virulence. mSphere 1, 10.1128/msphere.00217-16. 10.1128/msphere.00217-16.

25. Hanna, N., Kicka, S., Chiriano, G., Harrison, C., Sakouhi, H.O., Trofimov, V., Kranjc, A., Nitschke, J., Pagni, M., Cosson, P., et al. (2020). Identification of Anti-Mycobacterium and Anti-Legionella Compounds With Potential Distinctive Structural Scaffolds From an HD-PBL Using Phenotypic Screens in Amoebae Host Models. Front. Microbiol. 11. 10.3389/fmicb.2020.00266.

26. Clatworthy, A.E., Romano, K.P., and Hung, D.T. (2018). Whole-organism phenotypic screening for anti-infectives promoting host health. Nat Chem Biol 14, 331–341. 10.1038/s41589-018-0018-3.

27. Sykes, M.L., and Avery, V.M. (2013). Approaches to Protozoan Drug Discovery: Phenotypic Screening. J. Med. Chem. 56, 7727–7740. 10.1021/jm4004279.

28. Vincent, F., Nueda, A., Lee, J., Schenone, M., Prunotto, M., and Mercola, M. (2022). Phenotypic drug discovery: recent successes, lessons learned and new directions. Nat Rev Drug Discov 21, 899–914. 10.1038/s41573-022-00472-w.

29. Nakae, T. (1986). Outer-Membrane Permeability of Bacteria. CRC Critical Reviews in Microbiology 13, 1–62. 10.3109/10408418609108734.

30. Chen, P.B., Black, A.S., Sobel, A.L., Zhao, Y., Mukherjee, P., Molparia, B., Moore, N.E., Aleman Muench, G.R., Wu, J., Chen, W., et al. (2020). Directed remodeling of the mouse gut microbiome inhibits the development of atherosclerosis. Nature Biotechnology. 10.1038/s41587-020-0549-5.

31. Lukonin, I., Zinner, M., and Liberali, P. (2021). Organoids in image-based phenotypic chemical screens. Exp Mol Med 53, 1495–1502. 10.1038/s12276-021-00641-8.

32. Zeisel, S.H., Wishnok, J.S., and Blusztajn, J.K. (1983). Formation of Methylamines from Ingested Choline and Lecithin. The Journal of Pharmacology and Experimental Therapeutics 225, 320–324.

33. De La Huerga, J., and Popper, H. (1951). Urinary Excretion of Choline Metabolites Following Choline Administration in Normals and Patients With Hepatobiliary Dieseases. Journal of Clinical Investigation 30, 463–470.

34. Al-Waiz, M., Mikov, M., Mitchell, S.C., and Smith, R.L. (1992). The Exogenous Origin of Trimethylamine in the Mouse Animal Studies. Metabolism 41, 135–136.

35. Yu, Z., Zhang, L., Jiang, X., Xue, C., Chi, N., Zhang, T., and Wang, Y. (2020). Effects of dietary choline, betaine, and L-carnitine on the generation of trimethylamine-N-oxide in healthy mice. Journal of Food Science 85, 2207–2215. 10.1111/1750-3841.15186.

36. Bennett, B.J., Vallim, T.Q.D.A., Wang, Z., Shih, D.M., Meng, Y., Gregory, J., Allayee, H., Lee, R., Graham, M., Crooke, R., et al. (2013). Trimethylamine-N-Oxide, a metabolite associated with atherosclerosis, exhibits complex genetic and dietary regulation. Cell Metabolism 17, 49–60. 10.1016/j.cmet.2012.12.011.

37. Brewster, M.A., and Schedewie, H. (1983). Trimethylaminuria. Annals of Clinical and Laboratory Science 13, 20–24.

38. Wang, Z., Klipfell, E., Bennett, B.J., Koeth, R., Levison, B.S., Dugar, B., Feldstein, A.E., Britt, E.B., Fu, X., Chung, Y.M., et al. (2011). Gut flora metabolism of phosphatidylcholine promotes cardiovascular disease. Nature 472, 57–65. 10.1038/nature09922.

39. Roncal, C., Martínez-Aguilar, E., Orbe, J., Ravassa, S., Fernandez-Montero, A., Saenz-Pipaon, G., Ugarte, A., Estella-Hermoso De Mendoza, A., Rodriguez, J.A., Fernández-Alonso, S., et al. (2019). Trimethylamine-N-Oxide (TMAO) Predicts Cardiovascular Mortality in Peripheral Artery Disease. Sci Rep 9, 15580. 10.1038/s41598-019-52082-z.

40. Zeisel, S.H., and Warrier, M. (2017). Trimethylamine *N* -Oxide, the Microbiome, and Heart and Kidney Disease. Annu. Rev. Nutr. 37, 157–181. 10.1146/annurev-nutr-071816-064732.

41. Missailidis, C., Hällqvist, J., Qureshi, A.R., Barany, P., Heimbürger, O., Lindholm, B., Stenvinkel, P., and Bergman, P. (2016). Serum Trimethylamine-N-Oxide Is Strongly Related to Renal Function and Predicts Outcome in Chronic Kidney Disease. PLoS ONE 11, e0141738. 10.1371/journal.pone.0141738.

42. Xu, K.Y., Xia, G.H., Lu, J.Q., Chen, M.X., Zhen, X., Wang, S., You, C., Nie, J., Zhou, H.W., and Yin, J. (2017). Impaired renal function and dysbiosis of gut microbiota contribute to increased trimethylamine-N-oxide in chronic kidney disease patients. Scientific Reports 7. 10.1038/s41598-017-01387-y.

43. Shan, Z., Sun, T., Huang, H., Chen, S., Chen, L., Luo, C., Yang, W., Yang, X., Yao, P., Cheng, J., et al. (2017). Association between microbiota-dependent metabolite trimethylamine-N-oxide and type 2 diabetes. American Journal of Clinical Nutrition 106, 888–894. 10.3945/ajcn.117.157107.

44. Chen, Y.M., Liu, Y., Zhou, R.F., Chen, X.L., Wang, C., Tan, X.Y., Wang, L.J., Zheng, R.D., Zhang, H.W., Ling, W.H., et al. (2016). Associations of gut-flora-dependent metabolite trimethylamine-N-oxide, betaine and choline with non-alcoholic fatty liver disease in adults. Scientific Reports 6. 10.1038/srep19076.

45. Organ, C.L., Otsuka, H., Bhushan, S., Wang, Z., Bradley, J., Trivedi, R., Polhemus, D.J., Tang, W.H.W., Wu, Y., Hazen, S.L., et al. (2016). Choline Diet and Its Gut Microbe-Derived Metabolite, Trimethylamine N-Oxide, Exacerbate Pressure Overload-Induced Heart Failure. Circulation: Heart Failure 9, 1–10. 10.1161/CIRCHEARTFAILURE.115.002314.

46. Romano, K.A., Martinez-del Campo, A., Kasahara, K., Chittim, C.L., Vivas, E.I., Amador-Noguez, D., Balskus, E.P., and Rey, F.E. (2017). Metabolic, Epigenetic, and Transgenerational Effects of Gut Bacterial Choline Consumption. Cell Host and Microbe 22, 279–290.e7. 10.1016/j.chom.2017.07.021.

47. Thomas, M.S., and Fernandez, M.L. (2021). Trimethylamine N-Oxide (TMAO), Diet and Cardiovascular Disease. Curr Atheroscler Rep 23, 12. 10.1007/s11883-021-00910-x.

48. Martínez-del Campo, A., Bodea, S., Hamer, H.A., Marks, J.A., Haiser, H.J., Turnbaugh, P.J., and Balskusa, E.P. (2015). Characterization and detection of a widely distributed gene cluster that predicts anaerobic choline utilization by human gut bacteria. mBio 6, 1–12. 10.1128/mBio.00042-15.

49. Craciun, S., and Balskus, E.P. (2012). Microbial conversion of choline to trimethylamine requires a glycyl radical enzyme. Proceedings of the National Academy of Sciences 109, 21307–21312. 10.1073/pnas.1215689109.

50. Bodea, S., Funk, M.A., Balskus, E.P., and Drennan, C.L. (2016). Molecular Basis of C–N Bond Cleavage by the Glycyl Radical Enzyme Choline Trimethylamine-Lyase. Cell Chemical Biology 23, 1206–1216. 10.1016/j.chembiol.2016.07.020.

51. Craciun, S., Marks, J.A., and Balskus, E.P. (2014). Characterization of Choline Trimethylamine-Lyase Expands the Chemistry of Glycyl Radical Enzymes. ACS Chem. Biol. 9, 1408–1413. 10.1021/cb500113p.

52. Gupta, N., Buffa, J.A., Roberts, A.B., Sangwan, N., Skye, S.M., Li, L., Ho, K.J., Varga, J., DiDonato, J.A., Tang, W.H.W., et al. (2020). Targeted Inhibition of Gut Microbial TMAO Production Reduces Renal Tubulointerstitial Fibrosis and Functional Impairment in a Murine Model of Chronic Kidney Disease. Arteriosclerosis, Thrombosis, and Vascular Biology, 1239–1255. 10.1161/atvbaha.120.314139.

53. Zhang, W., Miikeda, A., Zuckerman, J., Jia, X., Charugundla, S., Zhou, Z., Kaczor-Urbanowicz, K.E., Magyar, C., Guo, F., Wang, Z., et al. (2021). Inhibition of microbiota-dependent TMAO production attenuates chronic kidney disease in mice. Sci Rep 11, 518. 10.1038/s41598-020-80063-0.

54. Nonlethal Inhibition of Gut Microbial Trimethylamine N-oxide Production Improves Cardiac Function and Remodeling in a Murine Model of Heart Failure 10.1161/JAHA.119.016223.

55. Mistry, J.S., Abraham, D.J., Kozikowski, A.P., and Hanin, I. (1991). Neurochemistry of aging. 2. Design, synthesis and biological evaluation of halomethyl analogs of choline with high affinity choline transport inhibitory activity. J. Med. Chem. *34*, 2031–2036. 10.1021/jm00111a016.

56. Winter, M., Bretschneider, T., Thamm, S., Kleiner, C., Grabowski, D., Chandler, S., Ries, R., Kley, J.T., Fowler, D., Bartlett, C., et al. (2019). Chemical Derivatization Enables MALDI-TOF-Based High-Throughput Screening for Microbial Trimethylamine (TMA)-Lyase Inhibitors. SLAS Discovery 24, 766–777. 10.1177/2472555219838216.

57. Švec, P., Nový, Z., Kučka, J., Petřík, M., Sedláček, O., Kuchař, M., Lišková, B., Medvedíková, M., Kolouchová, K., Groborz, O., et al. (2020). Iodinated Choline Transport-Targeted Tracers. J. Med. Chem. 63, 15960–15978. 10.1021/acs.jmedchem.0c01710.

58. Carlson, H.K., Mullan, M.R., Mosqueda, L.A., Chen, S., Arkin, M.R., and Coates, J.D. (2017). High-Throughput Screening To Identify Potent and Specific Inhibitors of Microbial Sulfate Reduction. Environ. Sci. Technol. 51, 7278–7285. 10.1021/acs.est.7b00686.

59. Simó, C., Fornari, T., García-Risco, M.R., Peña-Cearra, A., Abecia, L., Anguita, J., Rodríguez, H., and García-Cañas, V. (2022). Resazurin-based high-throughput screening method for the discovery of dietary phytochemicals to target microbial transformation of L - carnitine into trimethylamine, a gut metabolite associated with cardiovascular disease. Food Funct. 13, 5640–5653. 10.1039/D2FO00103A.

60. Campbell, J. (2010). High-Throughput Assessment of Bacterial Growth Inhibition by Optical Density Measurements. CP Chemical Biology 2, 195–208. 10.1002/9780470559277.ch100115.

61. Romano, K.A., Vivas, E.I., Amador-noguez, D., and Rey, F.E. (2015). Intestinal Microbiota Composition Modulates Choline Bioavailability. mBio 6, 1–8. 10.1128/mBio.02481-14.Editor.

62. Braissant, O., Astasov-Frauenhoffer, M., Waltimo, T., and Bonkat, G. (2020). A Review of Methods to Determine Viability, Vitality, and Metabolic Rates in Microbiology. Front. Microbiol. 11. 10.3389/fmicb.2020.547458.

63. Schmidt, A.C., and Leroux, J.-C.C. (2020). Treatments of trimethylaminuria: where we are and where we might be heading (Elsevier Ltd) 10.1016/j.drudis.2020.06.026.

64. Turnbaugh, P.J., Hamady, M., Yatsunenko, T., Cantarel, B.L., Duncan, A., Ley, R.E., Sogin, M.L., Jones, W.J., Roe, B.A., Affourtit, J.P., et al. (2009). A core gut microbiome in obese and lean twins. Nature 457, 480–484. 10.1038/nature07540.

65. Romano, K.A., Dill-McFarland, K.A., Kasahara, K., Kerby, R.L., Vivas, E.I., Amador-Noguez, D., Herd, P., and Rey, F.E. (2018). Fecal Aliquot Straw Technique (FAST) allows for easy and reproducible subsampling: assessing interpersonal variation in trimethylamine-N-oxide (TMAO) accumulation. Microbiome 6, 91. 10.1186/s40168-018-0458-8.

66. McNulty, N.P., Yatsunenko, T., Hsiao, A., Faith, J.J., Muegge, B.D., Goodman, A.L., Henrissat, B., Oozeer, R., Cools-Portier, S., Gobert, G., et al. (2011). The Impact of a Consortium of Fermented Milk Strains on the Gut Microbiome of Gnotobiotic Mice and Monozygotic Twins. Science Translational Medicine 3, 106ra106–106ra106. 10.1126/scitranslmed.3002701.

