## Supplementary material for "Phenotypic high-throughput screening identifies modulators of gut microbial choline metabolism": Document S1

**Supplementary Figures**

**Table S1**: $EC_{50}$ values of compounds **6–14** with varying heterocyclic cores

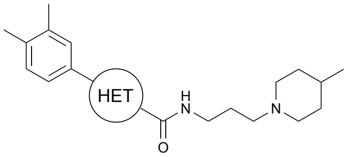


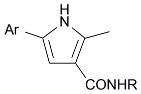

**6**

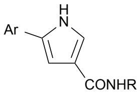

**7**

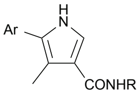

**8**

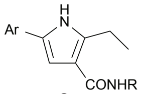

**9**

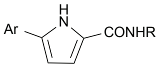

**10**

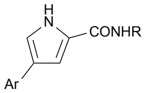

**11**

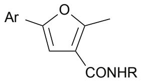

**12**

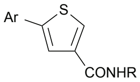

**13**

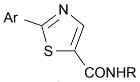

**14**

| | **$EC_{50}$ in whole cells (µM)** | | | |
|---|---|---|---|---|
| Compound | *E. coli* | *P. mirabilis* | *C. sporogenes* | *A. hydrogenalis* |
| **6** | 9 | 48 | 132 | 31 |
| **7** | 16 | 80 | 215 | 66 |
| **8** | >125 | 131 | 97 | N/A |
| **9** | 35 | N/A | 103 | N/A |
| **10** | 98 | 63 | 312 | N/A |
| **11** | 13 | 42 | >312 | N/A |
| **12** | 15 | >312 | 54 | 54 |
| **13** | 14 | 50 | 85 | N/A |
| **14** | 27 | 197 | 215 | N/A |

*N/A denotes that compound was not tested against bacterial strain.

**Table S2**: $EC_{50}$ values of compounds **6**, **15**–**19** with varying *N*-pyrrole substituents

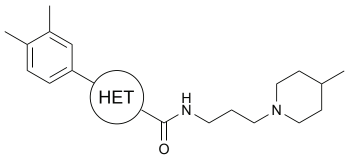


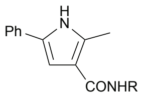


**6**

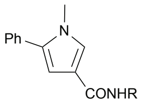


**15**

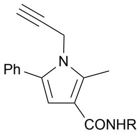


**16**

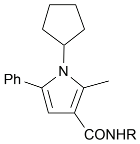


**17**

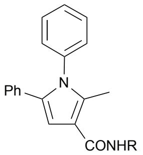


**18**

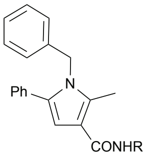


**19**

**$EC_{50}$ in whole cells (μM)**

| Compound | *E. coli* | *P. mirabilis* | *C. sporogenes* | *A. hydrogenalis* |
|---|---|---|---|---|
| **6** | 9 | 48 | 132 | 31 |
| **15** | 21 | 98 | 114 | 116 |
| **16** | 39 | N/A | 5 | N/A |
| **17** | 69 | N/A | 8 | N/A |
| **18** | 489 | N/A | 3 | N/A |
| **19** | 28 | N/A | 4 | N/A |

*N/A denotes that compound was not tested against bacterial strain.

**Table S3**: $EC_{50}$ values of compounds **6**, **20**–**26** with varying substituents on C5

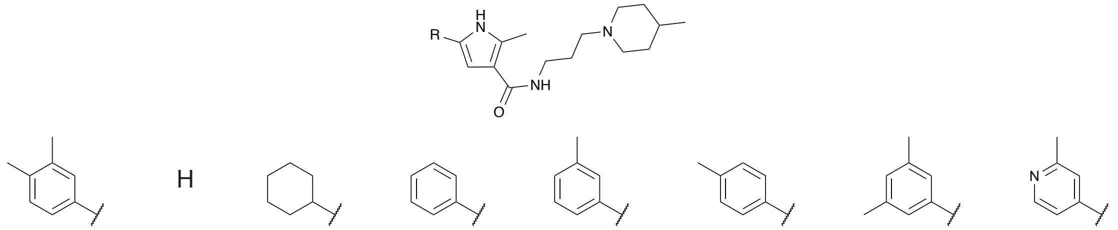


**6** **20** **21** **22** **23** **24** **25** **26**

| | **$EC_{50}$ in whole cells (μM)** | | | |
|---|---|---|---|---|
| Compound | *E. coli* | *P. mirabilis* | *C. sporogenes* | *A. hydrogenalis* |
| **6** | 9 | 48 | 132 | 31 |
| **20** | >125 | No inhibition at 312 | No inhibition at 312 | No inhibition at 312 |
| **21** | 51 | N/A | 8 | 32 |
| **22** | 7 | 157 | 147 | 24 |
| **23** | 5 | 46 | 95 | 35 |
| **24** | 92 | 131 | 117 | N/A |
| **25** | 6 | 28 | 133 | 19 |
| **26** | >125 | >312 | No inhibition at 312 | No inhibition at 312 |

**Table S4**: $EC_{50}$ values of compounds **6**, **27**, **28** with varying substituents on the C3 carboxamide

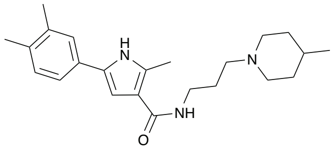

**6**

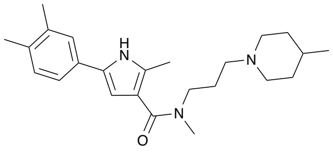

**27**

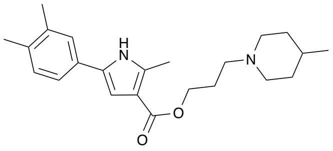

**28**

| | **$EC_{50}$ in whole cells (μM)** | | | |
|---|---|---|---|---|
| Compound | *E. coli* | *P. mirabilis* | *C. sporogenes* | *A. hydrogenalis* |
| **6** | 9 | 48 | 132 | 31 |
| **27** | >125 | >312 | 12 | 44 |
| **28** | 15 | >312 | 18 | 11 |

*N/A denotes that compound was not tested against bacterial strain.

**Table S5**: MIC values of select compounds against panel of commensal bacterial strains

| | MIC in MEGA medium (μM) | | | |
|---|---|---|---|---|
| Compound | *E. coli* | *P. mirabilis* | *C. sporogenes* | *A. hydrogenalis* |
| **6** | >200 | >200 | >200 | 200 |
| **19** | >200 | >200 | >200 | 25 |
| **23** | >200 | >200 | >200 | >200 |
| **44** | >200 | >200 | >200 | >200 |
| **45** | >200 | >200 | >200 | >200 |

| | MIC in MEGA medium (μM) | | | | |
|---|---|---|---|---|---|
| Compound | *B. thetaiotaomicron* | *B. ovatus* | *B. caccae* | *C. aerofaciens* | *E. rectale* |
| **6** | >200 | >200 | >200 | >200 | 200 |
| **19** | 50 | 25 | >200 | 6.25 | 25 |
| **23** | >200 | >200 | >200 | 200 | >200 |
| **44** | >200 | >200 | >200 | >200 | N/A |
| **45** | >200 | >200 | >200 | >200 | >200 |

*N/A denotes that compound was not tested against bacterial strain.

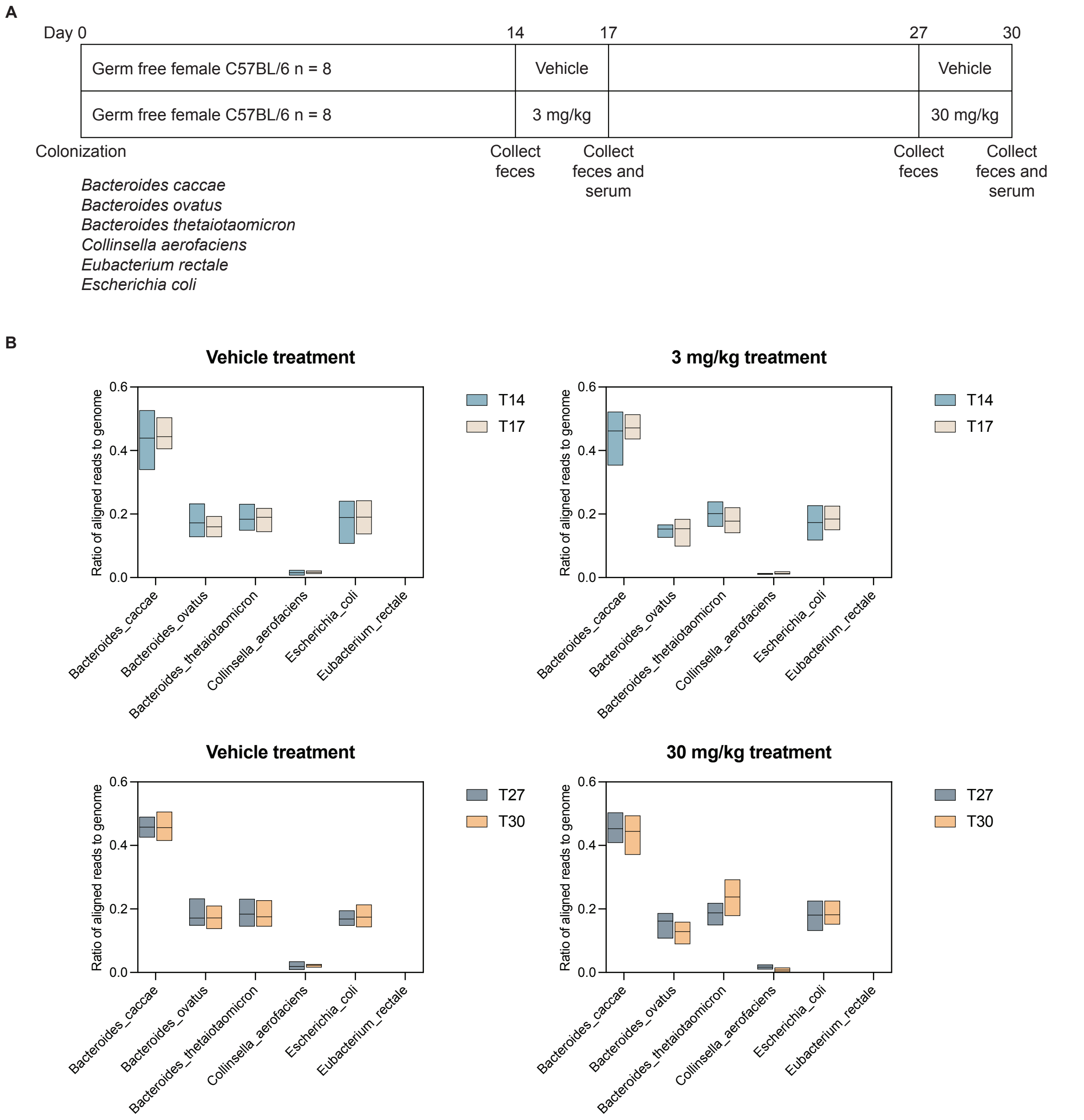


**Figure S1:** Inhibitor treatment in gnotobiotic mice. (A) Timeline of colonization, inhibitor dosage and sample collection. (B) Abundance ratios of bacterial species present in fecal samples as determined by COPRO-seq shows no statistically significant differences in abundance of each bacterial species before and after vehicle treatment, and inhibitor treatment at both dosages. Data are represented as mean ± SEM. Wilcoxon-sign ranked test, n = 8.
