## Supplementary material for "Phenotypic high-throughput screening identifies modulators of gut microbial choline metabolism": Document S2

### Supplementary Information Methods

#### General procedures:

**General procedure A — amine alkylation:** The phthalimide derivative (1.0 eq) was dissolved in DCM (0.25 M). The amine derivative as a free base or as a HCl salt (1.0–2.0 eq) and $NEt_3$ (2.4–5.0 eq) were added. The resulting mixture was heated at 50 °C overnight. The solution was concentrated under vacuum. The crude material was purified by chromatography on silica (Hex:EtOAc or EtOAc:MeOH).

**General procedure B — phthalimide deprotection:** The protected amine (1.0 eq) was dissolved in EtOH and $N_2H_4$ (2.4 eq) was added dropwise. The resulting mixture was heated at 70 °C for 2–6 h. Once cooled down, the solution was filtered through Celite. The filtrate was concentrated under vacuum and resuspended in DCM. The solution was filtered through Celite and the filtrate was concentrated under vacuum.

**General procedure C — bromination:** In an oven-dried round-bottom flask under argon, the reactant (1.0 eq) was dissolved in anhydrous DMF (0.33 M) and NBS (1.1 eq) was added portion wise. The resulting mixture was stirred at room temperature overnight.

**General procedure D — Suzuki reaction:** In an oven-dried round-bottom flask under argon, the halogenated derivative (1.0 eq), the boronic analog (1.2 eq), $Pd(OAc)_2$ (6 mol%), X-Phos (7 mol%), $Cs_2CO_3$ (2.5 eq) were added and the vial was properly capped. The mixture vessel was evacuated and backfilled with argon (process repeated 3 times). A mixture of *n*BuOH:$H_2O$ (4:1, 0.13 M) was added. The mixture vessel was evacuated and backfilled with argon (process repeated 3 times) and heated at 50 °C overnight. The reaction mixture was concentrated under vacuum. The residue was diluted with water and the aqueous phase was extracted 3 times with EtOAc. The combined organic layers were washed with brine, dried over $MgSO_4$ and concentrated under vacuum. The crude material was purified by chromatography on silica (Hex:EtOAc).

**General procedure E — saponification:** The ester derivative (1.0 eq) was dissolved in a mixture MeOH:$H_2O$ (4:1, 0.08 M) and LiOH or NaOH (5.0 eq) was added. The resulting mixture was heated at 70 °C overnight. The methanol was concentrated under vacuum. The basic aqueous phase was diluted and extracted 3 times with EtOAc. The aqueous phase was acidified to pH 1 using an aqueous 1M solution of HCl. The aqueous phase was reextracted 3 times with EtOAc. The combined organic layers were washed with brine, dried over $MgSO_4$ and concentrated under vacuum. The solid was used without further purification.

**General procedure F — amide coupling:** The free carboxylic acid (1.0 eq), HOBt (1.5 eq) and EDC•HCl (2.0 eq) were dissolved in DCM (0.1 M). The amine derivative (2.0 eq) and DIEA (4.0 eq) were added and the resulting mixture was stirred overnight. Water was added and the aqueous phase was extracted 3 times with DCM. The combined organic layers were washed with brine, dried over $MgSO_4$ and concentrated under vacuum. The crude material was purified by chromatography using a pretreated silica using a solution $Et_2O$:$NEt_3$ 95:5 (EtOAc:MeOH:$NEt_3$).

**General procedure G — *tert*-butyloxycarbonyl (Boc) deprotection:** The protected amine (1.0 eq) was dissolved in DCM (0.35 M) and cooled to 0 °C. TFA (14.6 eq) was added dropwise, and the resulting mixture was stirred at room temperature for 4 h. The reaction mixture was concentrated under vacuum, and the residue was resuspended in EtOAc. The organic phase was washed twice with a saturated solution of $NaHCO_3$ and brine, dried over $MgSO_4$ and concentrated under vacuum. The residue was used without further purification.

**General procedure H — pyrrole alkylation:** To the suspension of NaH (60% in mineral oil dispersion) (2.5 eq) in THF (2 M) at 0 °C was added a solution of pyrrole-3-carboxylate (1.0 eq) in THF (0.7 M) and stirred at 0 °C for 30 min followed by addition of alkyl halide (4.0 eq) and heated to 50 °C for 3 h. The reaction mixture was carefully quenched with water at 0 °C and extracted with EtOAc. Combined organic layer was dried over $Na_2SO_4$ and concentrated *in vacuo*. Purification by flash column chromatography ($SiO_2$, 25 g, Hex:EtOAc 0–15%).

**General procedure I — amide coupling:** To the solution of a carboxylic acid (1.0 eq) dissolved in MeCN (0.2 M) were added DIPEA (4.0 eq) followed by HATU (2.0 eq) and amine (2.0 eq) and stirred for 20 min. The reaction mixture was concentrated under reduced pressure. The residue was diluted with EtOAc and water, phases were separated; combined organic layers were washed with brine, dried over Na2SO4, and concentrated *in vacuo*. Purification was performed by column chromatography on silica with DCM to DCM:MeOH 5:1.

**General procedure J — pyrrole synthesis:** To a suspension of NaH (60% in mineral oil) (393 mg, 10.25 mmol, 1.2 eq) in THF (8 mL) was carefully added reactant (1.0 eq) followed by addition of the solution of 2-bromo-1-(3,4-dimethylphenyl)ethanoate (1.0 eq) in THF (3 mL) and the reaction was stirred at room temperature for 2 h. The reaction media was acidified with 1 M HCl, extracted with EtOAc (3 x 10 mL), washed with brine, dried over $MgSO_4$, and the solvent was removed under reduced pressure. Purification was performed by flash column chromatography ($SiO_2$, 50 g, Hex:EtOAc 0–20%).

**General procedure K — pyrrole formation:** To the solution of reactant (1.0 eq) dissolved in EtOH (4.3 mL) were added the amine (1.01 eq) and AcOH (5.0 eq) and stirred at 70 °C for 2 h. The solvent was removed under reduced pressure. Purification was performed by flash column chromatography ($SiO_2$, 25 g, Hex:EtOAc 0–50%).

### Preparation of amines

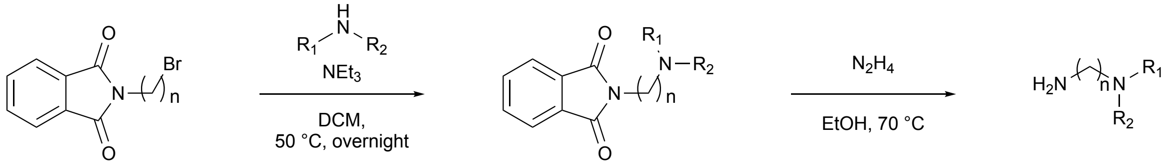


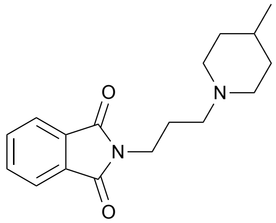

**2-(3-(4-methylpiperidin-1-yl)propyl)isoindoline-1,3-dione (S1)**

Following general procedure A, starting from 2-(3-bromopropyl)isoindoline-1,3-dione (2.8 g, 10.44 mmol, 1.0 eq), 4-methylpiperidine (1.2 mL, 10.44 mmol, 1.0 eq) and $NEt_3$ (3.5 mL, 25.06 mmol, 2.4 eq) and purifying by chromatography on silica gel using 1:1 Hex:EtOAc as eluent, 2-(3-(4-methylpiperidin-1-yl)propyl)isoindoline-1,3-dione was obtained as a yellow oil (1.58 g, 5.52 mmol, 53%). $^1$H NMR (400 MHz, $CDCl_3$) δ 7.83 (dd, 2H, *J* = 5.4 Hz, *J* = 3.1 Hz), 7.69 (dd, 2H, *J* = 5.5 Hz, *J* = 3.1 Hz), 3.74 (t, 2H, *J* = 6.9 Hz), 2.80 (d, 2H, *J* = 11.5 Hz), 2.36 (t, 2H, *J* = 7.1 Hz), 1.85 (quint., 2H, *J* = 7.0 Hz), 1.79 (td, 2H, *J* = 11.5 Hz, *J* = 2.3 Hz), 1.50 (d, 2H, *J* = 12.9 Hz), 1.26-1.21 (m, 1H), 0.98 (qd, 2H, *J* = 12.1 Hz, *J* = 3.8 Hz), 0.80 (d, 3H, *J* = 6.5 Hz); $^{13}$C NMR (101 MHz, $CDCl_3$) δ 168.6, 133.9, 132.5, 123.3, 56.7, 54.1, 37.0, 34.4, 30.8, 25.7, 22.0.

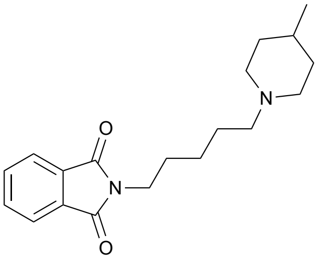

**2-(5-(4-methylpiperidin-1-yl)pentyl)isoindoline-1,3-dione (S2)**

Following general procedure A, starting from 2-(5-bromopentyl)isoindoline-1,3-dione (700 mg, 2.36 mmol, 1.0 eq), 4-methylpiperidine (559 µL, 4.73 mmol, 2.0 eq) and $NEt_3$ (791 µL, 5.67 mmol, 2.4 eq) and purifying by chromatography on silica gel using EtOAc as eluent, 2-(5-(4-methylpiperidin-1-yl)pentyl)isoindoline-1,3-dione was obtained as a colorless oil (540 mg, 1.72 mmol, 73%). $^1$H NMR (400 MHz, $CDCl_3$) δ 7.83 (dd, 2H, *J* = 5.4 Hz, *J* = 3.2 Hz), 7.70 (dd, 2H, *J* = 5.4 Hz, *J* = 3.1 Hz), 3.67 (t, 2H, *J* = 7.2 Hz), 2.84 (d, 2H, *J* = 8.0 Hz), 2.27 (t, 2H, *J* = 7.7 Hz), 1.84 (d, 2H, *J* = 11.5 Hz), 1.68 (quint., 2H, *J* = 7.3 Hz), 1.60-1.56 (m, 2H), 1.55-1.49 (m, 2H), 1.35-1.28 (m, 3H), 1.20 (t, 2H, *J* = 11.7 Hz), 0.89 (d, 3H, *J* = 6.3 Hz); $^{13}$C NMR (101 MHz, $CDCl_3$) δ 168.8, 134.2, 132.5, 123.5, 59.3, 54.4, 52.9, 38.3, 34.7, 31.2, 28.9, 27.0, 25.4.

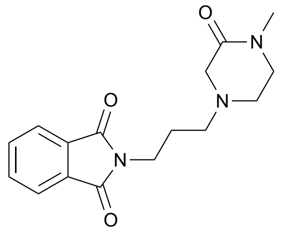

**2-(3-(4-methyl-3-oxopiperazin-1-yl)propyl)isoindoline-1,3-dione (S3)**
Following general procedure A, starting from 2-(3-bromopropyl)isoindoline-1,3-dione (600 mg, 2.24 mmol, 1.0 eq), 1-methylpiperazin-2-one (511 mg, 4.48 mmol, 2.0 eq) and $NEt_3$ (749 µL, 5.38 mmol, 2.4 eq) and purifying by chromatography on silica gel using 9:1 EtOAc:MeOH as eluent, 2-(3-(4-methyl-3-oxopiperazin-1-yl)propyl)isoindoline-1,3-dione was obtained as a yellow oil (105 mg, 0.35 mmol, 16%). $^1$H NMR (400 MHz, $CDCl_3$) δ 7.84-7.82 (m, 2H), 7.72-7.69 (m, 2H), 3.77 (t, 2H, *J* = 7.0 Hz), 3.19 (t, 2H, *J* = 5.1 Hz), 3.07 (s, 2H), 2.84 (s, 3H), 2.62 (t, 2H, *J* = 5.7 Hz), 2.46 (t, 2H, *J* = 6.7 Hz), 1.85 (quint., 2H, *J* = 6.9 Hz); $^{13}$C NMR (101 MHz, $CDCl_3$) δ 168.8, 167.3, 134.2, 132.6, 123.5, 57.7, 55.2, 49.7, 48.9, 26.6, 34.0, 25.6.

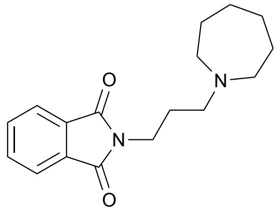

**2-(3-(azepan-1-yl)propyl)isoindoline-1,3-dione (S4)**
Following general procedure A, starting from 2-(3-bromopropyl)isoindoline-1,3-dione (1.0 g, 3.73 mmol, 1.0 eq), azepane (420 µL, 3.73 mmol, 1.0 eq) and $NEt_3$ (1.2 mL, 8.95 mmol, 2.4 eq) and purifying by chromatography on silica gel using 9:1 EtOAc:MeOH as eluent, 2-(3-(azepan-1-yl)propyl)isoindoline-1,3-dione was obtained as a yellow oil (446 mg, 1.56 mmol, 42%). $^1$H NMR (400 MHz, $CDCl_3$) δ 7.85-7.81 (m, 2H), 7.73-7.68 (m, 2H), 3.78-3.73 (m, 2H), 2.59-2.53 (m, 6H), 1.85 (quint., 2H, *J* = 6.6 Hz), 1.59 (br s, 4H), 1.52 (br s, 4H); $^{13}$C NMR (101 MHz, $CDCl_3$) δ 168.8, 134.2, 132.6, 123.5, 56.1, 55.8, 36.9, 27.3.

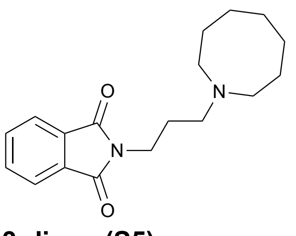

**2-(3-(azocan-1-yl)propyl)isoindoline-1,3-dione (S5)**
Following general procedure A, starting from 2-(3-bromopropyl)isoindoline-1,3-dione (1.0 g, 3.73 mmol, 1.0 eq), azocane (471 µL, 3.73 mmol, 1.0 eq) and $NEt_3$ (1.2 mL, 8.95 mmol, 2.4 eq) and purifying by chromatography on silica gel using EtOAc as eluent, 2-(3-(azocan-1-yl)propyl)isoindoline-1,3-dione was obtained as a yellow oil (792 mg, 2.64 mmol, 71%). $^1$H NMR (400 MHz, $CDCl_3$) δ 7.86-7.82 (m, 2H), 7.74-7.68 (m, 2H), 3.85 (t, 1H, *J* = 6.5 Hz), 3.78 (t, 2H, *J* = 6.5 Hz), 3.57 (t, 1H, *J* = 6.5 Hz), 2.51 (br s, 4H), 1.81 (br s, 2H), 1.63 (br s, 2H), 1.57 (br s, 8H); $^{13}$C NMR (101 MHz, $CDCl_3$) δ 168.8, 134.4, 134.2, 132.6, 123.7, 123.5, 54.7, 42.3, 36.9, 36.0, 31.8, 28.4, 27.7, 26.5.

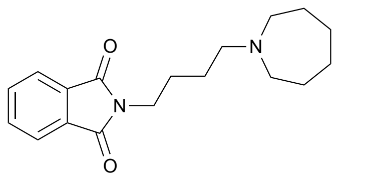

**2-(4-(azepan-1-yl)butyl)isoindoline-1,3-dione (S6)**

Following general procedure A, starting from 2-(4-bromobutyl)isoindoline-1,3-dione (4.8 g, 17.01 mmol, 1.0 eq), azepane (1.9 mL, 17.01 mmol, 1.0 eq) and $NEt_3$ (5.7 mL, 40.80 mmol, 2.4 eq) and purifying by chromatography on silica gel using 1:3 Hex:EtOAc as eluent, 2-(4-(azepan-1-yl)butyl)isoindoline-1,3-dione was obtained as a yellow oil (3.4 g, 11.42 mmol, 67%). $^1$H NMR (400 MHz, $CDCl_3$) δ 7.83 (dd, 2H, *J* = 4.7 Hz, *J* = 3.1 Hz), 7.70 (dd, 1H, *J* = 4.8 Hz, *J* = 3.1 Hz), 3.70 (t, 2H, *J* = 7.2 Hz), 2.61-2.58 (m, 4H), 2.48 (t, 2H, *J* = 7.5 Hz), 1.72-1.46 (m, 14H); $^{13}$C NMR (101 MHz, $CDCl_3$) δ 168.8, 134.3, 132.6, 58.0, 55.9, 38.3, 28.3, 27.3, 27.0, 25.3.

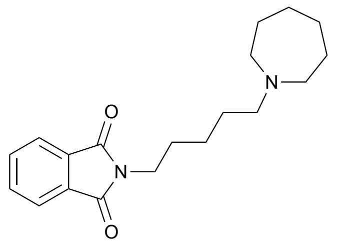

**2-(5-(azepan-1-yl)pentyl)isoindoline-1,3-dione (S7)**

Following general procedure A, starting from 2-(5-bromopentyl)isoindoline-1,3-dione (700 mg, 2.36 mmol, 1.0 eq), azepane (533 µL, 4.73 mmol, 2.0 eq) and $NEt_3$ (791 µL, 5.67 mmol, 2.4 eq) and purifying by chromatography on silica gel 9:1 EtOAc:MeOH as eluent, 2-(5-(azepan-1-yl)pentyl)isoindoline-1,3-dione was obtained as a yellow solid (458 mg, 1.46 mmol, 62%). $^1$H NMR (400 MHz, $CDCl_3$) δ 7.83 (dd, 2H, *J* = 5.5 Hz, *J* = 3.1 Hz), 7.70 (dd, 2H, *J* = 5.5 Hz, *J* = 3.1 Hz), 3.68 (t, 2H, *J* = 7.1 Hz), 2.82 (t, 4H, *J* = 5.4 Hz), 2.62 (t, 2H, *J* = 7.9 Hz), 1.75-1.60 (m, 12H), 1.34 (quint., 2H, *J* = 7.5 Hz); $^{13}$C NMR (101 MHz, $CDCl_3$) δ 168.8, 134.3, 132.4, 123.5, 58.1, 55.4, 38.0, 28.6, 27.3, 26.6, 26.0, 24.8.

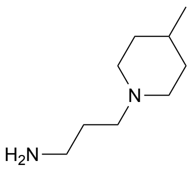

**3-(4-methylpiperidin-1-yl)propan-1-amine (S8)**

Following general procedure B, starting from 2-(3-(4-methylpiperidin-1-yl)propyl)isoindoline-1,3-dione (**S1**) (2.18 g, 7.61 mmol, 1.0 eq), 3-(4-methylpiperidin-1-yl)propan-1-amine was obtained as a yellow oil (932 mg, 5.97 mmol, 78%). $^1$H NMR (400 MHz, $CDCl_3$) δ 2.88 (d, 2H, *J* = 11.5 Hz), 2.75 (t, 2H, *J* = 6.7 Hz), 2.36 (t, 2H, *J* = 7.2 Hz), 1.99 (br s, 2H), 1.87 (dd, 2H, *J* = 11.9 Hz, *J* = 1.9 Hz), 1.68-1.59 (m, 4H), 1.33 (br s, 1H), 1.23 (td, 2H, *J* = 11.9 Hz, *J* = 3.7 Hz), 0.90 (d, 3H, *J* = 6.4 Hz); $^{13}$C NMR (101 MHz, $CDCl_3$) δ 57.4, 54.5, 41.4, 34.7, 31.2, 30.7, 22.2.

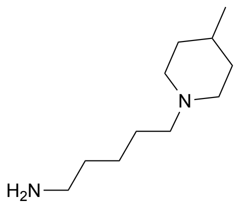

**5-(4-methylpiperidin-1-yl)pentan-1-amine (S9)**

Following general procedure B, starting from 2-(5-(4-methylpiperidin-1-yl)pentyl)isoindoline-1,3-dione (**S2**) (539 mg, 1.71 mmol, 1.0 eq), 5-(4-methylpiperidin-1-yl)pentan-1-amine was obtained as a yellow solid (316 mg, 0.95 mmol, 56%). $^1$H NMR (400 MHz, $CDCl_3$) δ 2.88 (d, 2H, *J* = 11.7 Hz), 2.68 (t, 2H, *J* = 7.0 Hz), 2.29 (t, 2H, *J* = 7.7 Hz), 1.87 (td, 2H, *J* = 11.8 Hz, *J* = 2.1 Hz), 1.60 (d, 2H, *J* = 12.2 Hz), 1.55-1.48 (m, 2H), 1.45 (quint., 2H, *J* = 7.3 Hz), 1.35-1.28 (m, 3H), 1.24 (qd, 2H, *J* = 12.2 Hz, *J* = 3.5 Hz), 0.91 (d, 3H, *J* = 6.2 Hz); $^{13}$C NMR (101 MHz, $CDCl_3$) δ 59.5, 54.4, 42.5, 34.6, 34.1, 31.2, 27.3, 25.4, 22.2.

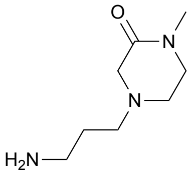

**4-(3-aminopropyl)-1-methylpiperazin-2-one (S10)**
Following general procedure B, starting from 2-(3-(4-methyl-3-oxopiperazin-1-yl)propyl)isoindoline-1,3-dione (**S3**) (104 mg, 0.35 mmol, 1.0 eq), 4-(3-aminopropyl)-1-methylpiperazin-2-one was obtained as an orange oil (55 mg, 0.32 mmol, 93%). $^1$H NMR (400 MHz, $CDCl_3$) δ 3.31 (t, 2H, *J* = 5.4 Hz), 3.12 (s, 2H), 2.93 (s, 3H), 2.81 (t, 2H, *J* = 6.7 Hz), 2.67 (t, 2H, *J* = 5.7 Hz), 2.62 (br s, 2H), 2.47 (t, 2H, *J* = 7.0 Hz), 1.67 (quint., 2H, *J* = 6.9 Hz); $^{13}$C NMR (101 MHz, $CDCl_3$) δ 167.5, 57.8, 55.8, 53.8, 50.1, 48.9, 40.6, 34.1.

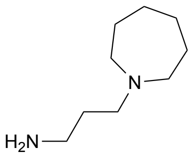

**3-(azepan-1-yl)propan-1-amine (S11)**
Following general procedure B, starting from 2-(3-(azepan-1-yl)propyl)isoindoline-1,3-dione (**S4**) (445 mg, 1.55 mmol, 1.0 eq), 3-(azepan-1-yl)propan-1-amine was obtained as an orange oil (218 mg, 1.40 mmol, 90%). $^1$H NMR (400 MHz, $CDCl_3$) δ 3.00 (br s, 2H), 2.79 (t, 2H, *J* = 6.6 Hz), 2.66-2.63 (m, 4H), 2.56 (t, 2H, *J* = 6.9 Hz), 1.67-1.62 (m, 6H), 1.59-1.56 (m, 4H)

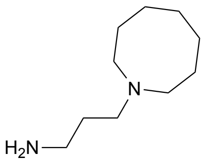

**3-(azocan-1-yl)propan-1-amine (S12)**
Following general procedure B, starting from 2-(3-(azocan-1-yl)propyl)isoindoline-1,3-dione (**S5**) (792 mg, 2.64 mmol, 1.0 eq), 3-(azocan-1-yl)propan-1-amine was obtained as an orange oil (449 mg, 2.64 mmol, 100%). $^1$H NMR (400 MHz, $CDCl_3$) δ 2.94 (br s, 2H), 2.82 (t, 2H, *J* = 6.6 Hz), 2.58-2.51 (m, 6H), 1.67-1.56 (m, 12)

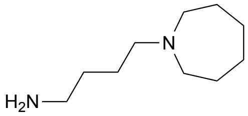

**4-(azepan-1-yl)butan-1-amine (S13)**
Following general procedure B, starting from 2-(4-(azepan-1-yl)butyl)isoindoline-1,3-dione (**S6**) (3.4 g, 11.42 mmol, 1.0 eq), 4-(azepan-1-yl)butan-1-amine was obtained as a yellow oil (1.9 g, 11.42 mmol, 100%). $^1$H NMR (400 MHz, $CDCl_3$) δ 4.49 (br s, 2H), 3.40 (t, 2H, *J* = 6.1 Hz), 3.34-3.30 (m, 4H), 3.16-3.12 (m, 2H), 2.29-2.27 (m, 6H), 2.23-2.19 (m, 6H); $^{13}$C NMR (101 MHz, $CDCl_3$) δ 57.8, 55.4, 41.4, 30.3, 27.3, 27.1, 25.5.

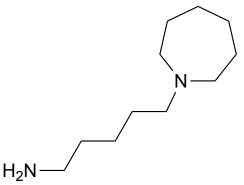

**5-(azepan-1-yl)pentan-1-amine (S14)**

Following general procedure B, starting from 2-(5-(azepan-1-yl)pentyl)isoindoline-1,3-dione (**S7**) (457 mg, 1.45 mmol, 1.0 eq), 5-(azepan-1-yl)pentan-1-amine was obtained as a yellow oil (142 mg, 0.769 mmol, 53%). $^{1}$H NMR (400 MHz, $CDCl_3$) δ 2.84 (br s, 2H), 2.74 (t, 4H, *J* = 5.4 Hz), 2.68 (t, 2H, *J* = 6.7 Hz), 2.56 (t, 2H, *J* = 7.5 Hz), 1.90 (br s, 4H), 1.60 (br s, 4H), 1.59-1.54 (m, 2H), 1.48 (quint., 2H, *J* = 7.3 Hz), 1.32 (quint., 2H, *J* = 7.3 Hz); $^{13}$C NMR (101 MHz, $CDCl_3$) δ 58.3, 55.6, 42.2, 33.4, 27.3, 27.2, 26.9, 25.0.

### Preparation of analogs

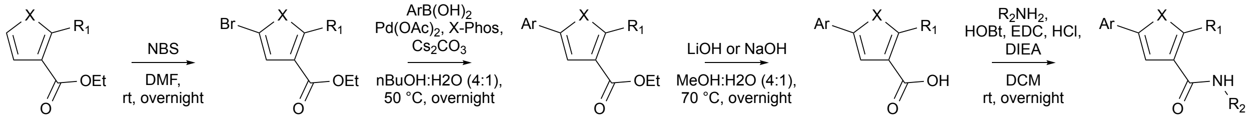


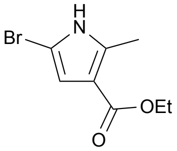

**Ethyl 5-bromo-2-methyl-1*H*-pyrrole-3-carboxylate (S15)**

Following general procedure C, starting from ethyl 2-methyl-1*H*-pyrrole-3-carboxylate (1.0 g, 6.53 mmol, 1.0 eq) and purifying by filtration after addition of water to the reaction mixture, ethyl 5-bromo-2-methyl-1*H*-pyrrole-3-carboxylate was obtained as a tan solid (1.0 g, 4.28 mmol, 66%). $^{1}$H NMR (400 MHz, DMSO-$d_6$) δ 11.88 (s, 1H), 6.31 (s, 1H), 4.13 (q, 2H, *J* = 7.1 Hz), 2.37 (s, 3H), 1.23 (t, 3H, *J* = 7.1 Hz); $^{13}$C NMR (101 MHz, DMSO-$d_6$) δ 163.6, 136.4, 112.3, 110.7, 96.1, 58.9, 14.3, 12.5.

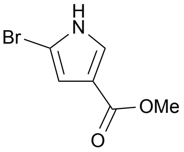

**Methyl 5-bromo-1*H*-pyrrole-3-carboxylate (S16)**

Following general procedure C, starting from methyl 1*H*-pyrrole-3-carboxylate (200 mg, 1.60 mmol, 1.0 eq) and purifying by chromatography on silica gel using 4:1 Hex:EtOAc as eluent, methyl 5-bromo-1*H*-pyrrole-3-carboxylate was obtained as a white solid (234 mg, 1.14 mmol, 72%). $^{1}$H NMR (400 MHz, DMSO-$d_6$) δ 12.19 (s, 1H), 7.45 (t, 1H, *J* = 1.9 Hz), 6.44 (t, 1H, *J* = 1.8 Hz), 3.69 (s, 3H); $^{13}$C NMR (101 MHz, DMSO-$d_6$) δ 163.5, 125.1, 116.3, 110.3, 99.5, 50.8.

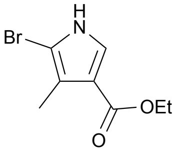

**Ethyl 5-bromo-4-methyl-1*H*-pyrrole-3-carboxylate (S17)**

Following general procedure C, starting from ethyl 4-methyl-1*H*-pyrrole-3-carboxylate (200 mg, 1.31 mmol, 1.0 eq) and purifying by chromatography on silica gel using 4:1 Hex:EtOAc as eluent, ethyl 5-bromo-4-methyl-1*H*-pyrrole-3-carboxylate was obtained as a white solid (192 mg, 0.83 mmol, 63%). $^{1}$H NMR (400

MHz, $CDCl_3$) δ 8.36 (s, 1H), 7.37 (d, 1H, *J* = 3.2 Hz), 3.80 (s, 3H), 2.23 (s, 3H); $^{13}$C NMR (101 MHz, $CDCl_3$) δ 165.2, 124.6, 120.4, 116.1, 100.1, 51.3, 11.5.

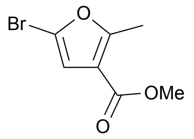


**Methyl 5-bromo-2-methylfuran-3-carboxylate (S18)**
Following general procedure C, starting from methyl 2-methylfuran-3-carboxylate (500 mg, 3.57 mmol, 1.0 eq) and purifying by chromatography on silica gel using 95:5 Hex:EtOAc as eluent, methyl 5-bromo-2-methylfuran-3-carboxylate was obtained as a yellow oil (552 mg, 2.52 mmol, 71%). $^1$H NMR (400 MHz, $CDCl_3$) δ 6.55 (s, 1H), 3.81 (s, 3H), 2.56 (s, 3H); $^{13}$C NMR (101 MHz, $CDCl_3$) δ 163.7, 161.1, 120.3, 112.4, 51.9, 14.1.

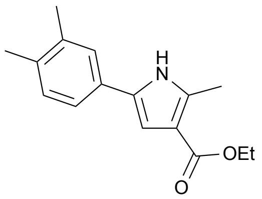


**Ethyl 5-(3,4-dimethylphenyl)-2-methyl-1*H*-pyrrole-3-carboxylate (S19)**
Following general procedure D, starting from ethyl 5-bromo-2-methyl-1*H*-pyrrole-3-carboxylate (**S15**) (500 mg, 1.96 mmol, 1.0 eq) and 3,4-dimethylphenylboronic acid (353 mg, 2.35 mmol, 1.2 eq) and purifying by chromatography on silica gel using 6:1 Hex:EtOAc as eluent, ethyl 5-(3,4-dimethylphenyl)-2-methyl-1*H*-pyrrole-3-carboxylate was obtained as a tan solid (453 mg, 1.96 mmol, 90%). $^1$H NMR (400 MHz, DMSO-$d_6$) δ 11.49 (s, 1H), 7.43 (s, 1H), 7.33 (d, 1H, *J* = 7.8 Hz), 7.10 (d, 1H, *J* = 7.8 Hz), 6.67 (s, 1H), 4.15 (q, 2H, *J* = 7.1 Hz), 2.46 (s, 3H), 2.23 (s, 3H), 2.19 (s, 3H), 1.26 (t, 3H, *J* = 7.1 Hz); $^{13}$C NMR (101 MHz, DMSO-$d_6$) δ 164.6, 136.4, 136.1, 134.0, 129.8, 129.5, 124.7, 120.9, 111.8, 105.7, 58.6, 19.4, 19.0, 14.5, 12.7. (1 peak missing, probably overlap)

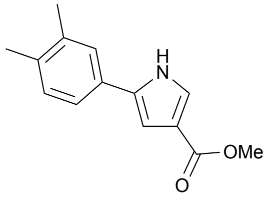


**Methyl 5-(3,4-dimethylphenyl)-1*H*-pyrrole-3-carboxylate (S20)**
Following general procedure D, starting methyl 5-bromo-1*H*-pyrrole-3-carboxylate (**S16**) (300 mg, 1.47 mmol, 1.0 eq) and (3,4-dimethylphenyl)boronic acid (265 mg, 1.77 mmol, 1.2 eq) and purifying by chromatography on silica gel using 4:1 Hex:EtOAc as eluent, methyl 5-(3,4-dimethylphenyl)-1*H*-pyrrole-3-carboxylate was obtained as a brown solid (305 mg, 1.33 mmol, 90%). $^1$H NMR (400 MHz, DMSO-$d_6$) δ 11.85 (s, 1H), 7.48 (s, 1H), 7.46 (br s, 1H), 7.38 (d, 1H, *J* = 7.8 Hz), 7.13 (d, 1H, *J* = 7.8 Hz), 6.80 (s, 1H), 3.71 (s, 3H), 2.24 (s, 3H), 2.21 (s, 3H)

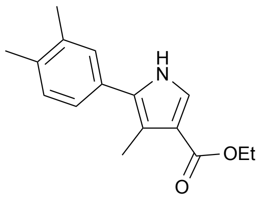


**Ethyl 5-(3,4-dimethylphenyl)-4-methyl-1*H*-pyrrole-3-carboxylate (S21)**

Following general procedure D, starting ethyl 5-bromo-4-methyl-1*H*-pyrrole-3-carboxylate (**S17**) (183 mg, 0.79 mmol, 1.0 eq) and (3,4-dimethylphenyl)boronic acid (142 mg, 0.95 mmol, 1.2 eq) and purifying by chromatography on silica gel using 2:1 Hex:EtOAc as eluent, ethyl 5-(3,4-dimethylphenyl)-4-methyl-1*H*-pyrrole-3-carboxylate was obtained as a brown oil. $^1$H NMR (400 MHz, $CDCl_3$) δ 8.42 (s, 1H), 7.42 (d, 1H, *J* = 3.2 Hz), 7.19 (d, 1H, *J* = 3.7 Hz), 7.18-7.14 (m, 2H), 3.67-3.63 (m, 2H), 2.41 (s, 3H), 2.30 (s, 3H), 2.29 (s, 3H), 0.94 (t, 3H, *J* = 7.3 Hz); $^{13}$C NMR (101 MHz, $CDCl_3$) δ 166.3, 137.4, 135.9, 130.4, 128.9, 128.6, 125.2, 124.7, 123.8, 117.4, 116.2, 63.1, 51.1, 35.2, 14.2, 11.4.

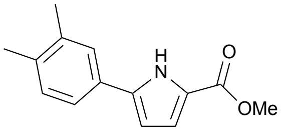


**Methyl 5-(3,4-dimethylphenyl)-1*H*-pyrrole-2-carboxylate (S22)**
Following general procedure D, starting methyl 5-bromo-1*H*-pyrrole-2-carboxylate (200 mg, 0.98 mmol, 1.0 eq) and (3,4-dimethylphenyl)boronic acid (176 mg, 1.18 mmol, 1.2 eq) and purifying by chromatography on silica gel using 4:1 Hex:EtOAc as eluent, methyl 5-(3,4-dimethylphenyl)-1*H*-pyrrole-2-carboxylate was obtained as a brown solid (86 mg, 0.37 mmol, 38%). $^1$H NMR (400 MHz, $CDCl_3$) δ 9.37 (s, 1H), 7.34 (s, 1H), 7.31 (dd, 1H, *J* = 7.8 Hz, *J* = 1.7 Hz), 7.16 (d, 1H, *J* = 7.8 Hz), 6.95 (dd, 1H, *J* = 3.8 Hz, *J* = 2.4 Hz), 6.49 (dd, 1H, *J* = 3.7 Hz, *J* = 2.9 Hz), 3.87 (s, 3H), 2.31 (s, 3H), 2.28 (s, 3H); $^{13}$C NMR (101 MHz, $CDCl_3$) δ 162.1, 137.6, 137.5, 136.8, 130.6, 129.3, 126.4, 122.9, 122.5, 119.2, 107.8, 51.8, 20.2, 19.9.

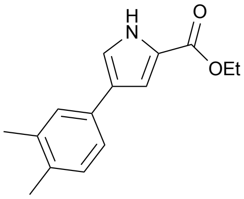


**Ethyl 4-(3,4-dimethylphenyl)-1*H*-pyrrole-2-carboxylate (S23)**
Following general procedure D, starting ethyl 4-bromo-1*H*-pyrrole-2-carboxylate (200 mg, 0.92 mmol, 1.0 eq) and (3,4-dimethylphenyl)boronic acid (165 mg, 1.10 mmol, 1.2 eq) and purifying by chromatography on silica gel using 4:1 Hex:EtOAc as eluent, ethyl 4-(3,4-dimethylphenyl)-1*H*-pyrrole-2-carboxylate was obtained as a yellow solid (196 mg, 0.81 mmol, 88%). $^1$H NMR (400 MHz, $CDCl_3$) δ 9.20 (s, 1H), 7.31 (s, 1H), 7.26 (dd, 1H, *J* = 7.8 Hz, *J* = 1.8 Hz), 7.20-7.17 (m, 2H), 7.12 (d, 1H, *J* = 7.8 Hz), 4.35 (q, 2H, *J* = 7.1 Hz), 2.29 (s, 3H), 2.27 (s, 3H), 1.38 (t, 3H, *J* = 7.2 Hz); $^{13}$C NMR (101 MHz, $CDCl_3$) δ 161.6, 137.2, 135.0, 132.5, 130.4, 127.3, 127.1, 123.9, 123.1, 119.5, 112.7, 60.8, 20.2, 19.8, 14.8.

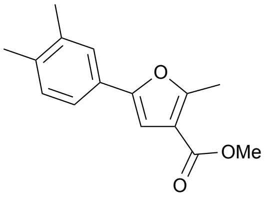


**Methyl 5-(3,4-dimethylphenyl)-2-methylfuran-3-carboxylate (S24)**
Following general procedure D, starting methyl 5-bromo-2-methylfuran-3-carboxylate (**S18**) (200 mg, 0.91 mmol, 1.0 eq) and (3,4-dimethylphenyl)boronic acid (164 mg, 1.10 mmol, 1.2 eq) and purifying by chromatography on silica gel using 9:1 Hex:EtOAc as eluent, methyl 5-(3,4-dimethylphenyl)-2-methylfuran-3-carboxylate was obtained as a white solid (223 mg, 0.91 mmol, 100%). $^1$H NMR (400 MHz, $CDCl_3$) δ 7.42 (s, 1H), 7.38 (d, 1H, *J* = 7.9 Hz), 7.14 (d, 1H, *J* = 7.9 Hz), 6.81 (s, 1H), 3.85 (s, 3H), 2.65 (s, 3H), 2.30 (s, 3H), 2.28 (s, 3H); $^{13}$C NMR (101 MHz, $CDCl_3$) δ 164.9, 158.6, 152.4, 137.2, 136.6, 130.3, 128.1, 125.2, 121.5, 115.3, 104.8, 51.6, 20.1, 19.9, 14.2.

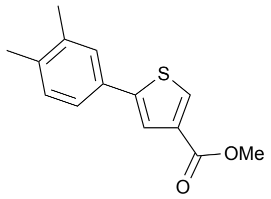

**Methyl 5-(3,4-dimethylphenyl)thiophene-3-carboxylate (S25)**

Following general procedure D, starting methyl 5-bromothiophene-3-carboxylate (100 mg, 0.45 mmol, 1.0 eq) and (3,4-dimethylphenyl)boronic acid (81 mg, 0.54 mmol, 1.2 eq) and purifying by chromatography on silica gel using 9:1 Hex:EtOAc as eluent, methyl 5-(3,4-dimethylphenyl)thiophene-3-carboxylate was obtained as a tun solid (73 mg, 0.30 mmol, 66%). $^{1}$H NMR (400 MHz, $CDCl_3$) δ 7.99 (d, 1H, *J* = 1.3 Hz), 7.66 (d, 1H, *J* = 1.3 Hz), 7.38 (br s, 1H), 7.34 (dd, 1H, *J* = 7.8 Hz, *J* = 1.8 Hz), 7.15 (d, 1H, *J* = 7.8 Hz), 3.88 (s, 3H), 2.31 (s, 3H), 2.28 (s, 3H); $^{13}$C NMR (101 MHz, $CDCl_3$) δ 163.6, 145.7, 137.6, 137.2, 134.4, 131.5, 131.4, 130.6, 127.5, 123.7, 123.0, 52.2, 20.1, 19.9.

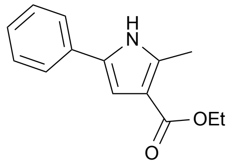

**Ethyl 2-methyl-5-phenyl-1*H*-pyrrole-3-carboxylate (S26)**

Following general procedure D, starting from ethyl 5-bromo-2-methyl-1*H*-pyrrole-3-carboxylate (**S15**) (100 mg, 0.43 mmol, 1.0 eq) and phenylboronic acid (63 mg, 0.52 mmol, 1.2 eq) and purifying by chromatography on silica gel using 95:5 to 9:1 Hex:EtOAc as eluent, ethyl 2-methyl-5-phenyl-1*H*-pyrrole-3-carboxylate was obtained as a white solid (92 mg, 0.40 mmol, 93%). $^{1}$H NMR (400 MHz, $CDCl_3$) δ 8.35 (br s, 1H), 7.45 (d, 2H, *J* = 7.3 Hz), 7.37 (t, 2H, *J* = 7.3 Hz), 7.23 (t, 1H, *J* = 7.3 Hz), 6.83 (d, 1H, *J* = 3.0 Hz), 4.30 (q, 2H, *J* = 7.1 Hz), 2.60 (s, 3H), 1.37 (t, 3H, *J* = 7.1 Hz)

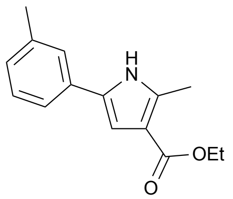

**Ethyl 2-methyl-5-(m-tolyl)-1*H*-pyrrole-3-carboxylate (S27)**

Following general procedure D, starting from ethyl 5-bromo-2-methyl-1*H*-pyrrole-3-carboxylate (**S15**) (100 mg, 0.39 mmol, 1.0 eq) and m-tolylboronic acid (64 mg, 0.47 mmol, 1.2 eq) and purifying by chromatography on silica gel using 4:1 Hex:EtOAc as eluent, ethyl 2-methyl-5-(m-tolyl)-1*H*-pyrrole-3-carboxylate e was obtained as a brown oil (95 mg, 0.39 mmol, 100%). $^{1}$H NMR (400 MHz, $CDCl_3$) δ 8.43 (br s, 1H), 7.28 (br s, 1H), 7.25 (s, 1H), 7.06-7.04 (m, 2H), 6.82 (d, 1H, *J* = 3.0 Hz), 4.29 (q, 2H, *J* = 7.2 Hz), 2.59 (s, 3H), 2.37 (s, 3H), 1.36 (t, 3H, *J* = 7.2 Hz)

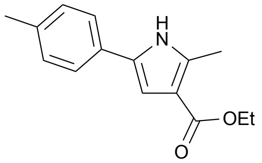

**Ethyl 2-methyl-5-(p-tolyl)-1*H*-pyrrole-3-carboxylate (S28)**

Following general procedure D, starting from ethyl 5-bromo-2-methyl-1*H*-pyrrole-3-carboxylate (**S15**) (120 mg, 0.47 mmol, 1.0 eq) and p-tolylboronic acid (77 mg, 0.56 mmol, 1.2 eq) and purifying by chromatography on silica gel using 6:1 Hex:EtOAc as eluent, ethyl 2-methyl-5-(p-tolyl)-1*H*-pyrrole-3-carboxylate was obtained as an orange solid (72 mg, 0.29 mmol, 63%). $^{1}$H NMR (400 MHz, DMSO-$d_6$) δ 11.52 (s, 1H), 7.52

(d, 2H, *J* = 8.1 Hz), 7.17 (d, 2H, *J* = 8.1 Hz), 6.68 (d, 1H, *J* = 2.9 Hz), 4.17 (q, 2H, *J* = 7.1 Hz), 2.47 (s, 3H), 2.28 (s, 3H), 1.26 (t, 3H, *J* = 7.1 Hz); $^{13}$C NMR (101 MHz, DMSO-$d_6$) δ 164.6, 136.1, 135.2, 129.7, 129.3, 129.1, 123.4, 111.8, 105.9, 58.6, 20.7, 14.4, 12.7.

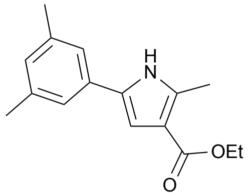


**Ethyl 5-(3,5-dimethylphenyl)-2-methyl-1*H*-pyrrole-3-carboxylate (S29)**
Following general procedure D, starting from ethyl 5-bromo-2-methyl-1*H*-pyrrole-3-carboxylate (**S15**) (150 mg, 0.65 mmol, 1.0 eq) and (3,5-dimethylphenyl)boronic acid (116 mg, 0.78 mmol, 1.2 eq) and purifying by chromatography on silica gel using 6:1 Hex:EtOAc as eluent, ethyl 5-(3,5-dimethylphenyl)-2-methyl-1*H*-pyrrole-3-carboxylate was obtained as an orange solid (102 mg, 0.39 mmol, 61%). $^{1}$H NMR (400 MHz, $CDCl_3$) δ 8.45 (s, 1H), 7.08 (s, 2H), 6.87 (s, 1H), 6.81 (d, 1H, *J* = 3.0 Hz), 4.29 (q, 2H, *J* = 7.1 Hz), 2.59 (s, 3H), 2.33 (s, 6H), 1.36 (t, 3H, *J* = 7.1 Hz); $^{13}$C NMR (101 MHz, $CDCl_3$) δ 165.7, 138.6, 136.0, 131.8, 130.3, 128.5, 121.7, 113.4, 107.3, 59.6, 21.5, 14.7, 13.5.

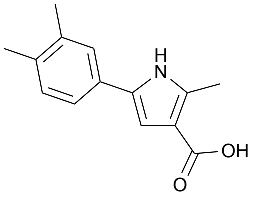


**5-(3,4-dimethylphenyl)-2-methyl-1*H*-pyrrole-3-carboxylic acid (S30)**
Following general procedure E, starting from ethyl 5-(3,4-dimethylphenyl)-2-methyl-1*H*-pyrrole-3-carboxylate (**S19**) (241 mg, 0.94 mmol, 1.0 eq) and LiOH (112 mg, 4.68 mmol, 5.0 eq), 5-(3,4-dimethylphenyl)-2-methyl-1*H*-pyrrole-3-carboxylic acid was obtained as an orange solid (138 mg, 0.60 mmol, 64%).

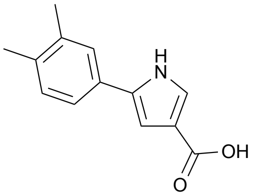


**5-(3,4-dimethylphenyl)-1*H*-pyrrole-3-carboxylic acid (S31)**
Following general procedure E, starting from methyl 5-(3,4-dimethylphenyl)-1*H*-pyrrole-3-carboxylate (**S20**) (67 mg, 0.29 mmol, 1.0 eq) and LiOH (35 mg, 1.46 mmol, 5.0 eq), 5-(3,4-dimethylphenyl)-1*H*-pyrrole-3-carboxylic acid was obtained as a greenish solid (50 mg, 023 mmol, 79%).

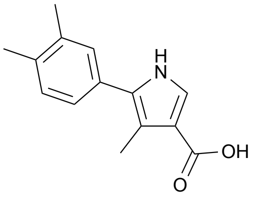


**5-(3,4-dimethylphenyl)-4-methyl-1*H*-pyrrole-3-carboxylic acid (S32)**
Following general procedure E, starting from methyl 5-(3,4-dimethylphenyl)-4-methyl-1*H*-pyrrole-3-carboxylate (**S21**) (203 mg, 0.84 mmol, 1.0 eq) and NaOH (167 mg, 4.17 mmol, 5.0 eq), 5-(3,4-dimethylphenyl)-4-methyl-1*H*-pyrrole-3-carboxylic acid was obtained as a brown solid (50 mg, 0.22 mmol, 26%).

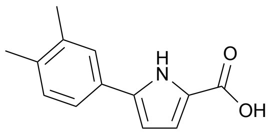

**5-(3,4-dimethylphenyl)-1*H*-pyrrole-2-carboxylic acid (S33)**
Following general procedure E, starting from methyl 5-(3,4-dimethylphenyl)-1*H*-pyrrole-2-carboxylate (**S22**) (85 mg, 0.37 mmol, 1.0 eq) and LiOH (44 mg, 1.85 mmol, 5.0 eq), 5-(3,4-dimethylphenyl)-1*H*-pyrrole-2-carboxylic acid was obtained as a purple solid (80 mg, 0.37 mmol, 100%).

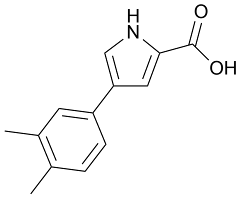

**4-(3,4-dimethylphenyl)-1*H*-pyrrole-2-carboxylic acid (S34)**
Following general procedure E, starting from ethyl 4-(3,4-dimethylphenyl)-1*H*-pyrrole-2-carboxylate (**S23**) (195 mg, 0.80 mmol, 1.0 eq) and LiOH (96 mg, 4.01 mmol, 5.0 eq), 4-(3,4-dimethylphenyl)-1*H*-pyrrole-2-carboxylic acid was obtained as a tan solid (149 mg, 0.69 mmol, 86%).

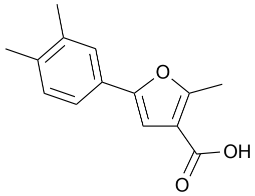

**5-(3,4-dimethylphenyl)-2-methylfuran-3-carboxylic acid (S35)**
Following general procedure E, starting from methyl 5-(3,4-dimethylphenyl)-2-methylfuran-3-carboxylate (**S24**) (72 mg, 0.30 mmol, 1.0 eq) and LiOH (35 mg, 1.47 mmol, 5.0 eq), 5-(3,4-dimethylphenyl)-2-methylfuran-3-carboxylic acid was obtained as a white powder (40 mg, 0.18 mmol, 59%).

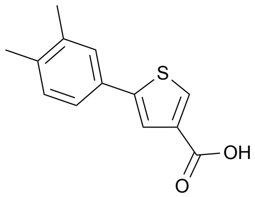

**5-(3,4-dimethylphenyl)thiophene-3-carboxylic acid (S36)**
Following general procedure E, starting from methyl 5-(3,4-dimethylphenyl)thiophene-3-carboxylate (**S25**) (73 mg, 0.30 mmol, 1.0 eq) and LiOH (36 mg, 1.48 mmol, 5.0 eq), 5-(3,4-dimethylphenyl)thiophene-3-carboxylic acid was obtained as a white solid (47 mg, 0.20 mmol, 69%).

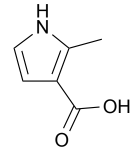

**2-methyl-1*H*-pyrrole-3-carboxylic acid (S37)**
Following general procedure E, starting from ethyl 2-methyl-1*H*-pyrrole-3-carboxylate (70 mg, 0.46 mmol, 1.0 eq) and LiOH (55 mg, 2.29 mmol, 5.0 eq), 2-methyl-1*H*-pyrrole-3-carboxylic acid was obtained as a yellow solid (50 mg, 0.40 mmol, 87%).

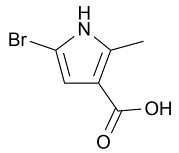

**5-bromo-2-methyl-1*H*-pyrrole-3-carboxylic acid (S38)**
Following general procedure E, starting from ethyl 5-bromo-2-methyl-1*H*-pyrrole-3-carboxylate (300 mg, 1.10 mmol, 1.0 eq) and NaOH (220 mg, 5.49 mmol, 5.0 eq), 5-bromo-2-methyl-1*H*-pyrrole-3-carboxylic acid was obtained as a brown solid (71 mg, 0.35 mmol, 32%).

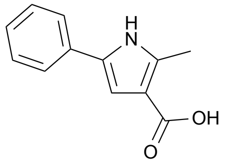

**2-methyl-5-phenyl-1*H*-pyrrole-3-carboxylic acid (S39)**
Following general procedure E, starting from ethyl 2-methyl-5-phenyl-1*H*-pyrrole-3-carboxylate (**S26**) (76 mg, 0.33 mmol, 1.0 eq) and LiOH (26 mg, 1.08 mmol, 5.0 eq), 2-methyl-5-phenyl-1*H*-pyrrole-3-carboxylic acid was obtained as a yellow solid (56 mg, 0.28 mmol, 84%).

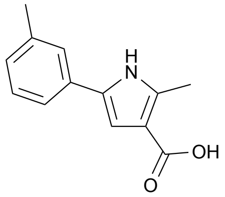

**2-methyl-5-(m-tolyl)-1*H*-pyrrole-3-carboxylic acid (S40)**
Following general procedure E, starting from ethyl 2-methyl-5-(m-tolyl)-1*H*-pyrrole-3-carboxylate (**S27**) (95 mg, 0.39 mmol, 1.0 eq) and LiOH (47 mg, 1.95 mmol, 5.0 eq), 2-methyl-5-(m-tolyl)-1*H*-pyrrole-3-carboxylic acid was obtained as an orange solid (38 mg, 0.17 mmol, 45%).

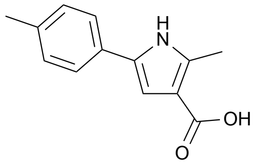

**2-methyl-5-(p-tolyl)-1*H*-pyrrole-3-carboxylic acid (S41)**
Following general procedure E, starting from ethyl 2-methyl-5-(p-tolyl)-1*H*-pyrrole-3-carboxylate (**S28**) (51 mg, 0.21 mmol, 1.0 eq) and LiOH (25 mg, 1.05 mmol, 5.0 eq), 2-methyl-5-(p-tolyl)-1*H*-pyrrole-3-carboxylic acid was obtained as a redish solid (23 mg, 0.11 mmol, 51%).

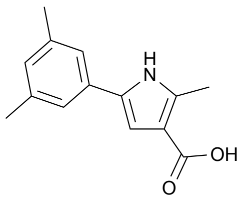

**5-(3,5-dimethylphenyl)-2-methyl-1*H*-pyrrole-3-carboxylic acid (S42)**
Following general procedure E, starting from ethyl 5-(3,5-dimethylphenyl)-2-methyl-1*H*-pyrrole-3-carboxylate (**S29**) (101 mg, 0.39 mmol, 1.0 eq) and LiOH (47 mg, 1.96 mmol, 5.0 eq), 5-(3,5-dimethylphenyl)-2-methyl-1*H*-pyrrole-3-carboxylic acid was obtained as an orange oil (44 mg, 0.19 mmol, 49%).

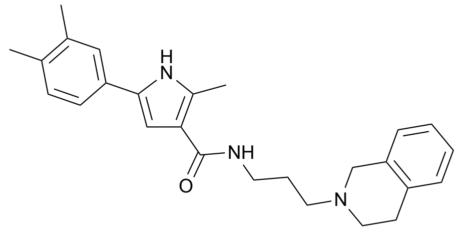

***N*-(3-(3,4-dihydroisoquinolin-2(1H)-yl)propyl)-5-(3,4-dimethylphenyl)-2-methyl-1*H*-pyrrole-3-carboxamide (Compound 5)**

Following general procedure F, starting from 5-(3,4-dimethylphenyl)-2-methyl-1*H*-pyrrole-3-carboxylic acid (**S30**) (50 mg, 0.22 mmol, 1.0 eq) and 3-(3,4-dihydroisoquinolin-2(1H)-yl)propan-1-amine (83 mg, 0.44 mmol, 2.0 eq) and purifying by chromatography on silica gel using 9:1:0.05 EtOAc:MeOH:$NEt_3$ as eluent, *N*-(3-(3,4-dihydroisoquinolin-2(1H)-yl)propyl)-5-(3,4-dimethylphenyl)-2-methyl-1*H*-pyrrole-3-carboxamide was obtained as a yellow solid (73 mg, 0.18 mmol, 84%). $^1$H NMR (400 MHz, $CDCl_3$) δ 7.09-7.01 (m, 5H), 7.00-6.92 (m, 2H), 6.82-6.80 (m, 1H), 6.24-6.21 (m, 1H), 3.43 (t, 2H, *J* = 6.3 Hz), 2.88 (q, 2H, *J* = 5.8 Hz), 2.73 (t, 2H, *J* = 6.2 Hz), 2.63 (t, 2H, *J* = 6.4 Hz), 2.47 (s, 3H), 2.20 (s, 3H), 2.18 (s, 3H), 1.82 (sext., 2H, *J* = 6.4 Hz); $^{13}$C NMR (101 MHz, $CDCl_3$) δ 167.1, 136.9, 134.7, 134.2, 134.0, 133.9, 130.3, 130.2, 130.0, 128.9, 126.8, 126.7, 126.1, 124.9, 121.5, 115.3, 103.1, 57.6, 51.2, 39.1, 29.0, 25.6, 19.9, 19.5, 12.8.

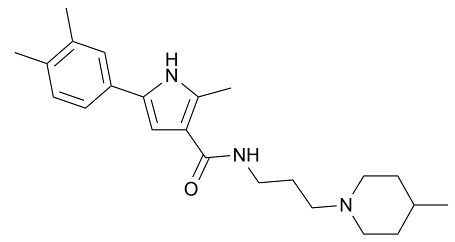

**5-(3,4-dimethylphenyl)-2-methyl-*N*-(3-(4-methylpiperidin-1-yl)propyl)-1*H*-pyrrole-3-carboxamide (Compound 6)**

Following general procedure F, starting from 5-(3,4-dimethylphenyl)-2-methyl-1*H*-pyrrole-3-carboxylic acid (**S30**) (38 mg, 0.17 mmol, 1.0 eq) and 3-(4-methylpiperidin-1-yl)propan-1-amine (**S8**) (52 mg, 0.33 mmol, 2.0 eq) and purifying by chromatography on silica gel using 9:1:0.05 EtOAc:MeOH:$NEt_3$ as eluent, 5-(3,4-dimethylphenyl)-2-methyl-*N*-(3-(4-methylpiperidin-1-yl)propyl)-1*H*-pyrrole-3-carboxamide was obtained as white solid (47 mg, 0.13 mmol, 77%). $^1$H NMR (400 MHz, $CDCl_3$) δ 8.63 (s, 1H), 7.56 (s, 1H), 7.25 (s, 1H), 7.20 (d, 1H, *J* = 7.6 Hz), 7.09 (d, 1H, *J* = 7.6 Hz), 6.55 (br s, 1H), 3.50 (t, 2H, *J* = 5.1 Hz), 3.02 (d, 2H, *J* = 10.5 Hz), 2.61 (s, 3H), 2.52 (q, 2H, *J* = 5.9 Hz), 2.26 (s, 3H), 2.45 (s, 3H), 1.95 (br s, 2H), 1.77 (q, 2H, *J* = 5.9 Hz), 1.67 (d, 2H, *J* = 9.1 Hz), 1.40-1.33 (m, 3H), 0.92 (d, 3H, *J* = 5.7 Hz); $^{13}$C NMR (101 MHz, $CDCl_3$) δ 166.3, 137.4, 135.2, 133.9, 130.4, 130.2, 130.1, 125.3, 121.5, 116.4, 103.9, 59.0, 54.6, 40.1, 34.6, 31.1, 25.3, 22.1, 20.2, 19.8, 13.5.

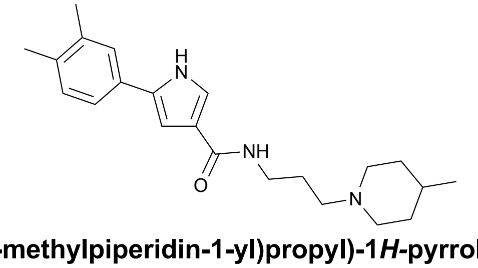

**5-(3,4-dimethylphenyl)-*N*-(3-(4-methylpiperidin-1-yl)propyl)-1*H*-pyrrole-3-carboxamide (Compound 7)**

Following general procedure F, starting from 5-(3,4-dimethylphenyl)-1*H*-pyrrole-3-carboxylic acid (**S31**) (49 mg, 0.23 mmol, 1.0 eq) and 3-(4-methylpiperidin-1-yl)propan-1-amine (**S8**) (71 mg, 0.46 mmol, 2.0 eq) and purifying by chromatography on silica gel using 9:1:0.05 EtOAc:MeOH:$NEt_3$ as eluent, 5-(3,4-dimethylphenyl)-*N*-(3-(4-methylpiperidin-1-yl)propyl)-1*H*-pyrrole-3-carboxamide was obtained as an orange solid (27 mg, 0.08 mmol, 33%). $^1$H NMR (400 MHz, DMSO-$d_6$) δ 11.54 (s, 1H), 7.84 (br s, 1H), 7.40 (br s, 1H), 7.31 (br s, 2H), 7.12 (d, 1H, *J* = 6.1 Hz), 6.80 (br s, 1H), 5.75 (br s, 1H), 3.03-2.87 (m, 4H), 2.37 (br s,

2H), 2.24 (s, 3H), 2.20 (s, 3H), 1.91 (q, 2H, *J* = 7.0 Hz), 1.65-1.57 (m, 7H), 1.14 (quint., 4H, *J* = 10.9 Hz), 0.88 (br s, 3H); $^{13}$C NMR (101 MHz, DMSO-$d_6$) δ 163.8, 136.4, 134.1, 131.8, 129.9, 129.8, 124.8, 121.1, 121.0, 104.0, 56.1, 55.7, 54.9, 53.3, 37.2, 37.0, 33.7, 30.2, 26.5, 26.4, 22.6, 21.8, 19.5, 19.0.

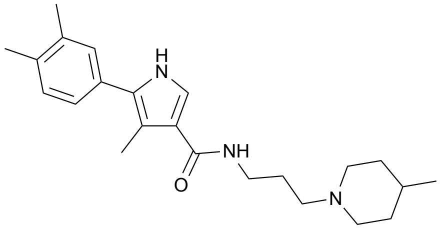


**5-(3,4-dimethylphenyl)-4-methyl-*N*-(3-(4-methylpiperidin-1-yl)propyl)-1*H*-pyrrole-3-carboxamide (Compound 8)**

Following general procedure F, starting from 5-(3,4-dimethylphenyl)-4-methyl-1*H*-pyrrole-3-carboxylic acid (**S32**) (50 mg, 0.22 mmol, 1.0 eq) and 3-(4-methylpiperidin-1-yl)propan-1-amine (**S8**) (68 mg, 0.44 mmol, 2.0 eq) and purifying by chromatography on silica gel using 9:1:0.05 EtOAc:MeOH:$NEt_3$ as eluent, 5-(3,4-dimethylphenyl)-4-methyl-*N*-(3-(4-methylpiperidin-1-yl)propyl)-1*H*-pyrrole-3-carboxamide was obtained as an orange solid (28 mg, 0.08 mmol, 35%). $^1$H NMR (400 MHz, $CDCl_3$) δ 8.74 (s, 1H), 7.17 (s, 1H), 7.15-7.13 (m, 2H), 7.12-7.08 (m, 1H), 3.47 (q, 2H, *J* = 6.4 Hz), 3.02 (d, 2H, *J* = 11.8 Hz), 2.56 (t, 2H, *J* = 6.6 Hz), 2.40 (s, 3H), 2.28 (s, 3H), 2.27 (s, 3H), 2.05 (t, 2H, *J* = 12.0 Hz), 1.82 (quint., 2H, *J* = 6.4 Hz), 1.69-1.64 (m, 2H), 1.42-1.38 (m, 1H), 1.31 (qd, 2H, *J* = 12.5 Hz, *J* = 3.2 Hz), 0.92 (d, 3H, *J* = 6.3 Hz); $^{13}$C NMR (101 MHz, $CDCl_3$) δ 166.7, 137.2, 135.6, 130.8, 130.7, 130.3, 128.9, 125.2, 120.7, 119.8, 115.3, 57.4, 54.2, 38.8, 34.6, 33.8, 25.8, 22.0, 20.2, 19.8, 11.7.

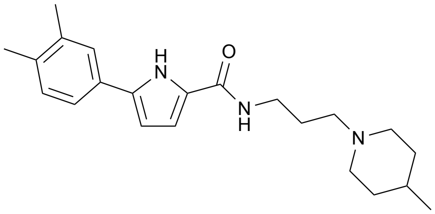


**5-(3,4-dimethylphenyl)-*N*-(3-(4-methylpiperidin-1-yl)propyl)-1*H*-pyrrole-2-carboxamide (Compound 10)**

Following general procedure F, starting from 5-(3,4-dimethylphenyl)-1*H*-pyrrole-2-carboxylic acid (**S33**) (80 mg, 0.37 mmol, 1.0 eq) and 3-(4-methylpiperidin-1-yl)propan-1-amine (**S8**) (116 mg, 0.74 mmol, 2.0 eq) and purifying by chromatography on silica gel using 9.5:0.5:0.05 EtOAc:MeOH:$NEt_3$ as eluent, 5-(3,4-dimethylphenyl)-*N*-(3-(4-methylpiperidin-1-yl)propyl)-1*H*-pyrrole-2-carboxamide was obtained as a pinkish solid (27 mg, 0.08 mmol, 20%). $^1$H NMR (400 MHz, $CDCl_3$) δ 9.78 (s, 1H), 8.09 (s, 1H), 7.36 (s, 1H), 7.32 (d, 1H, *J* = 7.9 Hz), 7.13 (d, 1H, *J* = 7.8 Hz), 6.70 (br s, 1H), 6.46 (br s, 1H), 3.53 (q, 2H, *J* = 5.6 Hz), 3.06 (d, 2H, *J* = 11.2 Hz), 2.67 (br s, 1H), 2.59 (t, 2H, *J* = 5.8 Hz), 2.29 (s, 3H), 2.26 (s, 3H), 2.04 (t, 2H, *J* = 11.0 Hz), 1.81 (quint., 2H, *J* = 5.6 Hz), 1.71 (d, 2H, *J* = 10.9 Hz), 1.42 (d, 2H, *J* = 7.5 Hz), 0.99 (d, 3H, *J* = 5.4 Hz); $^{13}$C NMR (101 MHz, $CDCl_3$) δ 161.6, 137.4, 136.1, 135.6, 130.5, 129.8, 126.9, 126.2, 122.3, 111.2, 106.9, 58.3, 54.4, 39.9, 34.1, 30.8, 24.9, 22.1, 20.2, 19.8.

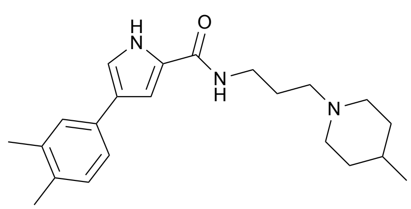


**4-(3,4-dimethylphenyl)-*N*-(3-(4-methylpiperidin-1-yl)propyl)-1*H*-pyrrole-2-carboxamide (Compound 11)**

Following general procedure F, starting from 4-(3,4-dimethylphenyl)-1*H*-pyrrole-2-carboxylic acid (**S34**) (148 mg, 0.69 mmol, 1.0 eq) and 3-(4-methylpiperidin-1-yl)propan-1-amine (**S8**) (215 mg, 1.38 mmol, 2.0

eq) and purifying by chromatography on silica gel using 9.5:0.5:0.05 EtOAc:MeOH:$NEt_3$ as eluent, 4-(3,4-dimethylphenyl)-*N*-(3-(4-methylpiperidin-1-yl)propyl)-1*H*-pyrrole-2-carboxamide was obtained as a yellow solid (72 mg, 0.20 mmol, 30%). $^1$H NMR (400 MHz, $CDCl_3$) δ 10.06 (s, 1H), 8.32 (s, 1H), 7.31-7.28 (m, 2H), 7.17 (s, 1H), 7.11 (d, 1H, *J* = 7.8 Hz), 6.90 (s, 1H), 3.57-3.54 (m, 2H), 3.08-3.01 (m, 4H), 2.56 (q, 2H, *J* = 5.3 Hz), 2.29 (s, 3H), 2.27 (s, 3H), 1.99-1.97 (m, 2H), 1.80-1.73 (m, 5H), 1.01 (d, 3H, *J* = 4.9 Hz); $^{13}$C NMR (101 MHz, $CDCl_3$) δ 161.5, 137.0, 134.5, 133.0, 130.2, 127.5, 126.8, 126.4, 123.0, 117.8, 106.6, 59.3, 54.7, 40.7, 34.8, 31.1, 24.6, 22.2, 20.2, 19.7.

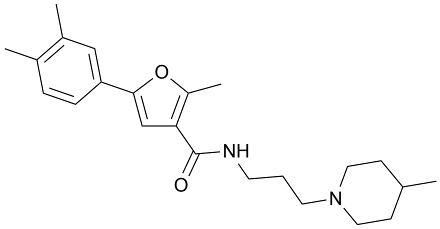

**5-(3,4-dimethylphenyl)-2-methyl-*N*-(3-(4-methylpiperidin-1-yl)propyl)furan-3-carboxamide (Compound 12)**

Following general procedure F, starting from 5-(3,4-dimethylphenyl)-2-methylfuran-3-carboxylic acid (**S35**) (40 mg, 0.17 mmol, 1.0 eq) and 3-(4-methylpiperidin-1-yl)propan-1-amine (**S8**) (54 mg, 0.35 mmol, 2.0 eq) and purifying by chromatography on silica gel using 10:0.05 EtOAc:$NEt_3$ as eluent, 5-(3,4-dimethylphenyl)-2-methyl-*N*-(3-(4-methylpiperidin-1-yl)propyl)furan-3-carboxamide was obtained as a yellow oil (54 mg, 0.15 mmol, 84%). $^1$H NMR (400 MHz, $CDCl_3$) δ 7.96 (br s, 1H), 7.42 (br s, 1H), 7.38 (dd, 1H, *J* = 7.9 Hz, *J* = 1.7 Hz), 7.12 (d, 1H, *J* = 7.9 Hz), 6.77 (s, 1H), 3.52 (q, 2H, *J* = 6.2 Hz), 3.15 (d, 2H, *J* = 10.5 Hz), 2.68-2.65 (m, 5H), 2.28 (s, 3H), 2.26 (s, 3H), 2.15-2.10 (m, 2H), 1.87 (quint., 2H, *J* = 5.6 Hz), 1.74 (d, 2H, *J* = 9.6 Hz), 1.51-1.45 (m, 3H), 0.97 (d, 3H, *J* = 5.4 Hz); $^{13}$C NMR (101 MHz, $CDCl_3$) δ 164.5, 156.2, 152.1, 137.2, 136.5, 130.3, 128.3, 125.1, 121.6, 117.7, 103.1, 54.3, 53.8, 39.4, 33.8, 30.7, 24.5, 21.6, 20.1, 19.9, 14.0.

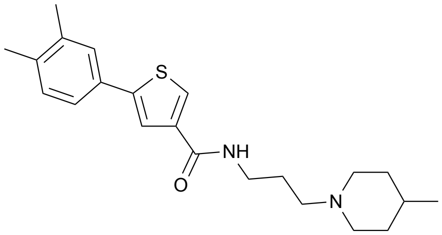

**5-(3,4-dimethylphenyl)-*N*-(3-(4-methylpiperidin-1-yl)propyl)thiophene-3-carboxamide (Compound 13)**

Following general procedure F, starting from 5-(3,4-dimethylphenyl)thiophene-3-carboxylic acid (**S36**) (47 mg, 0.20 mmol, 1.0 eq) and 3-(4-methylpiperidin-1-yl)propan-1-amine (**S8**) (63 mg, 0.41 mmol, 2.0 eq) and purifying by chromatography on silica gel using 9.5:0.5:0.05 EtOAc:MeOH:$NEt_3$ as eluent, 5-(3,4-dimethylphenyl)-*N*-(3-(4-methylpiperidin-1-yl)propyl)thiophene-3-carboxamide was obtained as a white solid (60 mg, 0.16 mmol, 80%). $^1$H NMR (400 MHz, DMSO-$d_6$) δ 8.30 (s, 1H), 7.99 (s, 1H), 7.78 (s, 1H), 7.43 (s, 1H), 7.35 (d, 1H, *J* = 6.1 Hz), 7.18 (d, 1H, *J* = 6.1 Hz), 3.41 (br s, 2H), 3.25 (br s, 2H), 2.82 (d, 2H, *J* = 8.1 Hz), 2.26 (s, 3H), 2.23 (s, 3H), 1.82 (t, 2H, *J* = 9.3 Hz), 1.66 (br s, 2H), 1.55 (d, 2H, *J* = 10.2 Hz), 1.29 (br s, 1H), 1.10 (t, 2H, *J* = 10.9 Hz), 0.85 (d, 3H, *J* = 4.8 Hz); $^{13}$C NMR (101 MHz, DMSO-$d_6$) δ 143.9, 138.8, 137.1, 136.3, 130.8, 130.2, 129.1, 126.3, 122.8, 122.0, 56.1, 53.5, 37.7, 34.0, 30.4, 26.5, 21.8, 29.3, 19.1.

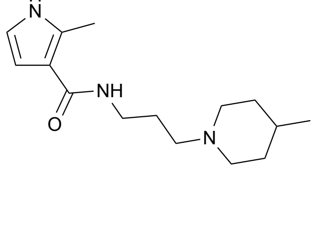

**2-methyl-*N*-(3-(4-methylpiperidin-1-yl)propyl)-1*H*-pyrrole-3-carboxamide (Compound 20)**
Following general procedure F, starting from 2-methyl-1*H*-pyrrole-3-carboxylic acid (**S37**) (50 mg, 0.40 mmol, 1.0 eq) and 3-(4-methylpiperidin-1-yl)propan-1-amine (**S8**) (124 mg, 0.79 mmol, 2.0 eq) and purifying by chromatography on silica gel using 9:1:0.05 EtOAc:MeOH:$NEt_3$ as eluent, 2-methyl-*N*-(3-(4-methylpiperidin-1-yl)propyl)-1*H*-pyrrole-3-carboxamide was obtained as a colorless oil (50 mg, 0.19 mmol, 48%). $^1$H NMR (400 MHz, $CDCl_3$) δ 8.17 (br s, 1H), 7.35 (s, 1H), 6.56 (t, 1H, *J* = 2.9 Hz), 6.35 (t, 1H, *J* = 2.9 Hz), 3.48 (q, 2H, *J* = 6.4 Hz), 3.00 (d, 2H, *J* = 11.2 Hz), 2.56 (s, 3H), 2.52 (t, 2H, *J* = 6.3 Hz), 1.97 (br s, 2H), 1.78 (t, 2H, *J* = 6.1 Hz), 1.66 (d, 2H, *J* = 11.8 Hz), 1.42-1.30 (m, 3H), 0.94 (d, 3H, *J* = 6.1 Hz); $^{13}$C NMR (101 MHz, DMSO) δ 165.13, 131.55, 115.12, 114.18, 106.94, 56.31, 53.44, 37.17, 33.89, 30.32, 26.71, 21.83, 12.55.

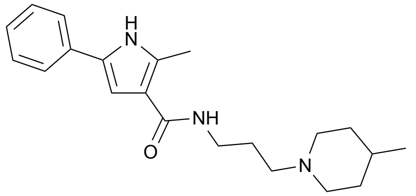


**2-methyl-*N*-(3-(4-methylpiperidin-1-yl)propyl)-5-phenyl-1*H*-pyrrole-3-carboxamide (Compound 22)**
Following general procedure F, starting from 2-methyl-5-phenyl-1*H*-pyrrole-3-carboxylic acid (**S39**) (59 mg, 0.29 mmol, 1.0 eq) and 3-(4-methylpiperidin-1-yl)propan-1-amine (**S8**) (92 mg, 0.59 mmol, 2.0 eq) and purifying by chromatography on silica gel using 9:1:0.05 EtOAc:MeOH:$NEt_3$ as eluent, 2-methyl-*N*-(3-(4-methylpiperidin-1-yl)propyl)-5-phenyl-1*H*-pyrrole-3-carboxamide was obtained as an orange solid (40 mg, 0.12 mmol, 40%). $^1$H NMR (400 MHz, DMSO-$d_6$) δ 11.32 (s, 1H), 7.70 (br s, 1H), 7.57 (d, 2H, *J* = 7.2 Hz), 7.35 (t, 2H, *J* = 7.2 Hz), 7.16 (t, 1H, *J* = 7.2 Hz), 6.83 (s, 1H), 3.20 (q, 2H, *J* = 5.6 Hz), 2.88 (d, 2H, *J* = 9.0 Hz), 2.47 (s, 3H), 2.36 (t, 2H, *J* = 5.9 Hz), 1.90 (br s, 2H), 1.65 (t, 2H, *J* = 6.2 Hz), 1.59 (d, 2H, *J* = 13.2 Hz), 1.33 (br s, 1H), 1.15 (q, 2H, *J* = 11.1 Hz), 0.87 (d, 3H, *J* = 6.1 Hz); $^{13}$C NMR (101 MHz, DMSO-$d_6$) δ 164.8, 133.6, 132.3, 128.7, 128.6, 125.6, 123.1, 115.7, 104.5, 56.4, 53.4, 37.3, 33.8, 30.2, 26.4, 21.7, 12.6.

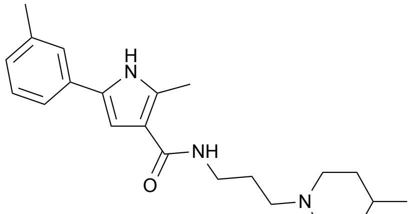


**2-methyl-*N*-(3-(4-methylpiperidin-1-yl)propyl)-5-(m-tolyl)-1*H*-pyrrole-3-carboxamide (Compound 23)**
Following general procedure F, starting from 2-methyl-5-(m-tolyl)-1*H*-pyrrole-3-carboxylic acid (**S40**) (37 mg, 0.17 mmol, 1.0 eq) and 3-(4-methylpiperidin-1-yl)propan-1-amine (**S8**) (54 mg, 0.34 mmol, 2.0 eq) and purifying by chromatography on silica gel using 9:1:0.05 EtOAc:MeOH:$NEt_3$ as eluent, 2-methyl-*N*-(3-(4-methylpiperidin-1-yl)propyl)-5-(m-tolyl)-1*H*-pyrrole-3-carboxamide was obtained as a yellow solid (22 mg, 0.06 mmol, 37%). $^1$H NMR (400 MHz, $CDCl_3$) δ 8.70 (s, 1H), 7.59 (br s, 1H), 7.27 (d, 2H, *J* = 7.3 Hz), 7.22 (t, 1H, *J* = 7.3 Hz), 7.00 (d, 1H, *J* = 7.3 Hz), 6.61 (d, 1H, *J* = 2.5 Hz), 3.48 (t, 2H, *J* = 5.7 Hz), 3.03 (d, 2H, *J* = 11.2 Hz), 2.61 (s, 3H), 2.54 (t, 2H, *J* = 5.7 Hz), 2.34 (s, 3H), 1.98 (t, 2H, *J* = 10.5 Hz), 1.78 (quint., 2H, *J* = 5.9 Hz), 1.67 (d, 2H, *J* = 9.9 Hz), 1.41-1.35 (m, 3H), 0.91 (d, 3H, *J* = 5.4 Hz); $^{13}$C NMR (101 MHz, $CDCl_3$) δ 166.3, 138.8, 134.3, 132.3, 130.1, 129.1, 127.5, 124.7, 121.2, 116.5, 104.5, 58.8, 54.5, 39.9, 34.4, 31.0, 25.2, 22.0, 21.8, 13.5.

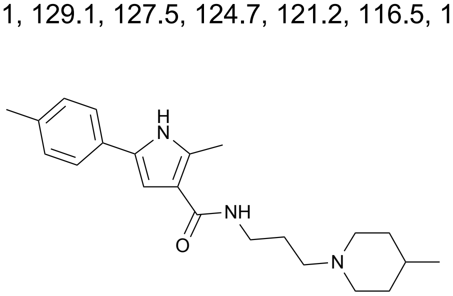


**2-methyl-*N*-(3-(4-methylpiperidin-1-yl)propyl)-5-(p-tolyl)-1*H*-pyrrole-3-carboxamide (Compound 24)**

Following general procedure F, starting from 2-methyl-5-(p-tolyl)-1*H*-pyrrole-3-carboxylic acid (**S41**) (23 mg, 0.11 mmol, 1.0 eq) and 3-(4-methylpiperidin-1-yl)propan-1-amine (**S8**) (33 mg, 0.21 mmol, 2.0 eq) and purifying by chromatography on silica gel using 9:1:0.05 EtOAc:MeOH:$NEt_3$ as eluent, 2-methyl-*N*-(3-(4-methylpiperidin-1-yl)propyl)-5-(p-tolyl)-1*H*-pyrrole-3-carboxamide was obtained as a white solid (25 mg, 0.07 mmol, 66%). $^1$H NMR (400 MHz, $CDCl_3$) δ 8.83 (s, 1H), 7.57 (br s, 1H), 7.37 (d, 1H, *J* = 5.6 Hz), 7.14 (d, 1H, *J* = 5.6 Hz), 6.59 (s, 1H), 3.48 (br s, 2H), 3.04 (d, 2H, *J* = 9.4 Hz), 2.61 (s, 3H), 2.54 (br s, 2H), 2.33 (s, 3H), 2.00 (br s, 2H), 1.78 (br s, 2H), 1.68 (d, 2H, *J* = 6.9 Hz), 1.40 (br s, 3H), 0.94 (br s, 3H); $^{13}$C NMR (101 MHz, $CDCl_3$) δ 166.3, 136.4, 134.1, 130.2, 129.8, 129.7, 124.0, 116.3, 104.0, 58.7, 54.5, 39.9, 34.3, 31.0, 25.2, 21.9, 21.4, 13.5.

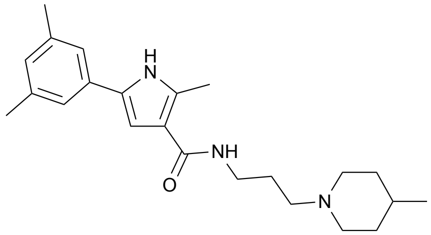


**5-(3,5-dimethylphenyl)-2-methyl-*N*-(3-(4-methylpiperidin-1-yl)propyl)-1*H*-pyrrole-3-carboxamide (Compound 25)**

Following general procedure F, starting from 5-(3,5-dimethylphenyl)-2-methyl-1*H*-pyrrole-3-carboxylic acid (**S42**) (43 mg, 0.19 mmol, 1.0 eq) and 3-(4-methylpiperidin-1-yl)propan-1-amine (**S8**) (59 mg, 0.38 mmol, 2.0 eq) and purifying by chromatography on silica gel using 9:1:0.05 EtOAc:MeOH:$NEt_3$ as eluent, 5-(3,5-dimethylphenyl)-2-methyl-*N*-(3-(4-methylpiperidin-1-yl)propyl)-1*H*-pyrrole-3-carboxamide was obtained as a white solid (26 mg, 0.07 mmol, 38%). $^1$H NMR (400 MHz, $CDCl_3$) δ 8.59 (s, 1H), 7.59 (s, 1H), 7.09 (s, 2H), 6.85 (s, 1H), 6.57 (s, 1H), 3.49 (q, 2H, *J* = 6.2 Hz), 3.02 (d, 2H, *J* = 11.1 Hz), 2.62 (s, 3H), 2.53 (t, 2H, *J* = 6.3 Hz), 2.31 (s, 6H), 1.96 (t, 2H, *J* = 10.6 Hz), 1.77 (t, 2H, *J* = 5.7 Hz), 1.68 (d, 2H, *J* = 10.1 Hz), 1.39 (d, 2H, *J* = 7.3 Hz), 1.26 (br s, 1H), 0.91 (d, 3H, *J* = 5.7 Hz); $^{13}$C NMR (101 MHz, $CDCl_3$) δ 166.2, 138.7, 134.1, 132.3, 130.2, 128.5, 121.9, 116.4, 104.4, 59.0, 54.6, 40.1, 34.6, 31.1, 25.3, 22.1, 21.7, 13.5.

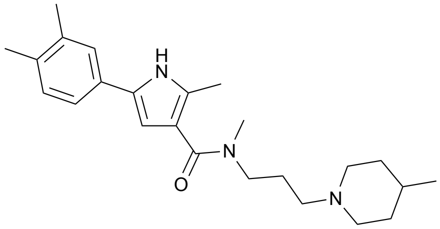


**5-(3,4-dimethylphenyl)-N,2-dimethyl-*N*-(3-(4-methylpiperidin-1-yl)propyl)-1*H*-pyrrole-3-carboxamide (Compound 27)**

Following general procedure F, starting from 5-(3,4-dimethylphenyl)-2-methyl-1*H*-pyrrole-3-carboxylic acid (**S30**) (48 mg, 0.21 mmol, 1.0 eq) and *N*-methyl-3-(4-methylpiperidin-1-yl)propan-1-amine (71 mg, 0.42 mmol, 2.0 eq) and purifying by chromatography on silica gel using 9:1:0.05 EtOAc:MeOH:$NEt_3$ as eluent, 5-(3,4-dimethylphenyl)-N,2-dimethyl-*N*-(3-(4-methylpiperidin-1-yl)propyl)-1*H*-pyrrole-3-carboxamide was obtained as white solid (58 mg, 0.15 mmol, 72%). $^1$H NMR (400 MHz, $CDCl_3$) δ 8.98 (s, 1H), 7.22 (s, 1H), 7.17 (d, 1H, *J* = 6.7 Hz), 7.07 (d, 1H, *J* = 7.2 Hz), 6.35 (s, 1H), 3.52 (t, 2H, *J* = 6.7 Hz), 3.08 (s, 3H), 2.85 (br s, 2H), 2.29 (s, 3H), 2.25 (s, 3H), 2.23 (s, 3H), 1.88-1.82 (m, 4H), 1.58 (d, 2H, *J* = 9.6 Hz), 1.32 (br s, 2H), 1.22 (t, 2H, *J* = 12.8 Hz=, 0.95-0.87 (m, 1H), 0.88 (d, 3H, *J* = 4.1 Hz); $^{13}$C NMR (101 MHz, $CDCl_3$) δ 169.2, 137.2, 134.8, 131.2, 130.4, 130.3, 130.2, 125.4, 121.5, 116.8, 105.3, 56.4, 54.4, 49.6, 46.2, 34.6, 31.1, 25.6, 22.2, 20.2, 19.7, 12.7.

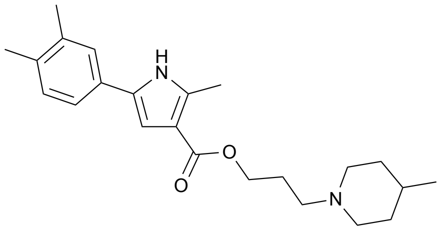

**3-(4-methylpiperidin-1-yl)propyl)-5-(3,4-dimethylphenyl)-2-methyl-1*H*-pyrrole-3-carboxylate (Compound 28)**

Following general procedure F, starting from 5-(3,4-dimethylphenyl)-2-methyl-1*H*-pyrrole-3-carboxylic acid (**S30**) (48 mg, 0.21 mmol, 1.0 eq) and 3-(4-methylpiperidin-1-yl)propan-1-ol (69 mg, 0.42 mmol, 2.0 eq) and purifying by chromatography on silica gel using 9:1:0.05 EtOAc:MeOH:$NEt_3$ as eluent, 3-(4-methylpiperidin-1-yl)propyl)-5-(3,4-dimethylphenyl)-2-methyl-1*H*-pyrrole-3-carboxylate was obtained as yellow solid (45 mg, 0.12 mmol, 58%). $^1$H NMR (400 MHz, $CDCl_3$) δ 8.44 (s, 1H), 7.24 (s, 1H), 7.18 (d, 1H, *J* = 7.8 Hz), 7.12 (d, 1H, *J* = 7.7 Hz), 6.76 (s, 1H), 4.26 (t, 2H, *J* = 6.3 Hz), 2.90 (d, 2H, *J* = 11.2 Hz), 2.57 (s, 3H), 2.47 (t, 2H, *J* = 7.4 Hz), 2.28 (s, 3H), 2.26 (s, 3H), 1.96-1.90 (m, 4H), 1.62 (d, 2H, *J* = 12.2 Hz), 1.36-1.31 (m, 1H), 1.26 (t, 2H, *J* = 9.4 Hz), 0.92 (d, 3H, *J* = 6.2 Hz); $^{13}$C NMR (101 MHz, $CDCl_3$) δ 165.9, 137.5, 136.0, 135.5, 130.5, 130.4, 129.8, 125.5, 121.5, 113.5, 107.1, 62.6, 56.2, 54.4, 34.7, 31.2, 27.0, 22.2, 20.2, 19.8, 13.8.

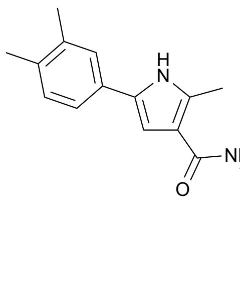

**5-(3,4-dimethylphenyl)-2-methyl-*N*-(2-(4-methylpiperidin-1-yl)ethyl)-1*H*-pyrrole-3-carboxamide (Compound 29)**

Following general procedure F, starting from 5-(3,4-dimethylphenyl)-2-methyl-1*H*-pyrrole-3-carboxylic acid (**S30**) (74 mg, 0.32 mmol, 1.0 eq) and 2-(4-methylpiperidin-1-yl)ethan-1-amine (92 mg, 0.65 mmol, 2.0 eq) and purifying by chromatography on silica gel using 10:0.05 EtOAc:$NEt_3$ as eluent, 5-(3,4-dimethylphenyl)-2-methyl-*N*-(2-(4-methylpiperidin-1-yl)ethyl)-1*H*-pyrrole-3-carboxamide was obtained as a white solid (61 mg, 0.17 mmol, 54%). $^1$H NMR (400 MHz, $CDCl_3$) δ 8.64 (s, 1H), 7.25 (s, 1H), 7.20 (d, 1H, *J* = 7.8 Hz), 7.11 (d, 1H, *J* = 7.8 Hz), 6.72 (br s, 1H), 6.58 (s, 1H), 3.53 (q, 2H, *J* = 5.5 Hz), 2.98 (d, 2H, *J* = 11.0 Hz), 2.62 (t, 2H, *J* = 5.7 Hz), 2.60 (s, 3H), 2.27 (s, 3H), 2.25 (s, 3H), 2.11 (t, 2H, *J* = 11.1 Hz), 1.66 (d, 2H, *J* = 12.1 Hz), 1.43-1.39 (m, 1H), 1.35 (t, 2H, *J* = 11.9 Hz), 0.94 (d, 3H, *J* = 5.9 Hz); $^{13}$C NMR (101 MHz, DMSO) δ 164.84, 136.31, 133.53, 132.99, 130.00, 129.81, 128.78, 124.36, 120.62, 115.51, 103.90, 57.78, 53.52, 36.19, 34.02, 30.34, 21.87, 19.56, 19.00, 12.58.

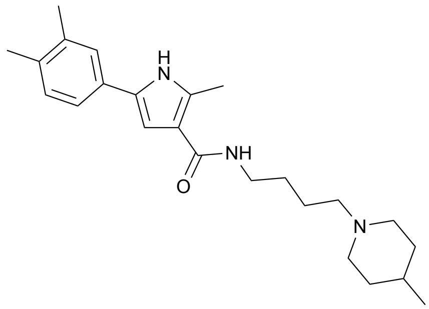

**5-(3,4-dimethylphenyl)-2-methyl-*N*-(4-(4-methylpiperidin-1-yl)butyl)-1*H*-pyrrole-3-carboxamide (Compound 30)**

Following general procedure F, starting from 5-(3,4-dimethylphenyl)-2-methyl-1*H*-pyrrole-3-carboxylic acid (**S30**) (84 mg, 0.37 mmol, 1.0 eq) and 4-(4-methylpiperidin-1-yl)butan-1-amine (125 mg, 0.73 mmol, 2.0 eq) and purifying by chromatography on silica gel using 9:1:0.05 EtOAc:MeOH:$NEt_3$ as eluent, 5-(3,4-dimethylphenyl)-2-methyl-*N*-(4-(4-methylpiperidin-1-yl)butyl)-1*H*-pyrrole-3-carboxamide was obtained as a yellow solid (88 mg, 0.23 mmol, 63%). $^1$H NMR (400 MHz, $CDCl_3$) δ 8.93 (s, 1H), 7.20 (d, 1H, *J* = 5.4 Hz), 7.10 (d, 1H, *J* = 5.4 Hz), 6.47 (s, 1H), 6.08 (s, 1H), 3.39 (br s, 2H), 2.90-2.87 (m, 2H), 2.58 (s, 3H), 2.33 (br s, 3H), 2.25 (s, 6H), 1.90-1.88 (m, 2H), 1.64-1.58 (m, 7H), 1.34-1.26 (m, 4H); $^{13}$C NMR (101 MHz, $CDCl_3$) δ 166.3, 137.3, 135.2, 133.8, 130.4, 130.0, 125.4, 121.5, 116.2, 103.6, 58.8, 54.3, 39.4, 34.5, 31.1, 28.2, 24.8, 22.2, 20.2, 19.7, 13.4. (1 peak missing, probably overlap)

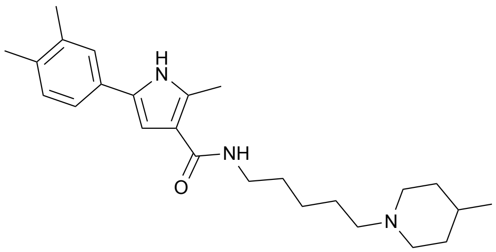


**5-(3,4-dimethylphenyl)-2-methyl-*N*-(4-(4-methylpiperidin-1-yl)pentyl)-1*H*-pyrrole-3-carboxamide (Compound 31)**

Following general procedure F, starting from 5-(3,4-dimethylphenyl)-2-methyl-1*H*-pyrrole-3-carboxylic acid (**S30**) (48 mg, 0.21 mmol, 1.0 eq) and 5-(4-methylpiperidin-1-yl)pentan-1-amine (**S9**) (77 mg, 0.42 mmol, 2.0 eq) and purifying by chromatography on silica gel using 9:1:0.05 EtOAc:MeOH:$NEt_3$ as eluent, 5-(3,4-dimethylphenyl)-2-methyl-*N*-(4-(4-methylpiperidin-1-yl)pentyl)-1*H*-pyrrole-3-carboxamide was obtained as a white solid (44 mg, 0.21 mmol, 53%). $^1$H NMR (400 MHz, $CDCl_3$) δ 8.70 (s, 1H), 7.24 (s, 1H), 7.18 (d, 1H, *J* = 7.5 Hz), 7.10 (d, 1H, *J* = 7.7 Hz), 6.45 (s, 1H), 5.83 (t, 1H, *J* = 6.1 Hz), 3.38 (q, 2H, *J* = 6.4 Hz), 2.88 (d, 2H, *J* = 10.7 Hz), 2.58 (s, 3H), 2.26 (s, 3H), 2.24 (s, 3H), 1.89 (t, 2H, *J* = 11.2 Hz), 1.61-1.49 (m, 7H), 1.35 (quint., 4H, *J* = 7.0 Hz), 1.29-1.20 (m, 4H), 0.90 (d, 3H, *J* = 5.9 Hz); $^{13}$C NMR (101 MHz, $CDCl_3$) δ 166.2, 137.4, 135.3, 133.8, 130.5, 129.9, 125.4, 121.5, 116.2, 103.5, 59.3, 54.3, 39.5, 34.5, 31.1, 30.3, 27.0, 25.5, 22.2, 20.2, 19.8, 13.4. (1 peak missing, probably overlap)

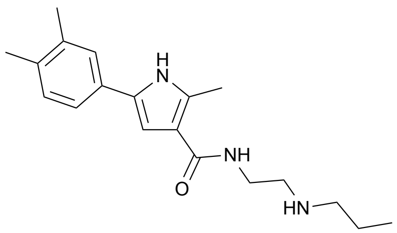


**5-(3,4-dimethylphenyl)-2-methyl-*N*-(2-(propylamino)ethyl)-1*H*-pyrrole-3-carboxamide (Compound 35)**

Following general procedure F, starting from 5-(3,4-dimethylphenyl)-2-methyl-1*H*-pyrrole-3-carboxylic acid (**S30**) (60 mg, 0.26 mmol, 1.0 eq) and *N1*-propylethane-1,2-diamine (64 µL, 0.52 mmol, 2.0 eq) and purifying by chromatography on silica gel using 8.5:1.5:0.05 EtOAc:MeOH:$NEt_3$ as eluent, 5-(3,4-dimethylphenyl)-2-methyl-*N*-(2-(propylamino)ethyl)-1*H*-pyrrole-3-carboxamide was obtained as a yellowish solid (47 mg, 0.15 mmol, 58%). $^1$H NMR (400 MHz, $CDCl_3$) δ 8.74 (s, 1H), 7.23 (s, 1H), 7.19 (d, 1H, *J* = 7.7 Hz), 7.08 (d, 1H, *J* = 7.7 Hz), 6.77 (s, 1H), 6.60 (s, 1H), 4.16 (br s, 1H), 3.54 (br s, 2H), 2.90 (br s, 2H), 2.66 (br s, 2H), 2.58 (s, 3H), 2.25 (s, 3H), 2.24 (s, 3H), 1.60 (br s, 2H), 0.92 (br s, 3H); $^{13}$C NMR (101 MHz, $CDCl_3$) δ 166.8, 137.4, 135.3, 134.0, 130.5, 129.9, 125.4, 121.5, 115.9, 104.1, 51.3, 49.2, 38.5, 22.5, 20.2, 19.7, 13.5, 11.9. (1 peak missing, probably overlap)

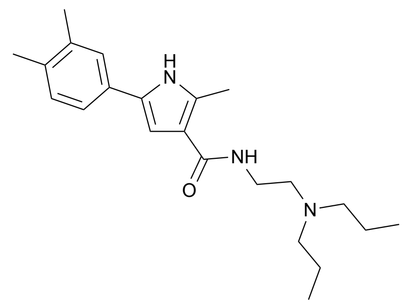

**5-(3,4-dimethylphenyl)-*N*-(3-(dipropylamino)propyl)-2-methyl-1*H*-pyrrole-3-carboxamide (Compound 36)**

Following general procedure F, starting from 5-(3,4-dimethylphenyl)-2-methyl-1*H*-pyrrole-3-carboxylic acid (**S30**) (61 mg, 0.27 mmol, 1.0 eq) and *N1,N1*-dipropylpropane-1,3-diamine (84 mg, 0.53 mmol, 2.0 eq) and purifying by chromatography on silica gel using 9:1:0.05 EtOAc:MeOH:$NEt_3$ as eluent, 5-(3,4-dimethylphenyl)-*N*-(3-(dipropylamino)propyl)-2-methyl-1*H*-pyrrole-3-carboxamide was obtained as a yellow solid (42 mg, 0.11 mmol, 43%).$^1$H NMR (400 MHz, $CDCl_3$) δ 8.80 (s, 1H), 7.22 (d, 1H, *J* = 1.7 Hz), 7.18 (dd, 1H, *J* = 7.8 Hz, *J* = 1.7 Hz), 7.08 (d, 1H, *J* = 7.9 Hz), 6.36 (s, 1H), 3.60 (t, 2H, *J* = 6.6 Hz), 3.44 (t, 2H, *J* = 7.7 Hz), 2.88 (br s, 2H), 2.59 (br s, 2H), 2.33 (s, 3H), 2.26 (s, 3H), 2.24 (s, 3H), 1.62 (sex., 2H, *J* = 7.4 Hz), 1.52 (br s, 2H), 0.94-0.86 (m, 6H); $^{13}$C NMR (101 MHz, CHLOROFORM-*D*) δ 166.84, 137.24, 135.06, 130.30, 130.19, 125.17, 122.42, 121.27, 104.91, 60.55, 48.24, 20.04, 19.58, 14.35, 12.64, 11.78, 11.31. (2 peaks missing, probably overlap)

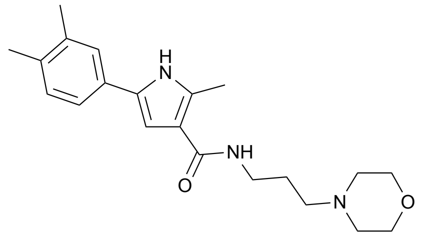

**5-(3,4-dimethylphenyl)-2-methyl-*N*-(3-morpholinopropyl)-1*H*-pyrrole-3-carboxamide (Compound 37)**

Following general procedure F, starting from 5-(3,4-dimethylphenyl)-2-methyl-1*H*-pyrrole-3-carboxylic acid (**S30**) (50 mg, 0.22 mmol, 1.0 eq) and 3-morpholinopropan-1-amine (63 mg, 0.44 mmol, 2.0 eq) and purifying by chromatography on silica gel using EtOAc:MeOH:$NEt_3$ 9.5:0.5:0.05, 5-(3,4-dimethylphenyl)-2-methyl-*N*-(3-morpholinopropyl)-1*H*-pyrrole-3-carboxamide was obtained as a yellow solid (58 mg, 0.16 mmol, 74%). $^1$H NMR (400 MHz, $CDCl_3$) δ 8.89 (s, 1H), 7.25 (s, 1H), 7.19 (d, 1H, *J* = 7.8 Hz), 7.10 (d, 1H, *J* = 7.9 Hz), 7.06 (t, 1H, *J* = 4.9 Hz), 6.53 (d, 1H, *J* = 2.6 Hz), 3.78 (t, 2H, *J* = 4.7 Hz), 3.48 (q, 2H, *J* = 6.1 Hz), 2.59 (s, 3H), 2.54-2.50 (m, 6H), 2.26 (s, 3H), 2.24 (s, 3H), 1.77 (quint., 2H, *J* = 6.3 Hz); $^{13}$C NMR (101 MHz, $CDCl_3$) δ 166.3, 137.4, 135.3, 134.0, 130.5, 130.4, 130.0, 125.4, 121.4, 116.1, 103.7, 67.1, 58.4, 54.1, 39.3, 25.4, 20.2, 19.7, 13.4.

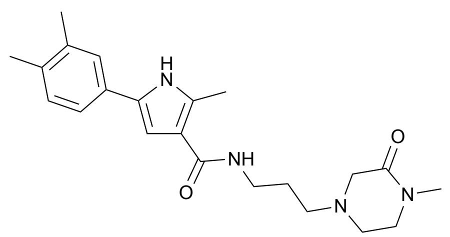

**5-(3,4-dimethylphenyl)-2-methyl-*N*-(3-(4-methyl-3-oxopiperazin-1-yl)propyl)-1*H*-pyrrole-3-carboxamide (Compound 39)**

Following general procedure F, starting from 5-(3,4-dimethylphenyl)-2-methyl-1*H*-pyrrole-3-carboxylic acid (**S30**) (41 mg, 0.18 mmol, 1.0 eq) and 4-(3-aminopropyl)-1-methylpiperazin-2-one (**S10**) (61 mg, 0.36 mmol, 2.0 eq) and purifying by chromatography on silica gel using EtOAc:MeOH:$NEt_3$ 9:1:0.05, 5-(3,4-dimethylphenyl)-2-methyl-*N*-(3-(4-methyl-3-oxopiperazin-1-yl)propyl)-1*H*-pyrrole-3-carboxamide was obtained as a yellow solid (26 mg, 0.07 mmol, 37%). $^1$H NMR (400 MHz, DMSO-$d_6$) δ 11.20 (s, 1H), 7.60 (t, 1H, *J* = 4.9 Hz), 7.36 (s, 1H), 7.28 (d, 1H, *J* = 7.7 Hz), 7.11 (d, 1H,*J* = 7.8 Hz), 6.76 (s, 1H), 3.26 (t, 2H, *J* = 6.1 Hz), 3.21 (sext., 2H, *J* = 6.1 Hz), 2.81 (s, 3H), 2.63 (t, 2H, *J* = 5.0 Hz), 2.44 (s, 3H), 2.38 (t, 2H, *J* = 6.9 Hz), 2.24 (s, 3H), 2.20 (s, 3H), 2.08 (s, 2H), 1.64 (t, 2H, *J* = 6.4 Hz); $^{13}$C NMR (101 MHz, DMSO-$d_6$) δ 166.0, 164.9, 136.3, 133.5, 133.0, 130.0, 129.8, 128.7, 124.4, 120.6, 115.5, 130.8, 57.1, 54.7, 49.2, 47.9, 36.7, 33.0, 30.7, 26.5, 19.5, 19.0.

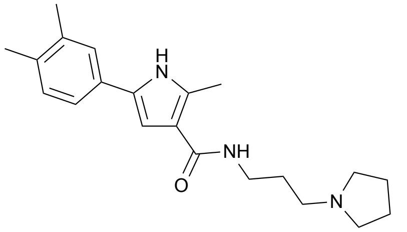

**5-(3,4-dimethylphenyl)-2-methyl-*N*-(3-(pyrrolidin-1-yl)propyl)-1*H*-pyrrole-3-carboxamide (Compound 40)**

Following general procedure F, starting from 5-(3,4-dimethylphenyl)-2-methyl-1*H*-pyrrole-3-carboxylic acid (**S30**) (59 mg, 0.26 mmol, 1.0 eq) and 3-(pyrrolidin-1-yl)propan-1-amine (65 µL, 0.52 mmol, 2.0 eq) and purifying by chromatography on silica gel using EtOAc:MeOH:$NEt_3$ 9:1:0.05, 5-(3,4-dimethylphenyl)-2-methyl-*N*-(3-(pyrrolidin-1-yl)propyl)-1*H*-pyrrole-3-carboxamide was obtained as a white solid (39 mg, 0.12 mmol, 45%). $^1$H NMR (400 MHz, DMSO-$d_6$) δ 11.23 (s, 1H), 7.71 (s, 1H), 7.36 (s, 1H), 7.28 (d, 1H, *J* = 7.9 Hz), 7.11 (d, 1H, *J* = 7.9 Hz), 6.76 (s, 1H), 3.22 (q, 2H, *J* = 5.8 Hz), 2.56 (br s, 6H), 2.45 (s, 3H), 2.23 (s, 3H), 2.20 (s, 3H), 1.75-1.67 (m, 6H); $^{13}$C NMR (101 MHz, DMSO-$d_6$) δ 164.9, 136.1, 133.5, 133.1, 130.0, 129.8, 128.8, 124.3, 120.6, 115.4, 103.7, 53.5, 37.0, 29.0, 28.1, 23.0, 19.5, 19.0, 12.6.

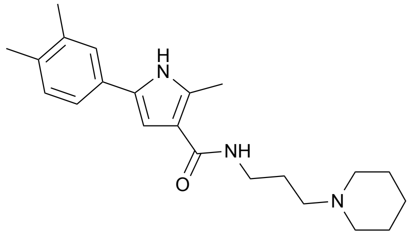

**5-(3,4-dimethylphenyl)-2-methyl-*N*-(3-(piperidin-1-yl)propyl)-1*H*-pyrrole-3-carboxamide (Compound 41)**

Following general procedure F, starting from 5-(3,4-dimethylphenyl)-2-methyl-1*H*-pyrrole-3-carboxylic acid (**S30**) (41 mg, 0.18 mmol, 1.0 eq) and 3-(piperidin-1-yl)propan-1-amine (57 µL, 0.36 mmol, 2.0 eq) and purifying by chromatography on silica gel using EtOAc:MeOH 9:1, 5-(3,4-dimethylphenyl)-2-methyl-*N*-(3-(piperidin-1-yl)propyl)-1*H*-pyrrole-3-carboxamide was obtained as a white solid (16 mg, 0.05 mmol, 26%). $^1$H NMR (400 MHz, DMSO-$d_6$) δ 11.30 (s, 1H), 7.82 (s, 1H), 7.40 (s, 1H), 7.33 (d, 1H, *J* = 7.8 Hz), 7.15 (d, 1H, *J* = 7.8 Hz), 6.84 (s, 1H), 3.28-3.20 (m, 4H), 2.73 (br s , 4H), 2.49 (s, 3H), 2.27 (s,3H), 2.24 (s, 3H), 1.81 (br s, 2H), 1.68 (br s, 2H), 1.51 (s, 2H); $^{13}$C NMR (151 MHz, DMSO) δ 165.58, 136.54, 133.85, 133.61, 129.99, 129.06, 124.48, 120.75, 115.17, 103.93, 52.89, 36.25, 29.12, 25.12, 23.70, 21.20, 19.70, 19.14, 12.73. (1 peak missing, probably overlap)

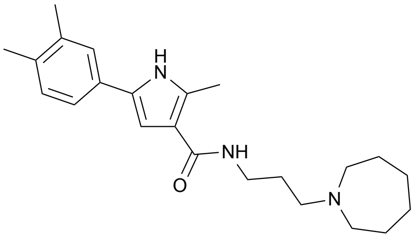

***N*-(3-(azepan-1-yl)propyl)-5-(3,4-dimethylphenyl)-2-methyl-1*H*-pyrrole-3-carboxamide (Compound 42)**

Following general procedure F, starting from 5-(3,4-dimethylphenyl)-2-methyl-1*H*-pyrrole-3-carboxylic acid (**S30**) (100 mg, 0.44 mmol, 1.0 eq) and 3-(azepan-1-yl)propan-1-amine (**S11**) (136 mg, 0.87 mmol, 2.0 eq) and purifying by chromatography on silica gel using 9:1:0.05 EtOAc:MeOH:$NEt_3$ as eluent, *N*-(3-(azepan-1-yl)propyl)-5-(3,4-dimethylphenyl)-2-methyl-1*H*-pyrrole-3-carboxamide was obtained as a white solid (114 mg, 0.31 mmol, 71%). $^1$H NMR (400 MHz, DMSO-$d_6$) δ 11.22 (s, 1H), 7.65 (s, 1H), 7.36 (s, 1H), 1.29 (d, 1H, *J* = 7.6 Hz), 7.11 (d, 1H, *J* = 7.6 Hz), 6.78 (s, 1H), 3.20 (q, 2H, *J* = 5.8 Hz), 2.68 (br s, 4H), 2.57 (br s, 2H), 2.45 (s, 3H), 2.23 (s, 3H), 2.20 (s, 3H), 1.62 (br s, 6H), 1.55 (br s, 4H); $^{13}$C NMR (101 MHz, DMSO-$d_6$) δ 165.0, 136.3, 133.5, 133.1, 130.0, 129.8, 128.7, 124.3, 120.6, 115.4, 103.8, 55.4, 54.7, 36.8, 27.1, 27.0, 26.4, 19.5, 19.0, 12.6.

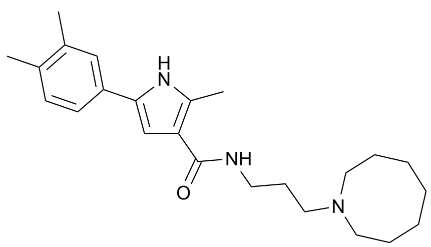

**_N_-(3-(azocan-1-yl)propyl)-5-(3,4-dimethylphenyl)-2-methyl-1_H_-pyrrole-3-carboxamide (Compound 43)**

Following general procedure F, starting from 5-(3,4-dimethylphenyl)-2-methyl-1*H*-pyrrole-3-carboxylic acid (**S30**) (35 mg, 0.15 mmol, 1.0 eq) and 3-(azocan-1-yl)propan-1-amine (**S12**) (52 mg, 0.31 mmol, 2.0 eq) and purifying by chromatography on silica gel using 9.5:0.5:0.05 EtOAc:MeOH:$NEt_3$ as eluent, *N*-(3-(azocan-1-yl)propyl)-5-(3,4-dimethylphenyl)-2-methyl-1*H*-pyrrole-3-carboxamide was obtained as a yellow solid (27 mg, 0.07 mmol, 47%). $^1$H NMR (400 MHz, DMSO-$d_6$) δ 11.19 (s, 1H), 7.54 (t, 1H, *J* = 5.6 Hz), 7.36 (s, 1H), 7.29 (dd, 1H, *J* = 7.8 Hz, J = 2.7 Hz), 7.11 (d, 1H, *J* = 7.8 Hz), 6.80 (d, 1H, *J* = 2.7 Hz), 3.32 (br s, 2H), 3.20 (q, 2H, *J* = 6.9 Hz), 2.45 (s, 3H), 2.42 (d, 2H, *J* = 7.1 Hz), 2.23 (s, 3H), 2.20 (s, 3H), 1.61 (quint., 4H, *J* = 7.0 Hz), 1.53 (br s, 10H); $^{13}$C NMR (101 MHz, DMSO-$d_6$) δ 164.9, 136.3, 133.5, 133.0, 130.0, 129.8, 129.7, 128.7, 124.3, 124.2, 120.6, 115.6, 103.9, 56.6, 53.8, 36.9, 28.4, 27.7, 27.1, 25.7, 19.5, 19.0, 12.6.

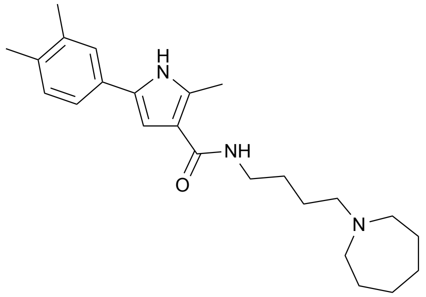

**_N_-(4-(azepan-1-yl)butyl)-5-(3,4-dimethylphenyl)-2-methyl-1_H_-pyrrole-3-carboxamide (Compound 44)**

Following general procedure F, starting from 5-(3,4-dimethylphenyl)-2-methyl-1*H*-pyrrole-3-carboxylic acid (**S30**) (50 mg, 0.22 mmol, 1.0 eq) and 4-(azepan-1-yl)butan-1-amine (**S13**) (74 mg, 0.44 mmol, 2.0 eq) and purifying by chromatography on silica gel using 9:1:0.05 EtOAc:MeOH:$NEt_3$ as eluent, *N*-(4-(azepan-1-yl)butyl)-5-(3,4-dimethylphenyl)-2-methyl-1*H*-pyrrole-3-carboxamide was obtained as a yellow solid (35 mg, 0.09 mmol, 43%). $^1$H NMR (400 MHz, $CDCl_3$) δ 8.82 (s, 1H), 7.25 (s, 1H), 7.20 (d, 1H, *J* = 7.8 Hz), 7.10 (d, 1H, *J* = 7.8 Hz), 6.51 (d, 1H, *J* = 2.6 Hz), 6.16 (t, 1H, *J* = 5.4 Hz), 3.39 (q, 2H, *J* = 5.9 Hz), 3.28 (br s, 1H), 2.72 (t, 4H, *J* = 5.6 Hz), 2.59 (s, 3H), 2.58-2.55 (m, 2H), 2.26 (s, 3H), 2.24 (s, 3H), 1.68 (br s 4H), 1.59 (br s, 8H); $^{13}$C NMR (101 MHz, $CDCl_3$) δ 166.3, 137.4, 135.3, 133.8, 130.5, 130.4, 130.0, 125.4, 121.5, 116.2, 103.7, 57.6, 55.5, 39.2, 28.0, 27.4, 27.1, 24.7, 20.2, 19.8, 13.4.

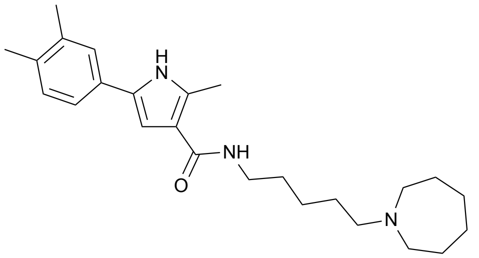

**_N_-(5-azepan-1-yl)pentyl)-5-(3,4-dimethylphenyl)-2-methyl-1_H_-pyrrole-3-carboxamide (Compound 45)**

Following general procedure F, starting from 5-(3,4-dimethylphenyl)-2-methyl-1*H*-pyrrole-3-carboxylic acid (**S30**) (48 mg, 0.21 mmol, 1.0 eq) and 5-(azepan-1-yl)pentan-1-amine (**S14**) (77 mg, 0.42 mmol, 2.0 eq) and purifying by chromatography on silica gel using 9:1:0.05 EtOAc:MeOH:$NEt_3$ as eluent, *N*-(5-azepan-1-yl)pentyl)-5-(3,4-dimethylphenyl)-2-methyl-1*H*-pyrrole-3-carboxamide was obtained as a white solid (48 mg, 0.12 mmol, 58%). $^1$H NMR (400 MHz, $CDCl_3$) δ 8.79 (s, 1H), 7.25 (s, 1H), 7.20 (d, 1H, *J* = 7.1 Hz), 7.10 (d, 1H, *J* = 7.5 Hz), 6.47 (s, 1H), 5.87 (s, 1H), 3.38 (q, 2H, *J* = 6.1 Hz), 2.66 (br s, 4H), 2.59 (s, 3H), 2.49 (t, 2H, *J* = 7.1 Hz), 2.26 (s, 3H), 2.24 (s, 3H), 1.66-1.57 (m, 12H), 1.37-1.34 (m, 2H); $^{13}$C NMR (101 MHz, $CDCl_3$) δ 166.2, 137.4, 135.3, 133.8, 130.5, 129.9, 125.4, 121.5, 116.2, 103.5, 58.3, 55.7, 39.5, 30.2, 27.6, 27.3, 27.1, 25.2, 20.2, 19.8, 13.4. (1 peak missing, probably overlap)

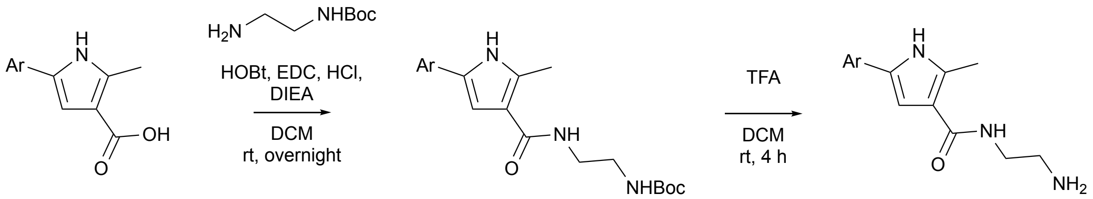


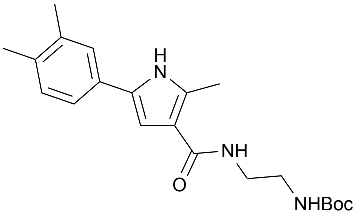

**_tert_-butyl (2-(5-(3,4-dimethylphenyl)-2-methyl-1_H_-pyrrole-3-carboxamido)ethyl)carbamate (S43)**

Following general procedure F, starting from 5-(3,4-dimethylphenyl)-2-methyl-1*H*-pyrrole-3-carboxylic acid (**S30**) (60 mg, 0.26 mmol, 1.0 eq) and *tert*-butyl (2-aminoethyl)carbamate (83 µL, 0.52 mmol, 2.0 eq) and purifying by chromatography on silica gel using 9.5:0.5:0.05 EtOAc:MeOH:$NEt_3$ as eluent, *tert*-butyl (2-(5-(3,4-dimethylphenyl)-2-methyl-1*H*-pyrrole-3-carboxamido)ethyl)carbamate was obtained as a yellowish solid (60 mg, 0.16 mmol, 62%). $^1$H NMR (400 MHz, $CDCl_3$) δ 8.30 (s, 1H), 7.22 (s, 1H), 7.16 (d, 1H, *J* = 7.8 Hz), 7.12 (d, 1H, *J* = 7.8 Hz), 6.52 (d, 1H, *J* = 1.9 Hz), 6.47 (br s, 1H), 3.51-3.49 (m, 2H), 3.37 (t, 2H, *J* = 5.7 Hz), 2.61 (s, 3H), 2.28 (s, 3H), 2.26 (s, 3H), 1.45 (s, 9H)

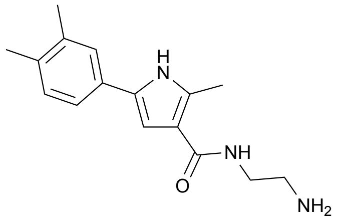

**_N_-(2-aminoethyl)-5-(3,4-dimethylphenyl)-2-methyl-1_H_-pyrrole-3-carboxamide (Compound 34)**

Following general procedure G, starting from tert-butyl (2-(5-(3,4-dimethylphenyl)-2-methyl-1*H*-pyrrole-3-carboxamido)ethyl)carbamate (**S43**) (59 mg, 0.16 mmol, 1.0 eq), *N*-(2-aminoethyl)-5-(3,4-dimethylphenyl)-2-methyl-1*H*-pyrrole-3-carboxamide was obtained as a white solid (34 mg, 0.12 mmol, 78%). $^1$H NMR (400

MHz, DMSO-$d_6$) δ 11.24 (s, 1H), 7.63 (t, 1H, *J* = 4.3 Hz), 7.37 (s, 1H), 7.29 (d, 1H, *J* = 7.8 Hz), 7.11 (d, 1H, *J* = 7.8 Hz), 6.83 (s, 1H), 3.67 (br s, 2H), 3.24 (q, 2H, *J* = 5.6 Hz), 2.72 (t, 2H, *J* = 5.8 Hz), 2.46 (s, 3H), 2.23 (s, 3H), 2.20 (s, 3H); $^{13}$C NMR (101 MHz, DMSO-$d_6$) δ 165.3, 136.3, 133.6, 133.3, 130.0, 129.8, 128.8, 124.4, 120.6, 115.3, 104.0, 41.0, 40.6, 19.6, 19.0, 12.6.

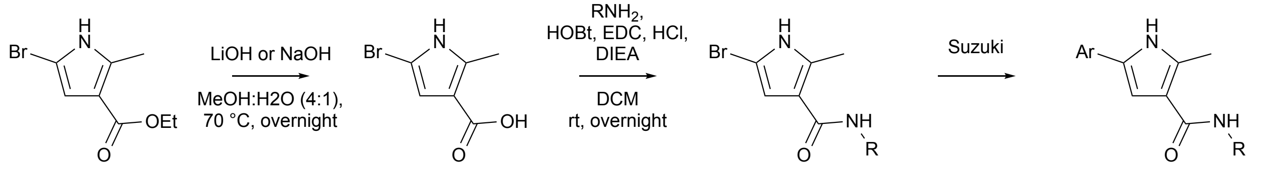


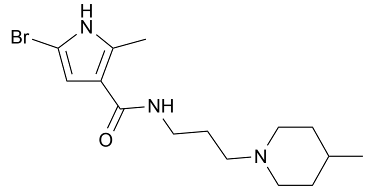


**5-bromo-2-methyl-*N*-(3-(4-methylpiperidin-1-yl)propyl)-1*H*-pyrrole-3-carboxamide (S44)**

Following general procedure F, starting from 5-bromo-2-methyl-1*H*-pyrrole-3-carboxylic acid (**S38**) (71 mg, 0.35 mmol, 1.0 eq) and 3-(4-methylpiperidin-1-yl)propan-1-amine (**S8**) (109 mg, 0.70 mmol, 2.0 eq) and purifying by chromatography on silica gel using 9:1:0.05 EtOAc:MeOH:$NEt_3$ as eluent, 5-bromo-2-methyl-*N*-(3-(4-methylpiperidin-1-yl)propyl)-1*H*-pyrrole-3-carboxamide was obtained as an orange solid (62 mg, 0.18 mmol, 52%). $^1$H NMR (400 MHz, $CDCl_3$) δ 9.32 (s, 1H), 7.76 (br s, 1H), 6.30 (s, 3H), 3.45 (q, 2H, *J* = 6.2 Hz), 2.99 (d, 2H, *J* = 11.8 Hz), 2.49 (s, 3H), 2.51-2.47 (m, 2H), 1.92 (t, 2H, *J* = 11.7 Hz), 1.73 (quint., 2H, *J* = 6.1 Hz), 1.65 (d, 2H, *J* = 12.9 Hz), 1.42-1.36 (m, 1H), 1.31 (td, 2H, *J* = 12.2 Hz, *J* = 3.3 Hz), 0.96 (d, 3H, *J* = 6.2 Hz); $^{13}$C NMR (101 MHz, $CDCl_3$) δ 165.5, 134.4, 116.6, 109.4, 96.1, 58.9, 54.5, 40.3, 34.4, 31.0, 24.9, 22.1, 13.1.

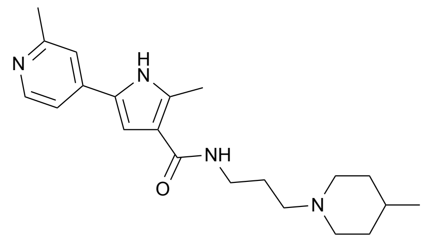


**2-methyl-*N*-(3-(4-methylpiperidin-1-yl)propyl)-5-(2-methylpyridin-4-yl)-1*H*-pyrrole-3-carboxamide (Compound 26)**

In an oven-dried round-bottom flask under argon, 5-bromo-2-methyl-*N*-(3-(4-methylpiperidin-1-yl)propyl)-1*H*-pyrrole-3-carboxamide (**S44**) (62 mg, 0.18 mmol, 1.0 eq), (2-methylpyridin-4-yl) boronic acid (30 mg, 0.22 mmol, 1.2 eq), $PdCl_2$(dppf) (0.05 eq), $Na_2CO_3$ (3.0 eq) were added and the vial was properly capped. The mixture vessel was evacuated and backfilled with argon (process repeated 3 times). A mixture of dioxane:$H_2O$ (4:1, 0.2 M) was added. The mixture vessel was evacuated and backfilled with argon (process repeated 3 times) and heated at 90 °C overnight. The reaction mixture was concentrated under vacuum. The residue was diluted with water and the aqueous phase was extracted 3 times with EtOAc. The combined organic layers were washed with brine, dried over $MgSO_4$ and concentrated under vacuum. The crude material was purified by chromatography on silica gel pretreated with a solution of $Et_2O$:$NEt_3$ 95:5 (AcOEt:MeOH:$NEt_3$ 9:1:0.05). 2-methyl-*N*-(3-(4-methylpiperidin-1-yl)propyl)-5-(2-methylpyridin-4-yl)-1*H*-pyrrole-3-carboxamide was obtained as a brown solid (16 mg, 0.045 mmol, 25%). $^1$H NMR (400 MHz, $CD_3OD$) δ 8.30 (d, 1H, *J* = 5.5 Hz), 7.45 (s, 1H), 7.37 (d, 1H, *J* = 5.4 Hz), 7.08 (s, 1H), 3.45-3.38 (m, 4H), 2.97 (t, 2H, *J* = 7.3 Hz), 2.74 (t, 2H, *J* = 11.3 Hz), 2.56 (s, 3H), 2.53 (s, 3H), 1.98 (quint., 2H, *J* = 7.0 Hz), 1.90 (d, 2H, *J* = 18.0 Hz), 1.67 (br s, 1H), 1.45 (q, 2H, *J* = 11.5 Hz), 1.01 (d, 3H, *J* = 6.3 Hz); $^{13}$C NMR (101 MHz, $CD_3OD$) δ 159.6, 149.7, 142.0, 138.5, 128.3, 118.4, 116.4, 109.0, 55.8, 53.9, 37.2, 33.0, 30.3, 26.4, 23.8, 21.4, 13.0. (2 peaks missing, probably overlap)

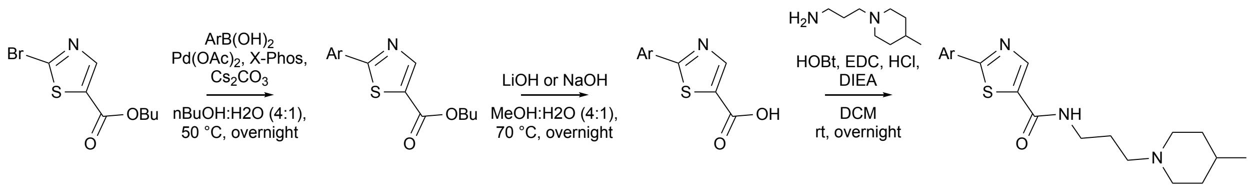


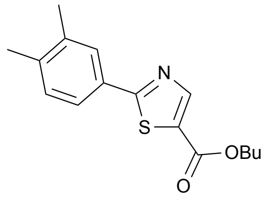

**butyl 2-(3,4-dimethylphenyl)thiazole-5-carboxylate (Compound 9) (S45)**

Following general procedure D, starting from methyl 2-bromothiazole-5-carboxylate (200 mg, 0.90 mmol, 1.0 eq) and (3,4-dimethylphenyl)boronic acid (162 mg, 1.08 mmol, 1.2 eq) and purifying by chromatography on silica gel using 8:1 Hex:EtOAc as eluent, butyl 2-(3,4-dimethylphenyl)thiazole-5-carboxylate was obtained as an orange oil (170 mg, 0.69 mmol, 76%). $^{1}H$ NMR (400 MHz, $CDCl_3$) δ 8.40 (s, 1H), 7.80 (s, 1H), 7.73 (d, 1H, *J* = 7.9 Hz), 7.23 (d, 1H, *J* = 7.9 Hz), 4.33 (t, 2H, *J* = 6.7 Hz), 2.34 (s, 3H), 2.32 (s, 3H), 1.82-1.71 (m, 2H), 1.47 (q, 2H, *J* = 7.4 Hz), 0.99 (t, 3H, *J* = 7.4 Hz)

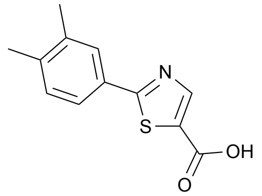

**2-(3,4-dimethylphenyl)thiazole-5-carboxylic acid (Compound 9) (S46)**

Following general procedure E, starting from butyl 2-(3,4-dimethylphenyl)thiazole-5-carboxylate (**S45**) (170 mg, 0.69 mmol, 1.0 eq) and LiOH (82 mg, 3.44 mmol, 5.0 eq), 2-(3,4-dimethylphenyl)thiazole-5-carboxylic acid was obtained as a yellow solid (104 mg, 0.45 mmol, 65%).

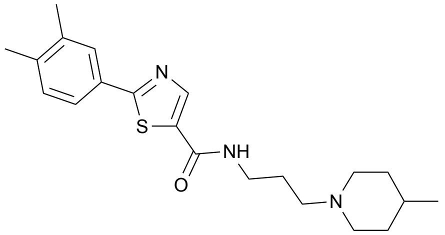

**2-(3,4-dimethylphenyl)-*N*-(3-(4-methylpiperidin-1-yl)propyl)thiazole-5-carboxamide (Compound 14)**

Following general procedure F, starting from 2-(3,4-dimethylphenyl)thiazole-5-carboxylic acid (**S46**) (104 mg, 0.45 mmol, 1.0 eq) and 3-(4-methylpiperidin-1-yl)propan-1-amine (**S8**) (139 mg, 0.89 mmol, 2.0 eq) and purifying by chromatography on silica gel using 9:1:0.05 EtOAc:MeOH:$NEt_3$ as eluent, 2-(3,4-dimethylphenyl)-*N*-(3-(4-methylpiperidin-1-yl)propyl)thiazole-5-carboxamide was obtained as a white solid (102 mg, 0.27 mmol, 61%). $^{1}H$ NMR (400 MHz, $CDCl_3$) δ 8.63 (br s, 1H), 8.34 (s, 1H), 7.76 (s, 1H), 7.69 (d, 1H, *J* = 7.8 Hz), 7.20 (d, 1H, *J* = 7.8 Hz), 3.59 (q, 2H, *J* = 5.8 Hz), 3.25 (d, 2H, *J* = 8.8 Hz), 2.79 (br s, 2H), 2.32 (s, 3H), 2.30 (s, 3H), 1.96 (br s, 2H), 1.80-1.76 (m, 3H), 1.63-1.56 (m, 3H), 1.02 (d, 3H, *J* = 5.9 Hz); $^{13}C$ NMR (101 MHz, CHLOROFORM-*D*) δ 171.30, 161.14, 144.87, 140.11, 137.60, 133.54, 131.05, 130.42, 127.94, 124.46, 77.48, 77.36, 77.16, 76.84, 72.20, 60.54, 30.91, 23.73, 21.19, 19.97, 19.84, 14.34, 13.82.

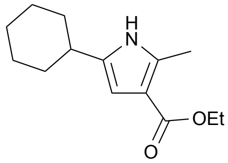

**Ethyl 5-cyclohexyl-2-methyl-1*H*-pyrrole-3-carboxylate (S47)**
To a solution of ethyl 5-(cyclohex-1-en-1-yl)-2-methyl-1*H*-pyrrole-3-carboxylate (680 mg, 2.91 mmol,1.0 eq) in EtOH (12 mL) was added Palladium, (10% on activated carbon wet, 155 mg, 1.46 mmol, 0.5 eq) and the reaction mixture was stirred under $H_2$ atmosphere for 30 min. The solution was filtered on a short plug of Celite and rinsed with methanol. Filtrate was concentrated to obtain ethyl 5-cyclohexyl-2-methyl-1*H*-pyrrole-3-carboxylate (685 mg, 2.91 mmol, quant) as a yellow gum. LRMS (ESI) [M+H]+ = 236.3.

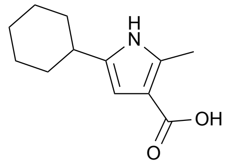

**5-cyclohexyl-2-methyl-1*H*-pyrrole-3-carboxylic acid (S48)**
To the solution of ethyl 5-cyclohexyl-2-methyl-1*H*-pyrrole-3-carboxylate (**S47**) (700 mg, 2.97 mmol, 1.0 eq) in MeOH:H2O (1:1) (0.2 M) was added LiOH (5.0 eq) and the reaction heated to 80 °C overnight. Acidified with 1 M HCl to precipitate solid that was collected by filtration. If a colloidal suspension formed, the mixture was instead extracted 3 times with EtOAc, washed with brine, fried and $Na_2SO_4$ and concentrated *in vacuo*. Product 5-cyclohexyl-2-methyl-1*H*-pyrrole-3-carboxylic acid (580 mg, 2.80 mmol, 94%) obtained as a beige solid. LRMS (ESI) [M+H]+ = 208.2.

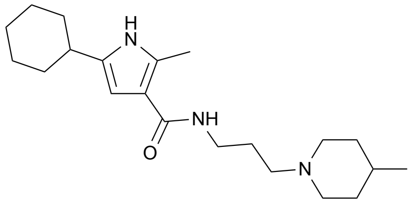

**5-cyclohexyl-2-methyl-*N*-(3-(4-methylpiperidin-1-yl)propyl)-1*H*-pyrrole-3-carboxamide (Compound 21)**

Following general procedure **I**, starting from 5-cyclohexyl-2-methyl-1*H*-pyrrole-3-carboxylic acid (**S48**) (200 mg, 0.96 mmol, 1.0 eq) and 3-(4-methylpiperidin-1-yl)propan-1-amine (**S8**) (181 mg, 1.16 mmol, 1.2 eq). After purification described in general procedure **I**, sample was re-purified on a C18 reverse phase column (40 g, AmFor:MeCN 5–70%) to afford 5-cyclohexyl-2-methyl-*N*-(3-(4-methylpiperidin-1-yl)propyl)-1*H*-pyrrole-3- carboxamide (190 mg, 0.55 mmol, 56%) obtained as a beige solid. LRMS (ESI) [M+H]+ =346.5; 1H NMR (400 MHz, DMSO-d6) δ 10.60 (s, 1H), 7.55 (t, J = 5.2 Hz, 1H), 6.01 (d, J = 2.2 Hz, 1H), 3.42 – 3.24 (m, 1H), 3.17 (q, J = 6.4 Hz, 2H), 3.01 (d, J = 8.3 Hz, 2H), 2.56 – 2.51 (m, 1H), 2.46 – 2.36 (m, 1H), 2.34 (s, 3H), 2.26 – 2.04 (m, 2H), 1.95 – 1.82 (m, 2H),1.79 – 1.58 (m, 7H), 1.37 – 1.09 (m, 8H), 0.90 (d, J = 6.5 Hz, 3H); 0.75 eq of formate salt (8.18 ppm). $^{13}$C NMR (101 MHz, $CDCl_3$) δ 168.91, 167.69, 136.62, 132.10, 112.99, 102.04, 54.60, 52.64, 36.55, 36.48, 33.10, 31.50, 29.36, 26.29, 26.11, 24.58, 21.08, 13.12.

### <u>Preparation of *N*-alkyl pyrroles</u>

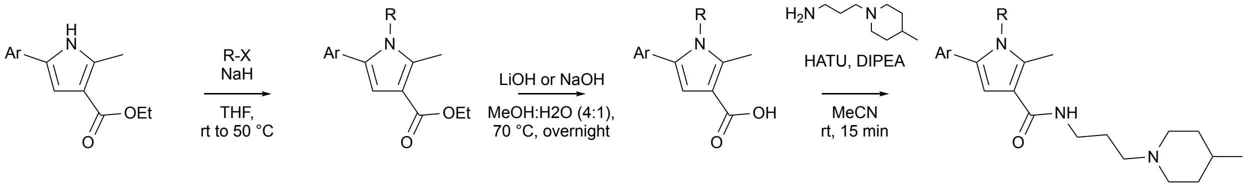

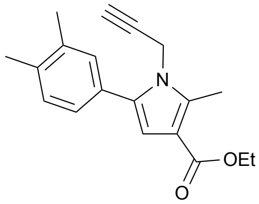

**Ethyl 5-(3,4-dimethylphenyl)-2-methyl-1-(prop-2-yn-1-yl)-1*H*-pyrrole-3-carboxylate (S49)**
Following the general procedure H, starting from ethyl 5-(2,3-dimethylphenyl)-2-methyl-1Hpyrrole-3-carboxylate (**S19**) (850 mg, 3.30 mmol, 1.0 eq) and 3-bromoprop-1-yne (2.46 g, 16.5 mmol, 5.0 eq). Purification by flash column chromatography ($SiO_2$, 25 g, Hex:Et:OAc 0–15%) afforded ethyl 5-(3,4-dimethylphenyl)-2-methyl-1-(prop-2-yn-1-yl)-1*H*-pyrrole-3-carboxylate (900 mg, 3.05 mmol, 92%) as a yellow oil. LRMS (ESI) [M+H]+ = 296.2.

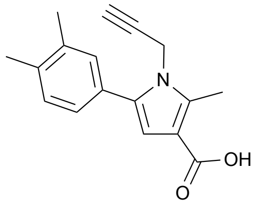

**5-(2,3-dimethylphenyl)-2-methyl-1-(prop-2-yn-1-yl)-1*H*-pyrrole-3-carboxylic acid (S50)**
Following general procedure E, starting from ethyl 5-(2,3-dimethylphenyl)-2-methyl-1-(prop-2-yn-1-yl)-1*H*-pyrrole-3-carboxylate (**S50**) (900 mg, 3.05 mmol, 1.0 eq). Product 5-(2,3-dimethylphenyl)-2-methyl-1-(prop-2-yn-1-yl)-1*H*-pyrrole-3-carboxylic acid (430 mg, 1.61mmol, 52%) obtained as a tan solid. LRMS (ESI) [M+H]+ = 268.3.

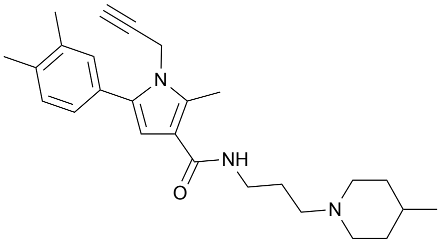

**5-(3,4-dimethylphenyl)-2-methyl-*N*-(3-(4-methylpiperidin-1-yl)propyl)-1-(prop-2-yn-1-yl)-1*H*-pyrrole-3-carboxamide (Compound 16)**
Following general procedure **I**, starting from 5-(3,4-dimethylphenyl)-2-methyl-1-(prop-2-yn-1-yl)-1*H*-pyrrole-3-carboxylic acid (**S50**) (400 mg, 1.50 mmol, 1.0 eq) and 3-(4-methylpiperidin-1-yl)propan-1-amine (**S8**) (351 mg, 2.24 mmol, 1.5 eq). After purification described in general procedure **I**, sample was re-purified on a C18 reverse phase column (40 g, AmFor:MeCN 5–55%) to afford 5-(3,4-dimethylphenyl)-2-methyl-*N*-(3-(4-methylpiperidin-1-yl)propyl)-1-(prop-2-yn-1-yl)-1*H*-pyrrole-3-carboxamide (107 mg, 0.21 mmol, 14%) as an off-white solid. LRMS (ESI) [M+H]+ = 406.6; 1H NMR (400 MHz, CDCl3) δ 7.55 (app. br. s, 1H), 7.20 (s, 1H), 7.18 – 7.10 (m, 2H), 6.66 (t, J = 6.5 Hz, 1H), 6.36 (s, 1H), 5.21 (d, J = 6.5 Hz, 2H), 3.49 (dd, J = 11.6, 5.4 Hz, 2H), 3.00 (d, J = 11.2 Hz, 2H), 2.67 (s, 3H), 2.51 (t, J = 5.8 Hz, 2H), 2.28 (s, 6H), 1.93 (t, J = 9.8 Hz, 2H), 1.77 (dt, J = 11.6, 5.8 Hz, 2H), 1.63 (d, J = 10.9 Hz, 2H), 1.45 – 1.07 (m, 3H), 0.82 (d, J = 5.9 Hz, 3H). $^{13}$C NMR (101 MHz, DMSO) δ 164.42, 136.19, 135.06, 134.09, 131.98, 129.82, 129.44, 129.24, 125.54, 116.11, 107.52, 95.44, 84.75, 56.57, 53.57, 37.58, 34.06, 30.45, 26.45, 21.78, 19.48, 19.12, 11.92, 1.17.

### Preparation of *N*-aryl and *N*-alkyl pyrroles

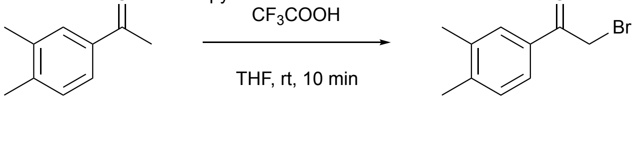

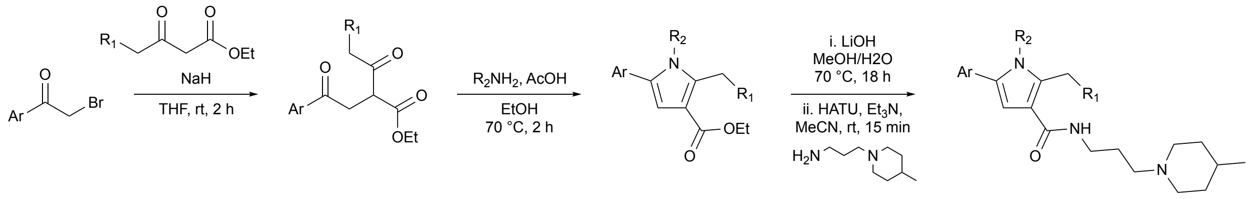

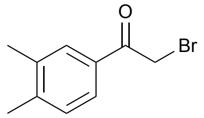

**2-bromo-1-(3,4-dimethylphenyl)ethenone (S51)**
To a solution of 1-(3,4-dimethylphenyl)ethanone (2 g, 13.50 mmol, 2.0 mL, 1.0 eq) in THF (40 mL) at RT were added $CF_3COOH$ (1.5 g, 13.50 mmol, 1.04 mL, 1.0 eq) followed by pyridinium tribromide (4.7 g, 14.85 mmol, 1.1 eq) and stirred at room temperature for 10 min. The reaction mixture was quenched by adding 50 mL of water; aqueous was extracted with EtOAc (2 x 50 mL); combined organic layer was washed with 40 mL of a sat. aq. solution of $CuSO_4$, brine, dried on Na2SO4 and concentrated *in vacuo*. Purification by a flash column chromatography ($SiO_2$, 100 g, Hex:EtOAc 0–5%) afforded 2-bromo-1-(3,4-dimethylphenyl)ethanone (2.6 g, 11.36 mmol, 84%) as a white solid. 1H NMR (400 MHz, DMSO-d6) δ 7.78 (d, J = 1.1 Hz, 1H), 7.74 (dd, J = 7.9, 1.8 Hz, 1H), 7.30 (d, J = 7.9 Hz, 1H), 4.87 (s, 2H), 2.29 (s, 6H).

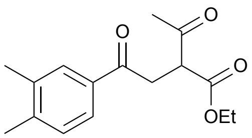

**Ethyl 2-acetyl-4-(3,4-dimethylphenyl)-4-oxobutanoate (S52)**
Following general procedure J, starting from ethyl 3-oxobutanoate (1.4 g, 1.39 mL, 11.01 mmol, 1.0 eq) and 2-bromo-1-(3,4-dimethylphenyl)ethanone (**S51**) (2.5 g, 11.01 mmol, 1.0 eq). Purification by a flash column chromatography ($SiO_2$; 50 g, Hex:EtOAc 0–40%) afforded ethyl 2-acetyl-4-(3,4-dimethylphenyl)-4-oxobutanoate (2.9 g, 10.49 mmol, 95%) as a clear oil. LRMS (ESI) [M+H]- = 277.3; 1H NMR (400 MHz, CDCl3) δ 7.74 (s, 1H), 7.71 (d, J = 7.8 Hz, 1H), 7.21 (d, J = 7.8 Hz, 1H), 4.24 – 4.20 (m, 3H), 3.68 (dd, J = 18.3, 8.3 Hz, 1H), 3.49 (dd, J = 18.3, 5.6 Hz, 1H), 2.44 (s, 3H), 2.31 (s, 6H), 1.29 (t, J = 7.1 Hz, 3H).

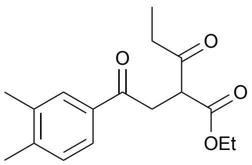

**Ethyl 2-(2-(3,4-dimethylphenyl)-2-oxoethyl)-3-oxopentanoate (S53)**
Following general procedure J, starting from ethyl 3-oxopentanoate (698 mg, 4.84 mmol, 1.1 eq) and 2-bromo-1-(3,4-dimethylphenyl)ethenone (**S51**) (1 g, 4.40 mmol, 1.0 eq). Purification by a flash column chromatography ($SiO_2$; 50 g, Hex:EtOAc 0–20%) afforded ethyl 2-(2-(3,4-dimethylphenyl)-2-oxoethyl)-3-oxopentanoate (1.27 g, 4.37 mmol, quant) as a pale yellow oil. LRMS (ESI) [M-*H*- = 289.3; 1H NMR (400 MHz, CDCl3) δ 7.78 – 7.66 (m, 2H), 7.21 (d, J = 7.9 Hz, 1H), 4.26 – 4.15 (m, 3H), 3.70 (dd, J = 18.3, 8.4 Hz, 1H), 3.50 (dd, J = 18.3, 5.5 Hz, 1H), 2.83 – 2.78 (m, 1H), 2.31 (s, 6H), 2.31 – 2.28 (m, 1H), 1.28 (t, J = 7.1 Hz, 3H), 1.11 (t, J = 7.2 Hz, 3H).

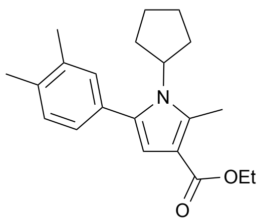

**Ethyl 1-cyclopentyl-5-(3,4-dimethylphenyl)-2-methyl-1*H*-pyrrole-3-carboxylate (S54)**

Following general procedure K, starting from ethyl 2-acetyl-4-(3,4-dimethylphenyl)-4-oxobutanoate (**S52**) (600 mg, 2.17 mmol, 1.0 eq) and cyclopentanamine (187 mg, 2.19 mmol, 1.01 eq). Purification by a flash column chromatography ($SiO_2$, 25 g, Hex:Et:OAc 0–15%) afforded ethyl 1-cyclopentyl-5-(3,4-dimethylphenyl)-2-methyl-1Hpyrrole-3-carboxylate (706 mg, 2.17 mmol, quant) as a clear gum. LRMS (ESI) [M+H]+ = 326.2; 1H NMR (400 MHz, CDCl3) δ 7.15 (br. d, J = 7.7 Hz, 1H), 7.11 (br. s, 1H), 7.05 (br. dd, J = 7.7, 1.6 Hz, 1H), 6.44 (s, 1H), 4.26 (q, J = 7.1 Hz, 2H), 2.69 (s, 3H), 2.30 (s, 3H), 2.29 (s, 3H), 2.08 – 1.93 (m, 4H), 1.94 – 1.75 (m, 2H), 1.66 – 1.48 (m, 3H), 1.32 (t, J = 7.1 Hz, 3H).

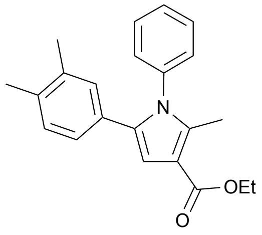

**Ethyl 5-(3,4-dimethylphenyl)-2-methyl-1-phenyl-1*H*-pyrrole-3-carboxylate (S55)**

Following general procedure K, starting from ethyl 2-acetyl-4-(3,4-dimethylphenyl)-4-oxobutanoate (**S52**) (600 mg, 2.17 mmol, 1.0 eq) and aniline (204 mg, 2.19 mmol, 0.20 mL, 1.01 eq). Product ethyl 5-(3,4-dimethylphenyl)-2-methyl-1-phenyl-1*H*-pyrrole-3-carboxylate (723 mg mg, 2.17 mmol, 99%) obtained as a yellow solid. LRMS (ESI) [M+H]+ = 334.2; 1H NMR (400 MHz, CDCl3) δ 7.42 – 7.33 (m, 3H), 7.19 – 7.12 (m, 2H), 6.90 (br. s, 1H), 6.87 (d, J = 7.9 Hz, 1H), 6.75 (s, 1H), 6.68 (dd, J = 7.8, 1.7 Hz, 1H), 4.32 (q, J = 7.1 Hz, 2H), 2.39 (s, 3H), 2.16 (s, 3H), 2.11 (s, 3H), 1.37 (t, J = 7.1 Hz, 3H).

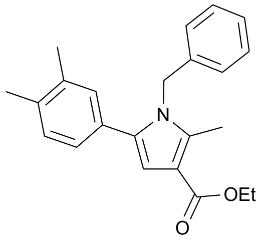

**Ethyl 1-benzyl-5-(3,4-dimethylphenyl)-2-methyl-1*H*-pyrrole-3-carboxylate (S56)**

Following general procedure K, starting from ethyl 2-acetyl-4-(3,4-dimethylphenyl)-4-oxobutanoate (**S52**) (600 mg, 2.17 mmol, 1.0 eq) and phenylmethanamine (302 mg, 2.82 mmol, 1.3 eq). Purification by a flash column chromatography ($SiO_2$, 25 g, Hex:Et:OAc 0–15%) afforded ethyl 1-benzyl-5-(3,4-dimethylphenyl)-2-methyl-1*H*-pyrrole-3-carboxylate (750 mg, 2.17 mmol, quant) as a clear gum. LRMS (ESI) [M+H]+ = 348.4; 1H NMR (400 MHz, CDCl3) δ 7.33 – 7.21 (m, 3H), 7.10 – 7.02 (m, 2H), 6.99 (dd, J = 7.8, 1.6 Hz, 2H), 6.91 (d, J = 7.3 Hz, 2H), 6.64 (s, 1H), 5.12 (s, 2H), 4.29 (q, J = 7.1 Hz, 2H), 2.45 (s, 3H), 2.24 (s, 3H), 2.18 (s, 3H), 1.36 (t, J = 7.1 Hz, 3H).

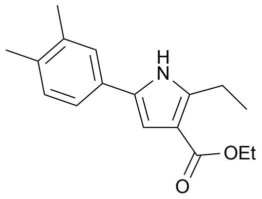

**Ethyl 5-(3,4-dimethylphenyl)-2-ethyl-1*H*-pyrrole-3-carboxylate (S57)**

Following general procedure K, starting from ethyl 2-(2-(3,4-dimethylphenyl)-2-oxoethyl)-3-oxopentanoate (**S53**) (1.3 g, 4.37 mmol, 1.0 eq) and ammonium acetate (1.4 g, 17.50 mmol, 4.0 eq). Purification by flash column chromatography ($SiO_2$, 50 g, Hex:Et:OAc 0–20%) afforded ethyl 5-(3,4-dimethylphenyl)-2-ethyl-1*H*-pyrrole-3-carboxylate (1 g, 3.69 mmol, 84%) as an off-white solid. LRMS (ESI) [M+H]+ = 271.2; 1H NMR (400 MHz, CDCl3) δ 8.49 (br. s, 1H), 7.25 (s, 1H), 7.20 (dd, J = 7.7, 1.9 Hz, 1H), 7.12 (d, J = 7.8 Hz, 1H), 6.79 (d, J = 2.9 Hz, 1H), 4.29 (q, J = 7.1 Hz, 2H), 3.03 (q, J = 7.6 Hz, 2H), 2.28 (s, 3H), 2.26 (s, 3H), 1.36 (t, J = 7.1 Hz, 3H), 1.30 (t, J = 7.6 Hz, 3H).

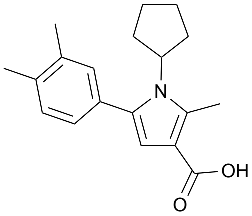


**1-cyclopentyl-5-(3,4-dimethylphenyl)-2-methyl-1*H*-pyrrole-3-carboxylic acid (S58)**

Following general procedure E, starting from ethyl 1-cyclopentyl-5-(3,4-dimethylphenyl)-2-methyl-1*H*-pyrrole-3-carboxylate (**S54**) (705 mg, 2.17 mmol, 1.0 eq). Product 1-cyclopentyl-5-(3,4-dimethylphenyl)-2-methyl-1*H*-pyrrole-3-carboxylic acid (470 mg, 0.64 mmol, 72%) obtained as a white solid. LRMS (ESI) [M+H]+ = 298.3.

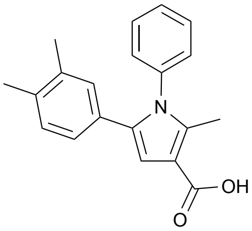


**5-(3,4-dimethylphenyl)-2-methyl-1-phenyl-1*H*-pyrrole-3-carboxylic acid (S59)**

Following general procedure E, starting from ethyl 2-methyl-1,5-diphenyl-1*H*-pyrrole-3-carboxylate (**S55**) (720 mg, 2.16 mmol, 1.0 eq). Product 5-(3,4-dimethylphenyl)-2-methyl-1-phenyl-1Hpyrrole-3-carboxylic acid (630 mg, 2.06 mmol, 95%) obtained as an off-white solid. LRMS (ESI) [M+H]+ = 306.3.

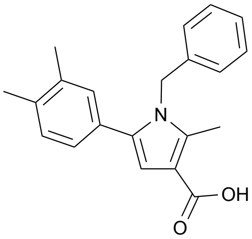


**1-benzyl-5-(3,4-dimethylphenyl)-2-methyl-1*H*-pyrrole-3-carboxylic acid (S60)**

Following general procedure E, starting from ethyl 1-cyclopentyl-5-(3,4-dimethylphenyl)-2-methyl-1*H*-pyrrole-3-carboxylate (**S56**) (750 mg, 2.16 mmol, 1.0 eq). Product 1-benzyl-5-(3,4-dimethylphenyl)-2-methyl-1*H*-pyrrole-3-carboxylic acid (595 mg, 1.86 mmol, 86%) obtained as a white solid. LRMS (ESI) [M+H]+ = 320.4.

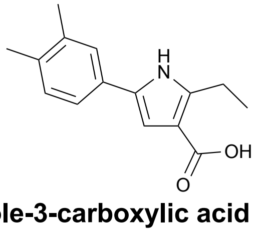


**5-(3,4-dimethylphenyl)-2-ethyl-1*H*-pyrrole-3-carboxylic acid (S61)**

Following general procedure E, starting from ethyl 5-(3,4-dimethylphenyl)-2-ethyl-1*H*-pyrrole-3-carboxylate (**S57**) (1 g, 3.69 mmol, 1.0 eq). Product 5-(3,4-dimethylphenyl)-2-ethyl-1*H*-pyrrole-3-carboxylic acid (750 mg, 3.08 mmol, 83%) obtained as a tan solid. LRMS (ESI) [M-H]- = 242.4.

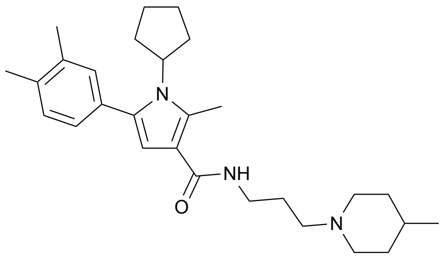

**1-cyclopentyl-5-(3,4-dimethylphenyl)-2-methyl-*N*-(3-(4-methylpiperidin-1-yl)propyl)-1*H*-pyrrole-3-carboxamide (Compound 17)**

Following general procedure **I**, starting from 1-cyclopentyl-5-(3,4-dimethylphenyl)-2-methyl-1*H*-pyrrole-3-carboxylic acid (**S58**) (200 mg, 0.67 mmol, 1.0 eq) and 3-(4-methylpiperidin-1-yl)propan-1-amine (**S8**) (158 mg, 1.01 mmol, 1.5 eq). After purification described in general procedure **I**, sample was re-purified on a C18 reverse phase column (40 g, AmFor:MeCN 5-66%) to afford 1-cyclopentyl-5-(3,4-dimethylphenyl)-2-methyl-N-(3-(4-methylpiperidin-1-yl)propyl)-1*H*-pyrrole-3-carboxamide (198 mg, 0.45 mmol, 67%) as a tan solid. LRMS (ESI) [M+H]+ = 466.6; 1H NMR (500 MHz, CDCl3) δ 7.30 (s, 1H), 7.13 (d, J = 7.7 Hz, 1H), 7.09 (s, 1H), 7.04 (dd, J = 7.7, 1.6 Hz, 1H), 6.21 (s, 1H), 4.62 (p, J = 9.3 Hz, 1H), 3.46 (dd, J = 11.9, 5.4 Hz, 2H), 3.02 (d, J = 11.5 Hz, 2H), 2.72 (s, 3H), 2.54 (t, J = 6.3 Hz, 2H), 2.29 (s, 3H), 2.28 (s, 3H), 2.09 – 1.92 (m, 6H), 1.90 – 1.81 (m, 2H), 1.78 (dt, J = 12.4, 6.3 Hz, 2H), 1.67 – 1.50 (m, 4H), 1.44 – 1.23 (m, 3H), 0.79 (d, J = 6.0 Hz, 3H). $^{13}C$ NMR (101 MHz, DMSO) δ 164.97, 136.29, 135.42, 133.41, 132.25, 131.08, 130.39, 129.48, 126.57, 115.81, 106.90, 56.54, 53.47, 37.52, 33.88, 31.11, 30.32, 26.33, 24.74, 21.68, 19.43, 19.09, 12.85. (1 peak missing, probably overlap)

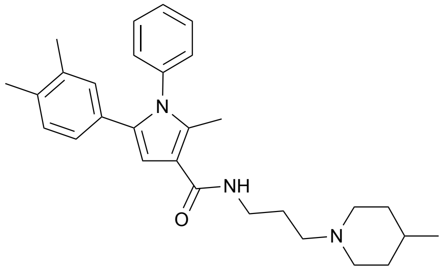

**5-(3,4-dimethylphenyl)-2-methyl-N-(3-(4-methylpiperidin-1-yl)propyl)-1-phenyl-1Hpyrrole-3-carboxamide (Compound 18)**

Following general procedure **I**, starting from 5-(3,4-dimethylphenyl)-2-methyl-1-phenyl-1*H*-pyrrole-3-carboxylic acid (**S59**) (200 mg, 0.65 mmol, 1.0 eq) and 3-(4-methylpiperidin-1-yl)propan-1-amine (**S8**) (205 mg, 1.31 mmol, 2.0 eqv). Product 5-(3,4-dimethylphenyl)-2-methyl-N-(3-(4-methylpiperidin-1-yl)propyl)-1-phenyl-1Hpyrrole-3-carboxamide (225 mg, 0.51 mmol, 77%) obtained as an off-white solid. LRMS (ESI) [M+H]+ = 444.6; 1H NMR (400 MHz, DMSO-d6) δ 8.02 (t, J = 5.8 Hz, 1H), 7.52 – 7.39 (m, 3H), 7.24 – 7.13 (m, 2H), 6.91 – 6.89 ( m, 1H), 6.88 (s, 1H), 6.77 (overlapping s, 1H), 6.61 (dd, J = 7.8, 1.7 Hz, 1H), 3.49 – 3.31 (m, 2H), 3.28 (dd, J = 12.4, 6.3 Hz, 2H), 3.07 – 2.89 (m, 2H), 2.92 – 2.62 (m, 2H), 2.30 (s, 3H), 2.11 (s, 3H), 2.07 (s, 3H), 1.90 – 1.82 (m, 2H), 1.82 – 1.74 (m, 2H), 1.69 – 1.47 (m, 1H), 1.44 – 1.18 (m, 2H), 0.92 (d, J = 6.5 Hz, 3H); 0.06 eq of formate salt (8.14 ppm). $^{13}C$ NMR (101 MHz, DMSO) δ 165.21, 137.84, 135.89, 134.76, 134.42, 132.70, 129.77, 129.30, 129.17, 128.67, 128.51, 128.29, 124.75, 115.19, 107.21, 54.38, 52.13, 35.98, 31.30, 28.39, 24.90, 21.10, 19.38, 18.89, 11.96.

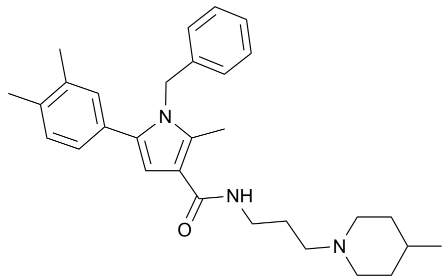

**1-benzyl-5-(3,4-dimethylphenyl)-2-methyl-N-(3-(4-methylpiperidin-1-yl)propyl)-1Hpyrrole-3-carboxamide (Compound 19)**

Following general procedure **I**, starting from 1-benzyl-5-(3,4-dimethylphenyl)-2-methyl-1*H*-pyrrole-3-carboxylic acid (**S60**) (200 mg, 0.63 mmol, 1.0 eq) and 3-(4-methylpiperidin-1-yl)propan-1-amine (**S8**) (196 mg, 1.25 mmol, 2.0 eq). After purification described in general procedure **I**, sample was re-purified on a C18 reverse phase column (40 g, AmFor:MeCN 5-80%) to afford 1-benzyl-5-(3,4-dimethylphenyl)-2-methyl-N-(3-(4-methylpiperidin-1-yl)propyl)-1*H*-pyrrole-3-carboxamide (70 mg, 0.15 mmol, 24%) as a beige solid. LRMS (ESI) [M+H]+ = 458.6; 1H NMR (400 MHz, CDCl3) δ 7.34 – 7.28 (m, 2H), 7.25 – 7.21 (m, 1H), 7.09 – 7.01 (m, 2H), 7.01 – 6.89 (m, 4H), 6.51 (s, 1H) 5.11 (s, 2H), 3.49 (dd, J = 11.6, 6.4 Hz, 2H), 3.36 (d, J = 11.8 Hz, 2H), 3.04 (t, J = 5.5 Hz, 2H), 2.87 (t, J = 12.3 Hz, 2H), 2.42 (s, 3H), 2.22 (s, 3H), 2.17 (s, 3H), 2.11 – 2.02 (m, 2H), 1.92 (d, J = 13.6 Hz, 2H), 1.84 – 1.61 (m, 1H), 1.61 – 1.39 (m, 2H), 0.98 (d, J = 6.3 Hz, 3H). $^{13}$C NMR (101 MHz, $CDCl_3$) δ 169.29, 137.73, 136.97, 136.50, 135.78, 135.45, 130.67, 129.89, 129.66, 129.04, 127.49, 126.45, 125.73, 113.07, 106.70, 53.65, 52.66, 47.93, 35.69, 31.85, 29.04, 24.70, 21.13, 19.87, 19.60, 11.84.

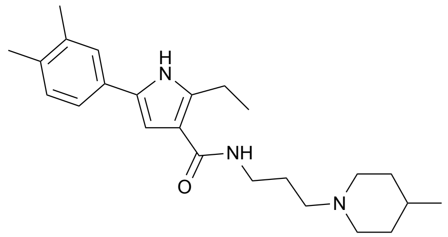

**5-(3,4-dimethylphenyl)-2-ethyl-N-(3-(4-methylpiperidin-1-yl)propyl)-1*H*-pyrrole-3-carboxamide (Compound 9)**

Following general procedure **I**, starting from 5-(3,4-dimethylphenyl)-2-ethyl-1*H*-pyrrole-3-carboxylic acid (200 mg, 0.82 mmol, 1.0 eq) (**S61**) and 3-(4-methylpiperidin-1-yl)propan-1-amine (**S8**) (257 mg, 1.64 mmol, 2.0 eq). After purification described in general procedure **I**, sample was re-purified on a C18 reverse phase column (40 g, AmBic:MeOH 0-85%) to afford 5-(3,4-dimethylphenyl)-2-ethyl-N-(3-(4-methylpiperidin-1-yl)propyl)-1Hpyrrole-3-carboxamide (158 mg, 0.41 mmol, 50%) as an off-white solid. LRMS (ESI) [M+H]+ = 382.5; 1H NMR (400 MHz, DMSO-d6) δ 11.24 (s, 1H), 7.90 (t, J = 5.7 Hz, 1H), 7.37 (br. s, 1H), 7.30 (dd, J = 7.8, 1.5 Hz, 1H), 7.13 (d, J = 7.9 Hz, 1H), 6.78 (d, J = 2.6 Hz, 1H), 3.44 (d, J = 10.0 Hz, 2H), 3.26 (q, J = 6.1 Hz, 2H), 3.11 – 2.97 (m, 2H), 2.92 (q, J = 7.4 Hz, 2H), 2.89 – 2.77 (m, 2H), 2.24 (s, 3H), 2.21 (s, 3H), 1.93 – 1.68 (m, 4H), 1.71 – 1.50 (m, 1H), 1.30 (dd, J = 24.6, 11.5 Hz, 2H), 1.17 (t, J = 7.4 Hz, 3H), 0.92 (d, J = 6.2 Hz, 3H); 0.53 eq of formate salt (9.02 ppm). $^{13}$C NMR (101 MHz, DMSO) δ 165.62, 140.01, 136.36, 133.76, 129.89, 129.85, 129.08, 124.49, 120.81, 114.01, 103.87, 54.03, 51.99, 35.50, 31.15, 28.14, 24.62, 21.09, 19.63, 19.56, 19.02, 14.90.

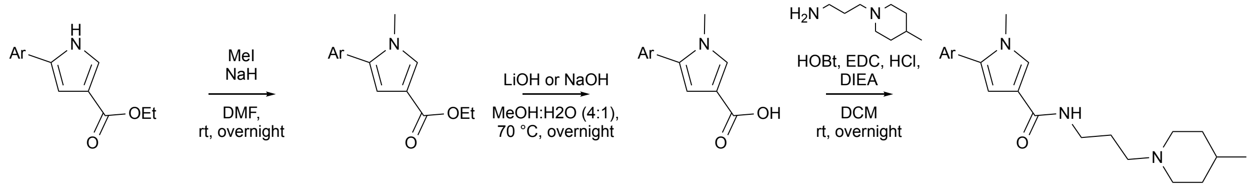

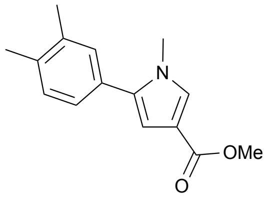

**methyl 5-(3,4-dimethylphenyl)-1-methyl-1*H*-pyrrole-3-carboxylate (S62)**
In an oven-dried round-bottom flask under argon, methyl 5-(3,4-dimethylphenyl)-1*H*-pyrrole-3-carboxylate (**S20**) (86 mg, 0.38 mmol, 1.0 eq) was dissolved in anhydrous DMF (3.7 mL, 0.1 M) at 0 °C and MeI (35 µL, 0.56 mmol, 1.1 eq) was added dropwise. NaH (60%, 23 mg, 0.56 mmol, 1.1 eq) was added portionwise and the resulting mixture was allowed to warm up slowly to room temperature and stirred overnight. The reaction was quenched with water and the aqueous phase was extracted 3 times with EtOAc. The combined organic layers were washed with brine, dried over $MgSO_4$ and concentrated under vacuum, to afford methyl 5-(3,4-dimethylphenyl)-1-methyl-1*H*-pyrrole-3-carboxylate as a colorless oil (40 mg, 0.17 mmol, 44%). $^{1}$H NMR (400 MHz, $CDCl_3$) δ 7.31 (d, 1H, *J* = 1.8 Hz), 7.18-7.16 (m, 2H), 7.11 (dd, 1H, *J* = 7.8 Hz, *J* = 1.9 Hz), 6.57 (d, 1H, *J* = 1.8 Hz), 3.81 (s, 3H), 3.64 (s, 3H), 2.30 (s, 6H); $^{13}$C NMR (101 MHz, $CDCl_3$) δ 165.7, 137.1, 136.5, 136.1, 130.5, 130.1, 128.4, 126.6, 115.3, 109.7, 51.4, 35.8, 20.2, 19.8.

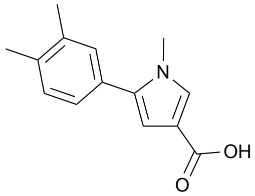

**5-(3,4-dimethylphenyl)-1-methyl-1*H*-pyrrole-3-carboxylic acid (S63)**
Following general procedure E, starting from methyl 5-(3,4-dimethylphenyl)-1-methyl-1*H*-pyrrole-3-carboxylate (**S62**) (40 mg, 0.17 mmol, 1.0 eq) and LiOH (20 mg, 0.82 mmol, 5.0 eq), 5-(3,4-dimethylphenyl)-1-methyl-1*H*-pyrrole-3-carboxylic acid was obtained as a white solid (37 mg, 0.16 mmol, 98%).

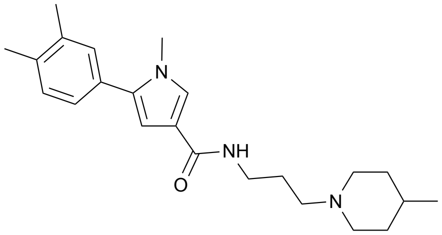

**5-(3,4-dimethylphenyl)-1-methyl-*N*-(3-(4-methylpiperidin-1-yl)propyl)-1*H*-pyrrole-3-carboxamide (Compound 15)**
Following general procedure F, starting from 5-(3,4-dimethylphenyl)-1-methyl-1*H*-pyrrole-3-carboxylic acid (**S63**) (37 mg, 0.16 mmol, 1.0 eq) and 3-(4-methylpiperidin-1-yl)propan-1-amine (**S8**) (50 mg, 0.32 mmol, 2.0 eq) and purifying by chromatography on silica gel using 9:1:0.05 EtOAc:MeOH:$NEt_3$ as eluent, 5-(3,4-dimethylphenyl)-1-methyl-*N*-(3-(4-methylpiperidin-1-yl)propyl)-1*H*-pyrrole-3-carboxamide was obtained as a white solid (32 mg, 0.09 mmol, 54%). $^{1}$H NMR (400 MHz, $CDCl_3$) δ 7.63 (br s, 1H), 7.31 (s, 1H), 7.16-7.09 (m, 3H), 6.46 (s, 1H), 3.63 (s, 3H), 3.51 (q, 2H, *J* = 5.4 Hz), 3.12 (d, 2H, *J* = 10.5 Hz), 2.65 (br s, 2H), 2.29 (s, 6H), 2.12 (br s, 2H), 1.87 (br s, 2H), 1.70 (d, 2H, *J* = 8.0 Hz), 1.45 (br s, 3H), 0.91 (br s, 3H); $^{13}$C NMR (101 MHz, $CDCl_3$) δ 165.3, 137.0, 136.3, 135.7, 130.0, 130.3, 130.0, 126.5, 126.1, 119.5, 107.2, 57.8, 54.2, 39.2, 35.7, 33.6, 30.7, 24.9, 21.8, 20.2, 19.8.
