## Supplementary material for "Phenotypic high-throughput screening identifies modulators of gut microbial choline metabolism": Document S3

Compound 5

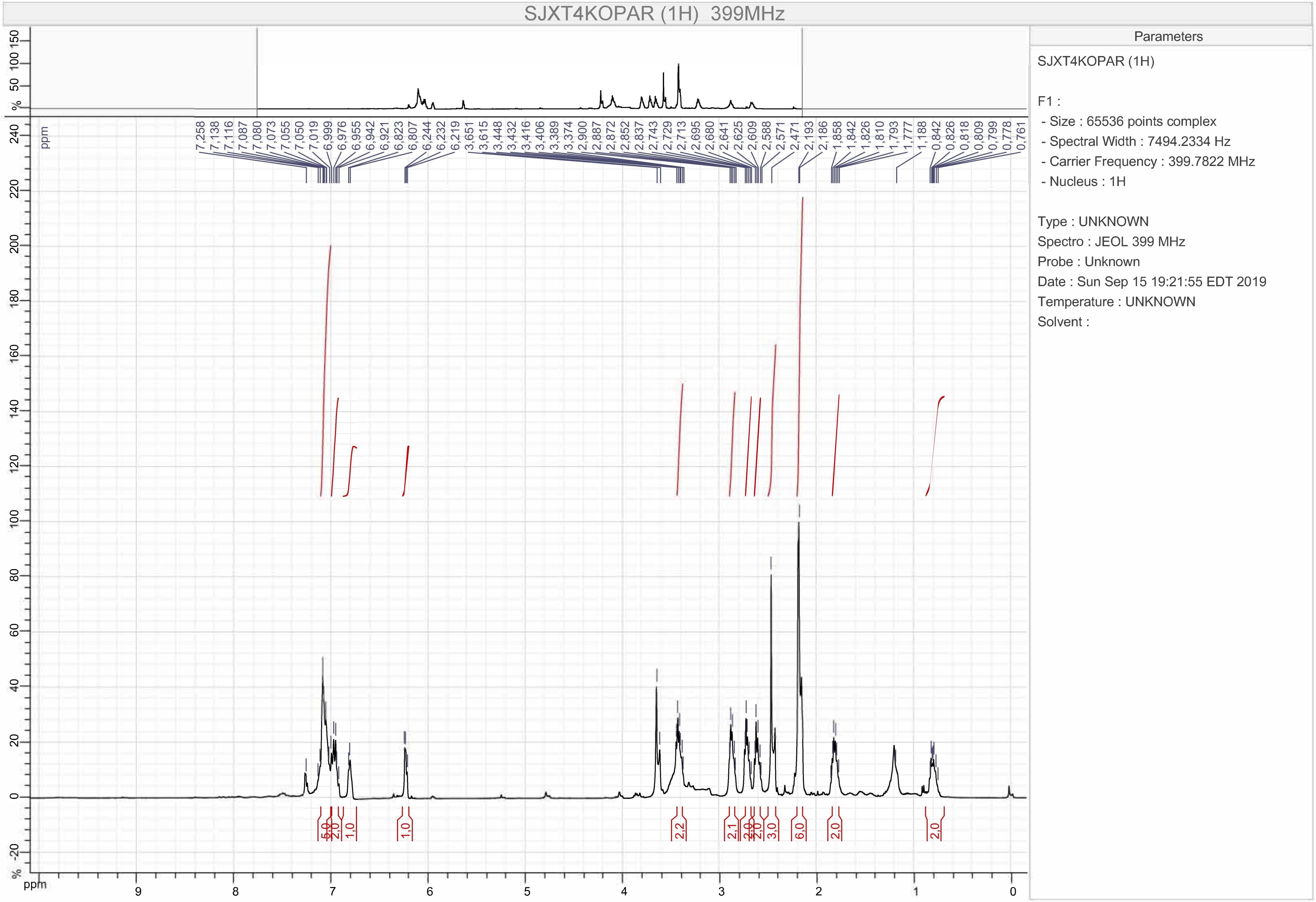
SJXT4KOPAR (1H) 399MHz
Parameters
SJXT4KOPAR (1H)
F1 :
- Size : 65536 points complex
- Spectral Width : 7494.2334 Hz
- Carrier Frequency : 399.7822 MHz
- Nucleus : 1H
Type : UNKNOWN
Spectro : JEOL 399 MHz
Probe : Unknown
Date : Sun Sep 15 19:21:55 EDT 2019
Temperature : UNKNOWN
Solvent :
ppm

Compound 5

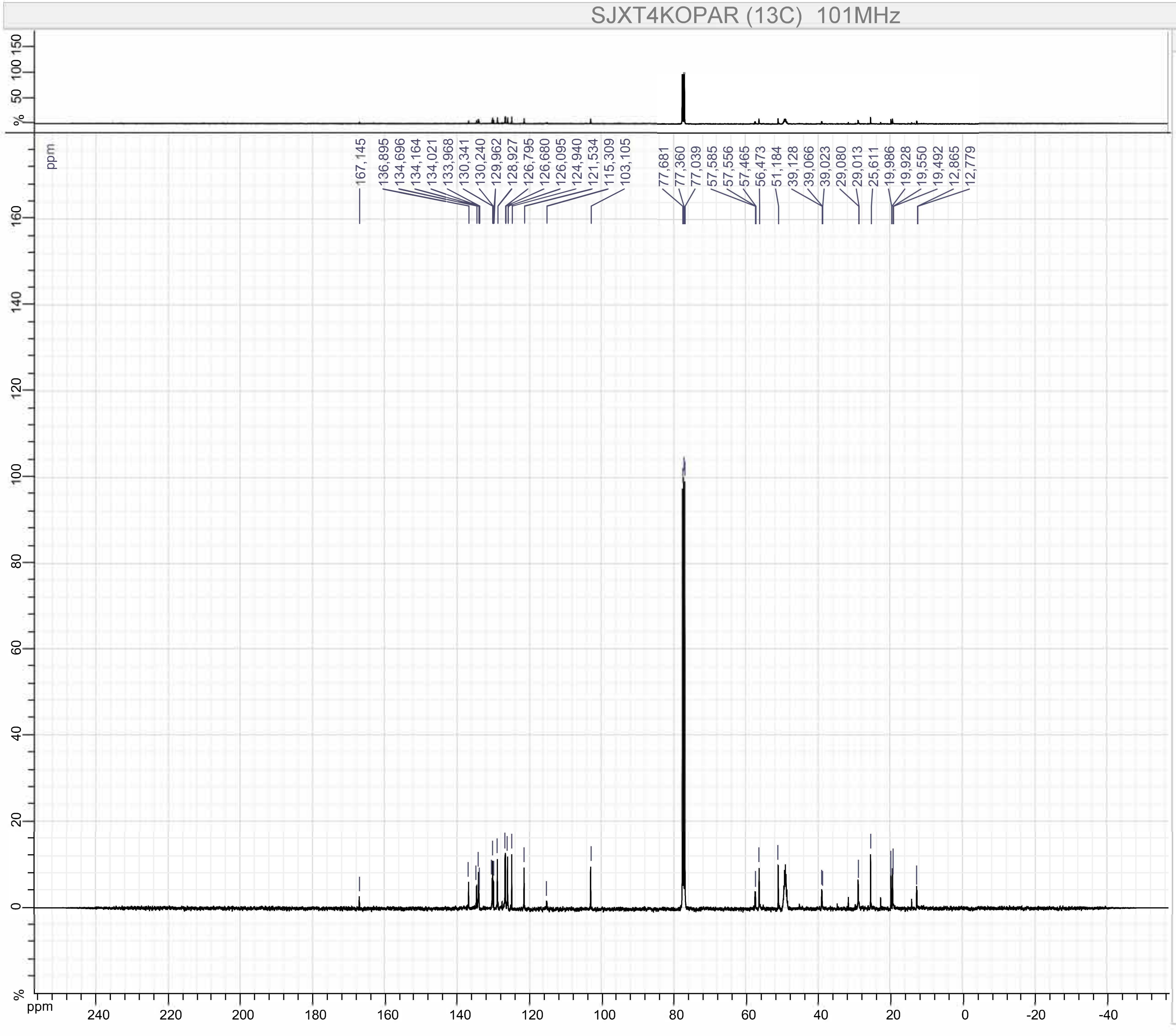


Parameters

SJXT4KOPAR (13C)

F1 :
- Size : 65536 points complex
- Spectral Width : 31566.62 Hz
- Carrier Frequency : 100.52531 MHz
- Nucleus : 13C

Type : UNKNOWN
Spectro : JEOL 101 MHz
Probe : Unknown
Date : Sun Sep 15 19:25:48 EDT 2019
Temperature : UNKNOWN
Solvent :

Compound 6

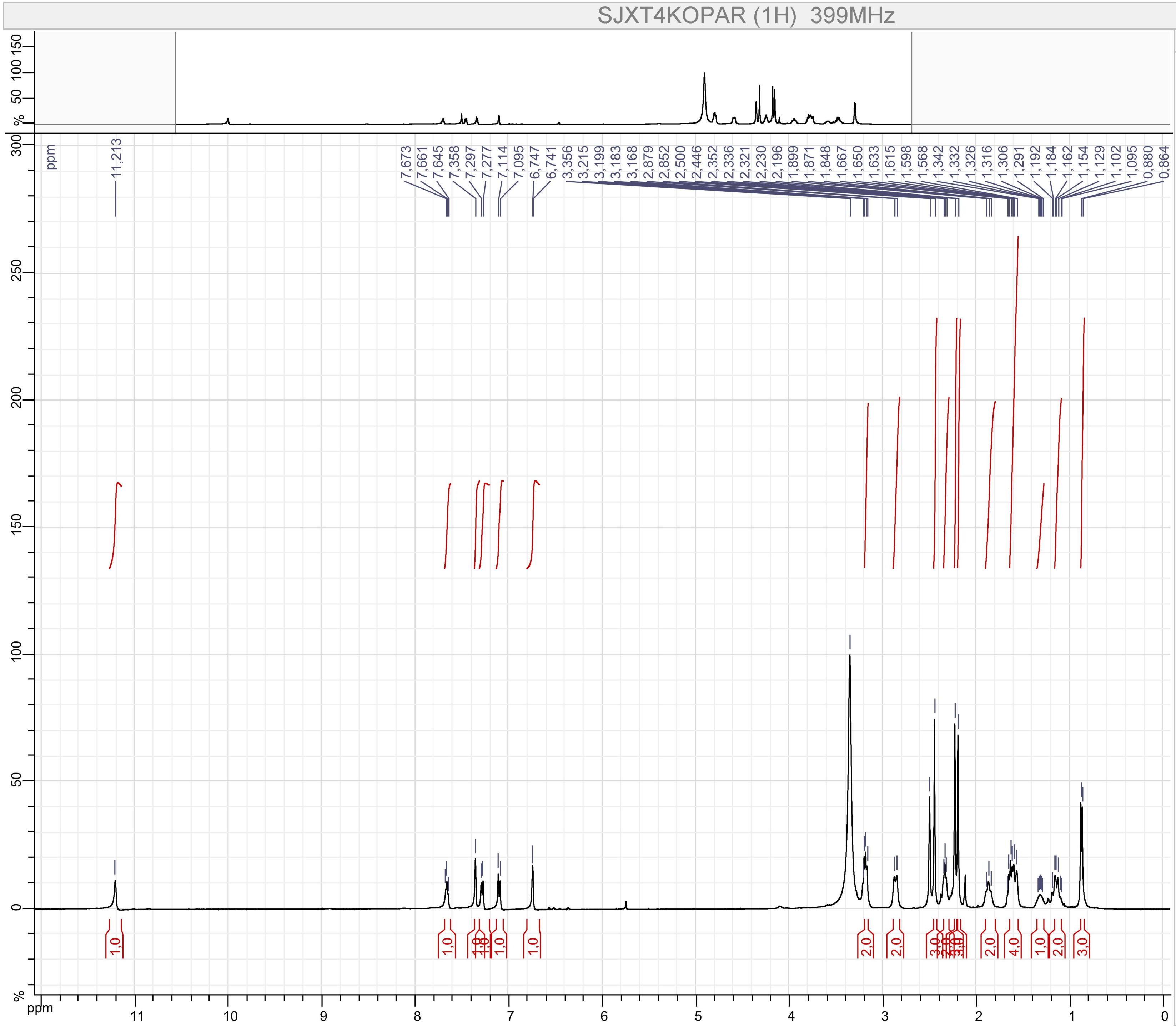


Parameters

SJXT4KOPAR (1H)

F1 :
- Size : 65536 points complex
- Spectral Width : 7494.2334 Hz
- Carrier Frequency : 399.7822 MHz
- Nucleus : 1H

Type : UNKNOWN
Spectro : JEOL 399 MHz
Probe : Unknown
Date : Fri Jul 12 23:06:06 EDT 2019
Temperature : UNKNOWN
Solvent :

Compound 6

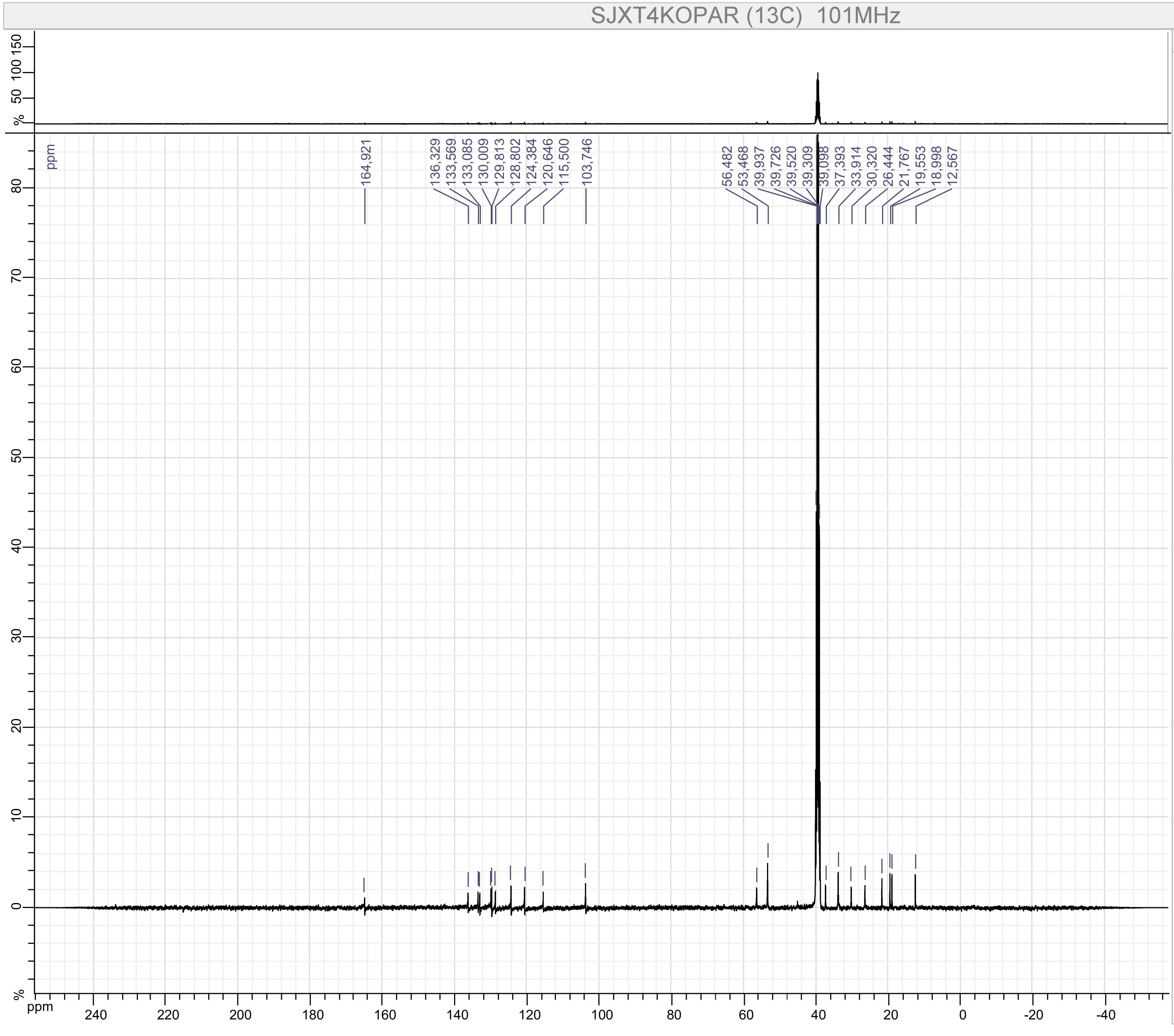


Parameters

SJXT4KOPAR (13C)

F1 :
- Size : 65536 points complex
- Spectral Width : 31566.62 Hz
- Carrier Frequency : 100.52531 MHz
- Nucleus : 13C

Type : UNKNOWN
Spectro : JEOL 101 MHz
Probe : Unknown
Date : Fri Jul 12 23:09:03 EDT 2019
Temperature : UNKNOWN
Solvent :

Compound 7

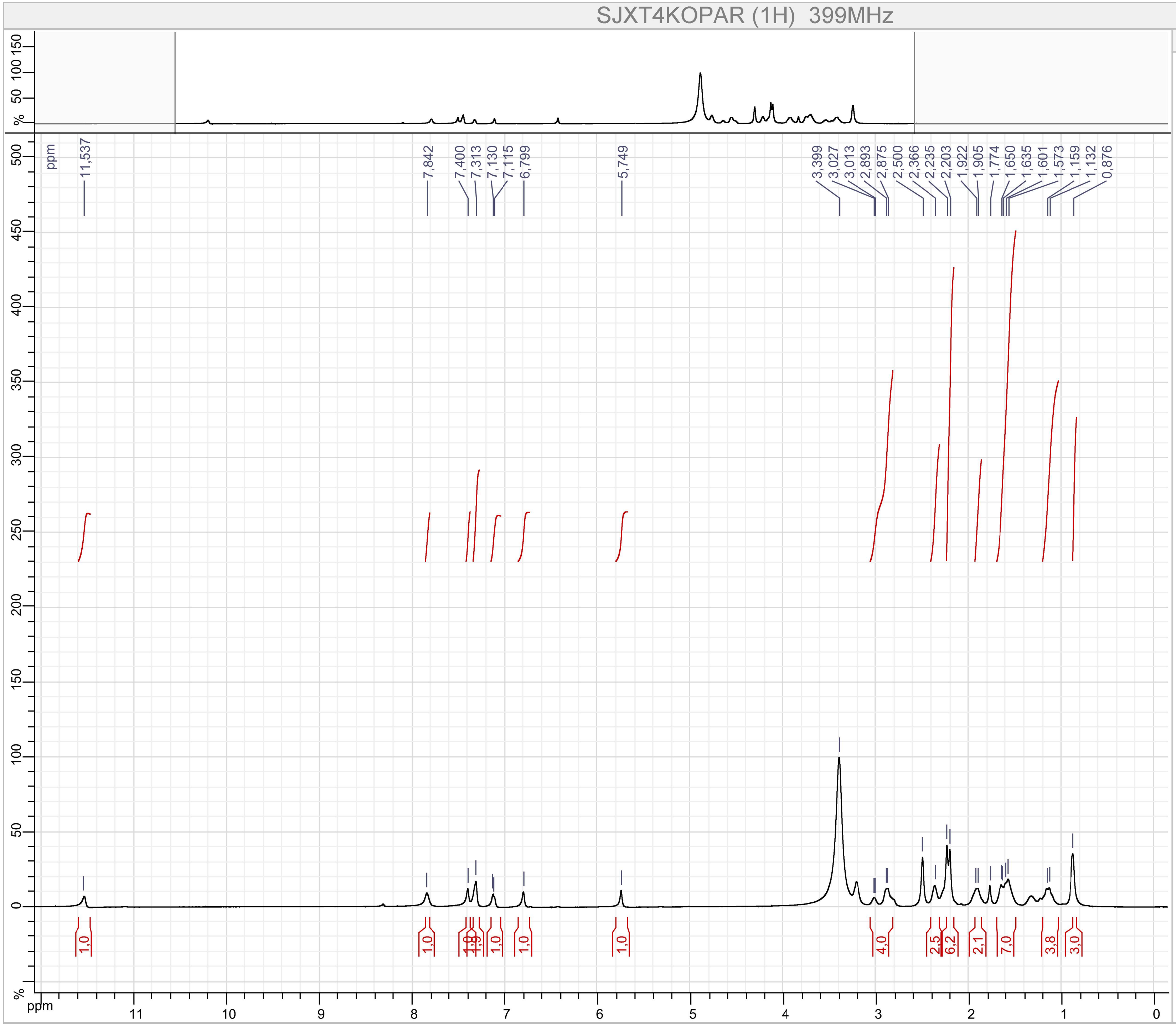


Parameters

SJXT4KOPAR (1H)

F1 :
- Size : 65536 points complex
- Spectral Width : 7494.2334 Hz
- Carrier Frequency : 399.7822 MHz
- Nucleus : 1H

Type : UNKNOWN
Spectro : JEOL 399 MHz
Probe : Unknown
Date : Mon Nov 05 21:04:32 EST 2018
Temperature : UNKNOWN
Solvent :

Compound 7

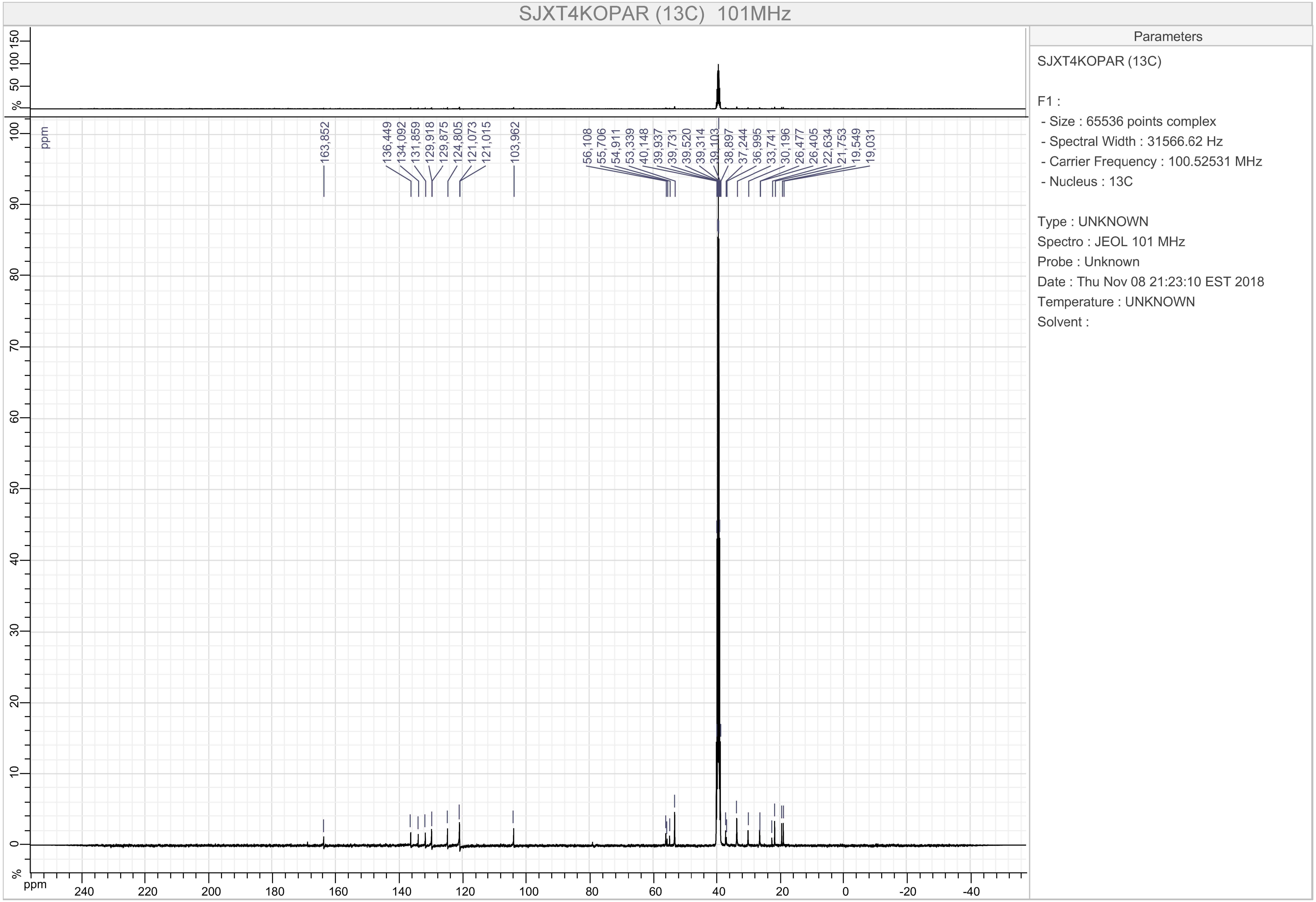

SJXT4KOPAR (13C) 101MHz
163,852
136,449
134,092
131,859
129,918
129,875
124,805
121,073
121,015
103,962
56,108
55,706
54,911
53,339
40,148
39,937
39,731
39,520
39,314
39,103
38,897
37,244
36,995
33,741
30,196
26,477
26,405
22,634
21,753
19,549
19,031
ppm
%
Parameters
SJXT4KOPAR (13C)
F1 :
- Size : 65536 points complex
- Spectral Width : 31566.62 Hz
- Carrier Frequency : 100.52531 MHz
- Nucleus : 13C
Type : UNKNOWN
Spectro : JEOL 101 MHz
Probe : Unknown
Date : Thu Nov 08 21:23:10 EST 2018
Temperature : UNKNOWN
Solvent :

Compound 8

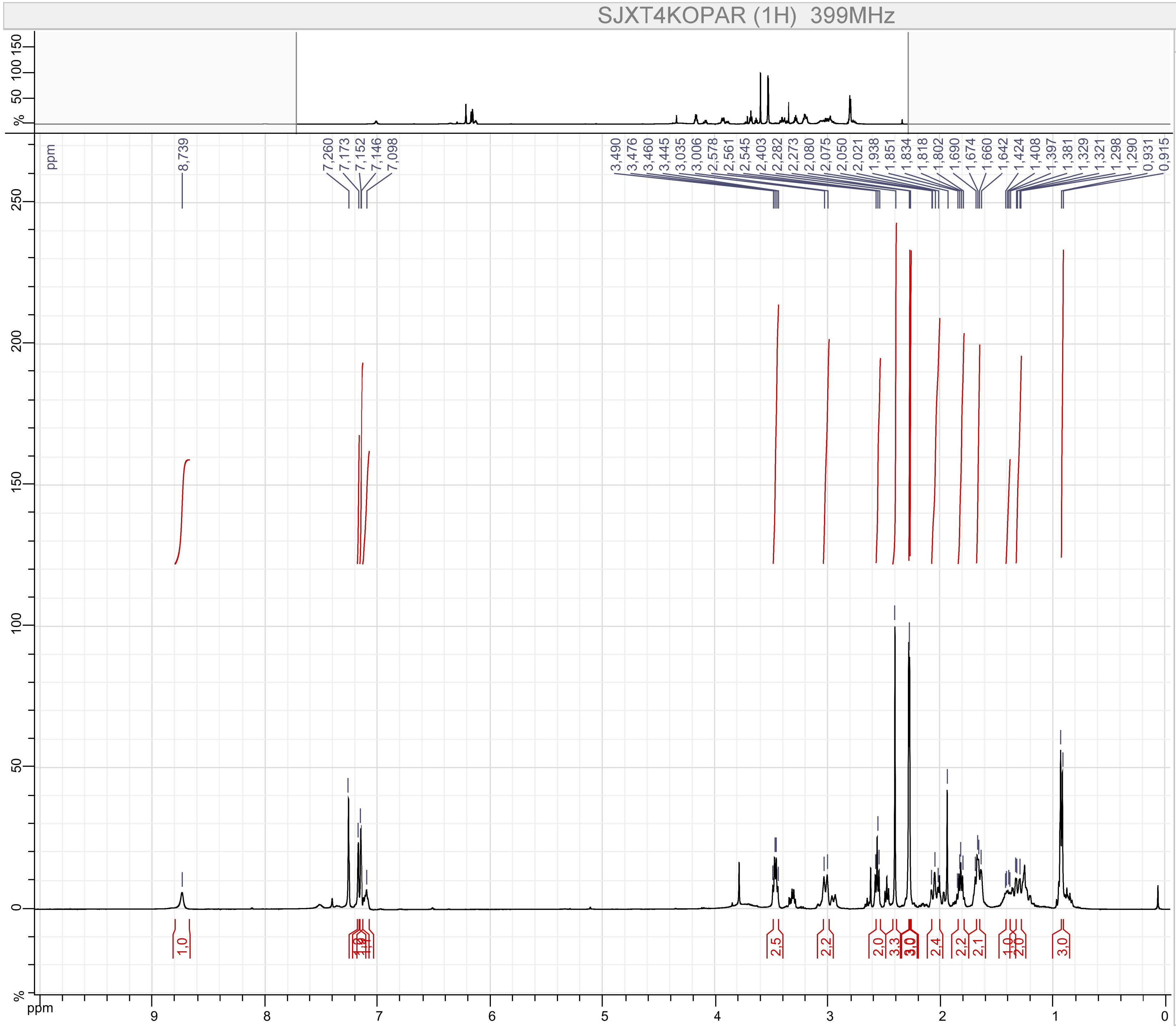


Parameters

SJXT4KOPAR (1H)

F1 :

- Size : 65536 points complex
- Spectral Width : 7494.2334 Hz
- Carrier Frequency : 399.7822 MHz
- Nucleus : 1H

Type : UNKNOWN

Spectro : JEOL 399 MHz

Probe : Unknown

Date : Fri Sep 06 22:04:35 EDT 2019

Temperature : UNKNOWN

Solvent :

Compound 8

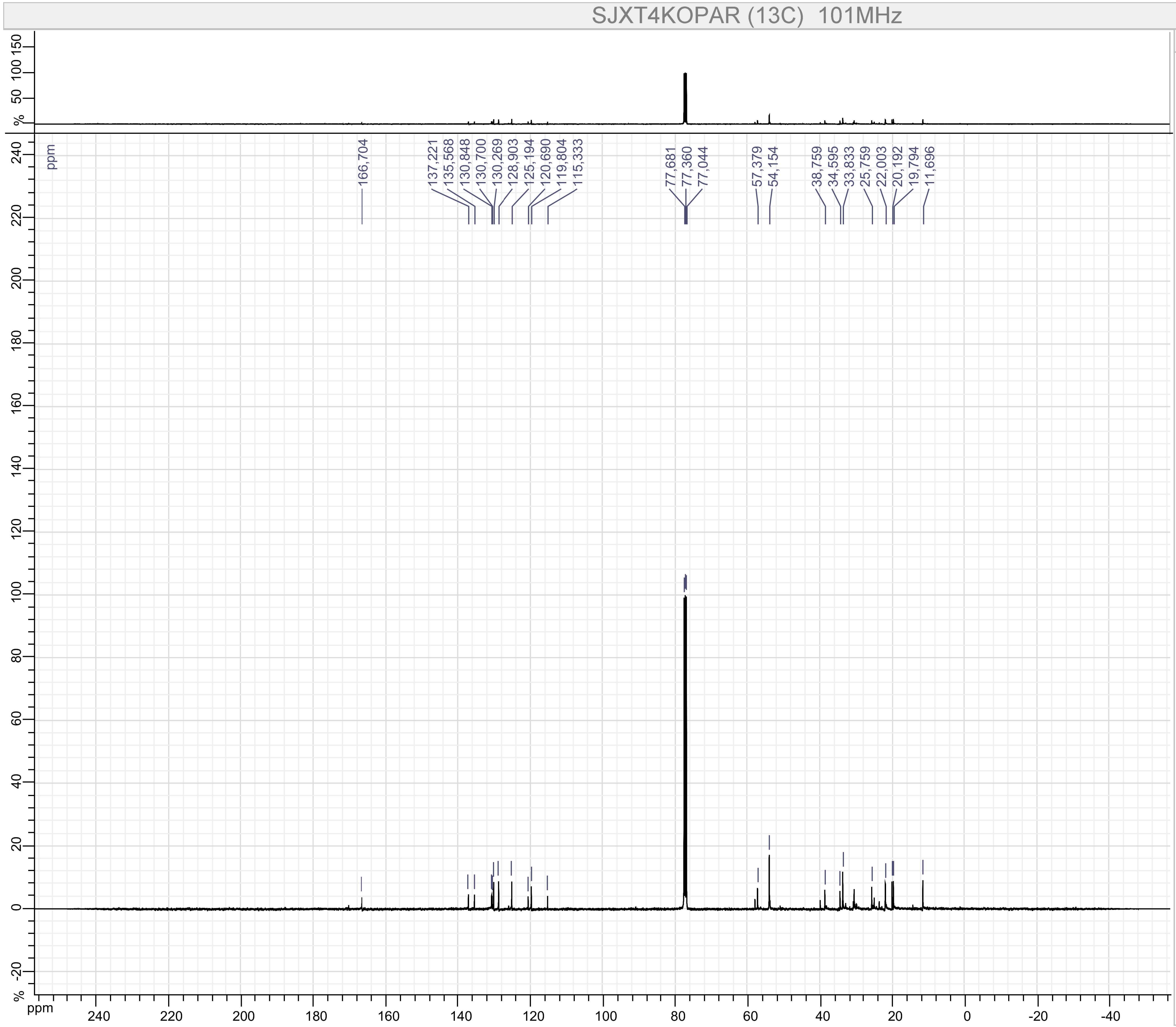


Parameters

SJXT4KOPAR (13C)

F1 :
- Size : 65536 points complex
- Spectral Width : 31566.62 Hz
- Carrier Frequency : 100.52531 MHz
- Nucleus : 13C

Type : UNKNOWN
Spectro : JEOL 101 MHz
Probe : Unknown
Date : Fri Sep 06 22:09:50 EDT 2019
Temperature : UNKNOWN
Solvent :

Compound 9

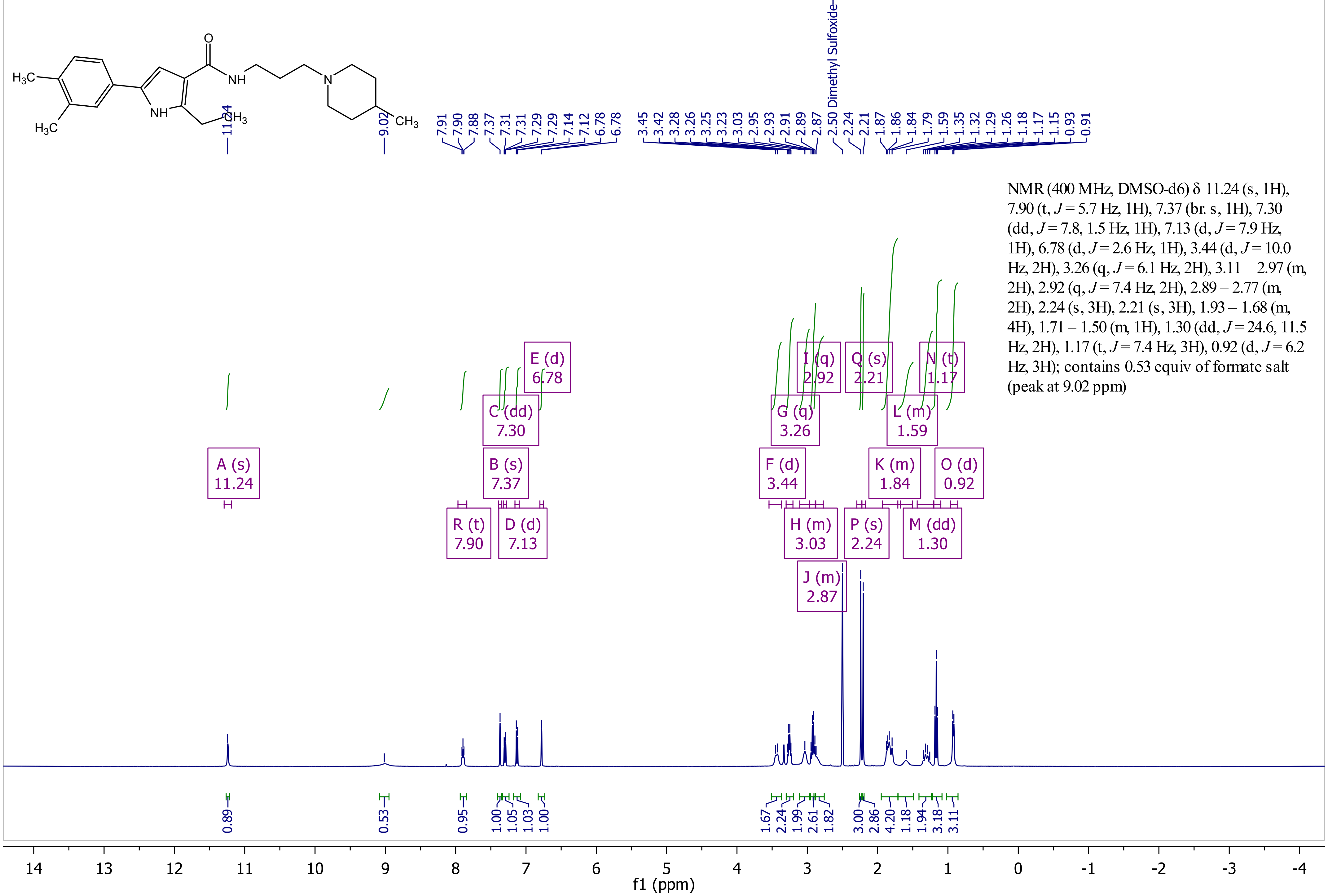


NMR (400 MHz, DMSO-d6) δ 11.24 (s, 1H), 7.90 (t, $J$ = 5.7 Hz, 1H), 7.37 (br. s, 1H), 7.30 (dd, $J$ = 7.8, 1.5 Hz, 1H), 7.13 (d, $J$ = 7.9 Hz, 1H), 6.78 (d, $J$ = 2.6 Hz, 1H), 3.44 (d, $J$ = 10.0 Hz, 2H), 3.26 (q, $J$ = 6.1 Hz, 2H), 3.11 – 2.97 (m, 2H), 2.92 (q, $J$ = 7.4 Hz, 2H), 2.89 – 2.77 (m, 2H), 2.24 (s, 3H), 2.21 (s, 3H), 1.93 – 1.68 (m, 4H), 1.71 – 1.50 (m, 1H), 1.30 (dd, $J$ = 24.6, 11.5 Hz, 2H), 1.17 (t, $J$ = 7.4 Hz, 3H), 0.92 (d, $J$ = 6.2 Hz, 3H); contains 0.53 equiv of formate salt (peak at 9.02 ppm)

Compound 9

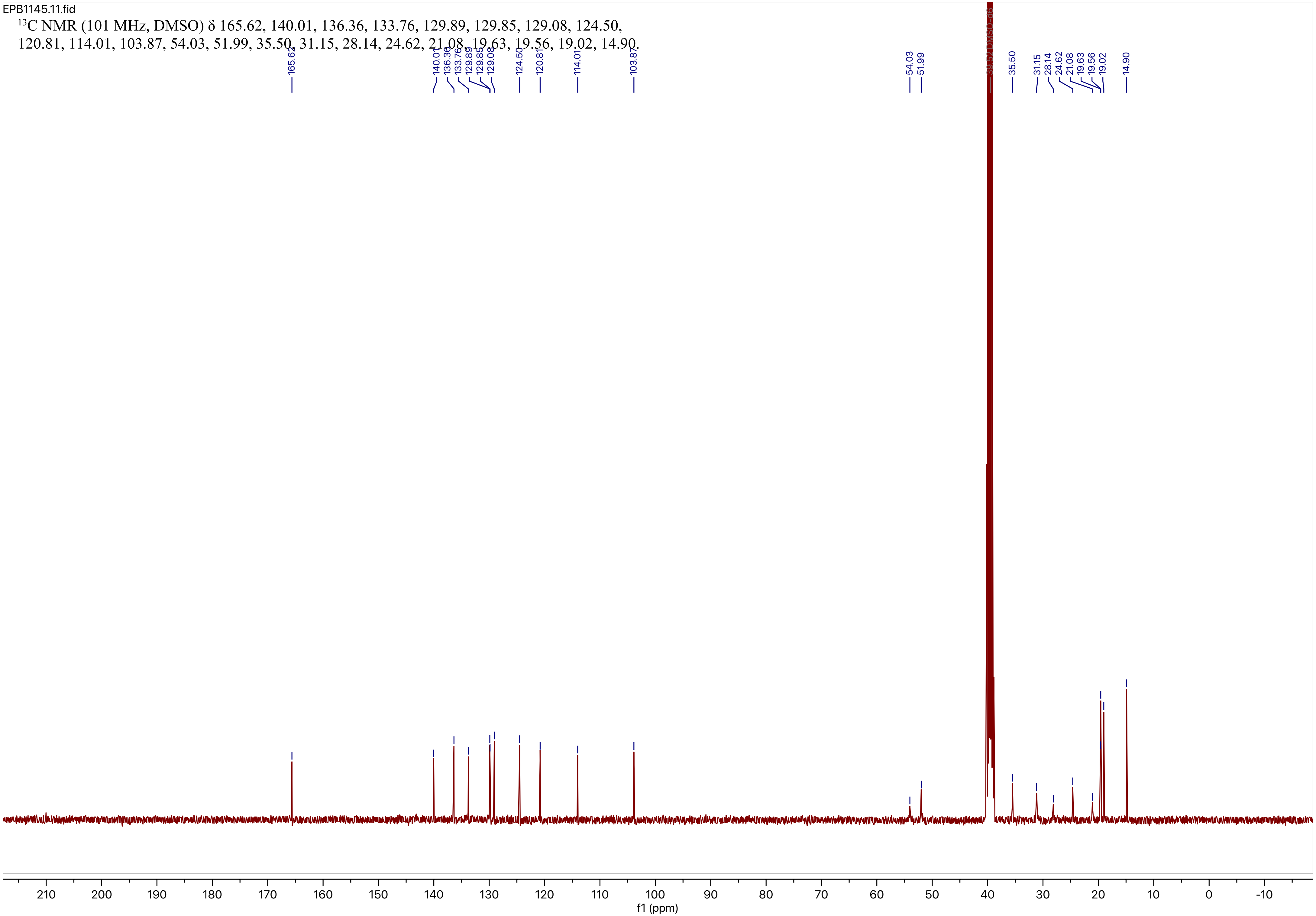
EPB1145.11.fid
$^{13}$C NMR (101 MHz, DMSO) δ 165.62, 140.01, 136.36, 133.76, 129.89, 129.85, 129.08, 124.50, 120.81, 114.01, 103.87, 54.03, 51.99, 35.50, 31.15, 28.14, 24.62, 21.08, 19.63, 19.56, 19.02, 14.90.
f1 (ppm)

Compound 10

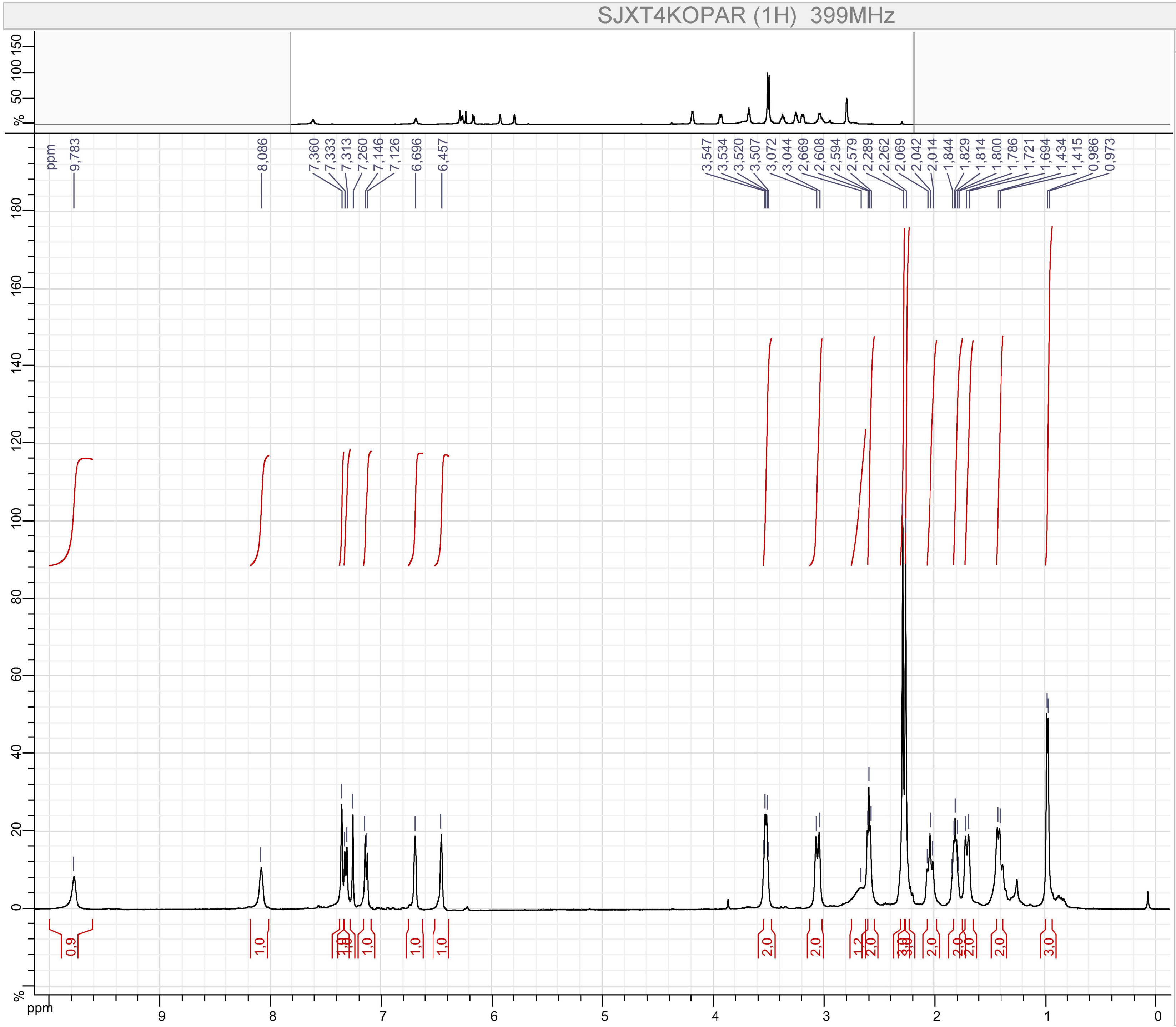


Parameters

SJXT4KOPAR (1H)

F1 :
- Size : 65536 points complex
- Spectral Width : 7494.2334 Hz
- Carrier Frequency : 399.7822 MHz
- Nucleus : 1H

Type : UNKNOWN
Spectro : JEOL 399 MHz
Probe : Unknown
Date : Fri Jul 12 22:04:09 EDT 2019
Temperature : UNKNOWN
Solvent :

Compound 10

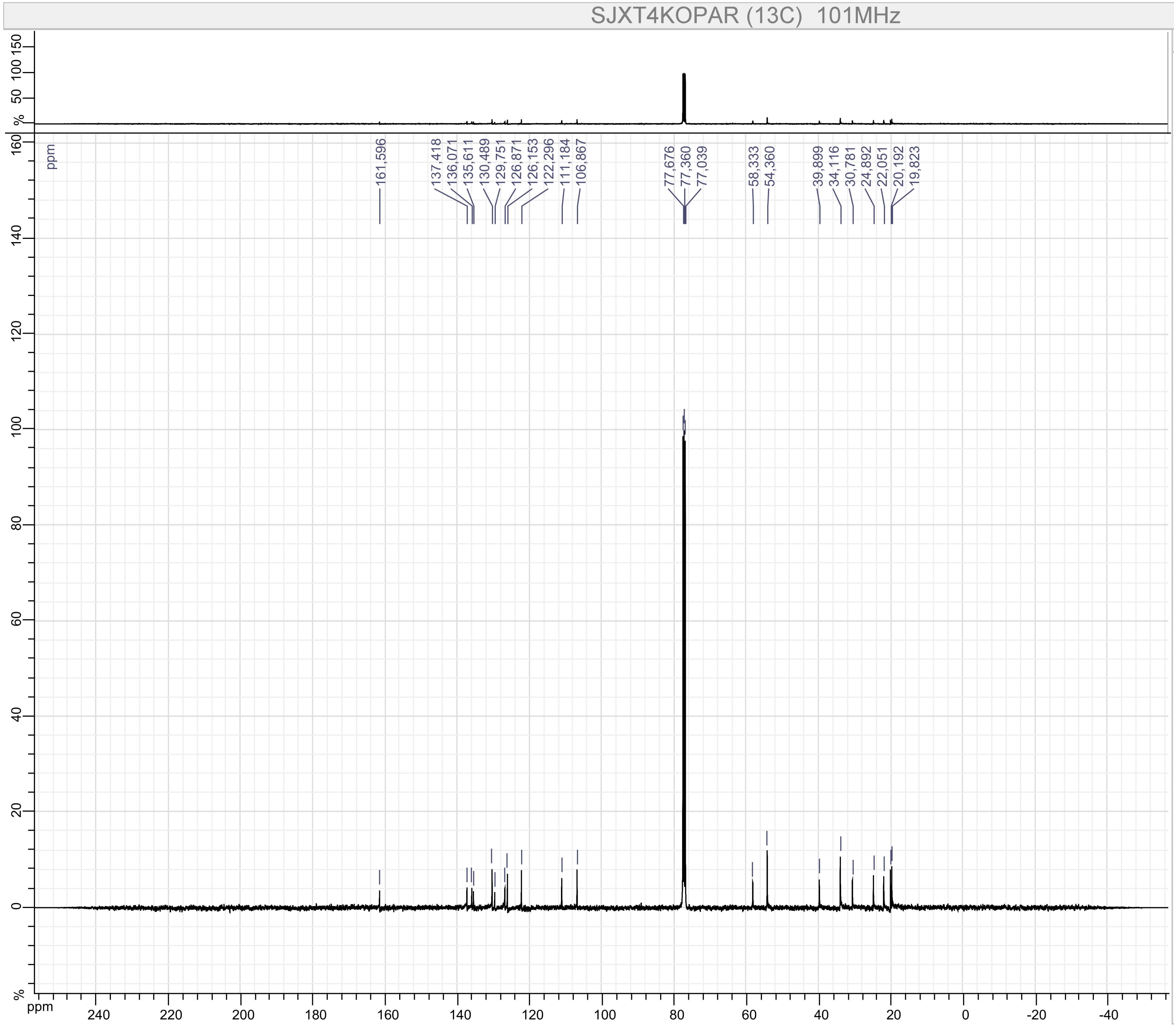


Parameters

SJXT4KOPAR (13C)

F1 :
- Size : 65536 points complex
- Spectral Width : 31566.62 Hz
- Carrier Frequency : 100.52531 MHz
- Nucleus : 13C

Type : UNKNOWN
Spectro : JEOL 101 MHz
Probe : Unknown
Date : Fri Jul 12 22:07:34 EDT 2019
Temperature : UNKNOWN
Solvent :

Compound 11

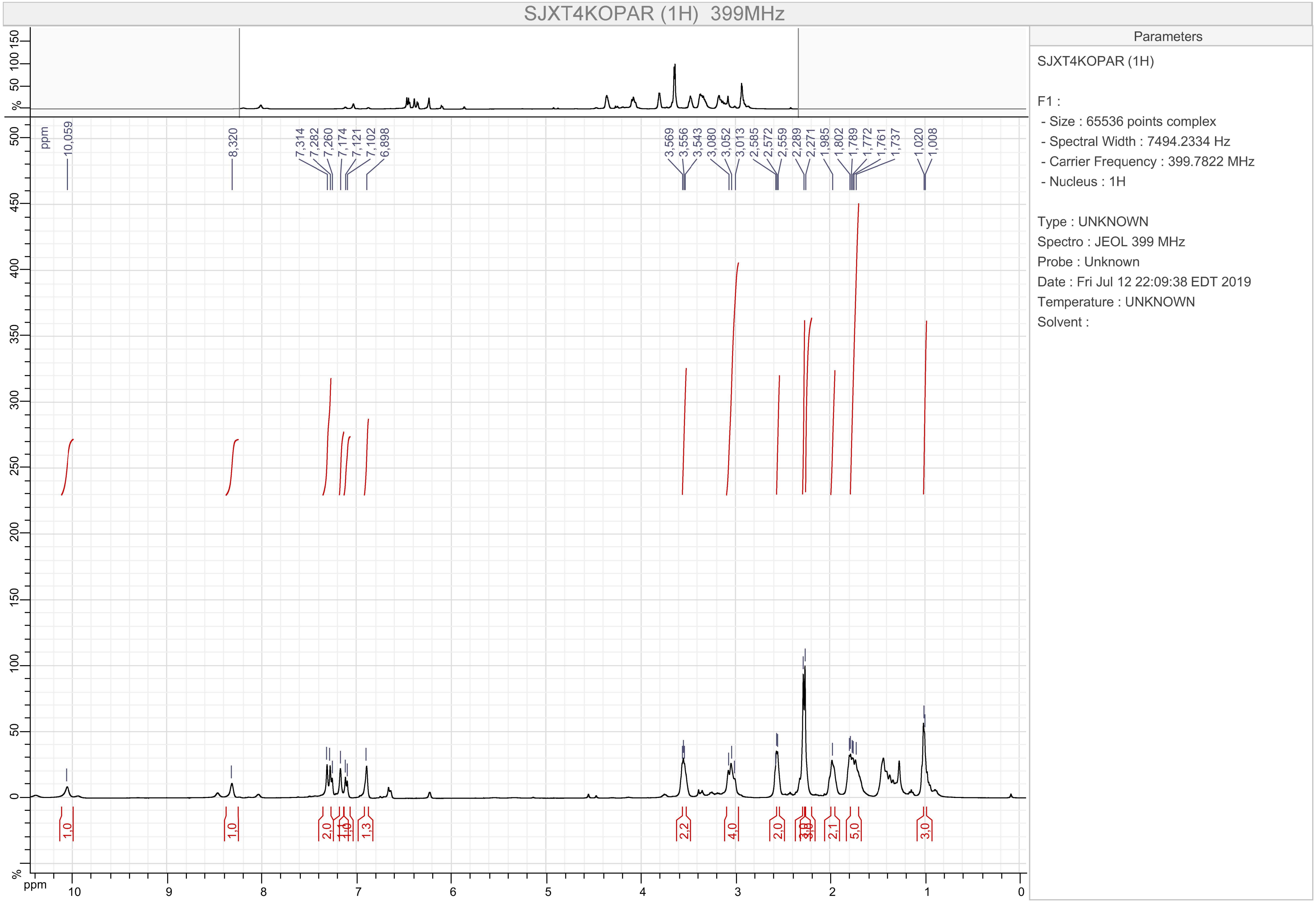
SJXT4KOPAR (1H) 399MHz
Parameters
SJXT4KOPAR (1H)
F1 :
- Size : 65536 points complex
- Spectral Width : 7494.2334 Hz
- Carrier Frequency : 399.7822 MHz
- Nucleus : 1H
Type : UNKNOWN
Spectro : JEOL 399 MHz
Probe : Unknown
Date : Fri Jul 12 22:09:38 EDT 2019
Temperature : UNKNOWN
Solvent :
10,059
8,320
7,314
7,282
7,260
7,174
7,121
7,102
6,898
3,569
3,556
3,543
3,080
3,052
3,013
2,585
2,572
2,559
2,289
2,271
1,985
1,802
1,789
1,772
1,761
1,737
1,020
1,008
1,0
1,0
2,0
1,1
1,0
1,3
2,2
4,0
2,0
3,0
3,0
2,1
5,0
3,0
ppm

Compound 11

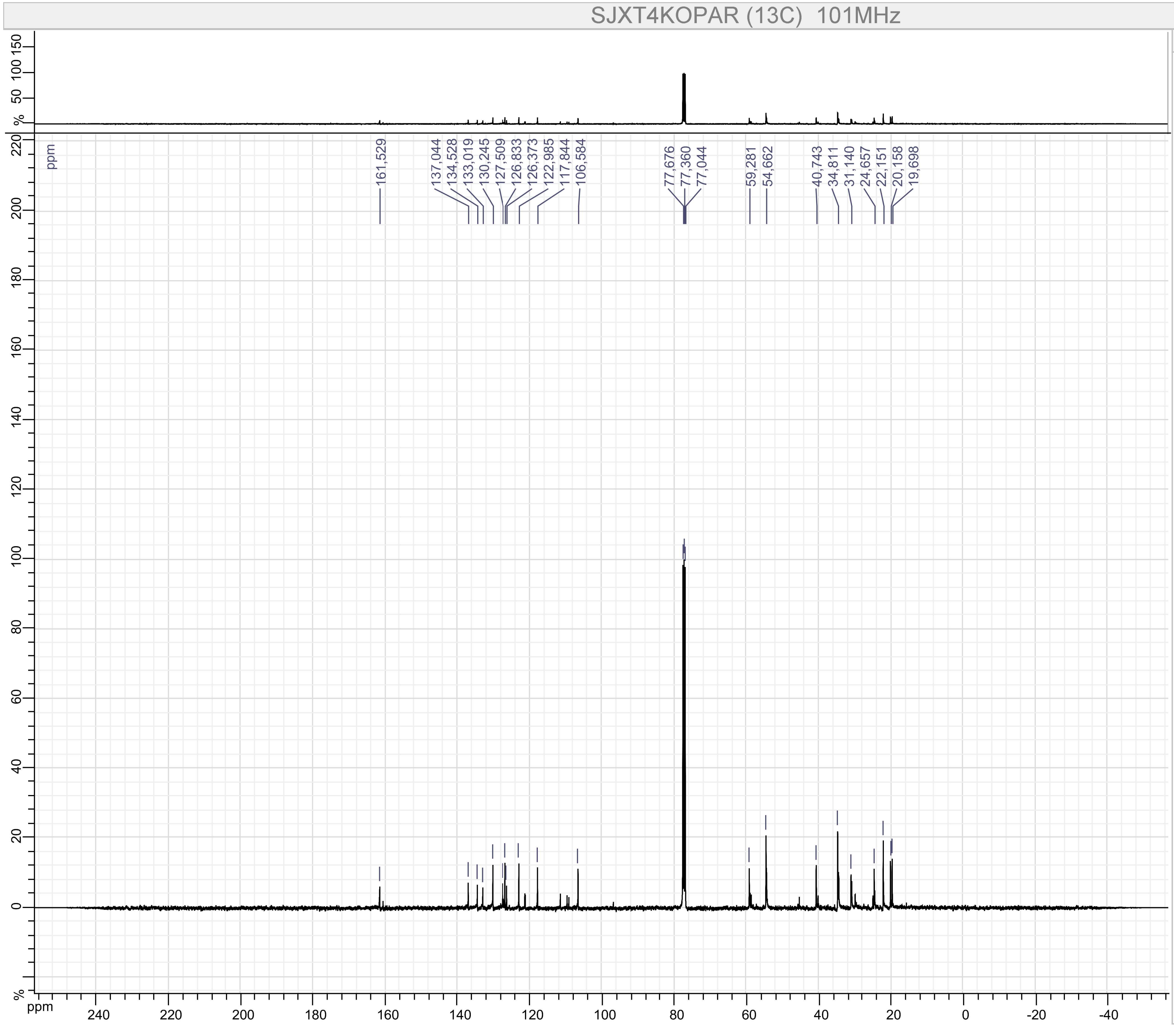


Parameters

SJXT4KOPAR (13C)

F1 :
- Size : 65536 points complex
- Spectral Width : 31566.62 Hz
- Carrier Frequency : 100.52531 MHz
- Nucleus : 13C

Type : UNKNOWN
Spectro : JEOL 101 MHz
Probe : Unknown
Date : Fri Jul 12 22:14:18 EDT 2019
Temperature : UNKNOWN
Solvent :

Compound 12

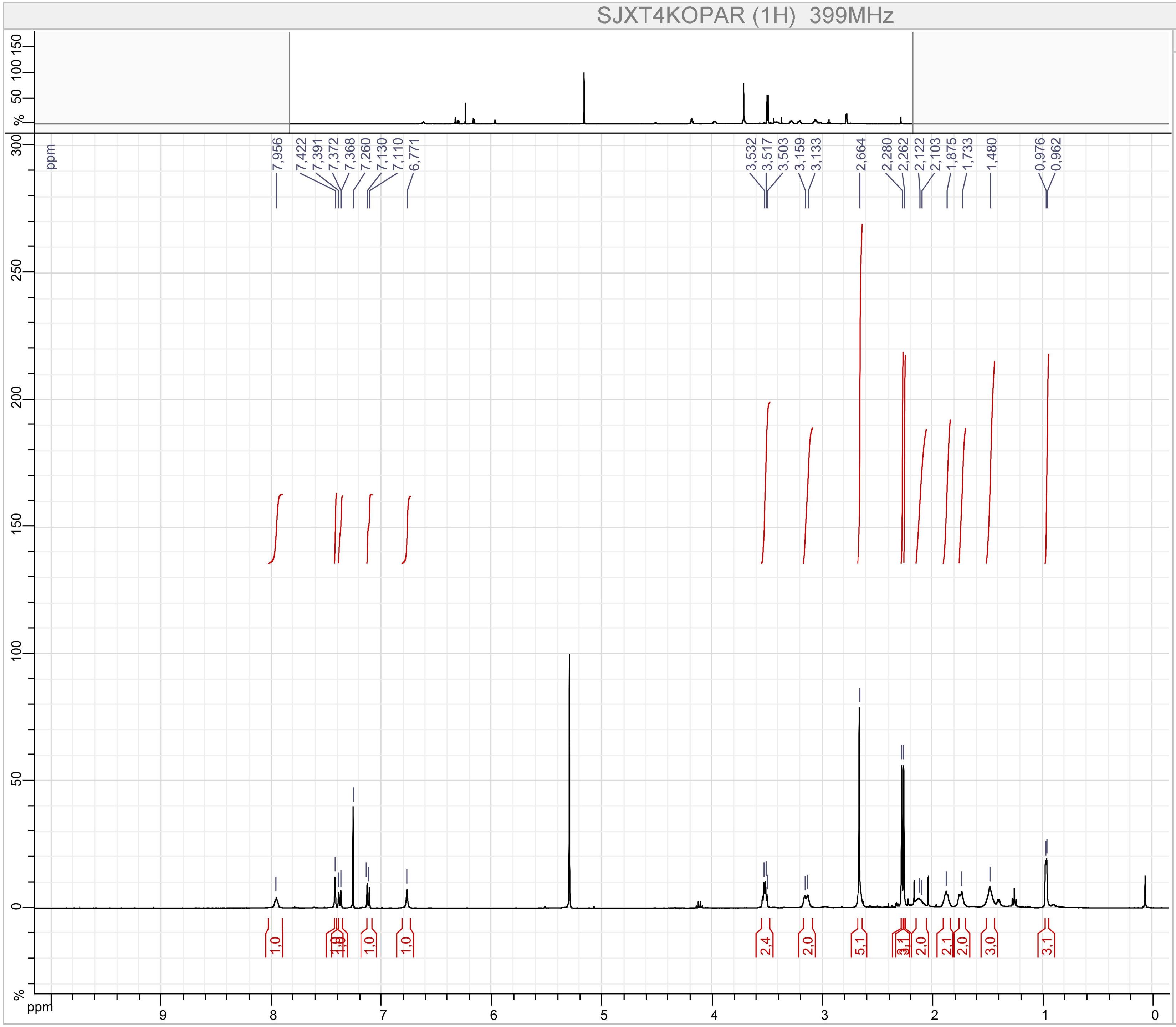


Parameters

SJXT4KOPAR (1H)

F1 :
- Size : 65536 points complex
- Spectral Width : 7494.2334 Hz
- Carrier Frequency : 399.7822 MHz
- Nucleus : 1H

Type : UNKNOWN
Spectro : JEOL 399 MHz
Probe : Unknown
Date : Thu Oct 04 18:15:08 EDT 2018
Temperature : UNKNOWN
Solvent :

Compound 12

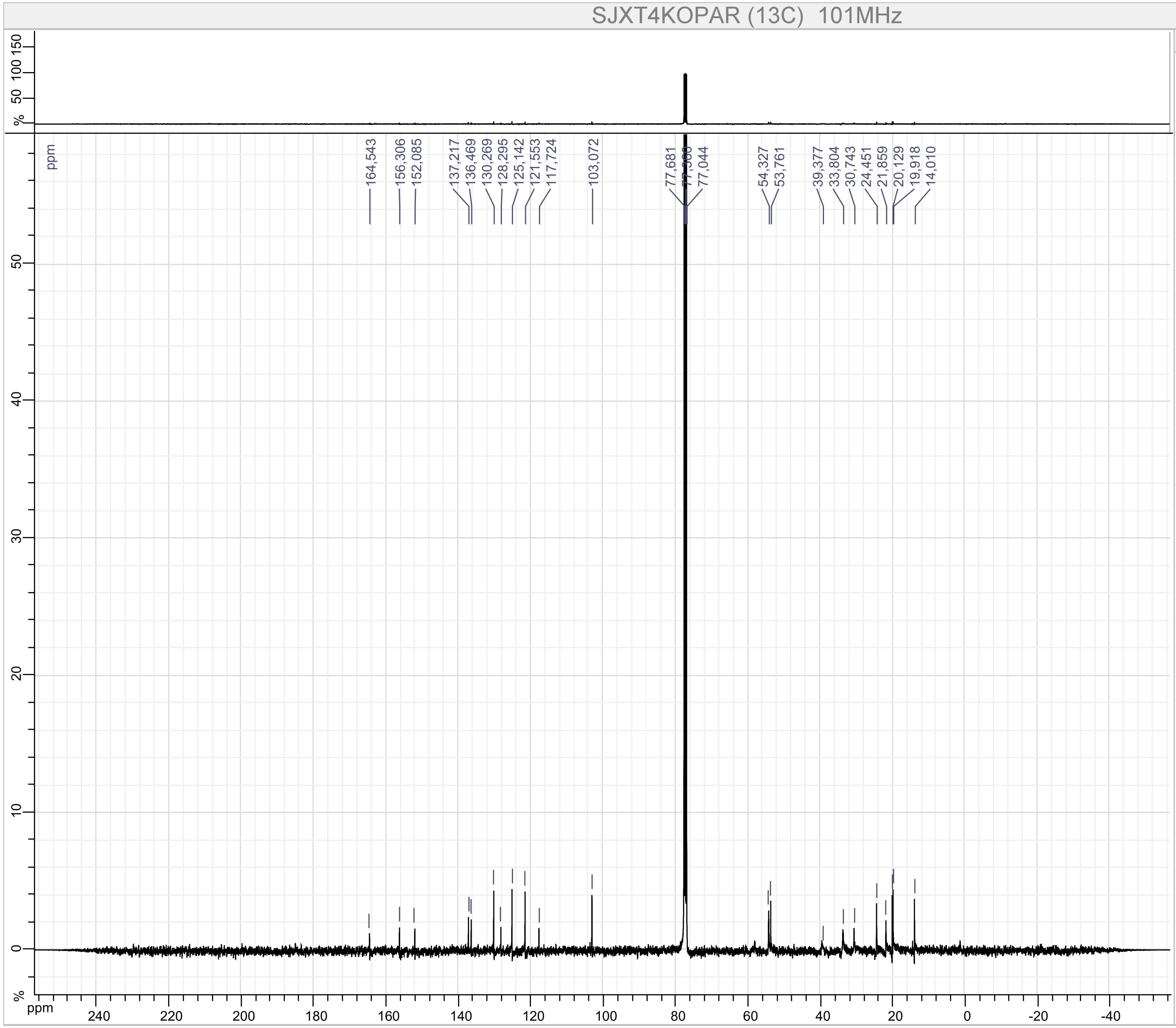


Parameters

SJXT4KOPAR (13C)

F1 :
- Size : 65536 points complex
- Spectral Width : 31566.62 Hz
- Carrier Frequency : 100.52531 MHz
- Nucleus : 13C

Type : UNKNOWN
Spectro : JEOL 101 MHz
Probe : Unknown
Date : Sun Oct 14 22:20:05 EDT 2018
Temperature : UNKNOWN
Solvent :

Compound 13

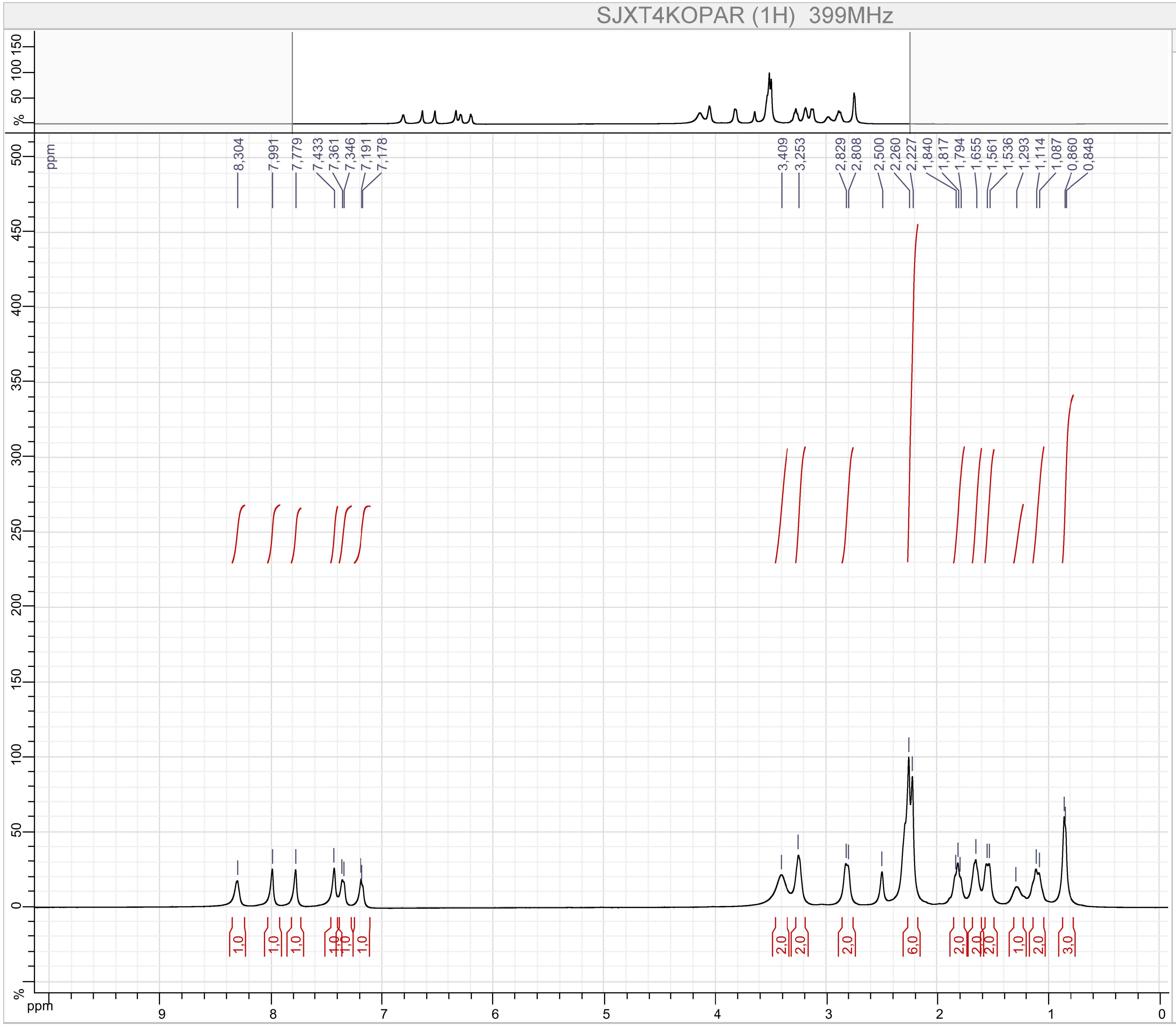


Parameters

SJXT4KOPAR (1H)

F1 :
- Size : 65536 points complex
- Spectral Width : 7494.2334 Hz
- Carrier Frequency : 399.7822 MHz
- Nucleus : 1H

Type : UNKNOWN
Spectro : JEOL 399 MHz
Probe : Unknown
Date : Mon Nov 05 21:11:32 EST 2018
Temperature : UNKNOWN
Solvent :

Compound 13

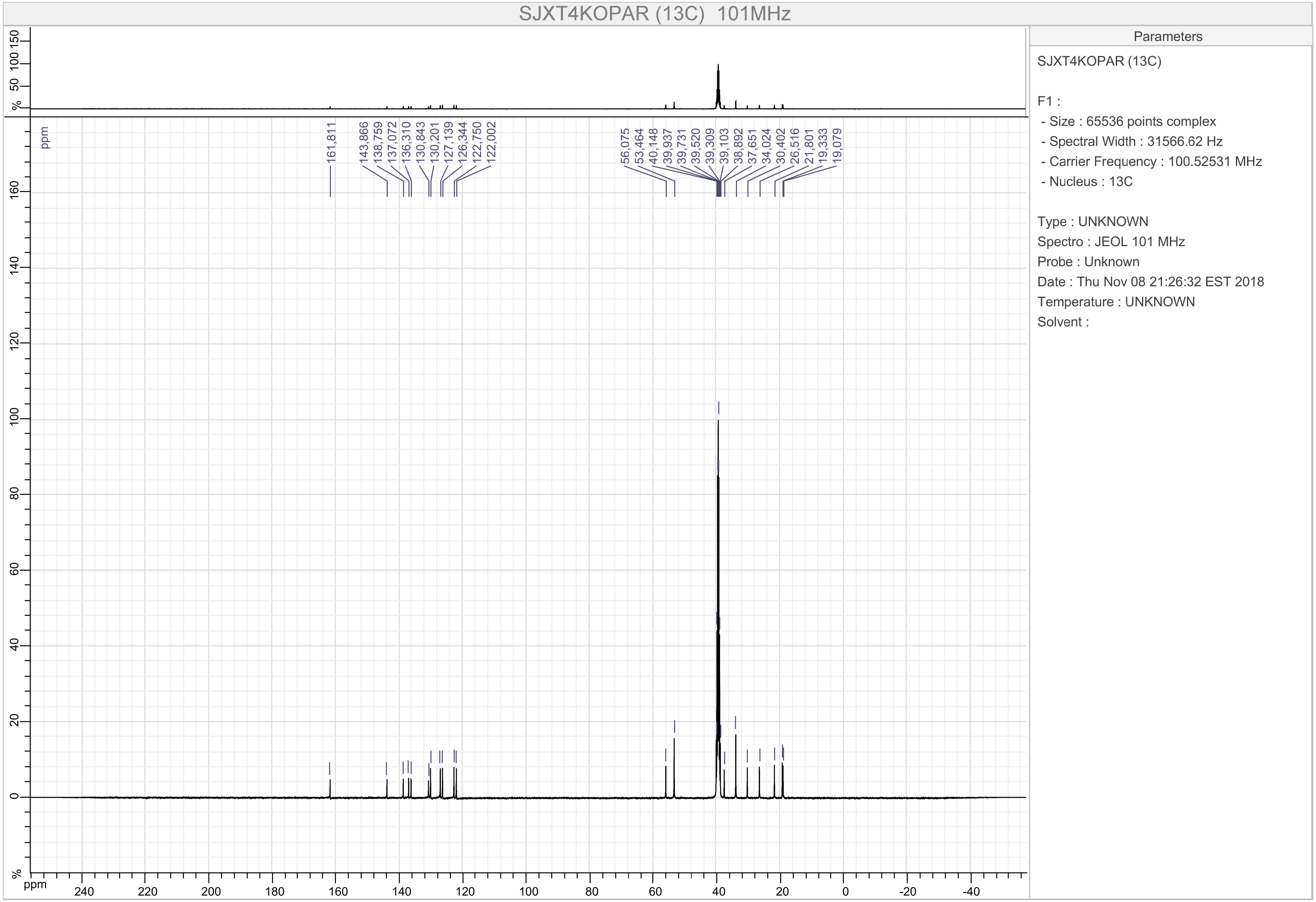
SJXT4KOPAR (13C) 101MHz
161,811
143,866
138,759
137,072
136,310
130,843
130,201
127,139
126,344
122,750
122,002
56,075
53,464
40,148
39,937
39,731
39,520
39,309
39,103
38,892
37,651
34,024
30,402
26,516
21,801
19,333
19,079
ppm
Parameters
SJXT4KOPAR (13C)
F1 :
- Size : 65536 points complex
- Spectral Width : 31566.62 Hz
- Carrier Frequency : 100.52531 MHz
- Nucleus : 13C
Type : UNKNOWN
Spectro : JEOL 101 MHz
Probe : Unknown
Date : Thu Nov 08 21:26:32 EST 2018
Temperature : UNKNOWN
Solvent :

Compound 14

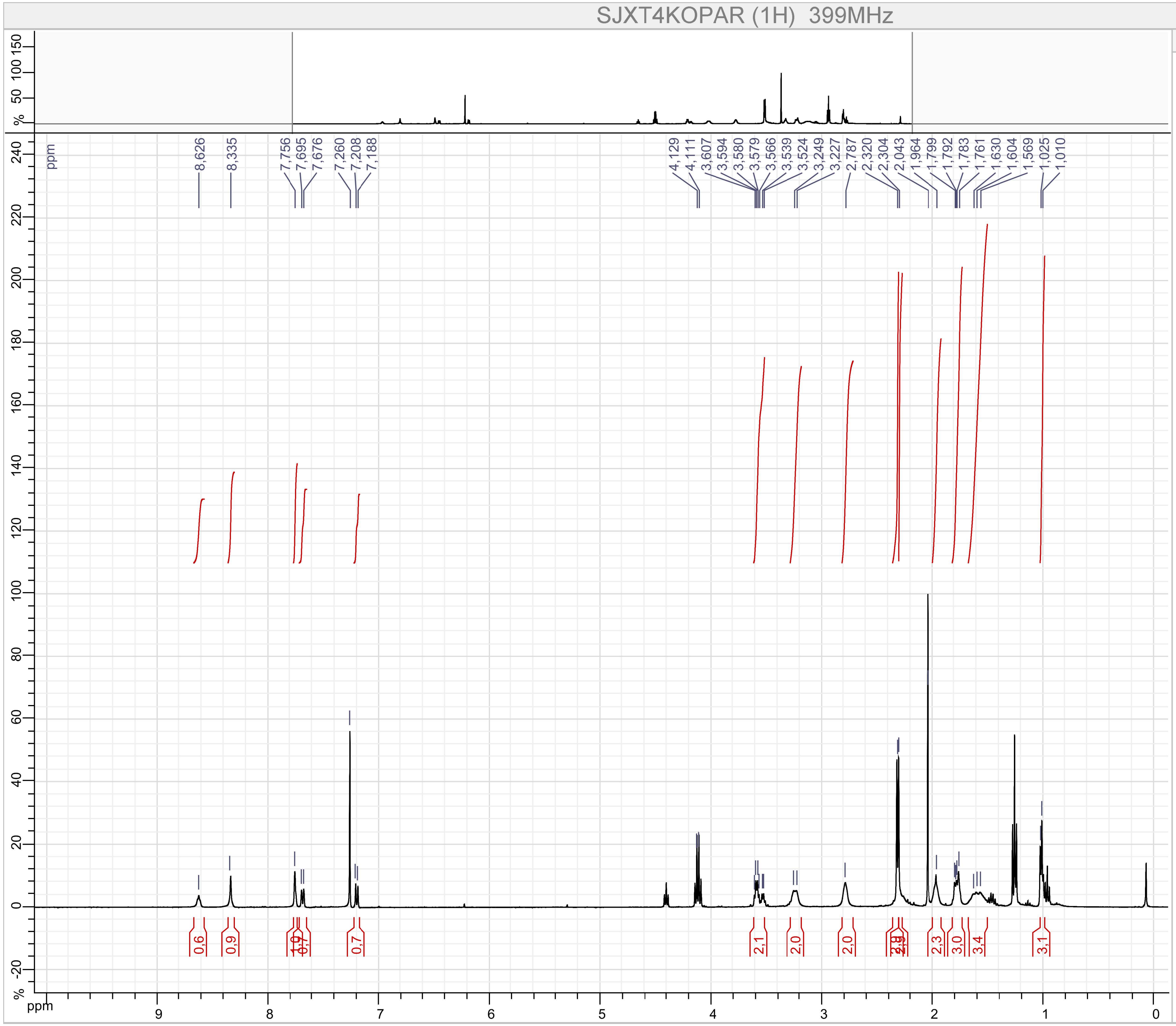


Parameters

SJXT4KOPAR (1H)

F1 :

- Size : 65536 points complex
- Spectral Width : 7494.2334 Hz
- Carrier Frequency : 399.7822 MHz
- Nucleus : 1H

Type : UNKNOWN

Spectro : JEOL 399 MHz

Probe : Unknown

Date : Thu Oct 04 18:30:32 EDT 2018

Temperature : UNKNOWN

Solvent :

Compound 14

MB-A-039

$^{13}$C NMR (101 MHz, CHLOROFORM-*D*) δ 171.30, 161.14, 144.87, 140.11, 137.60, 133.54, 131.05, 130.42, 127.94, 124.46, 77.48, 77.36, 77.16, 76.84, 72.20, 60.54, 30.91, 23.73, 21.19, 19.97, 19.84, 14.34, 13.82.

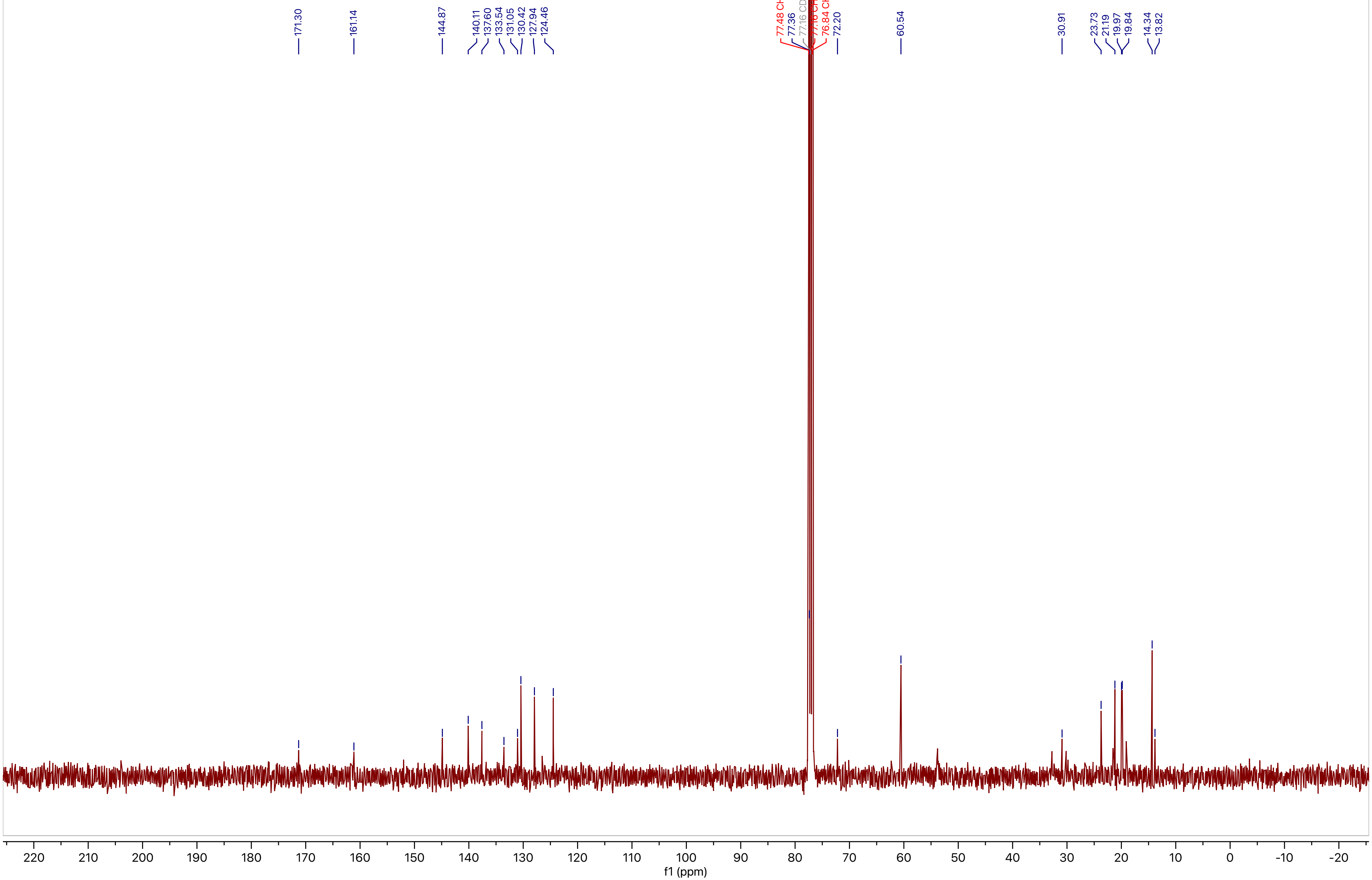

Compound 15

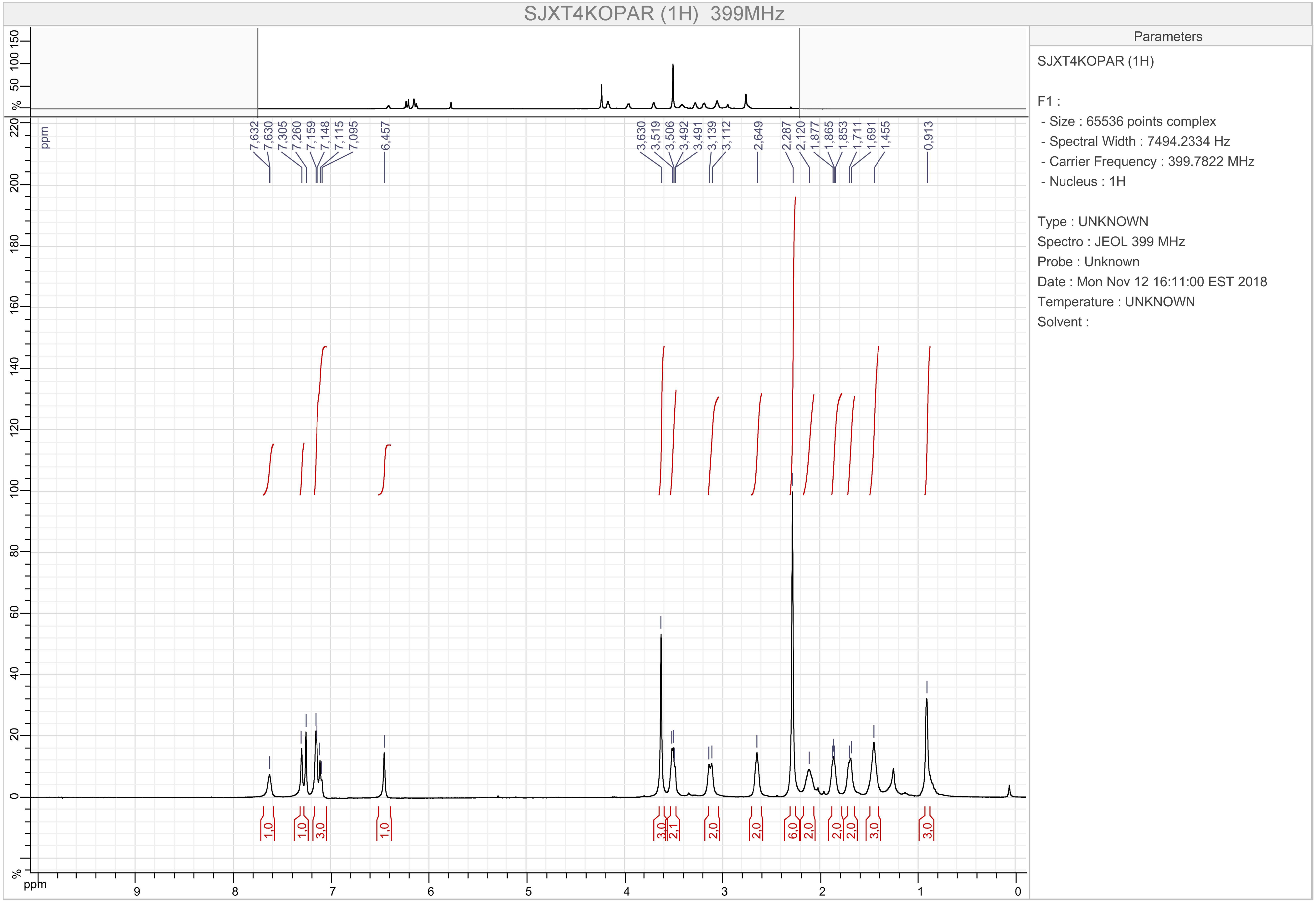

SJXT4KOPAR (1H) 399MHz
7,632 7,630 7,305 7,260 7,159 7,148 7,115 7,095 6,457 3,630 3,519 3,506 3,492 3,491 3,139 3,112 2,649 2,287 2,120 1,877 1,865 1,853 1,711 1,691 1,455 0,913
1,0 1,0 3,0 1,0 3,0 2,1 2,0 2,0 6,0 2,0 2,0 2,0 3,0 3,0
ppm
Parameters
SJXT4KOPAR (1H)
F1 :
- Size : 65536 points complex
- Spectral Width : 7494.2334 Hz
- Carrier Frequency : 399.7822 MHz
- Nucleus : 1H
Type : UNKNOWN
Spectro : JEOL 399 MHz
Probe : Unknown
Date : Mon Nov 12 16:11:00 EST 2018
Temperature : UNKNOWN
Solvent :

Compound 15

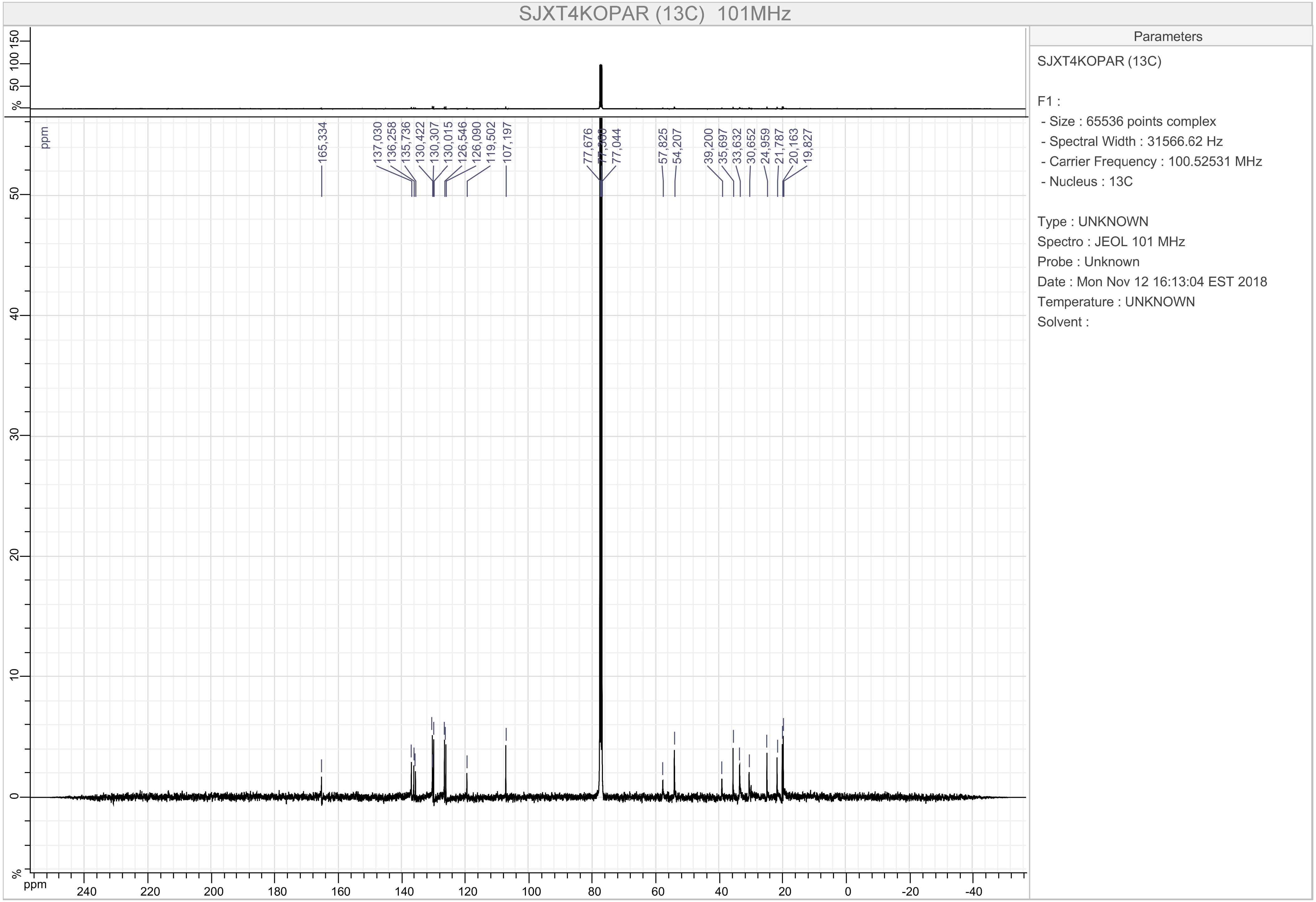

SJXT4KOPAR (13C) 101MHz
165,334
137,030
136,258
135,736
130,422
130,307
130,015
126,546
126,090
119,502
107,197
77,676
77,044
57,825
54,207
39,200
35,697
33,632
30,652
24,959
21,787
20,163
19,827
ppm
Parameters
SJXT4KOPAR (13C)
F1 :
- Size : 65536 points complex
- Spectral Width : 31566.62 Hz
- Carrier Frequency : 100.52531 MHz
- Nucleus : 13C
Type : UNKNOWN
Spectro : JEOL 101 MHz
Probe : Unknown
Date : Mon Nov 12 16:13:04 EST 2018
Temperature : UNKNOWN
Solvent :

Compound 16

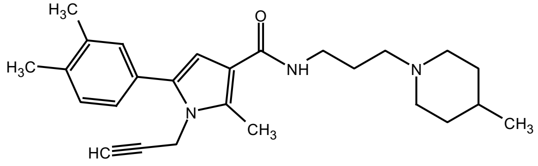

NMR (400 MHz, CDCl3 ) δ 7.55 (app. br. s, 1H), 7.20 (s, 1H), 7.18 – 7.10 (m, 2H), 6.66 (t, *J* = 6.5 Hz, 1H), 6.36 (s, 1H), 5.21 (d, *J* = 6.5 Hz, 2H), 3.49 (dd, *J* = 11.6, 5.4 Hz, 2H), 3.00 (d, *J* = 11.2 Hz, 2H), 2.67 (s, 3H), 2.51 (t, *J* = 5.8 Hz, 2H), 2.28 (s, 6H), 1.93 (t, *J* = 9.8 Hz, 2H), 1.77 (dt, *J* = 11.6, 5.8 Hz, 2H), 1.63 (d, *J* = 10.9 Hz, 2H), 1.45 – 1.07 (m, 3H), 0.82 (d, *J* = 5.9 Hz, 3H).

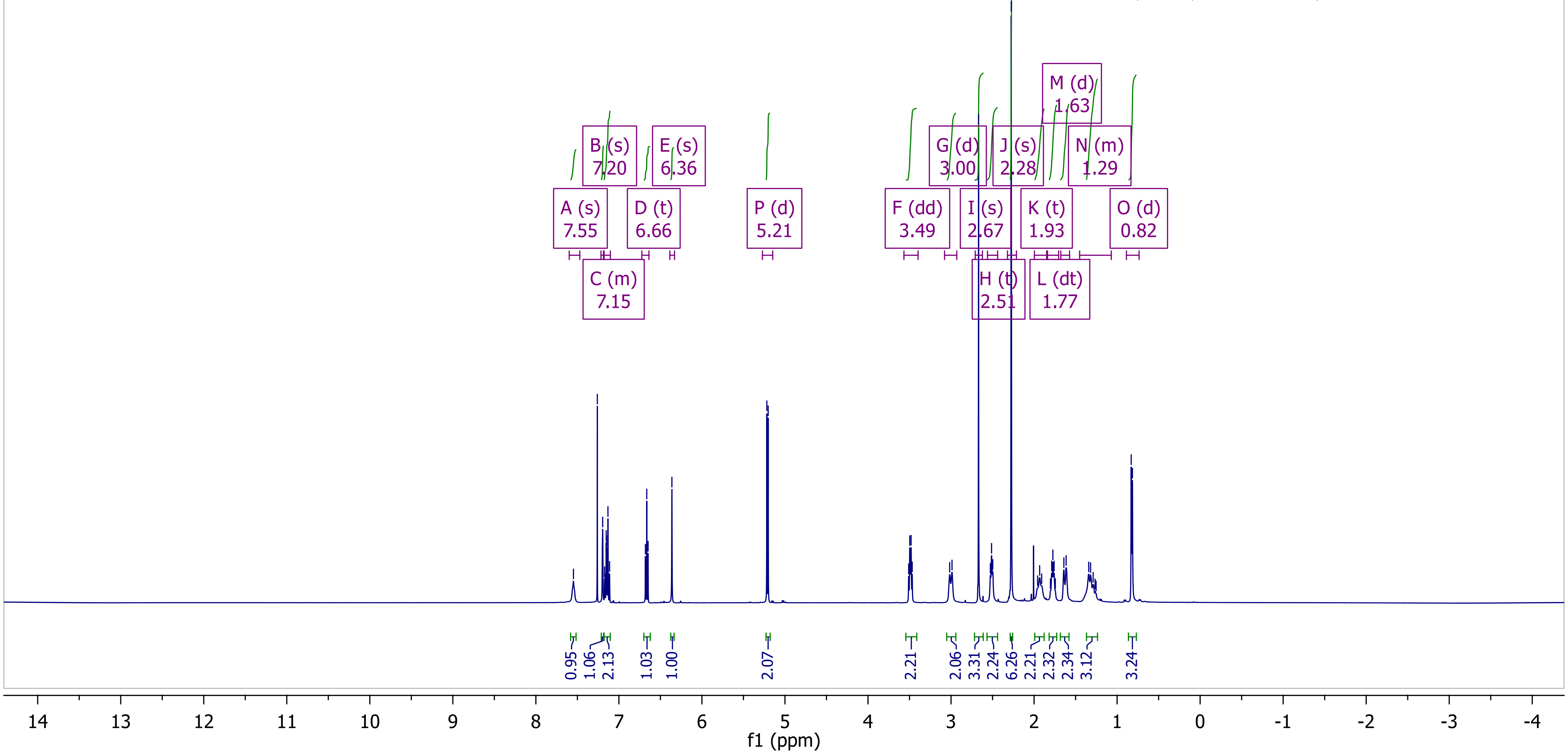

Compound 16

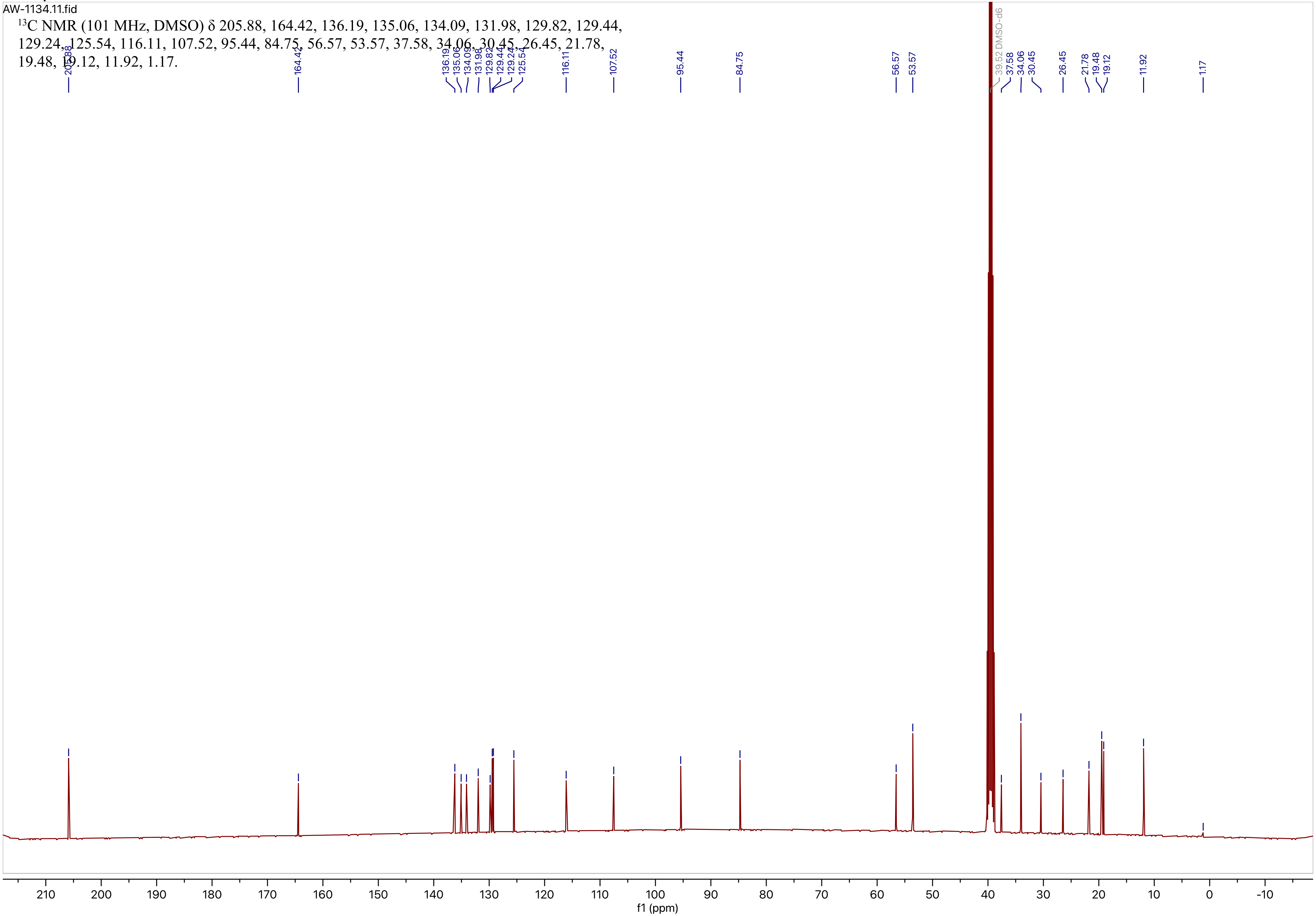

AW-1134.11.fid
$^{13}$C NMR (101 MHz, DMSO) δ 205.88, 164.42, 136.19, 135.06, 134.09, 131.98, 129.82, 129.44, 129.24, 125.54, 116.11, 107.52, 95.44, 84.75, 56.57, 53.57, 37.58, 34.06, 30.45, 26.45, 21.78, 19.48, 19.12, 11.92, 1.17.
39.52 DMSO-d6
f1 (ppm)

Compound 17

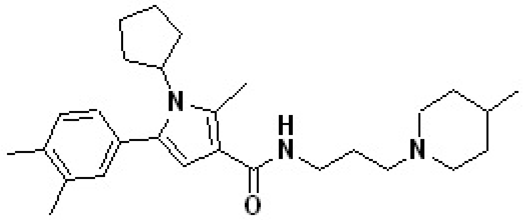

[1]H NMR (500 MHz, $CDCl_3$) δ 7.30 (s, 1H), 7.13 (d, $J$ = 7.7 Hz, 1H), 7.09 (s, 1H), 7.04 (dd, $J$ = 7.7, 1.6 Hz, 1H), 6.21 (s, 1H), 4.62 (p, $J$ = 9.3 Hz, 1H), 3.46 (dd, $J$ = 11.9, 5.4 Hz, 2H), 3.02 (d, $J$ = 11.5 Hz, 2H), 2.72 (s, 3H), 2.54 (t, $J$ = 6.3 Hz, 2H), 2.29 (s, 3H), 2.28 (s, 3H), 2.09 – 1.92 (m, 6H), 1.90 – 1.81 (m, 2H), 1.78 (dt, $J$ = 12.4, 6.3 Hz, 2H), 1.67 – 1.50 (m, 4H), 1.44 – 1.23 (m, 3H), 0.79 (d, $J$ = 6.0 Hz, 3H).

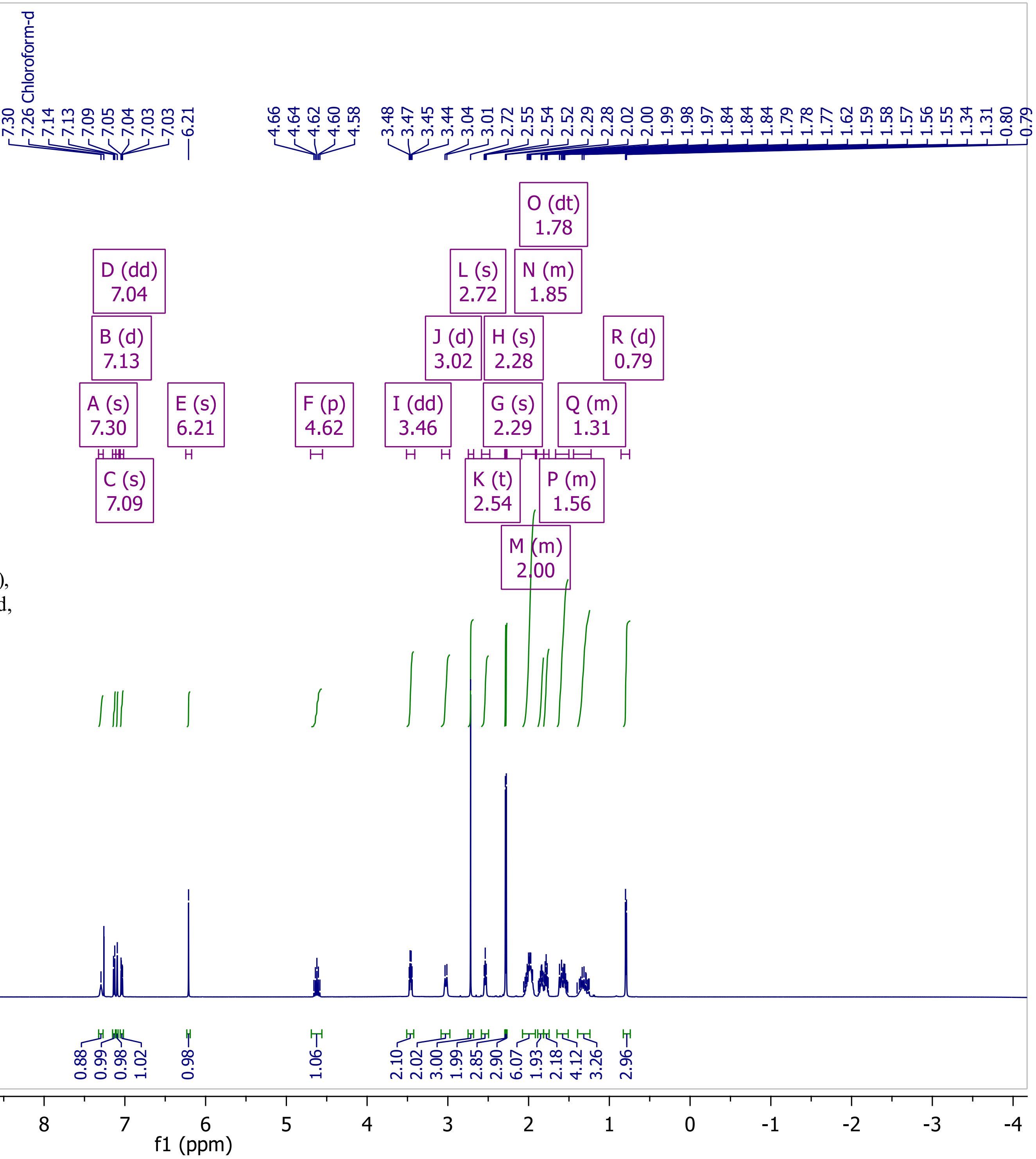

Compound 17

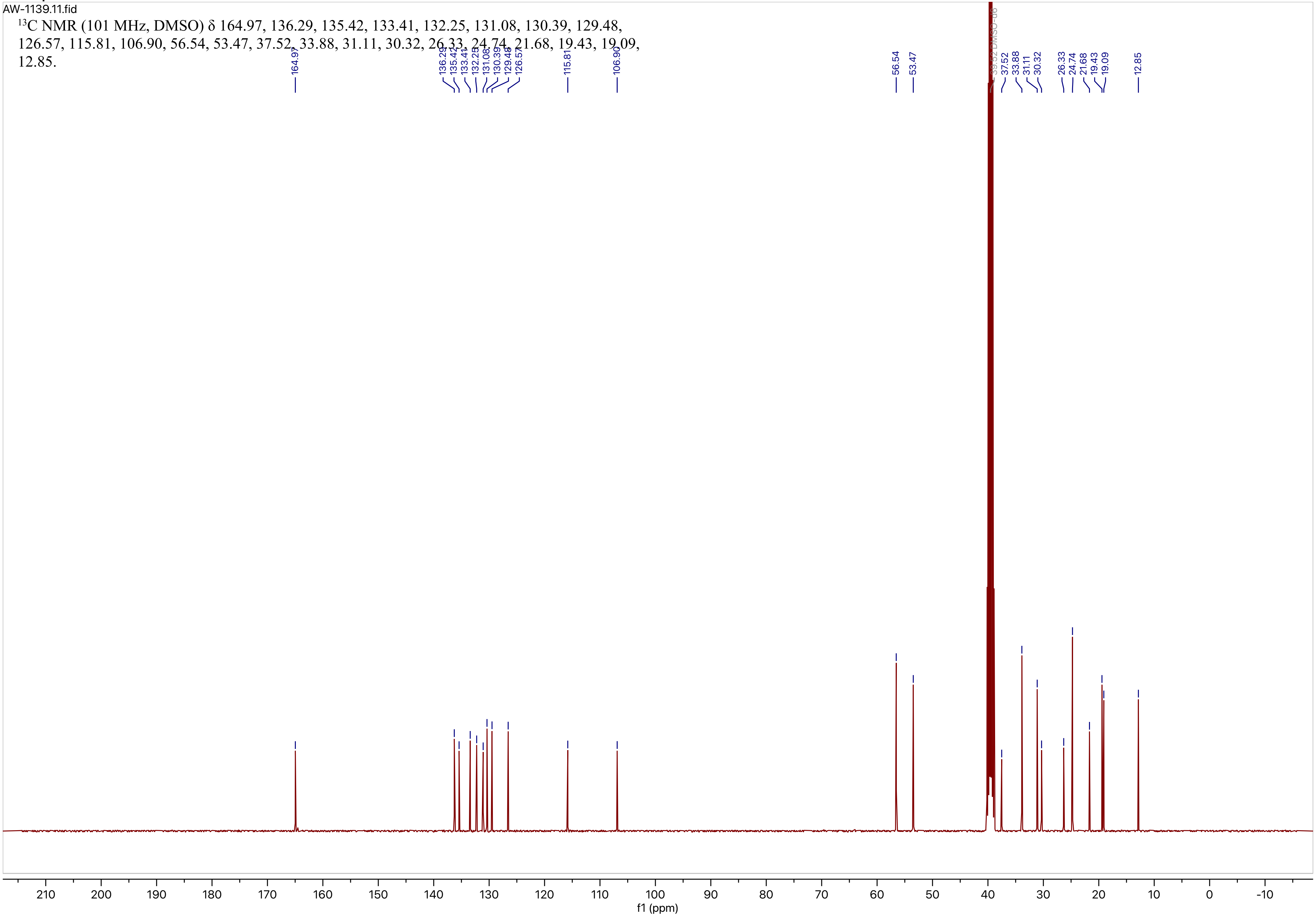
AW-1139.11.fid
$^{13}$C NMR (101 MHz, DMSO) δ 164.97, 136.29, 135.42, 133.41, 132.25, 131.08, 130.39, 129.48, 126.57, 115.81, 106.90, 56.54, 53.47, 37.52, 33.88, 31.11, 30.32, 26.33, 24.74, 21.68, 19.43, 19.09, 12.85.
164.97
136.29
135.42
133.41
132.25
131.08
130.39
129.48
126.57
115.81
106.90
56.54
53.47
39.52 DMSO-d6
37.52
33.88
31.11
30.32
26.33
24.74
21.68
19.43
19.09
12.85
210
200
190
180
170
160
150
140
130
120
110
100
90
80
70
60
50
40
30
20
10
0
-10
f1 (ppm)

Compound 18

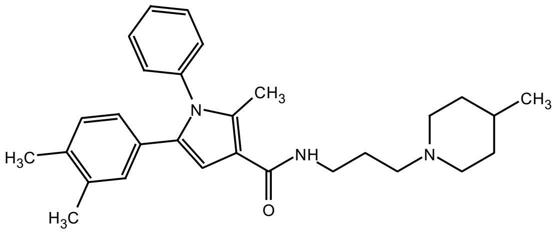


NMR (400 MHz, DMSO-d6) δ 8.02 (t, $J$ = 5.8 Hz, 1H), 7.52 – 7.39 (m, 3H), 7.24 – 7.13 (m, 2H), 6.91 – 6.89 ( m, 1H), 6.88 (s, 1H), 6.77 (overlapping s, 1H), 6.61 (dd, $J$ = 7.8, 1.7 Hz, 1H), 3.49 – 3.31 (m, 2H), 3.28 (dd, $J$ = 12.4, 6.3 Hz, 2H), 3.07 – 2.89 (m, 2H), 2.92 – 2.62 (m, 2H), 2.30 (s, 3H), 2.11 (s, 3H), 2.07 (s, 3H), 1.90 – 1.82 (m, 2H), 1.82 – 1.74 (m, 2H), 1.69 – 1.47 (m, 1H), 1.44 – 1.18 (m, 2H), 0.92 (d, $J$ = 6.5 Hz, 3H); contains 0.06 equiv of formate salt (8.14 ppm).

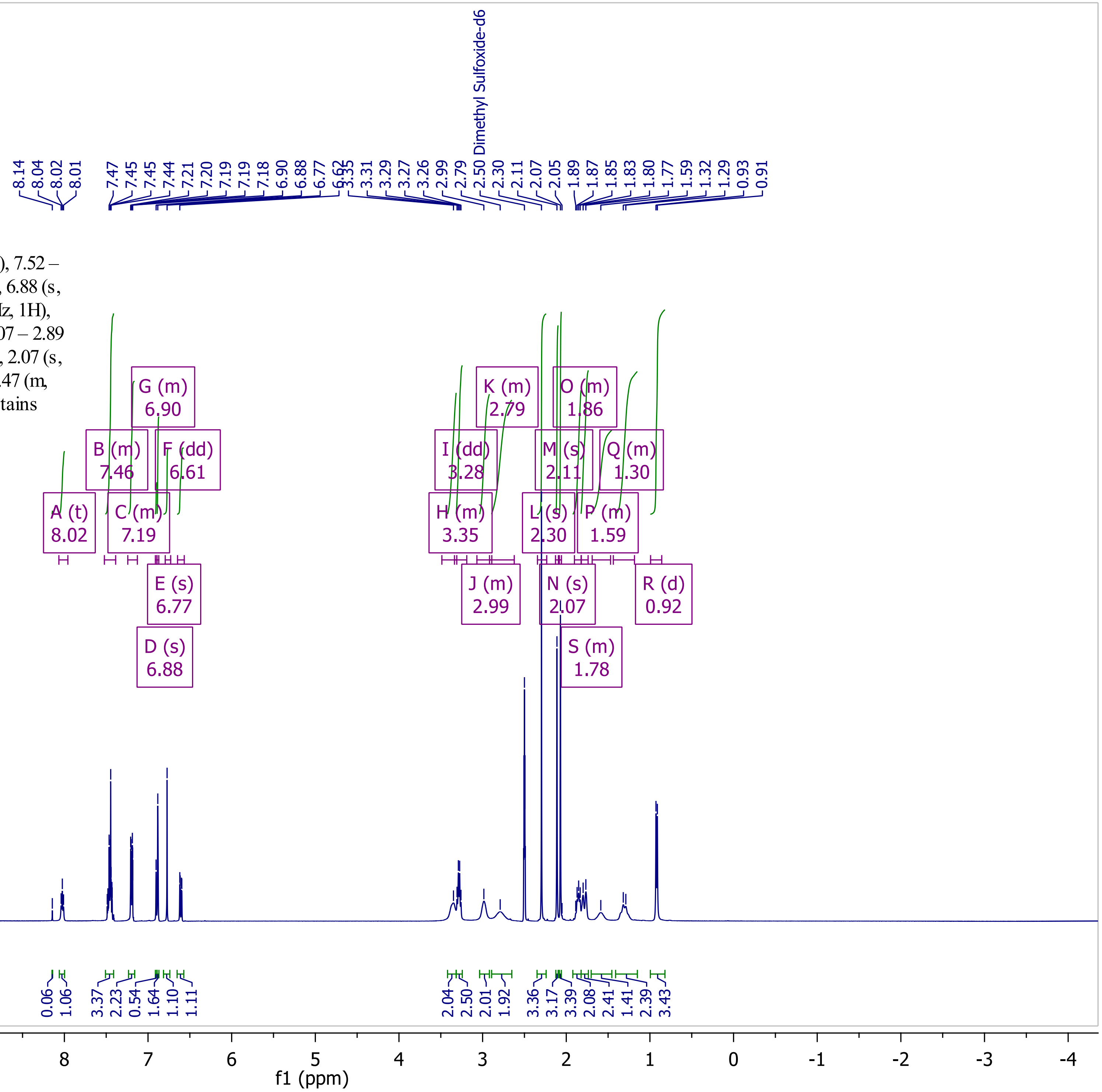

Compound 18

EPB1132.11.fid

$^{13}$C NMR (101 MHz, DMSO) δ 165.21, 137.84, 135.89, 134.76, 134.42, 132.70, 129.77, 129.30, 129.17, 128.67, 128.51, 128.29, 124.75, 115.19, 107.21, 52.13, 35.98, 24.90, 19.38, 18.89, 11.96.

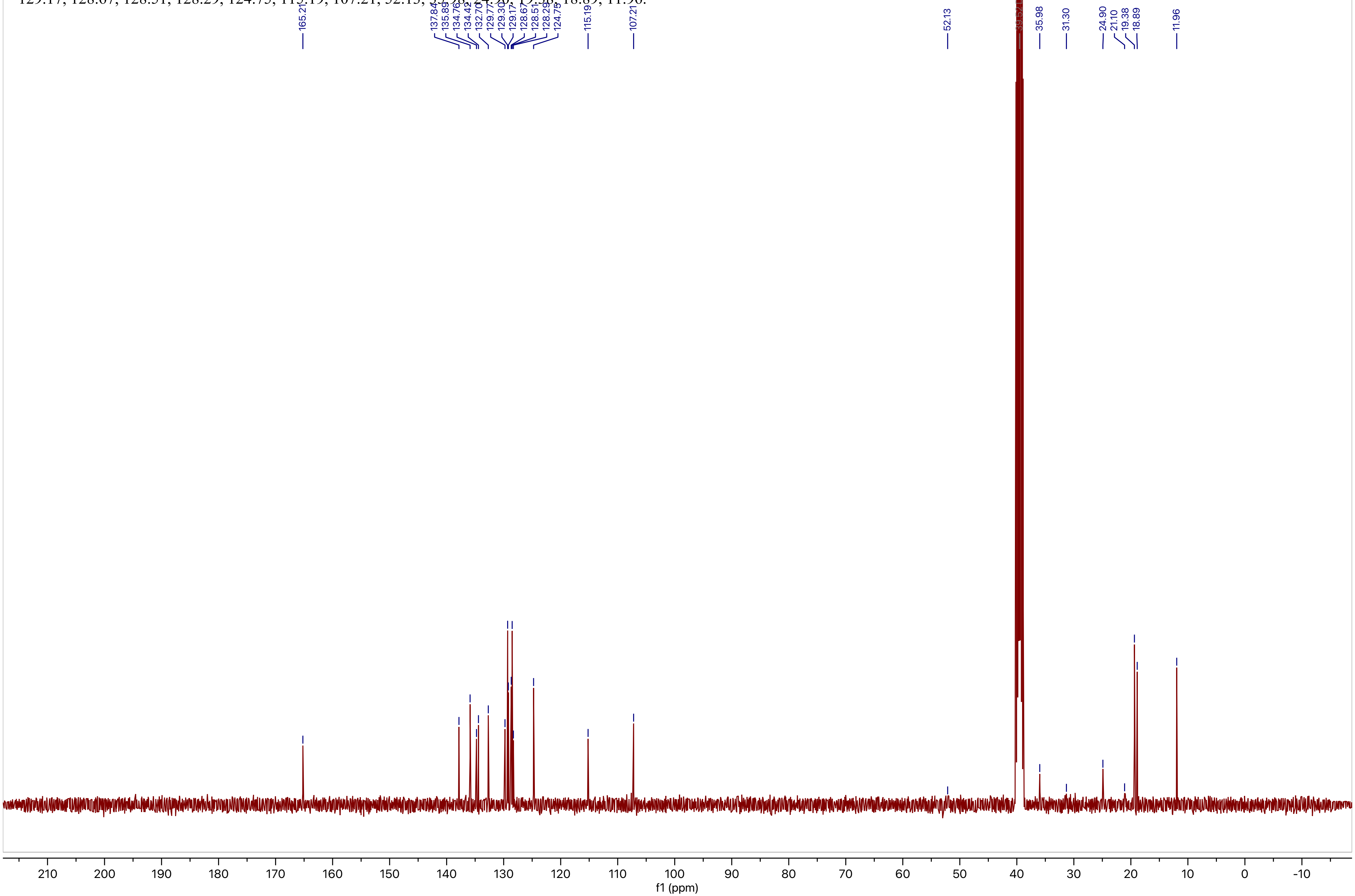

Compound 19

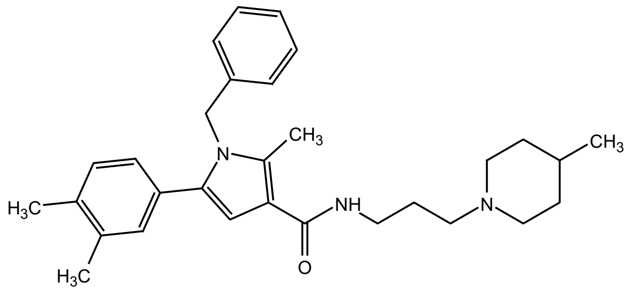


NMR (400 MHz, CDCl3 ) δ 7.34 – 7.28 (m, 2H), 7.25 – 7.21 (m, 1H), 7.09 – 7.01 (m, 2H), 7.01 – 6.89 (m, 4H), 6.51 (s, 1H) 5.11 (s, 2H), 3.49 (dd, $J$ = 11.6, 6.4 Hz, 2H), 3.36 (d, $J$ = 11.8 Hz, 2H), 3.04 (t, $J$ = 5.5 Hz, 2H), 2.87 (t, $J$ = 12.3 Hz, 2H), 2.42 (s, 3H), 2.22 (s, 3H), 2.17 (s, 3H), 2.11 – 2.02 (m, 2H), 1.92 (d, $J$ = 13.6 Hz, 2H), 1.84 – 1.61 (m, 1H), 1.61 – 1.39 (m, 2H), 0.98 (d, $J$ = 6.3 Hz, 3H).

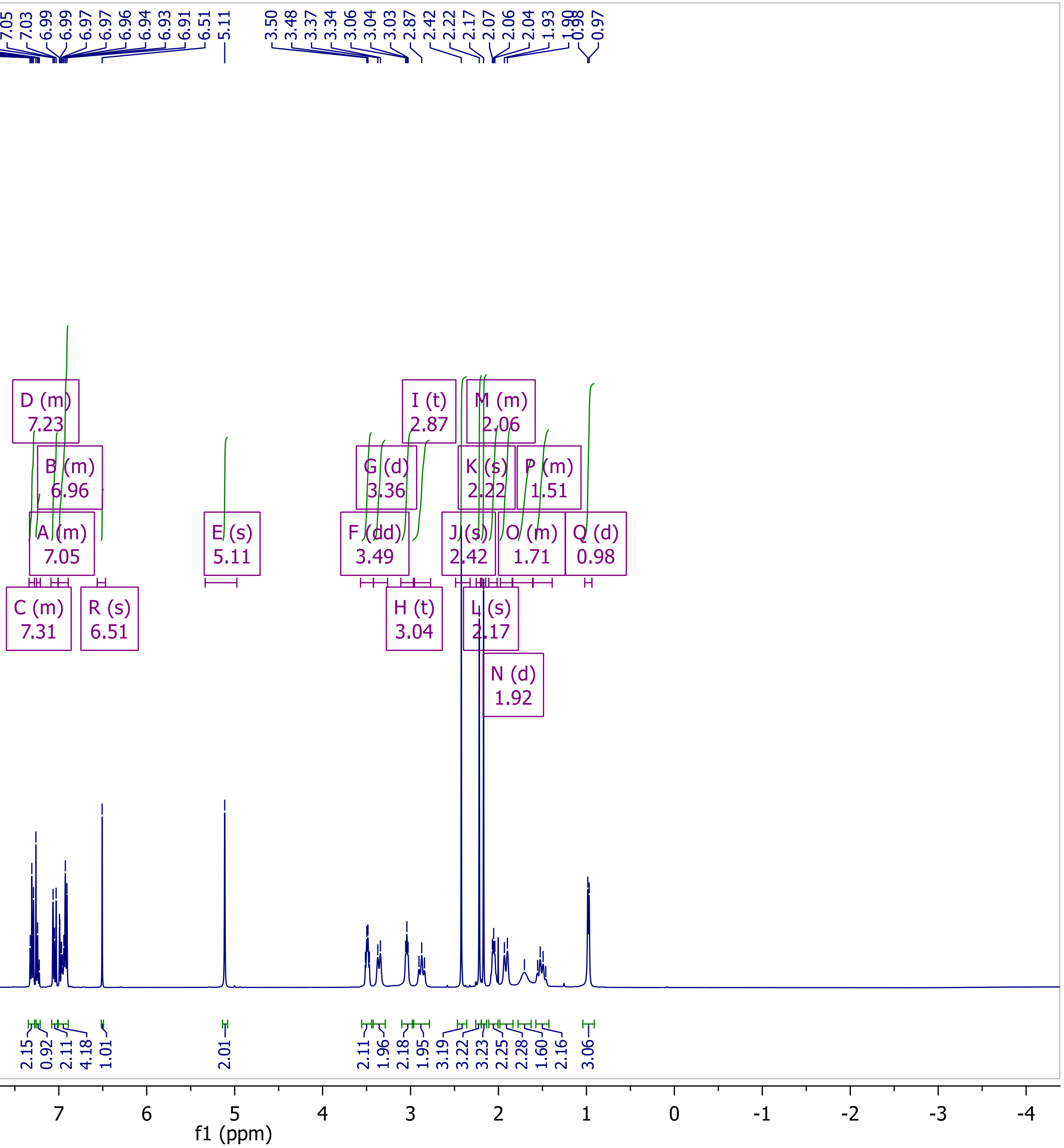

Compound 19

EPB1133-CDCl3.11.fid

$^{13}$C NMR (101 MHz, $CDCl_3$) δ 169.29, 137.73, 136.97, 136.50, 135.78, 135.45, 130.67, 129.89, 129.66, 129.04, 127.49, 126.45, 125.73, 113.07, 106.70, 53.65, 52.66, 47.93, 35.69, 31.85, 29.04, 24.70, 21.13, 19.87, 19.60, 11.84.

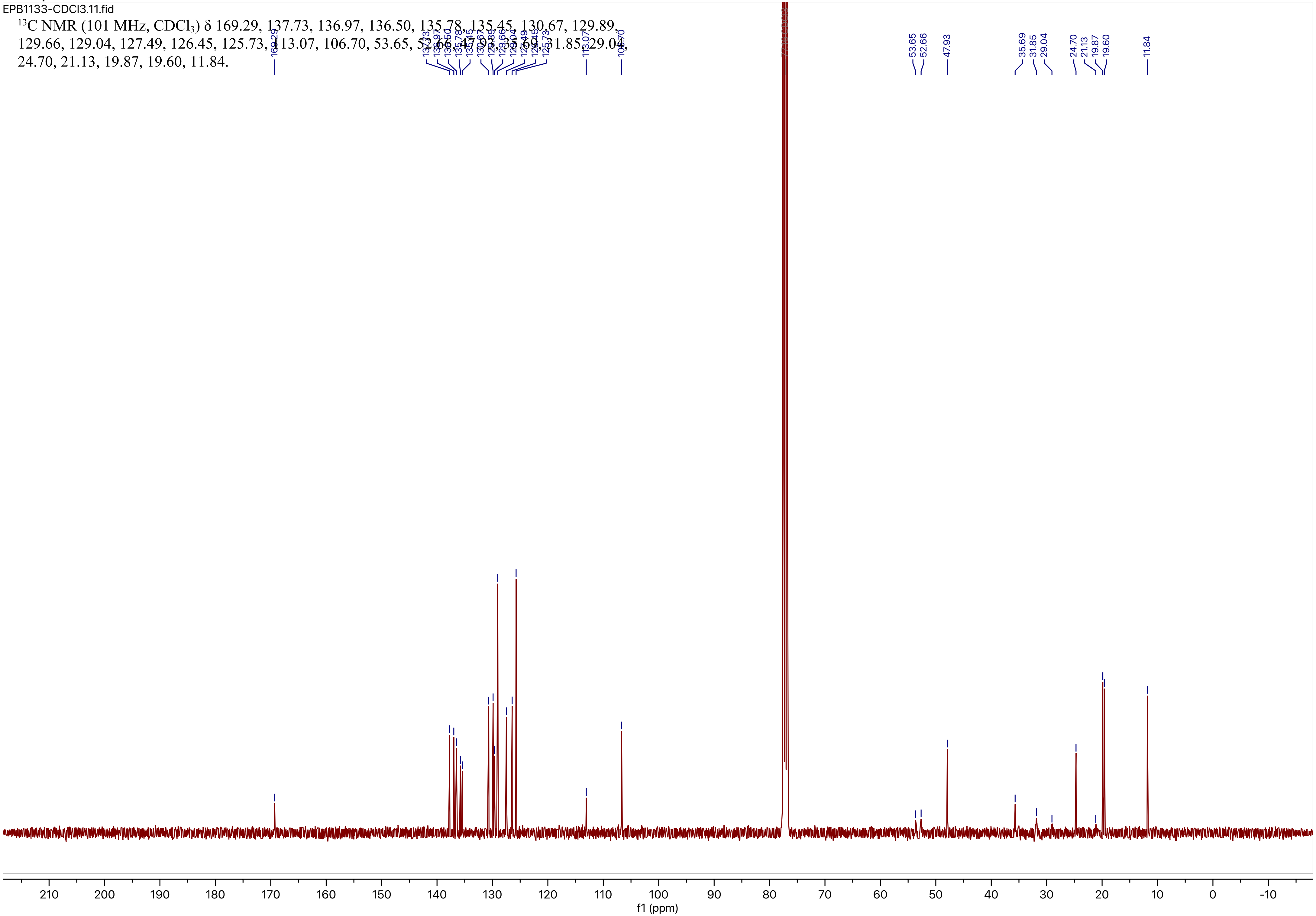

Compound 20

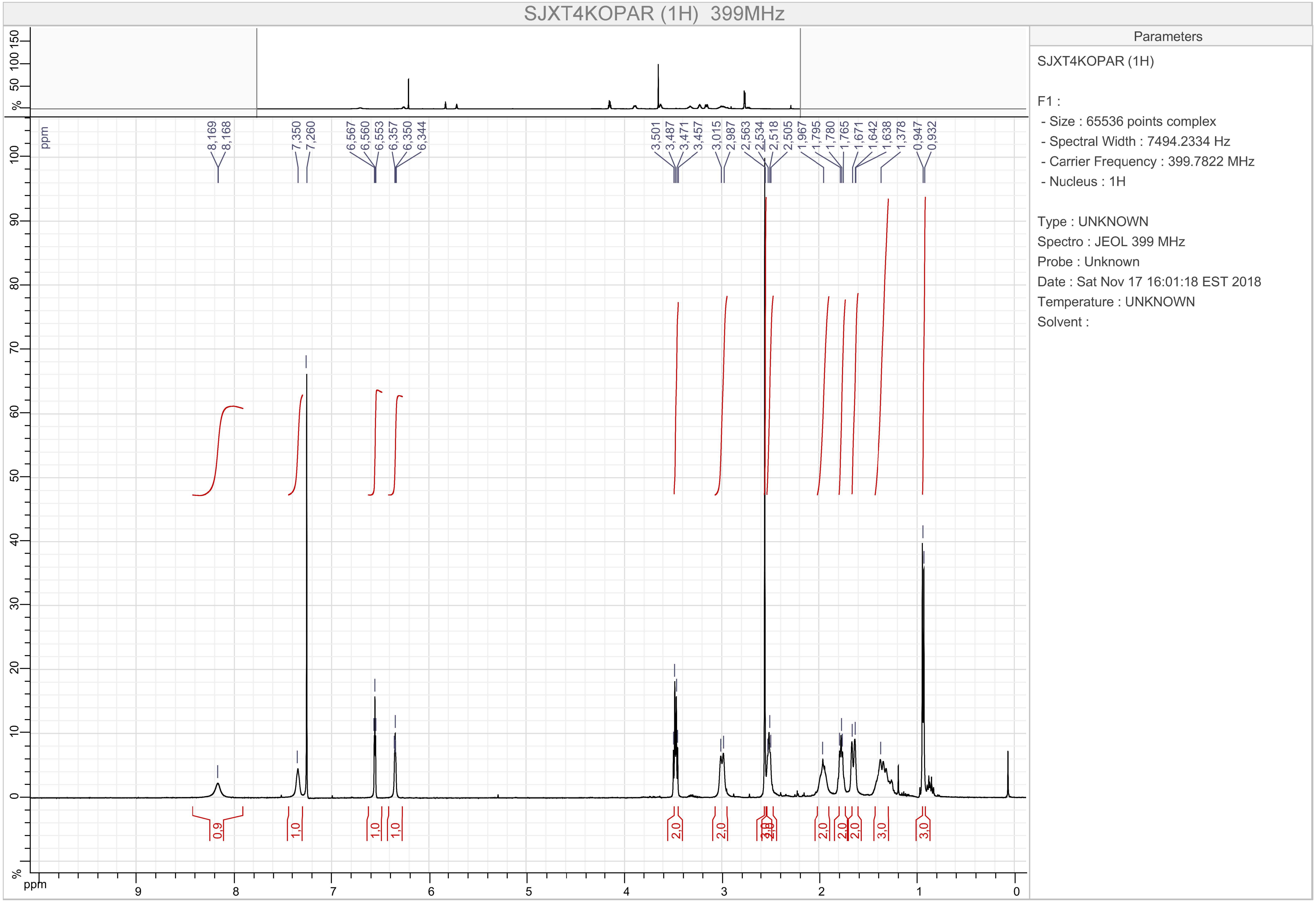
SJXT4KOPAR (1H) 399MHz
Parameters
SJXT4KOPAR (1H)
F1 :
- Size : 65536 points complex
- Spectral Width : 7494.2334 Hz
- Carrier Frequency : 399.7822 MHz
- Nucleus : 1H
Type : UNKNOWN
Spectro : JEOL 399 MHz
Probe : Unknown
Date : Sat Nov 17 16:01:18 EST 2018
Temperature : UNKNOWN
Solvent :
8,169
8,168
7,350
7,260
6,567
6,560
6,553
6,357
6,350
6,344
3,501
3,487
3,471
3,457
3,015
2,987
2,563
2,534
2,518
2,505
1,967
1,795
1,780
1,765
1,671
1,642
1,638
1,378
0,947
0,932
0,9
1,0
1,0
1,0
2,0
2,0
2,0
2,0
2,0
3,0
3,0
ppm
%

### Compound 20

EPB1018.11.fid

$^{13}$C NMR (101 MHz, DMSO) δ 165.13, 131.55, 115.12, 114.18, 106.94, 56.31, 53.44, 37.17, 33.89, 30.32, 26.71, 21.83, 12.55.

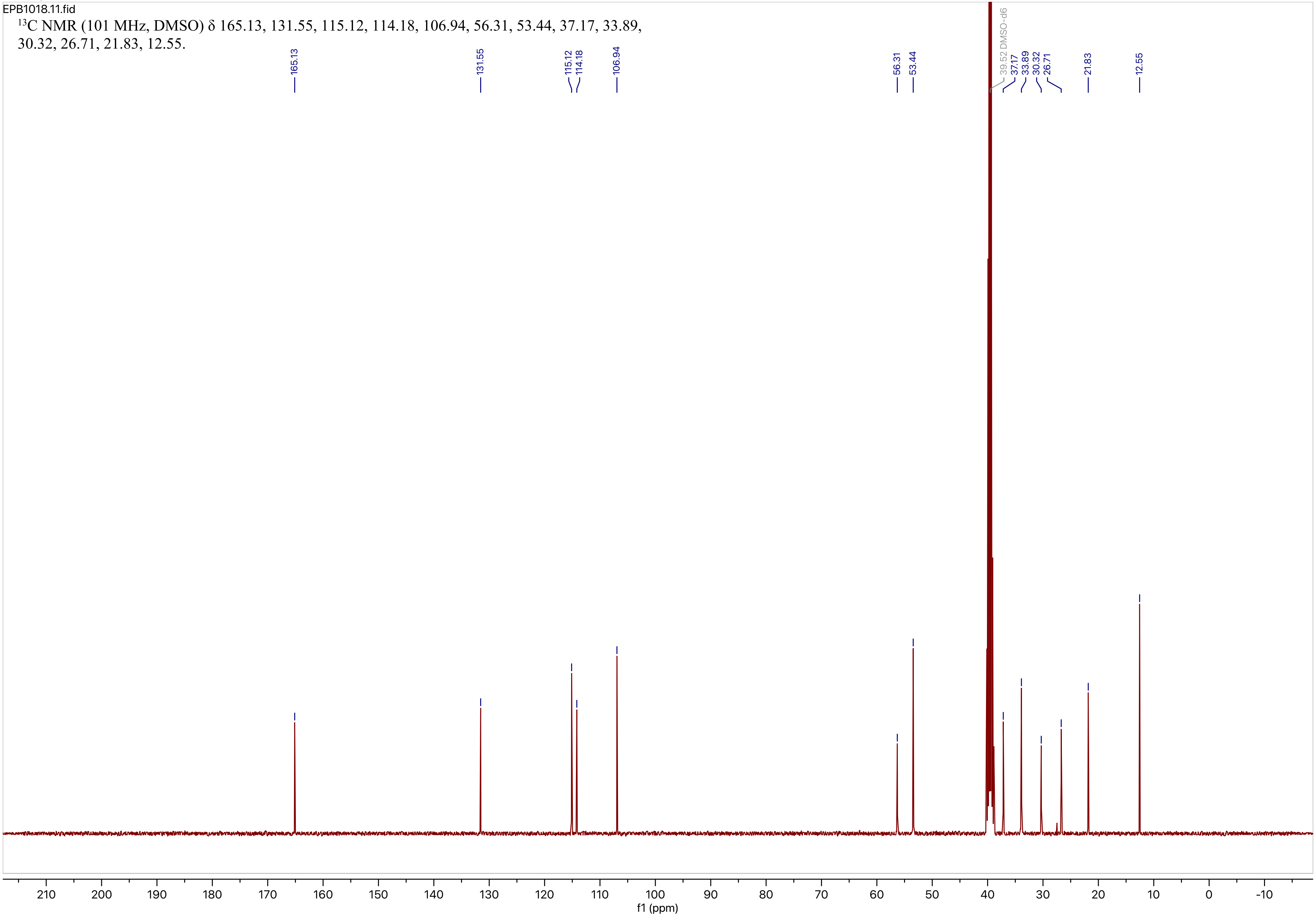

Compound 21

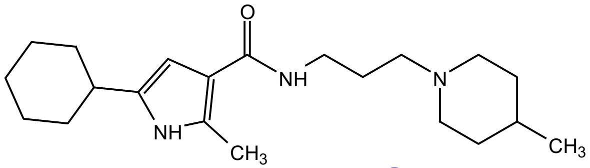

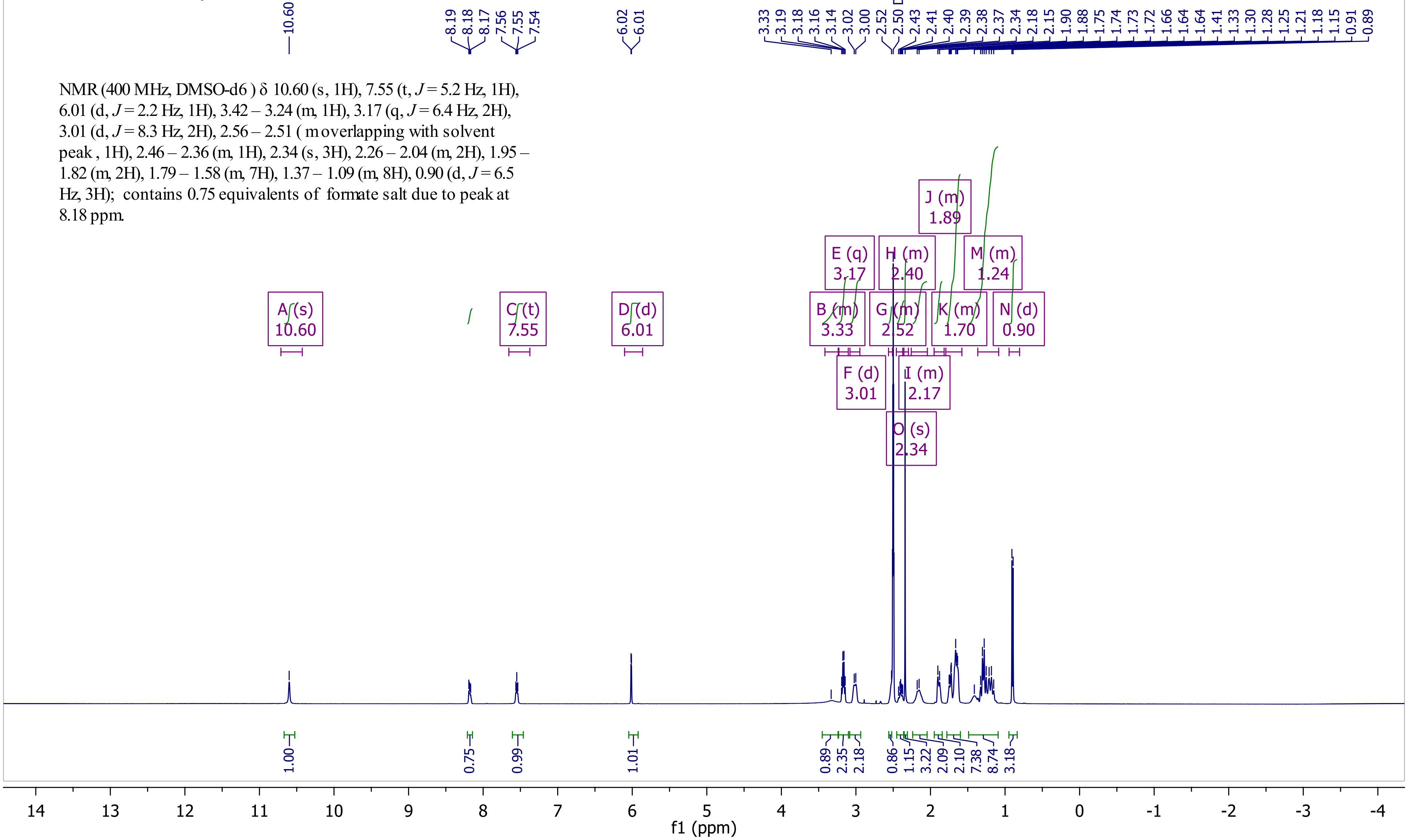


NMR (400 MHz, DMSO-d6 ) δ 10.60 (s, 1H), 7.55 (t, *J* = 5.2 Hz, 1H), 6.01 (d, *J* = 2.2 Hz, 1H), 3.42 – 3.24 (m, 1H), 3.17 (q, *J* = 6.4 Hz, 2H), 3.01 (d, *J* = 8.3 Hz, 2H), 2.56 – 2.51 ( m overlapping with solvent peak , 1H), 2.46 – 2.36 (m, 1H), 2.34 (s, 3H), 2.26 – 2.04 (m, 2H), 1.95 – 1.82 (m, 2H), 1.79 – 1.58 (m, 7H), 1.37 – 1.09 (m, 8H), 0.90 (d, *J* = 6.5 Hz, 3H); contains 0.75 equivalents of formate salt due to peak at 8.18 ppm.

Compound 21

EPB1141-CDCl3.11.fid

$^{13}$C NMR (101 MHz, $CDCl_3$) δ 168.91, 167.69, 136.62, 132.10, 112.99, 102.04, 54.60, 52.64, 36.55, 36.48, 33.10, 31.50, 29.36, 26.29, 26.11, 24.58, 21.08, 13.12.

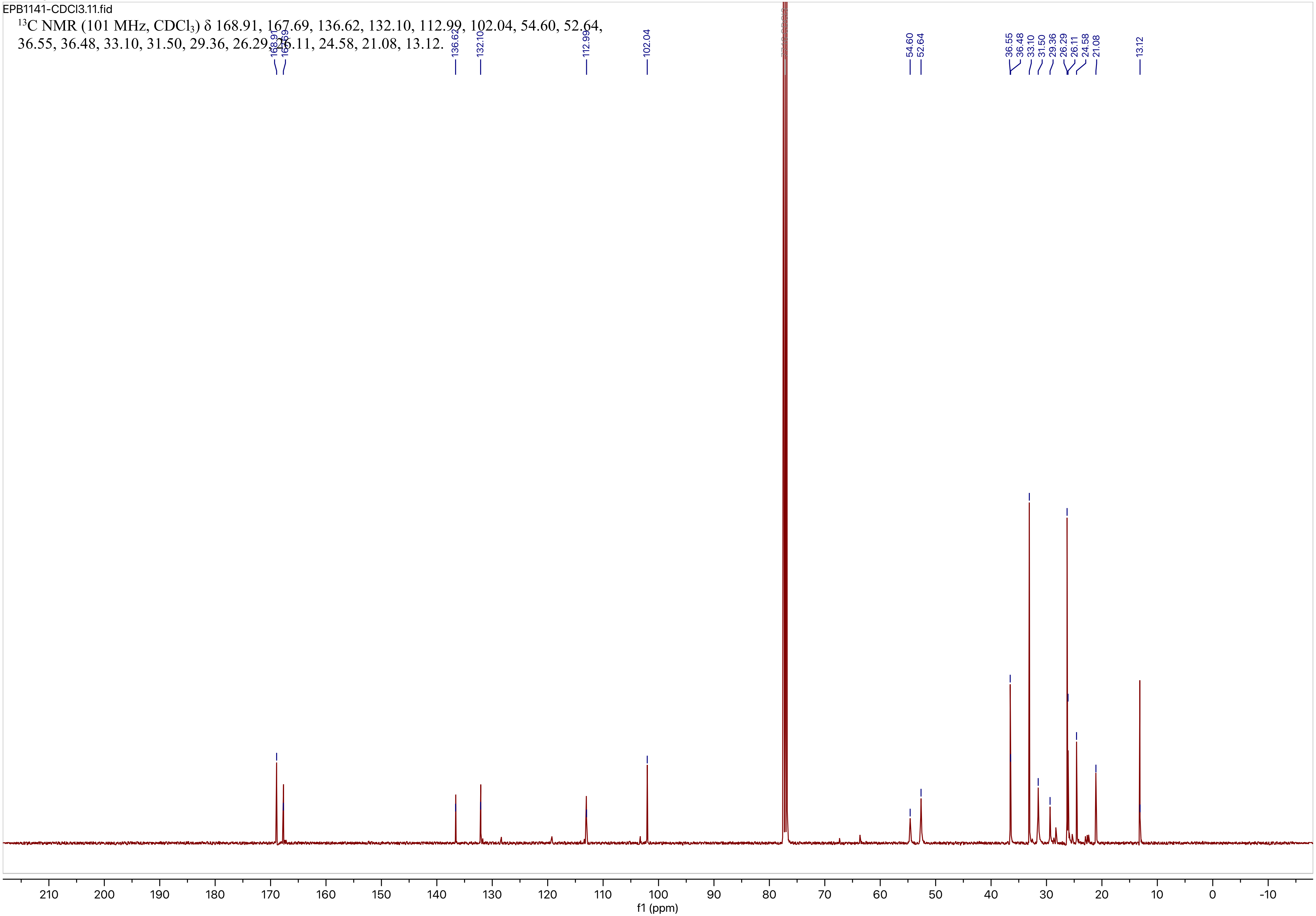

Compound 22

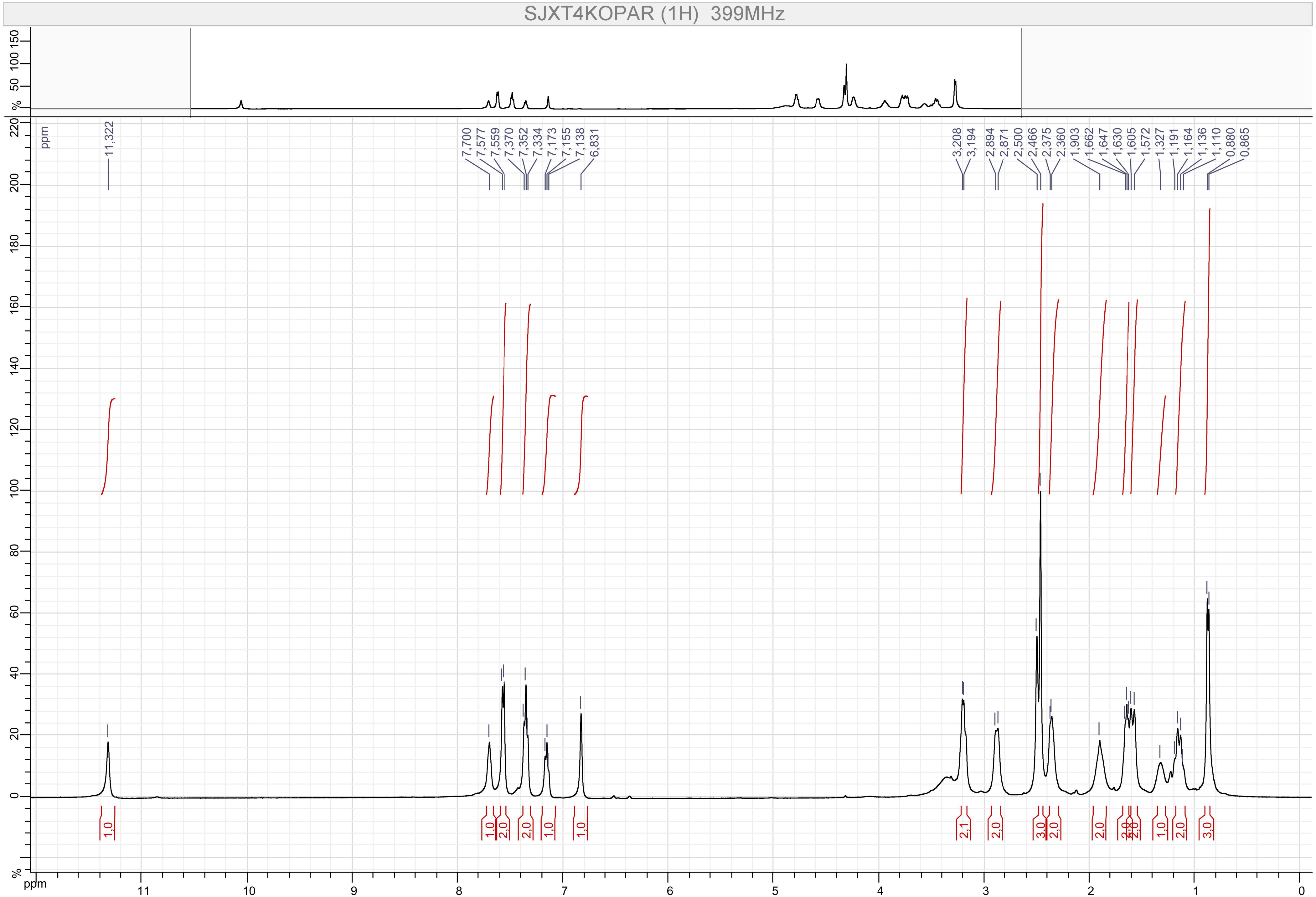

SJXT4KOPAR (1H) 399MHz
11,322
7,700
7,577
7,559
7,370
7,352
7,334
7,173
7,155
7,138
6,831
3,208
3,194
2,894
2,871
2,500
2,466
2,375
2,360
1,903
1,662
1,647
1,630
1,605
1,572
1,327
1,191
1,164
1,136
1,110
0,880
0,865
1,0
1,0
2,0
2,0
1,0
1,0
2,1
2,0
3,0
2,0
2,0
1,0
2,0
3,0
ppm
%

Compound 22

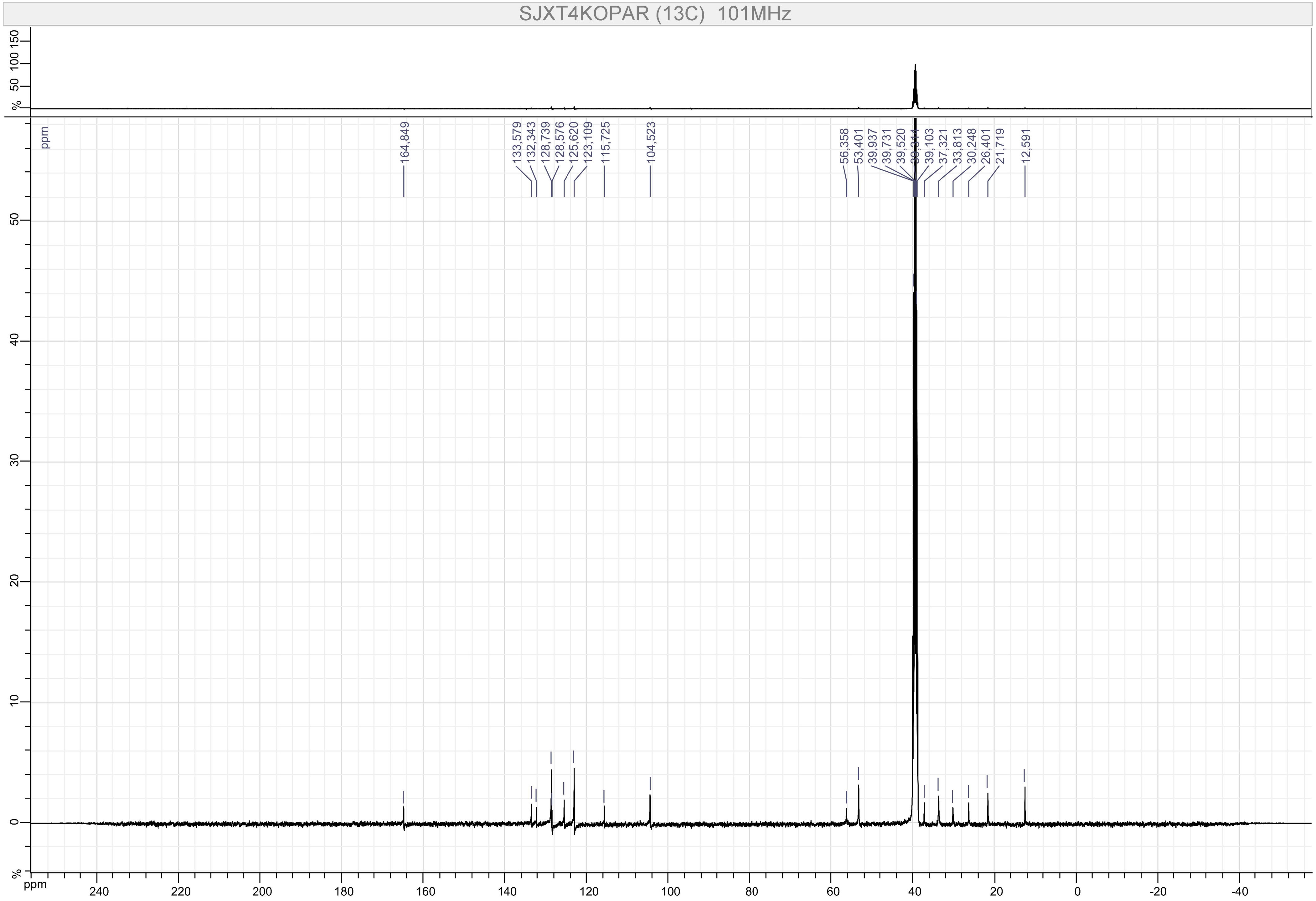

SJXT4KOPAR (13C) 101MHz
164,849
133,579
132,343
128,739
128,576
125,620
123,109
115,725
104,523
56,358
53,401
39,937
39,731
39,520
39,314
39,103
37,321
33,813
30,248
26,401
21,719
12,591
ppm
%

Compound 23

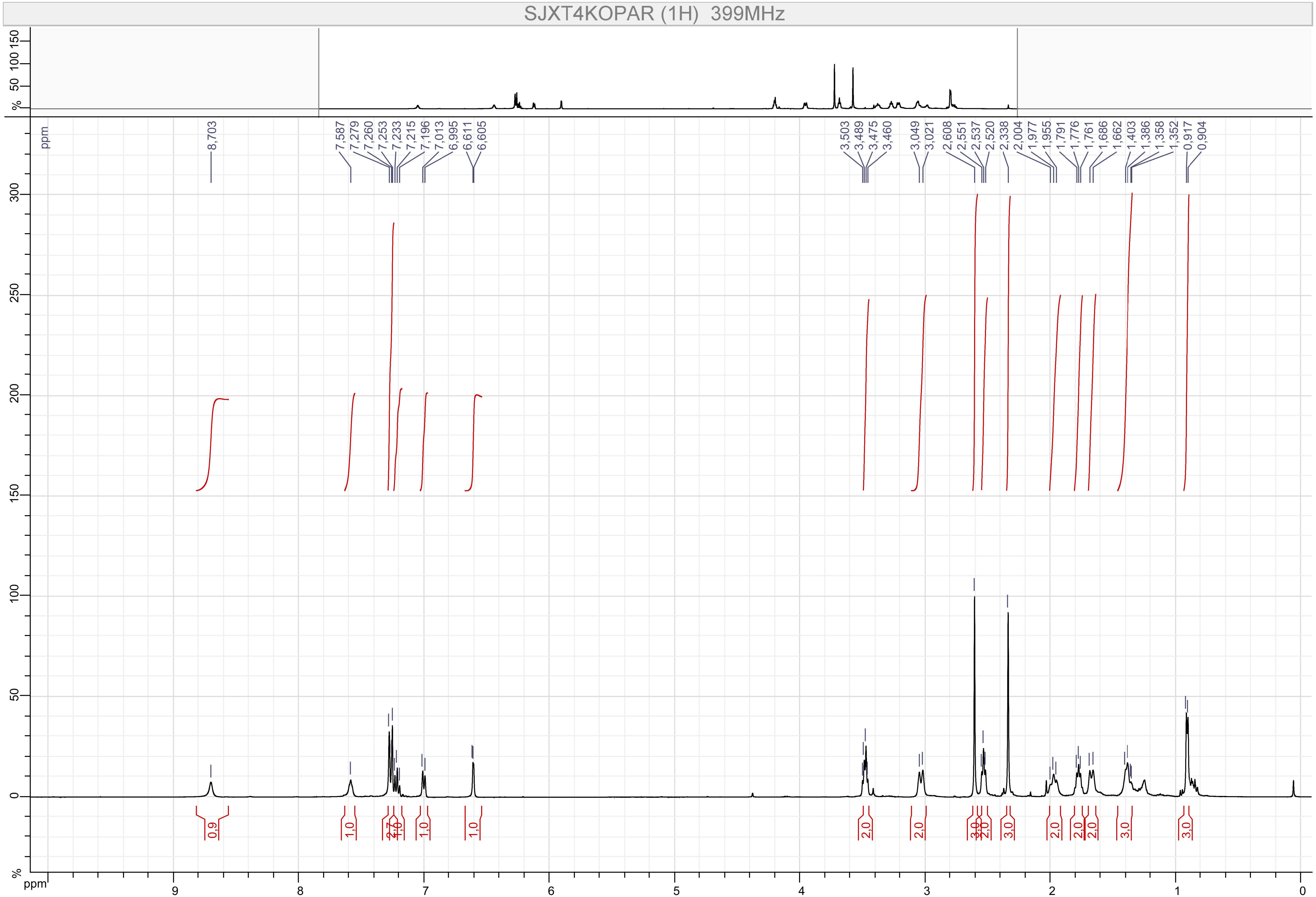

SJXT4KOPAR (1H) 399MHz
8,703
7,587 7,279 7,260 7,253 7,233 7,215 7,196 7,013 6,995 6,611 6,605
3,503 3,489 3,475 3,460 3,049 3,021 2,608 2,551 2,537 2,520 2,338 2,004 1,977 1,955 1,791 1,776 1,761 1,686 1,662 1,403 1,386 1,358 1,352 0,917 0,904
0,9 1,0 2,7 1,0 1,0 1,0 2,0 2,0 3,0 2,0 3,0 2,0 2,0 2,0 3,0 3,0
ppm

Compound 23

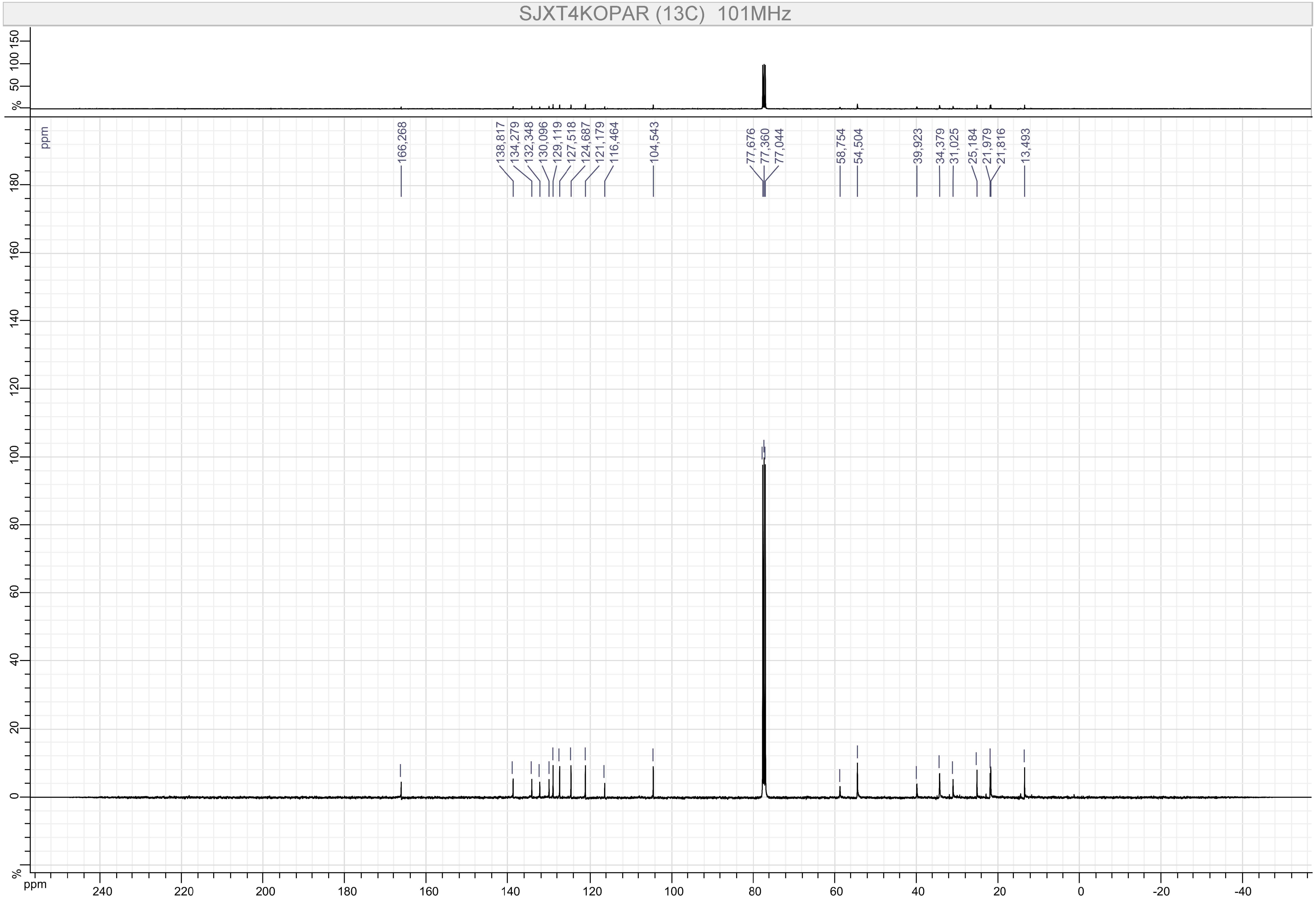
SJXT4KOPAR (13C) 101MHz
166,268
138,817
134,279
132,348
130,096
129,119
127,518
124,687
121,179
116,464
104,543
77,676
77,360
77,044
58,754
54,504
39,923
34,379
31,025
25,184
21,979
21,816
13,493
ppm
%

Compound 24

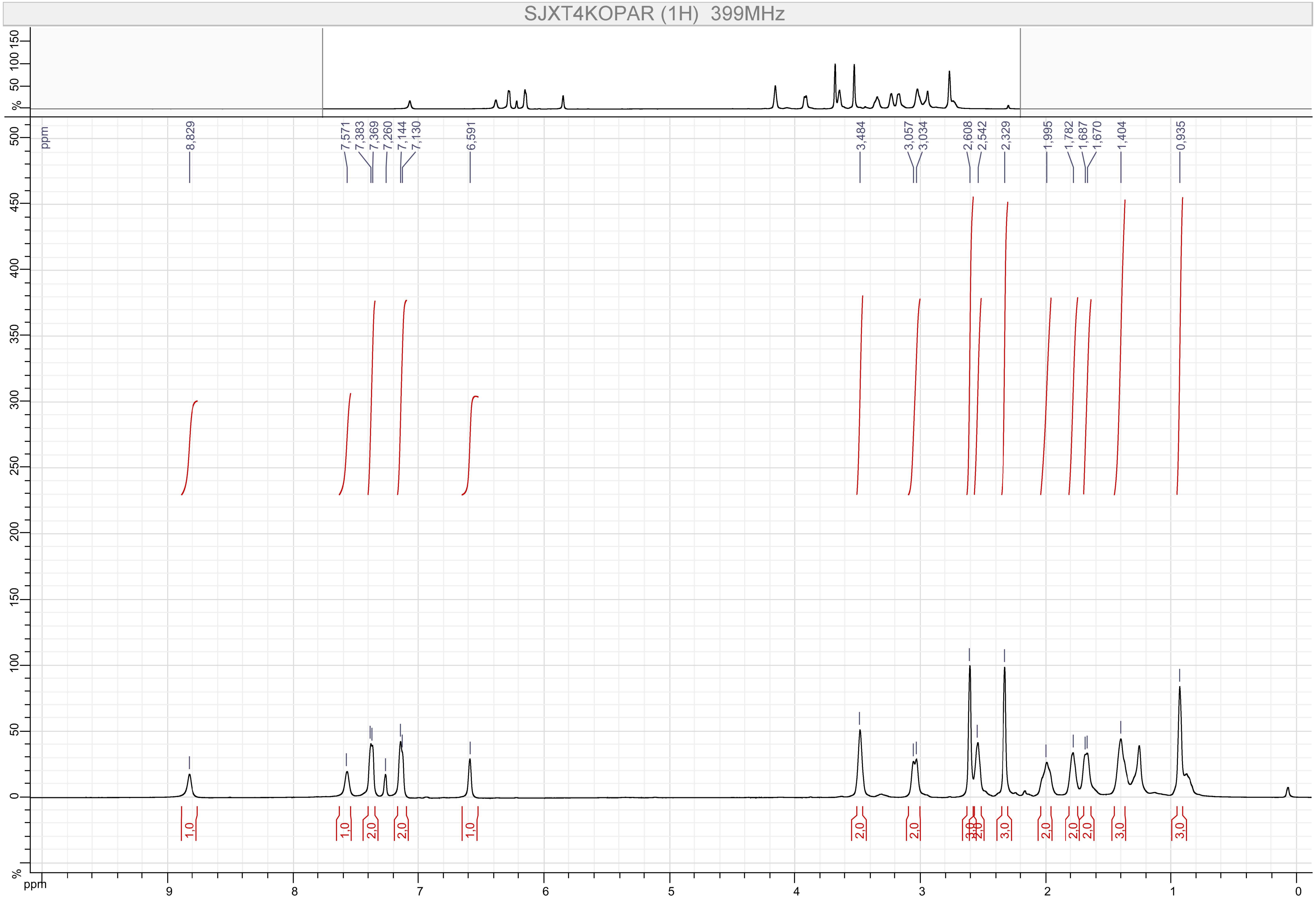
SJXT4KOPAR (1H) 399MHz
8,829
7,571
7,383
7,369
7,260
7,144
7,130
6,591
3,484
3,057
3,034
2,608
2,542
2,329
1,995
1,782
1,687
1,670
1,404
0,935
1,0
1,0
2,0
2,0
1,0
2,0
2,0
3,0
2,0
3,0
2,0
2,0
2,0
3,0
3,0
ppm
%

Compound 24

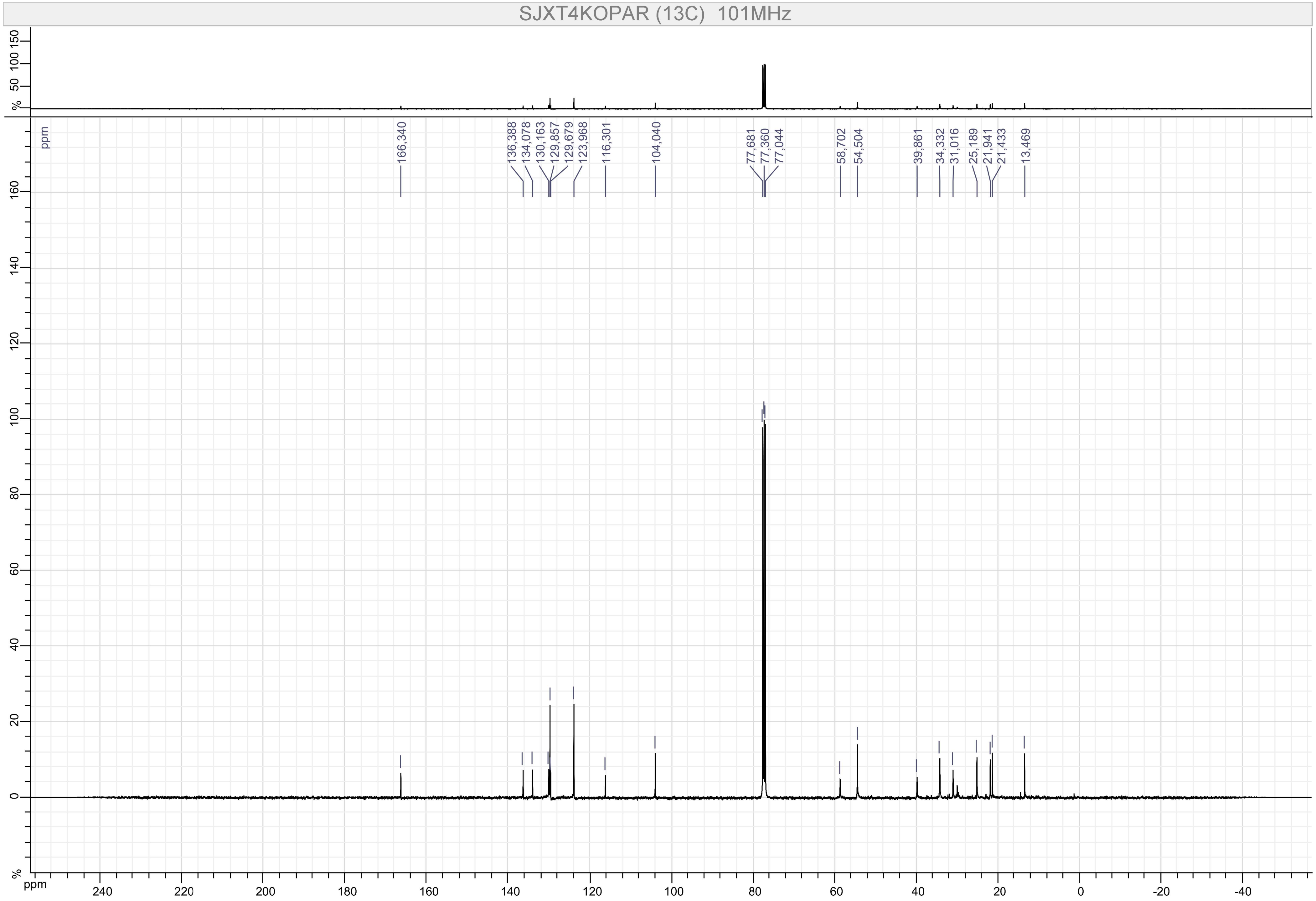
SJXT4KOPAR (13C) 101MHz
166,340
136,388
134,078
130,163
129,857
129,679
123,968
116,301
104,040
77,681
77,360
77,044
58,702
54,504
39,861
34,332
31,016
25,189
21,941
21,433
13,469
ppm
%

Compound 25

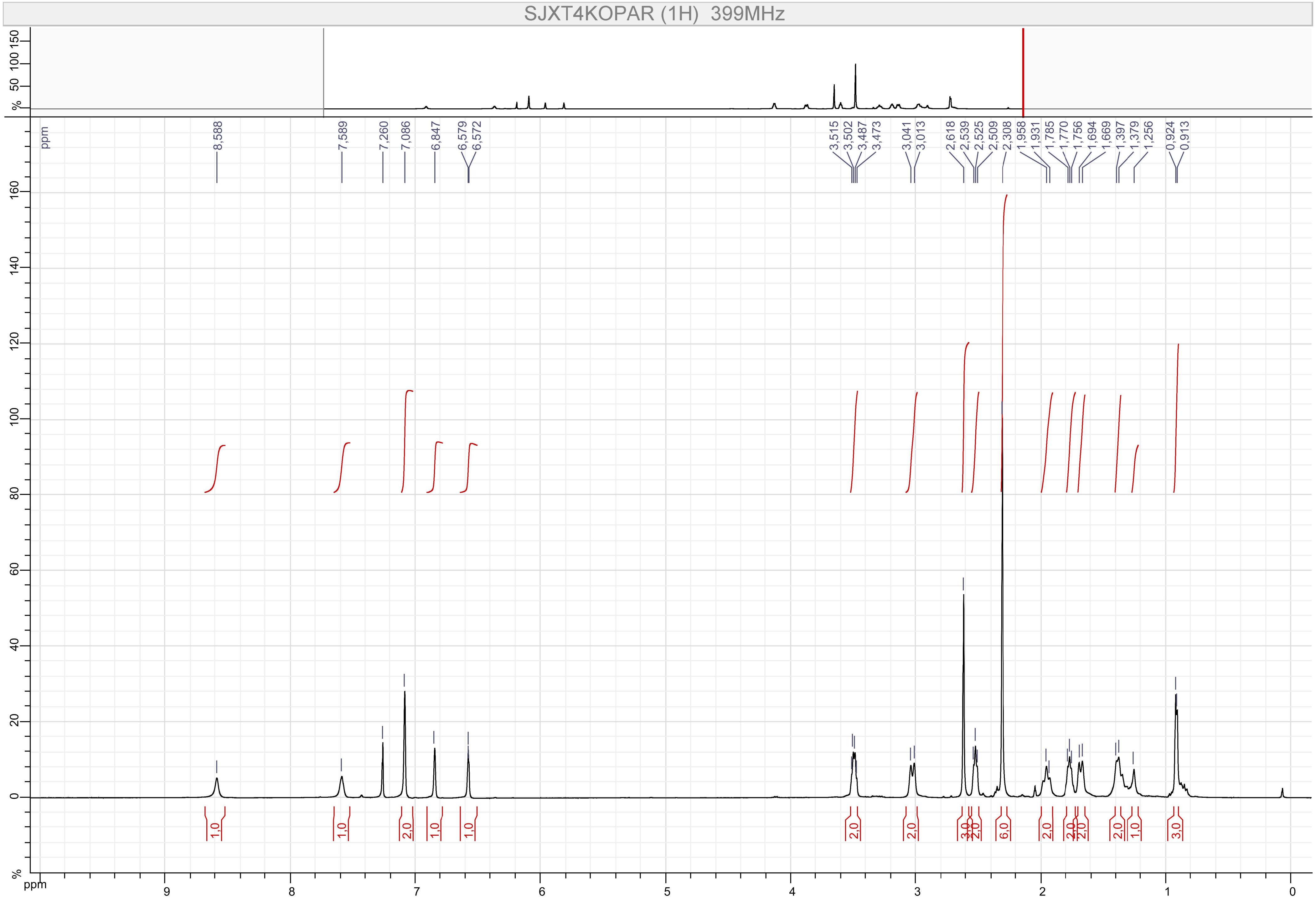
SJXT4KOPAR (1H) 399MHz
8,588
7,589
7,260
7,086
6,847
6,579
6,572
3,515
3,502
3,487
3,473
3,041
3,013
2,618
2,539
2,525
2,509
2,308
1,958
1,931
1,785
1,770
1,756
1,694
1,669
1,397
1,379
1,256
0,924
0,913
1,0
1,0
2,0
1,0
1,0
2,0
2,0
3,0
2,0
6,0
2,0
2,0
2,0
2,0
1,0
3,0
ppm
%

Compound 25

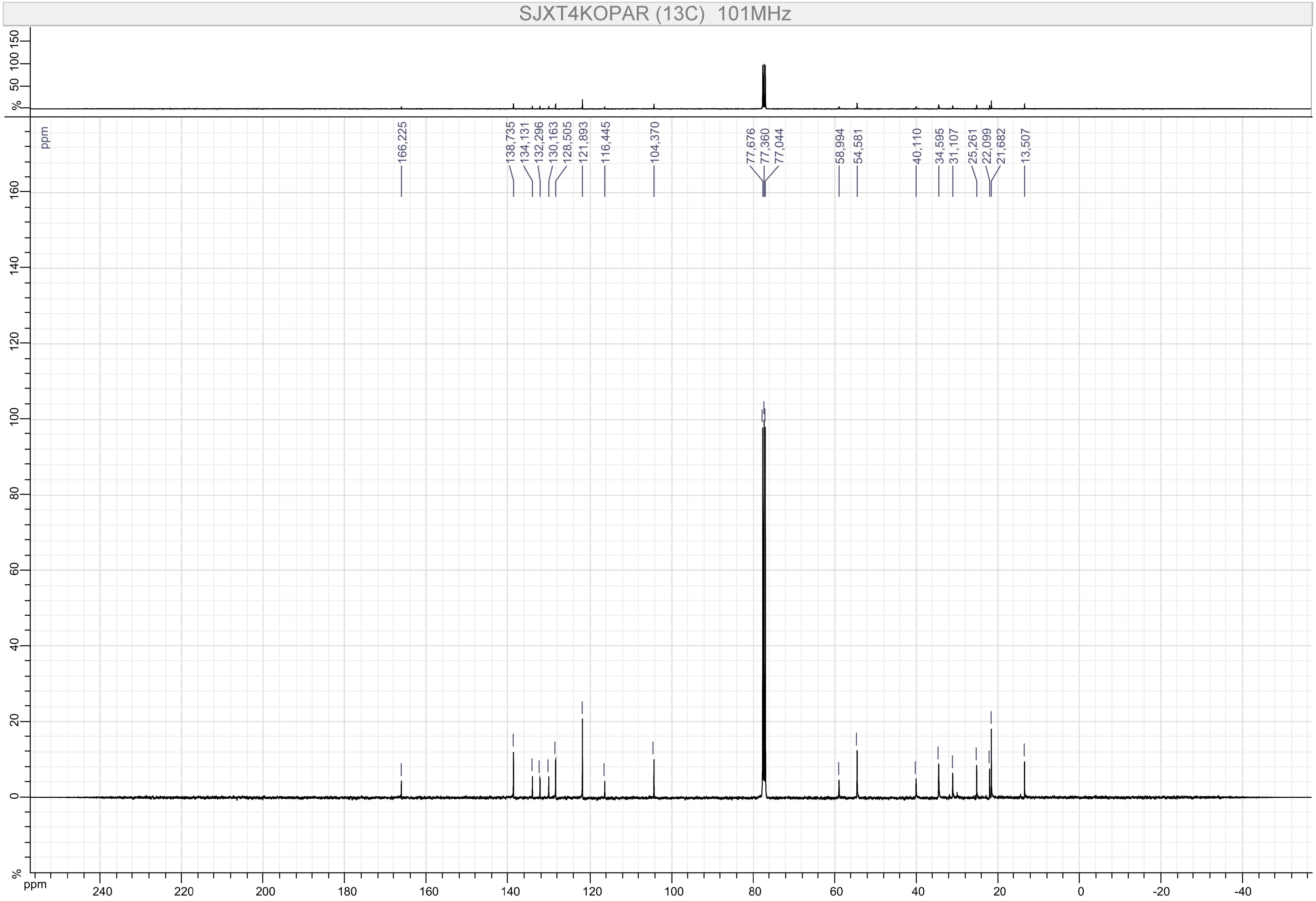
SJXT4KOPAR (13C) 101MHz
166,225
138,735
134,131
132,296
130,163
128,505
121,893
116,445
104,370
77,676
77,360
77,044
58,994
54,581
40,110
34,595
31,107
25,261
22,099
21,682
13,507
ppm
%

Compound 26

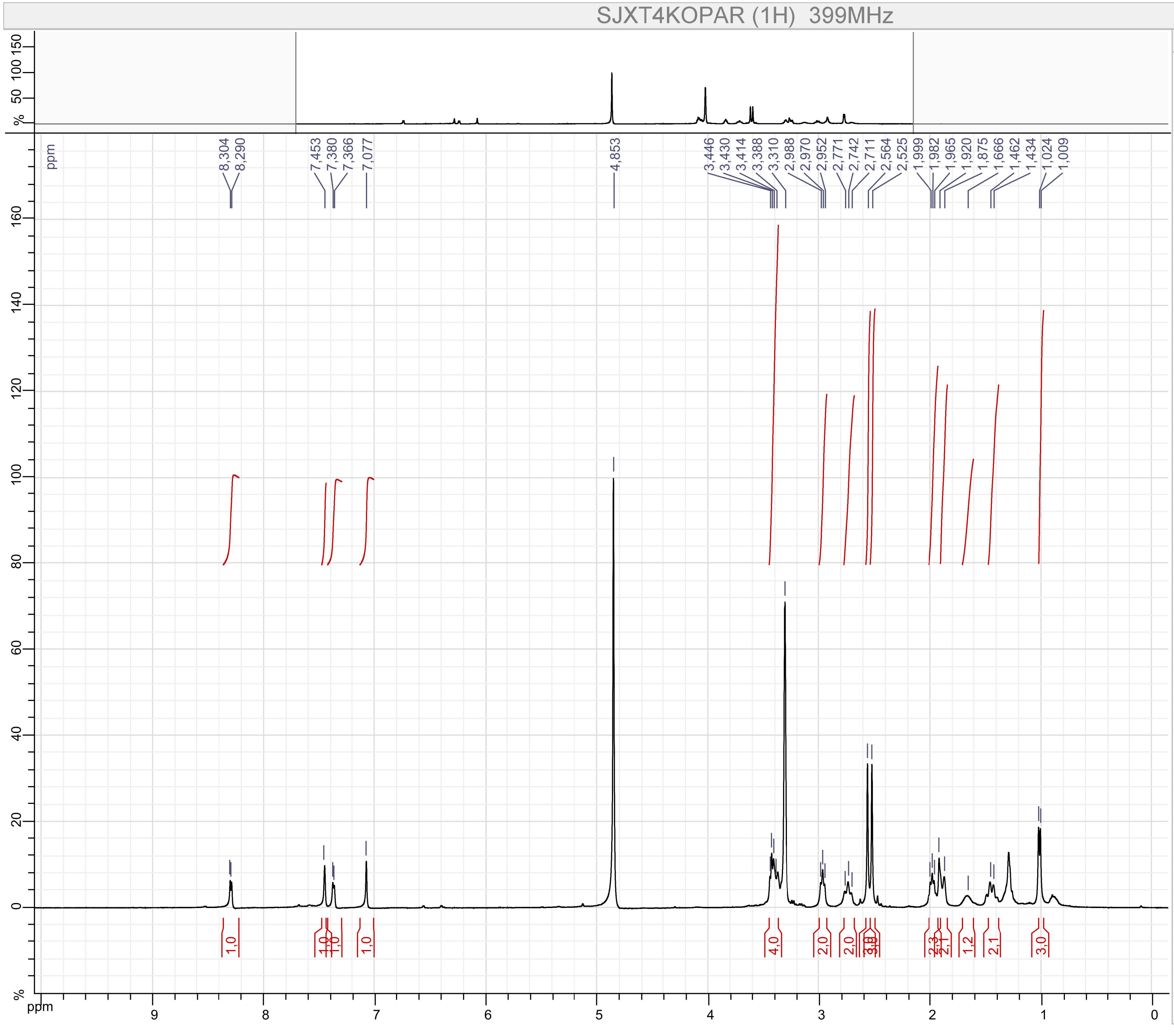


Parameters

SJXT4KOPAR (1H)

F1 :
- Size : 65536 points complex
- Spectral Width : 7494.2334 Hz
- Carrier Frequency : 399.7822 MHz
- Nucleus : 1H

Type : UNKNOWN
Spectro : JEOL 399 MHz
Probe : Unknown
Date : Fri Sep 06 21:36:54 EDT 2019
Temperature : UNKNOWN
Solvent :

Compound 26

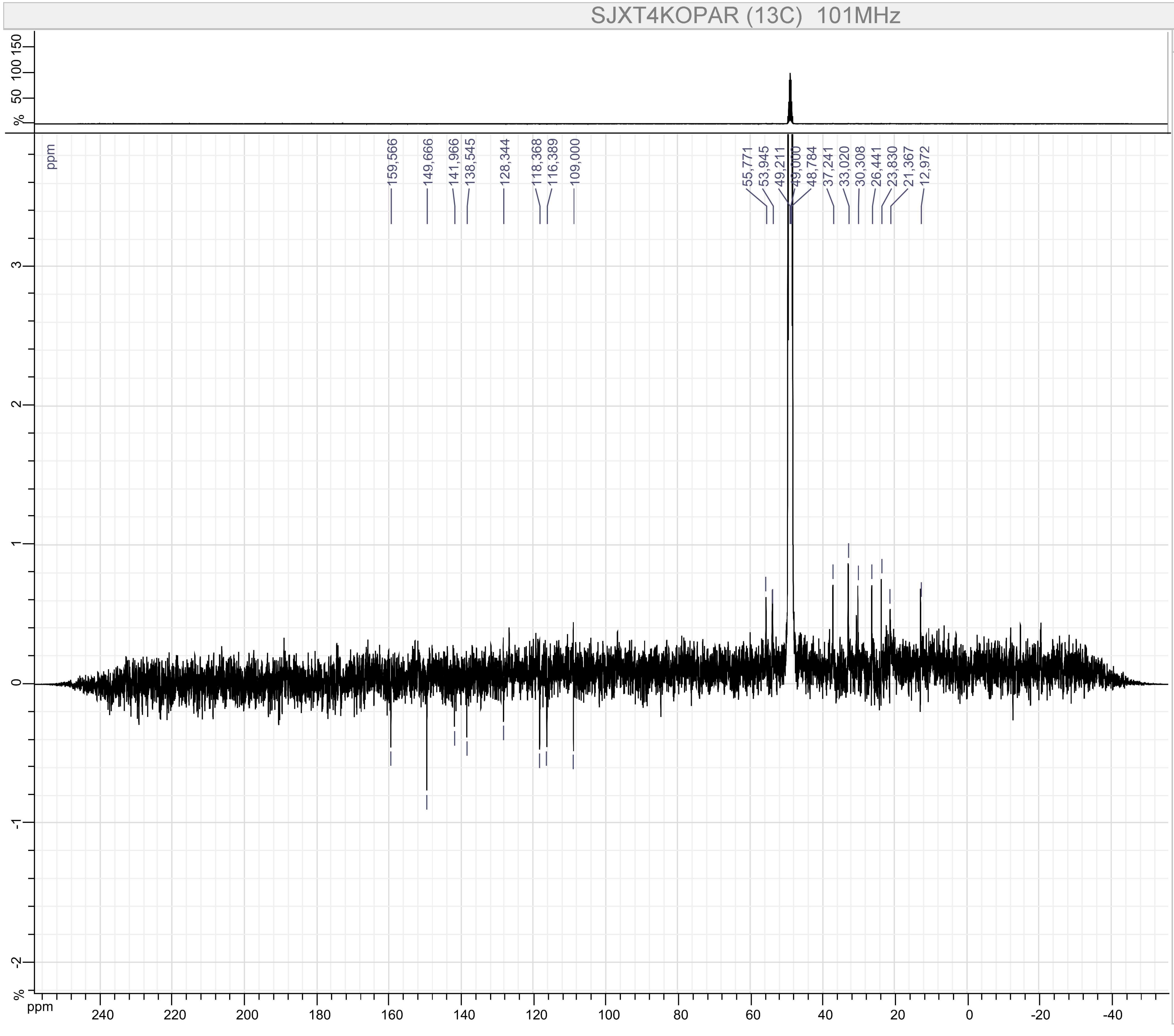


Parameters

SJXT4KOPAR (13C)

F1 :
- Size : 65536 points complex
- Spectral Width : 31566.62 Hz
- Carrier Frequency : 100.52531 MHz
- Nucleus : 13C

Type : UNKNOWN
Spectro : JEOL 101 MHz
Probe : Unknown
Date : Fri Sep 06 21:40:35 EDT 2019
Temperature : UNKNOWN
Solvent :

Compound 27

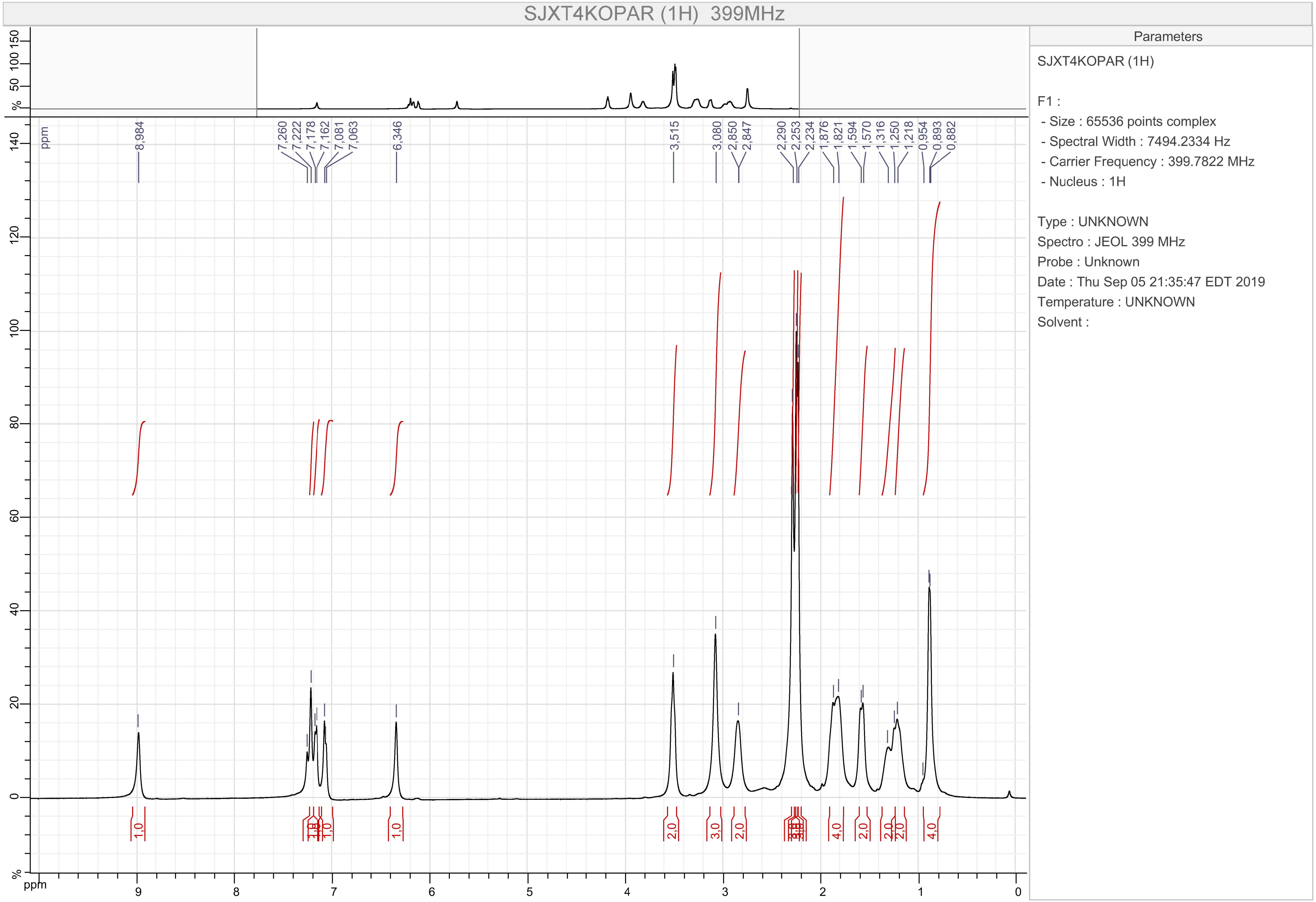

SJXT4KOPAR (1H) 399MHz
Parameters
SJXT4KOPAR (1H)
F1 :
- Size : 65536 points complex
- Spectral Width : 7494.2334 Hz
- Carrier Frequency : 399.7822 MHz
- Nucleus : 1H
Type : UNKNOWN
Spectro : JEOL 399 MHz
Probe : Unknown
Date : Thu Sep 05 21:35:47 EDT 2019
Temperature : UNKNOWN
Solvent :
8,984
7,260
7,222
7,178
7,162
7,081
7,063
6,346
3,515
3,080
2,850
2,847
2,290
2,253
2,234
1,876
1,821
1,594
1,570
1,316
1,250
1,218
0,954
0,893
0,882
ppm

Compound 27

SJXT4KOPAR (13C)  101MHz

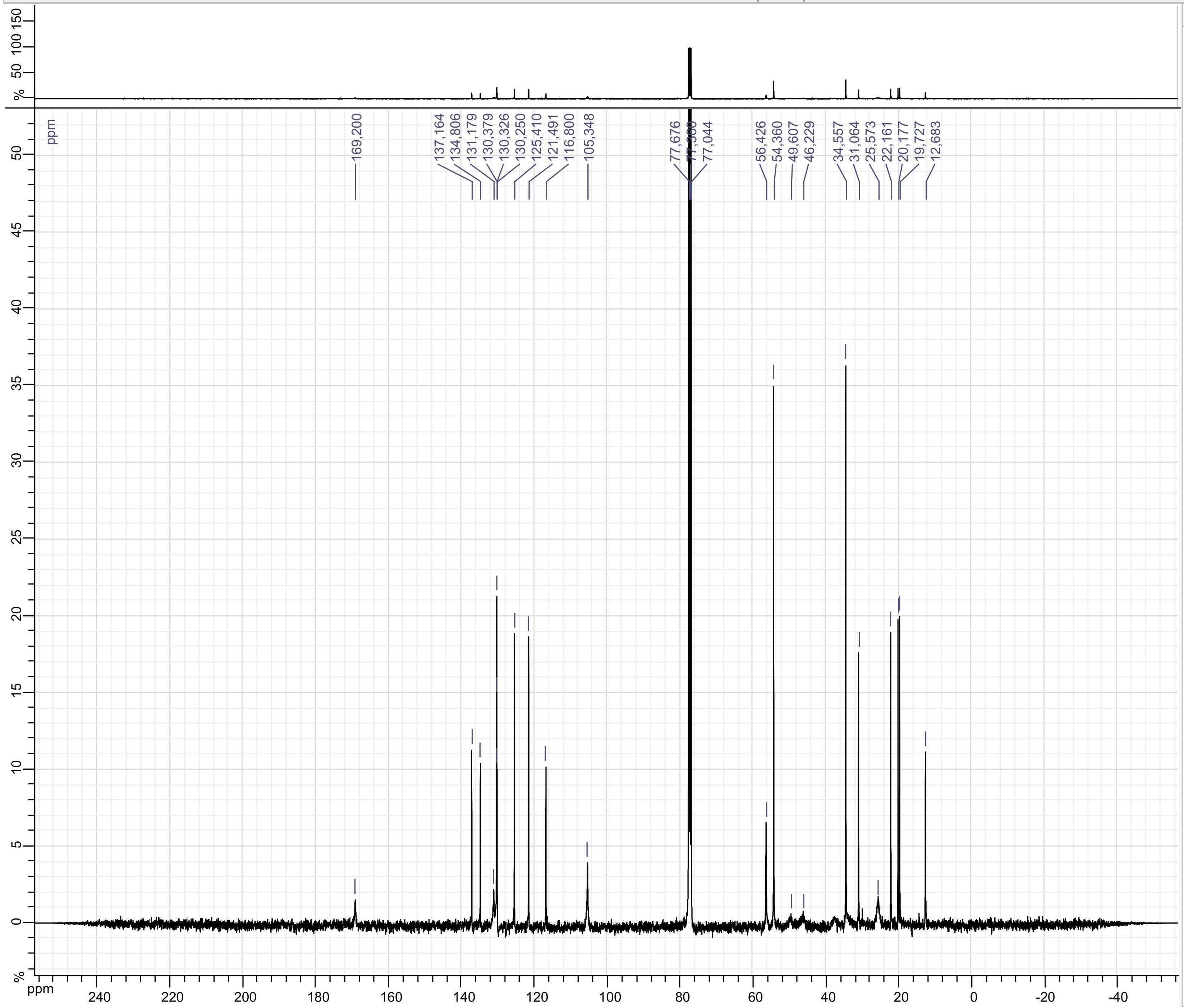


Parameters

SJXT4KOPAR (13C)

F1 :
- Size : 65536 points complex
- Spectral Width : 31566.62 Hz
- Carrier Frequency : 100.52531 MHz
- Nucleus : 13C

Type : UNKNOWN
Spectro : JEOL 101 MHz
Probe : Unknown
Date : Fri Sep 06 21:44:38 EDT 2019
Temperature : UNKNOWN
Solvent :

Compound 28

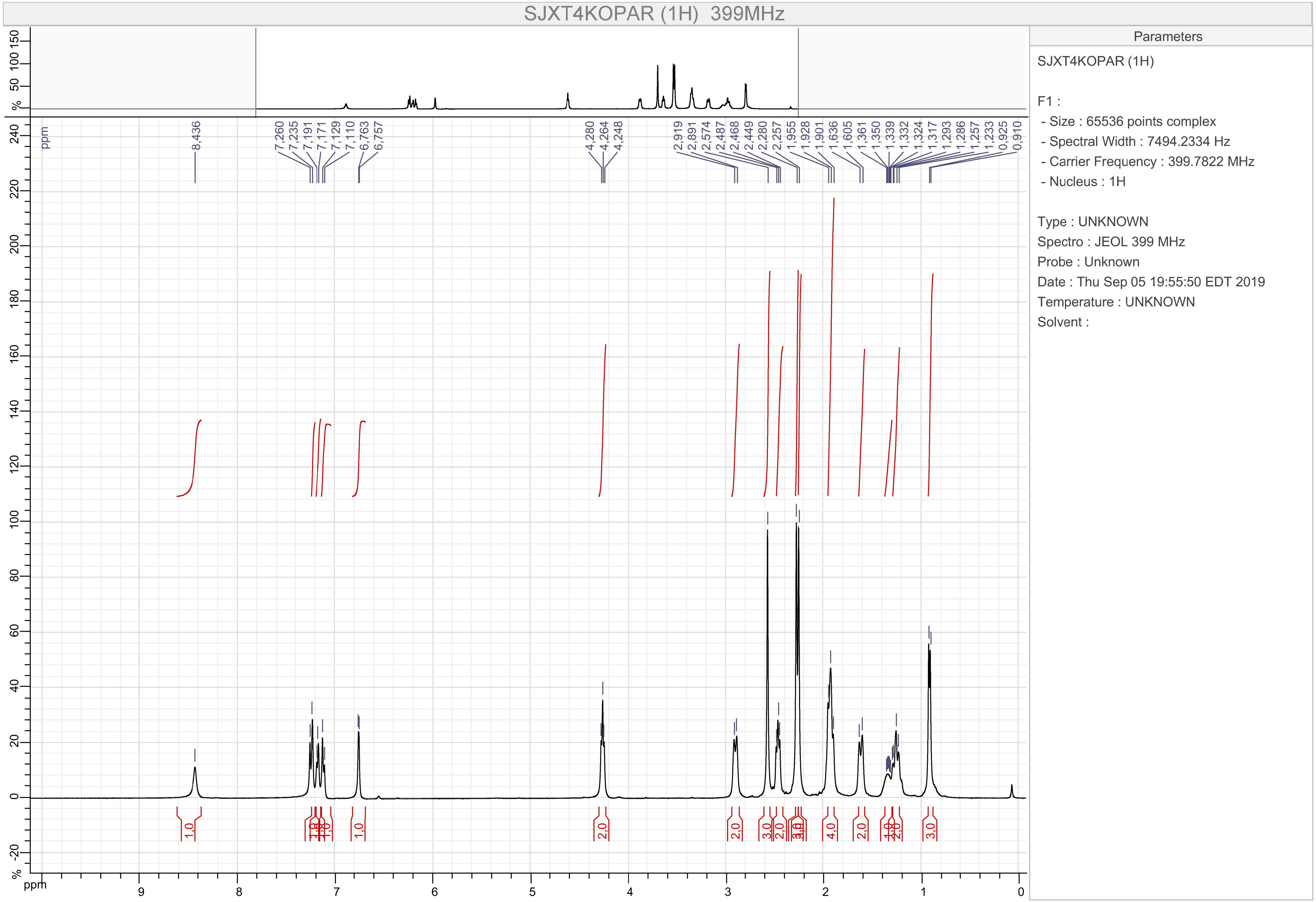

SJXT4KOPAR (1H) 399MHz
Parameters
SJXT4KOPAR (1H)
F1 :
- Size : 65536 points complex
- Spectral Width : 7494.2334 Hz
- Carrier Frequency : 399.7822 MHz
- Nucleus : 1H
Type : UNKNOWN
Spectro : JEOL 399 MHz
Probe : Unknown
Date : Thu Sep 05 19:55:50 EDT 2019
Temperature : UNKNOWN
Solvent :
8,436
7,260
7,235
7,191
7,171
7,129
7,110
6,763
6,757
4,280
4,264
4,248
2,919
2,891
2,574
2,487
2,468
2,449
2,280
2,257
1,955
1,928
1,901
1,636
1,605
1,361
1,350
1,339
1,332
1,324
1,317
1,293
1,286
1,257
1,233
0,925
0,910
ppm

Compound 28

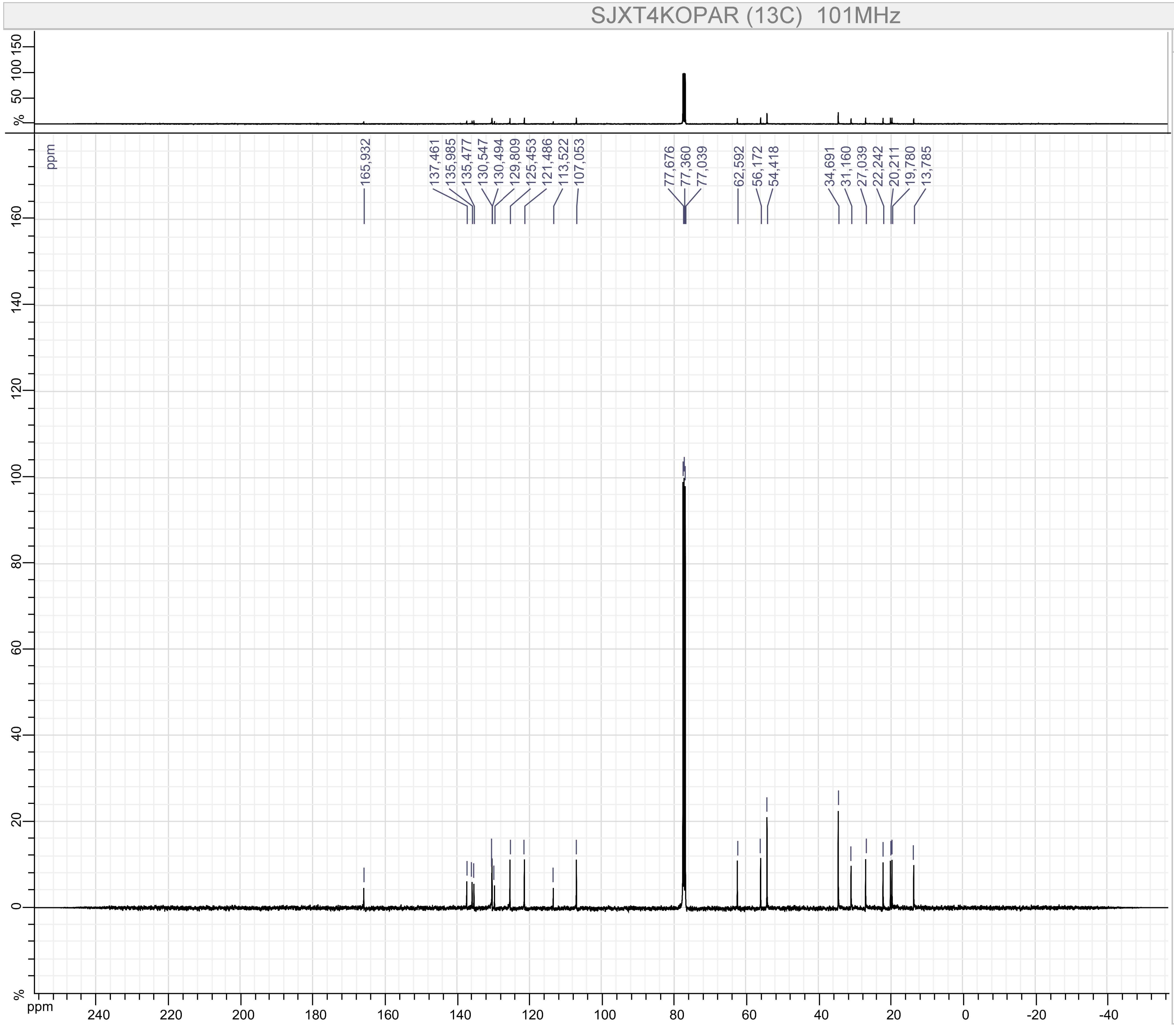


Parameters

SJXT4KOPAR (13C)

F1 :
- Size : 65536 points complex
- Spectral Width : 31566.62 Hz
- Carrier Frequency : 100.52531 MHz
- Nucleus : 13C

Type : UNKNOWN
Spectro : JEOL 101 MHz
Probe : Unknown
Date : Fri Sep 06 21:43:10 EDT 2019
Temperature : UNKNOWN
Solvent :

Compound 29

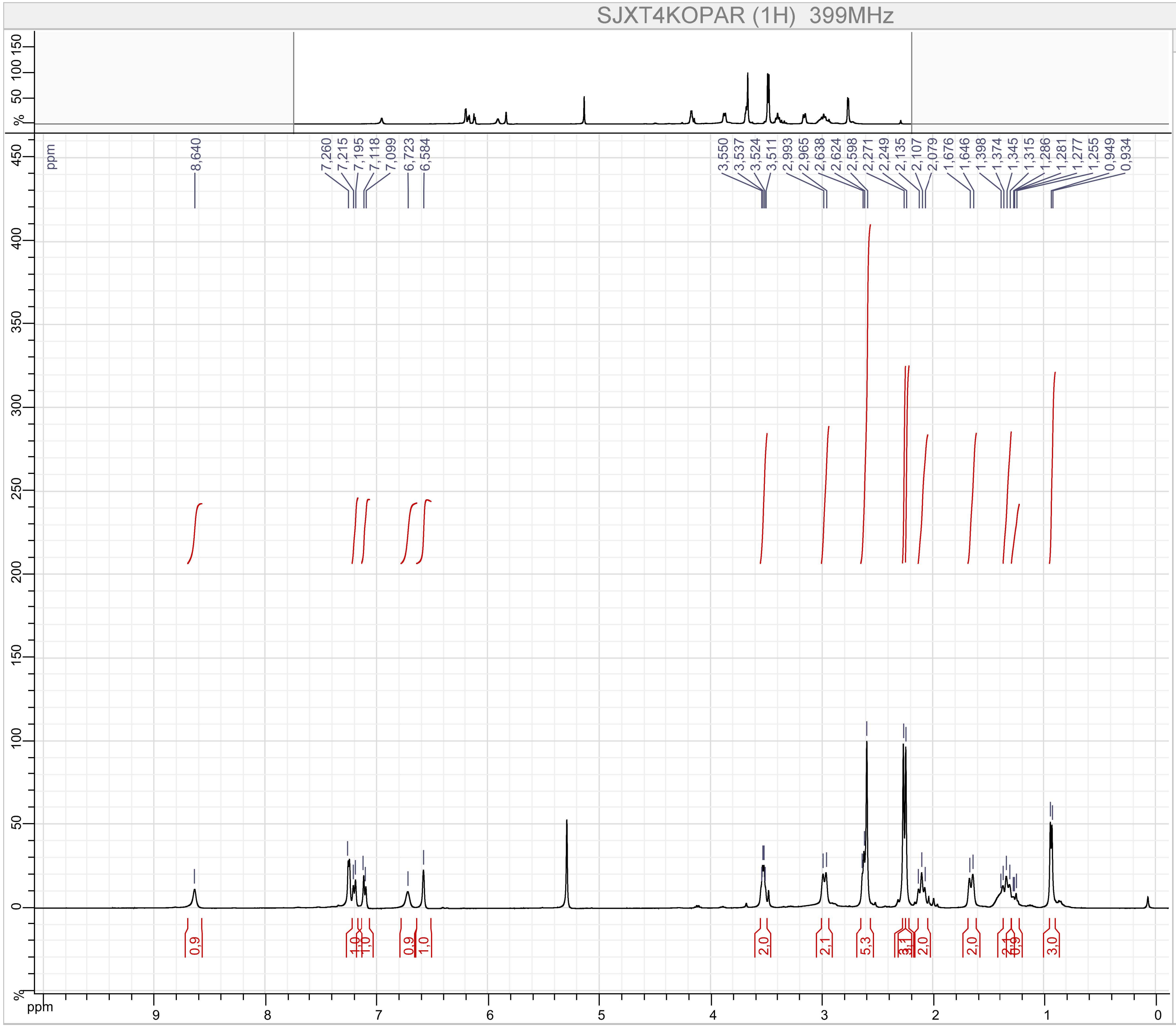


Parameters

SJXT4KOPAR (1H)

F1 :
- Size : 65536 points complex
- Spectral Width : 7494.2334 Hz
- Carrier Frequency : 399.7822 MHz
- Nucleus : 1H

Type : UNKNOWN
Spectro : JEOL 399 MHz
Probe : Unknown
Date : Thu Oct 04 18:36:24 EDT 2018
Temperature : UNKNOWN
Solvent :

Compound 29

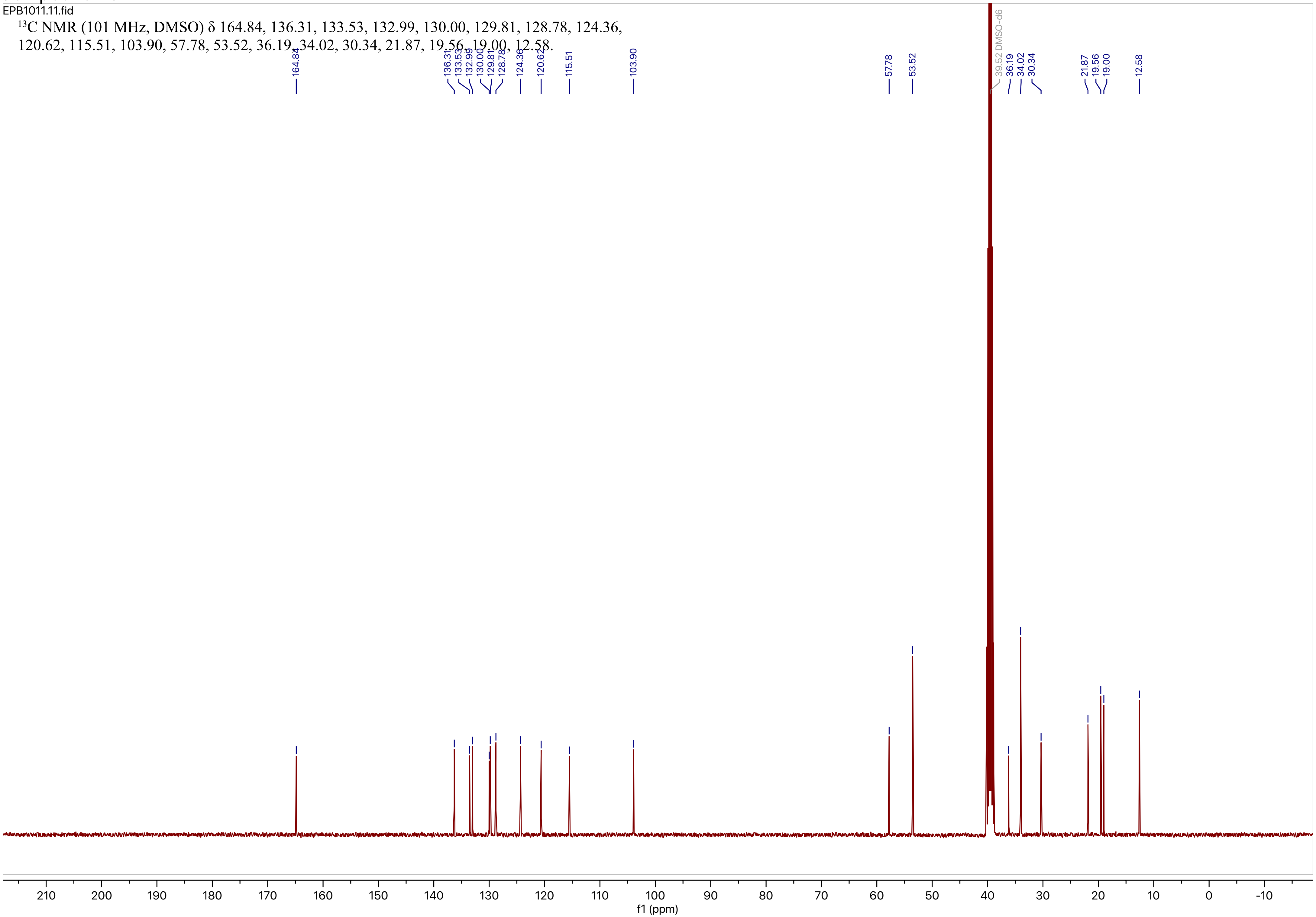


$^{13}C$ NMR (101 MHz, DMSO) δ 164.84, 136.31, 133.53, 132.99, 130.00, 129.81, 128.78, 124.36, 120.62, 115.51, 103.90, 57.78, 53.52, 36.19, 34.02, 30.34, 21.87, 19.56, 19.00, 12.58.

Compound 30

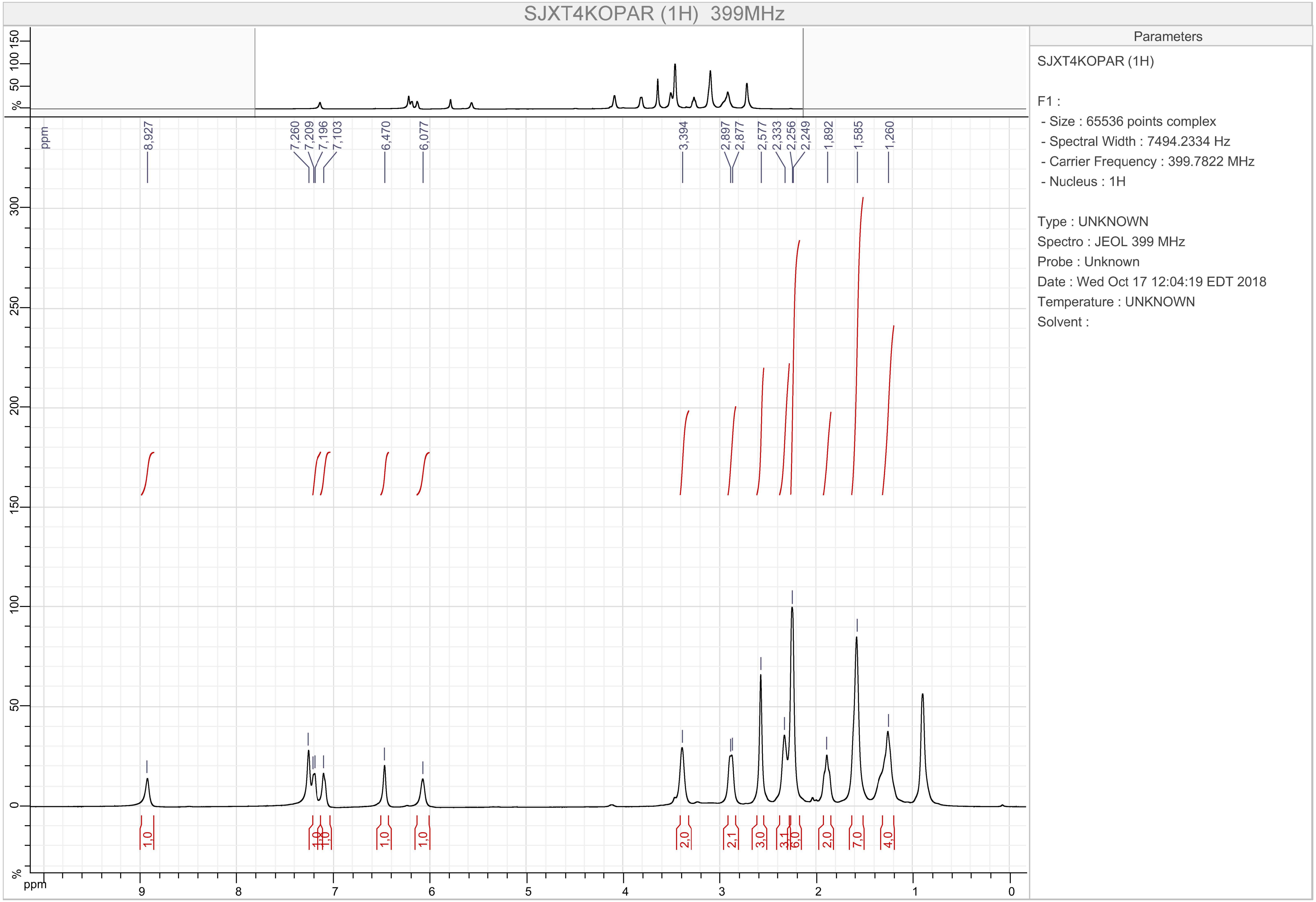

SJXT4KOPAR (1H) 399MHz
Parameters
SJXT4KOPAR (1H)
F1 :
- Size : 65536 points complex
- Spectral Width : 7494.2334 Hz
- Carrier Frequency : 399.7822 MHz
- Nucleus : 1H
Type : UNKNOWN
Spectro : JEOL 399 MHz
Probe : Unknown
Date : Wed Oct 17 12:04:19 EDT 2018
Temperature : UNKNOWN
Solvent :
8,927
7,260
7,209
7,196
7,103
6,470
6,077
3,394
2,897
2,877
2,577
2,333
2,256
2,249
1,892
1,585
1,260
1,0
1,0
1,0
1,0
1,0
1,0
2,0
2,1
3,0
3,1
6,0
2,0
7,0
4,0
ppm

Compound 30

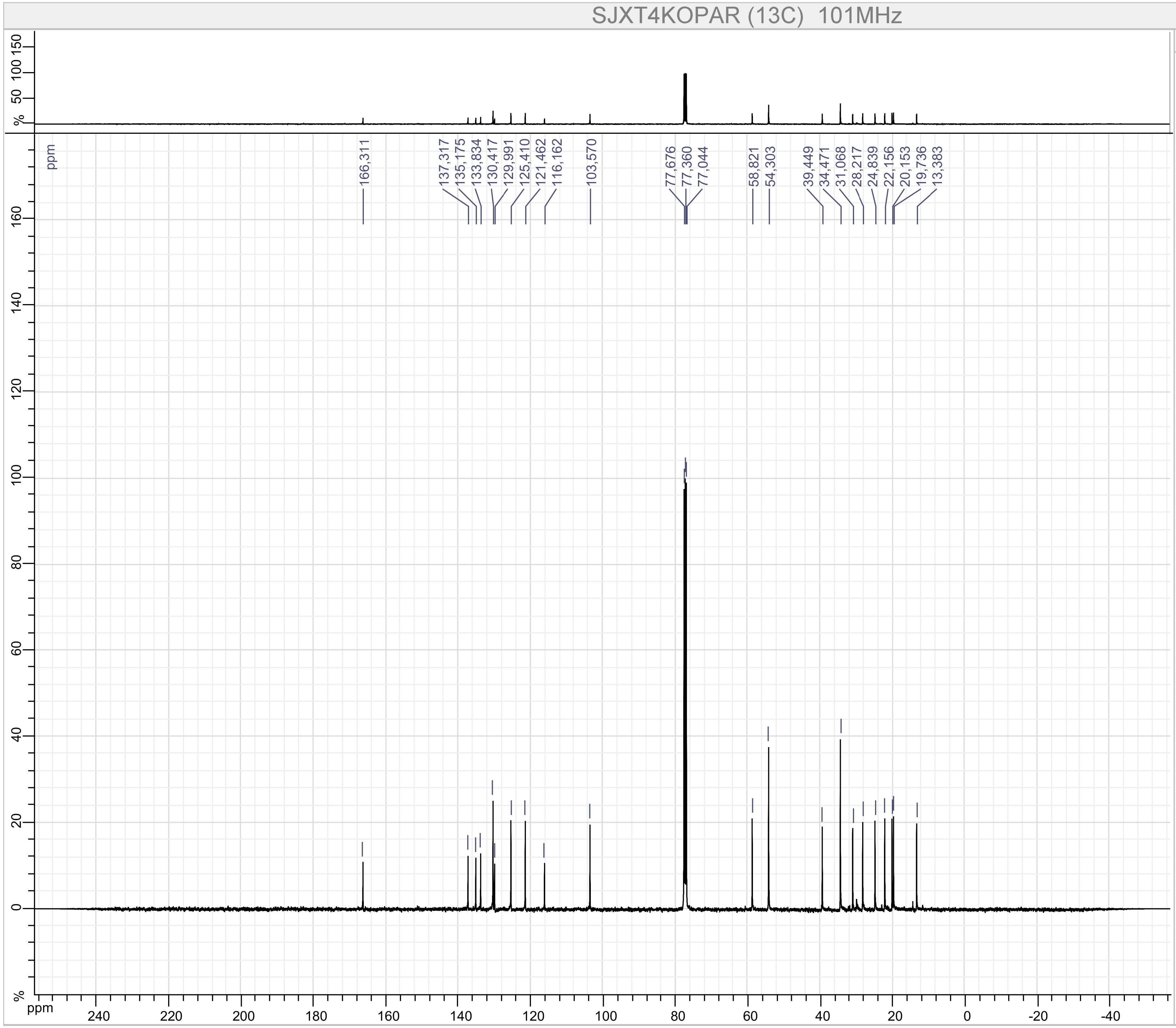


Parameters

SJXT4KOPAR (13C)

F1 :
- Size : 65536 points complex
- Spectral Width : 31566.62 Hz
- Carrier Frequency : 100.52531 MHz
- Nucleus : 13C

Type : UNKNOWN
Spectro : JEOL 101 MHz
Probe : Unknown
Date : Wed Oct 17 12:06:45 EDT 2018
Temperature : UNKNOWN
Solvent :

Compound 31

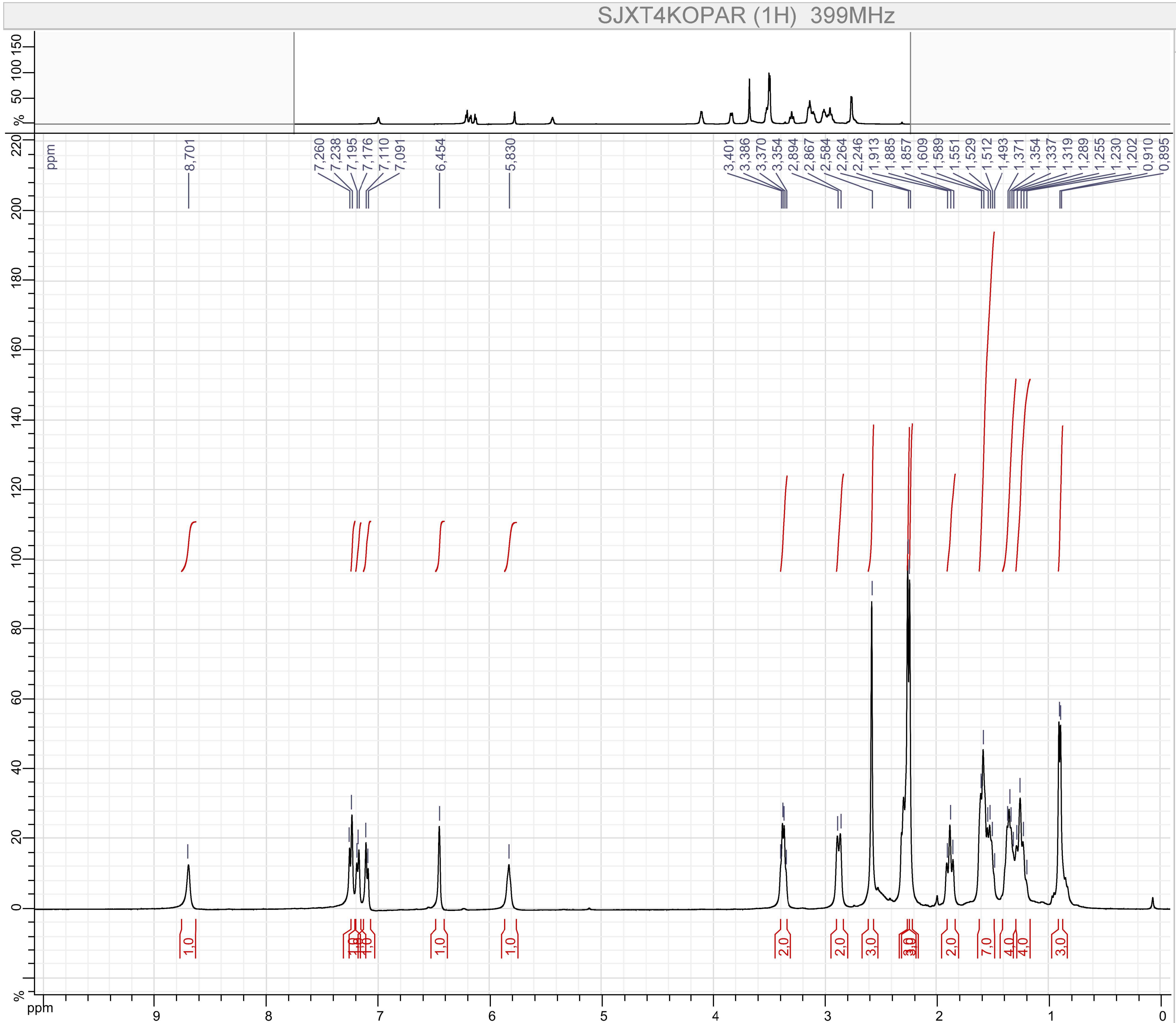


Parameters

SJXT4KOPAR (1H)

F1 :
- Size : 65536 points complex
- Spectral Width : 7494.2334 Hz
- Carrier Frequency : 399.7822 MHz
- Nucleus : 1H

Type : UNKNOWN
Spectro : JEOL 399 MHz
Probe : Unknown
Date : Fri Sep 06 21:55:53 EDT 2019
Temperature : UNKNOWN
Solvent :

Compound 31

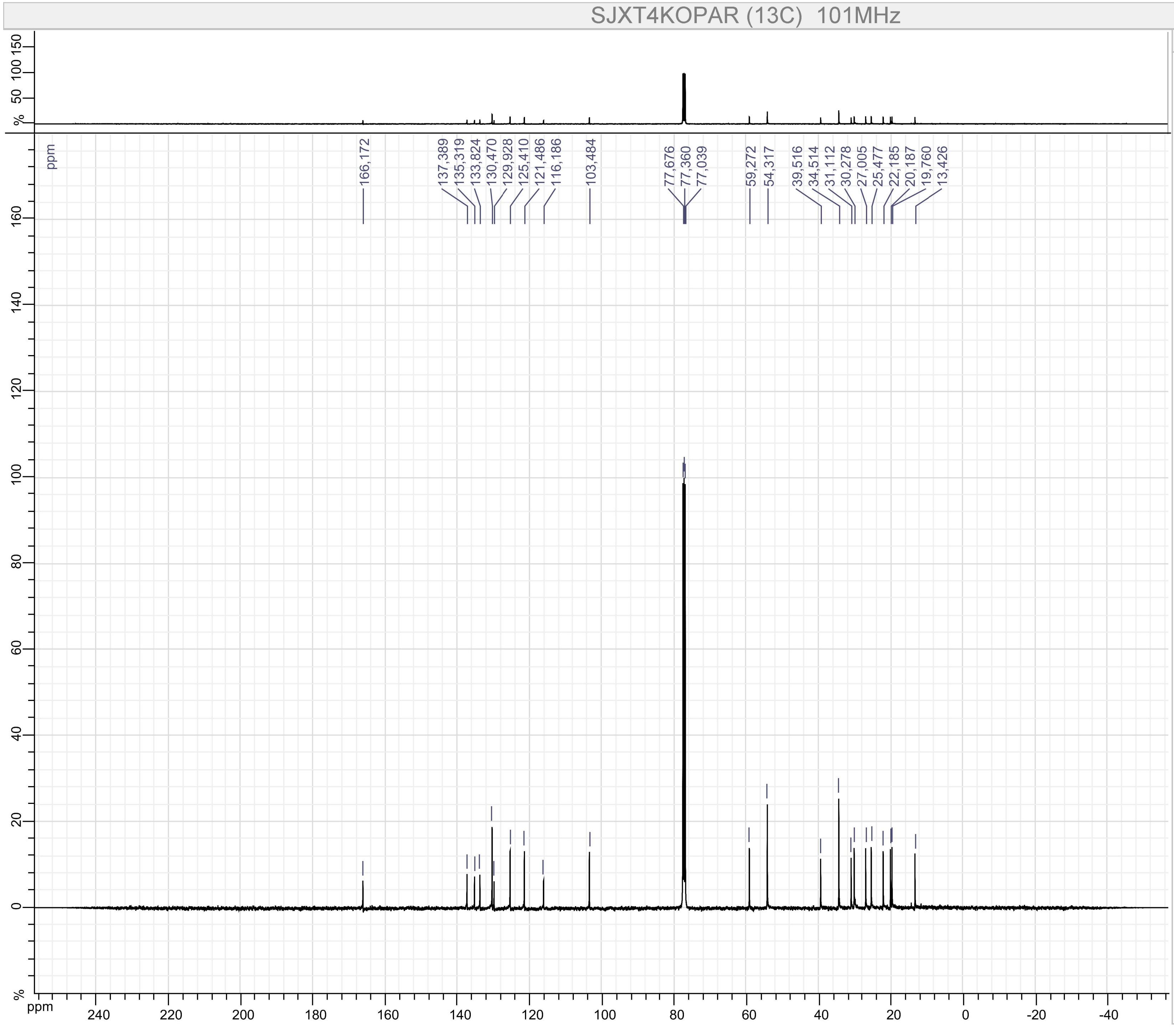


Parameters

SJXT4KOPAR (13C)

F1 :
- Size : 65536 points complex
- Spectral Width : 31566.62 Hz
- Carrier Frequency : 100.52531 MHz
- Nucleus : 13C

Type : UNKNOWN
Spectro : JEOL 101 MHz
Probe : Unknown
Date : Fri Sep 06 21:59:16 EDT 2019
Temperature : UNKNOWN
Solvent :

Compound 34

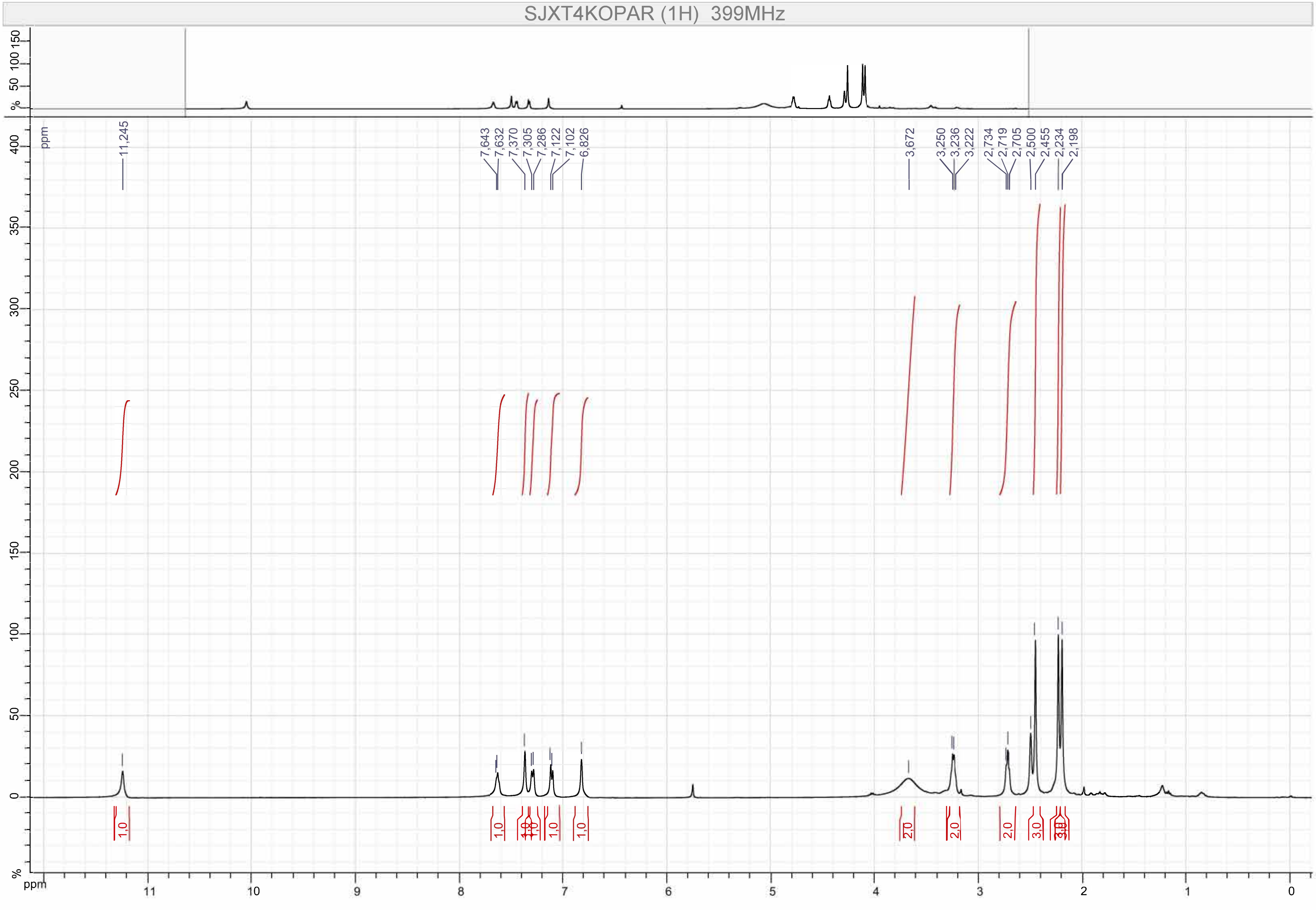

SJXT4KOPAR (1H) 399MHz
11,245
7,643
7,632
7,370
7,305
7,286
7,122
7,102
6,826
3,672
3,250
3,236
3,222
2,734
2,719
2,705
2,500
2,455
2,234
2,198
1,0
1,0
1,0
1,0
1,0
1,0
2,0
2,0
2,0
3,0
3,0
ppm

Compound 34

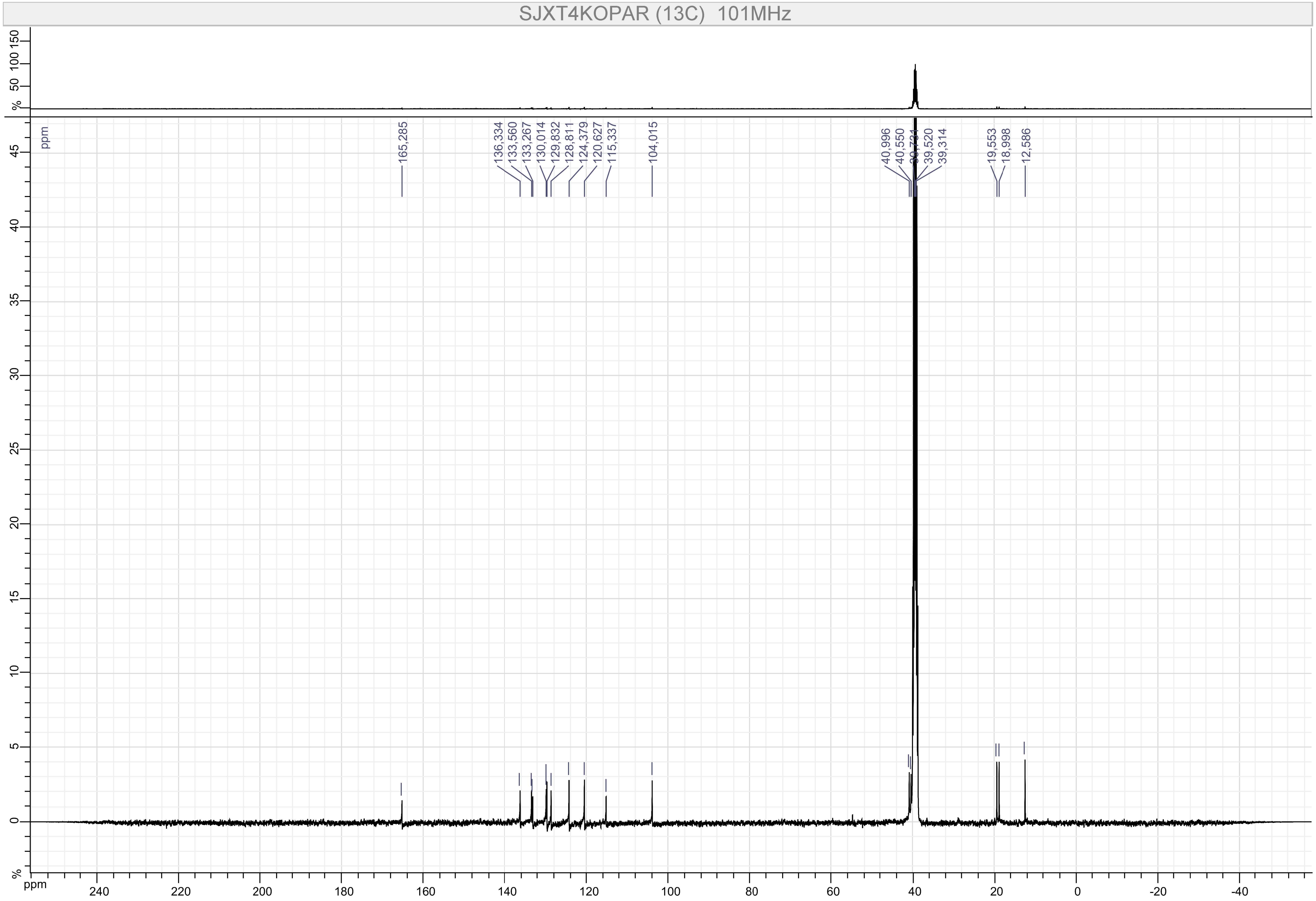

SJXT4KOPAR (13C) 101MHz
165,285
136,334
133,560
133,267
130,014
129,832
128,811
124,379
120,627
115,337
104,015
40,996
40,550
39,520
39,314
19,553
18,998
12,586
ppm

Compound 35

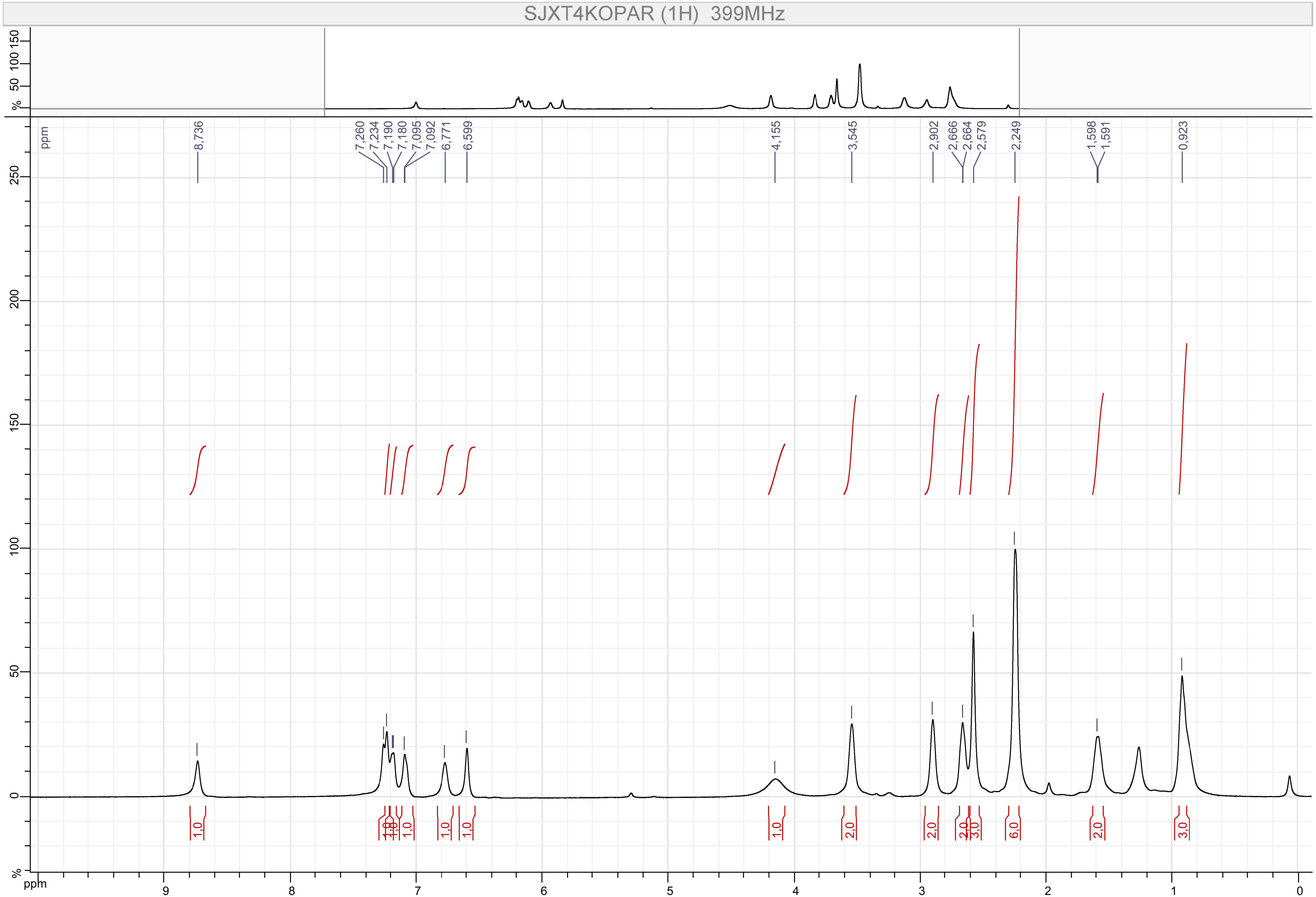

SJXT4KOPAR (1H) 399MHz
8,736
7,260
7,234
7,190
7,180
7,095
7,092
6,771
6,599
4,155
3,545
2,902
2,666
2,664
2,579
2,249
1,598
1,591
0,923
ppm
%

Compound 35

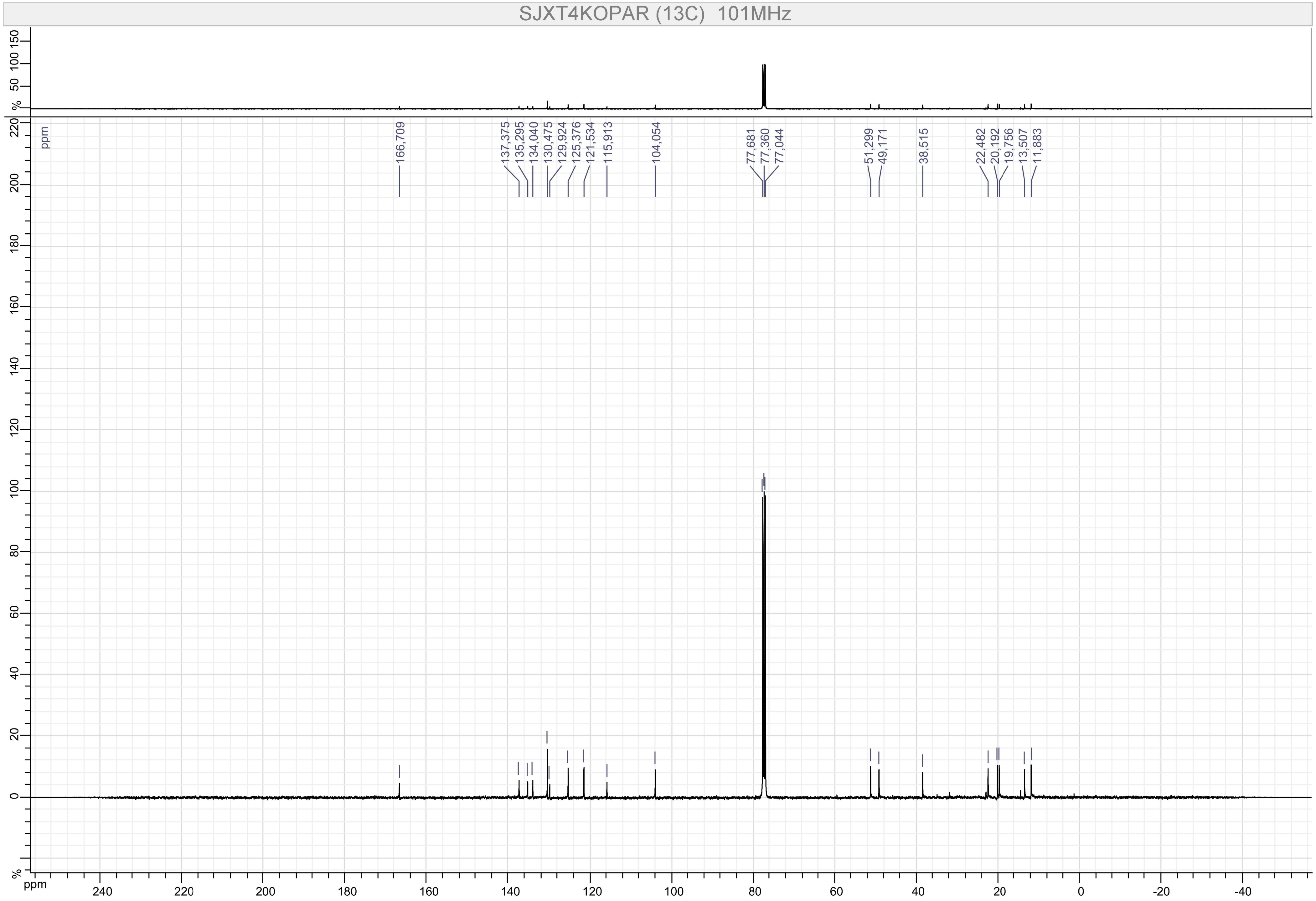

SJXT4KOPAR (13C) 101MHz
166,709
137,375
135,295
134,040
130,475
129,924
125,376
121,534
115,913
104,054
77,681
77,360
77,044
51,299
49,171
38,515
22,482
20,192
19,756
13,507
11,883
ppm
%

Compound 36

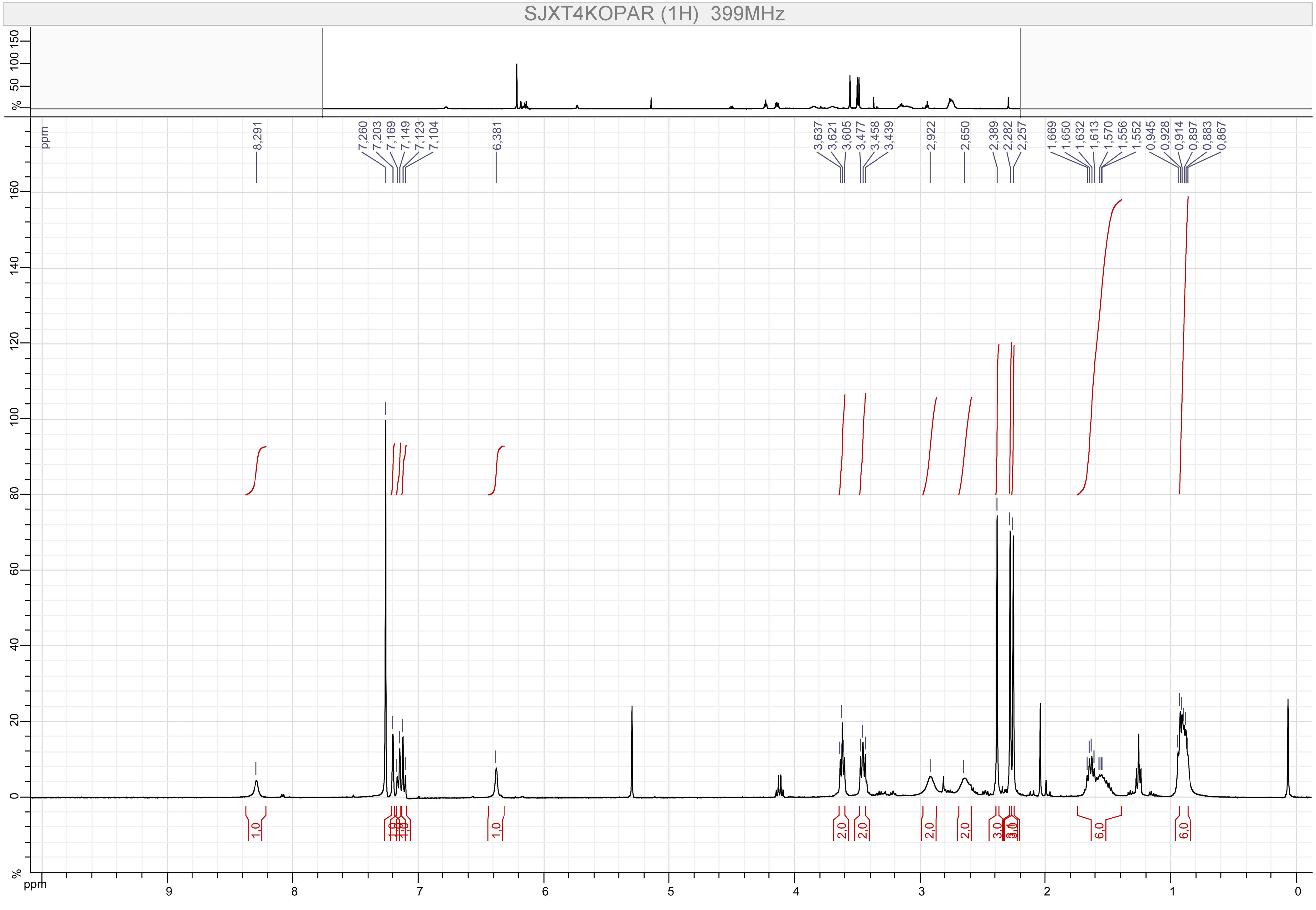

SJXT4KOPAR (1H) 399MHz
8,291
7,260
7,203
7,169
7,149
7,123
7,104
6,381
3,637
3,621
3,605
3,477
3,458
3,439
2,922
2,650
2,389
2,282
2,257
1,669
1,650
1,632
1,613
1,570
1,556
1,552
0,945
0,928
0,914
0,897
0,883
0,867
1,0
1,0
2,0
2,0
2,0
2,0
3,0
6,0
6,0
ppm

Compound 36

MB-A-099

$^{13}$C NMR (101 MHz, CHLOROFORM-*D*) δ 166.84, 137.24, 135.06, 130.30, 130.19, 125.17, 122.42, 121.27, 104.91, 60.55, 48.24, 20.04, 19.58, 14.35, 12.64, 11.78, 11.31.

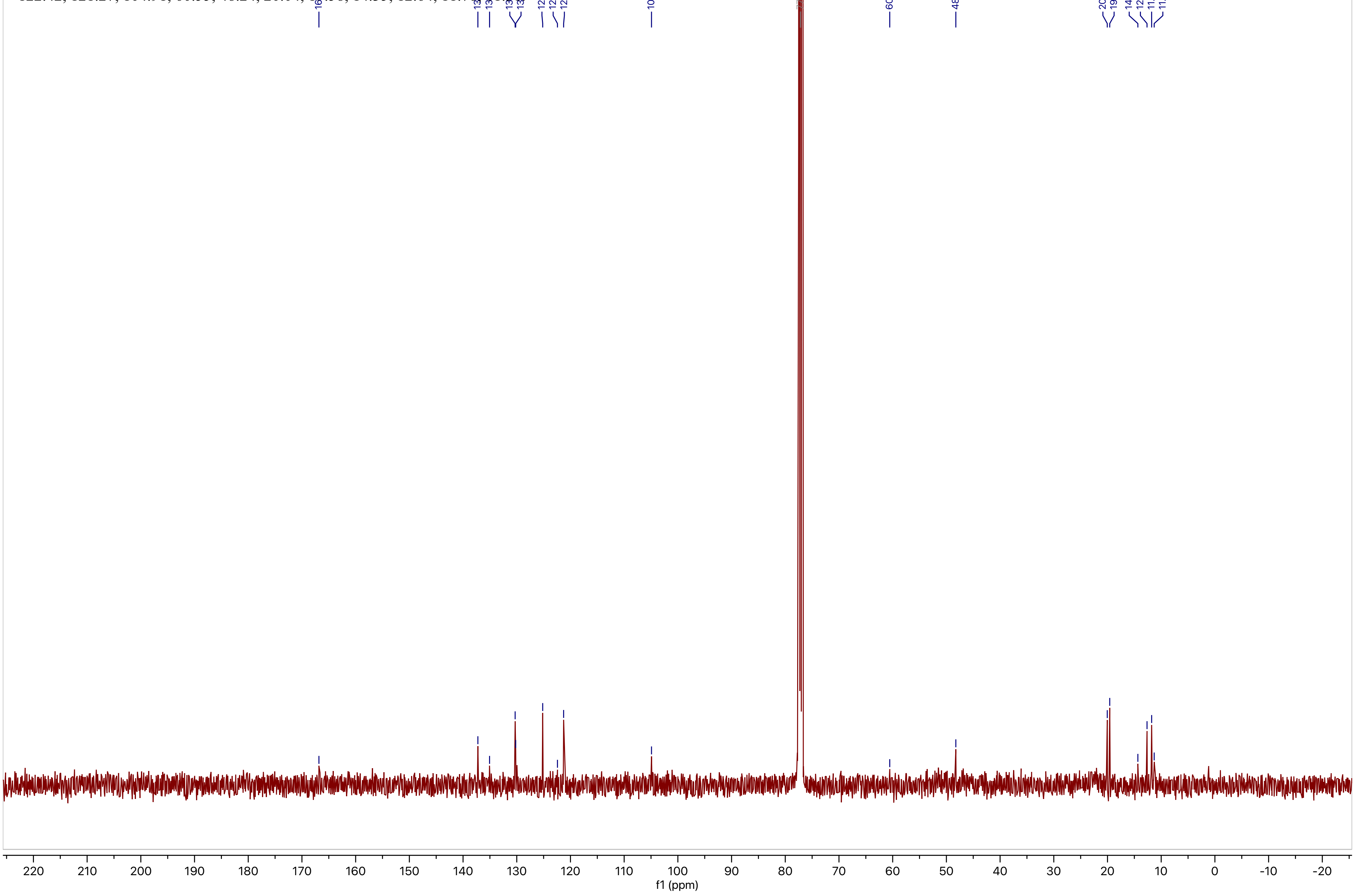

Compound 37

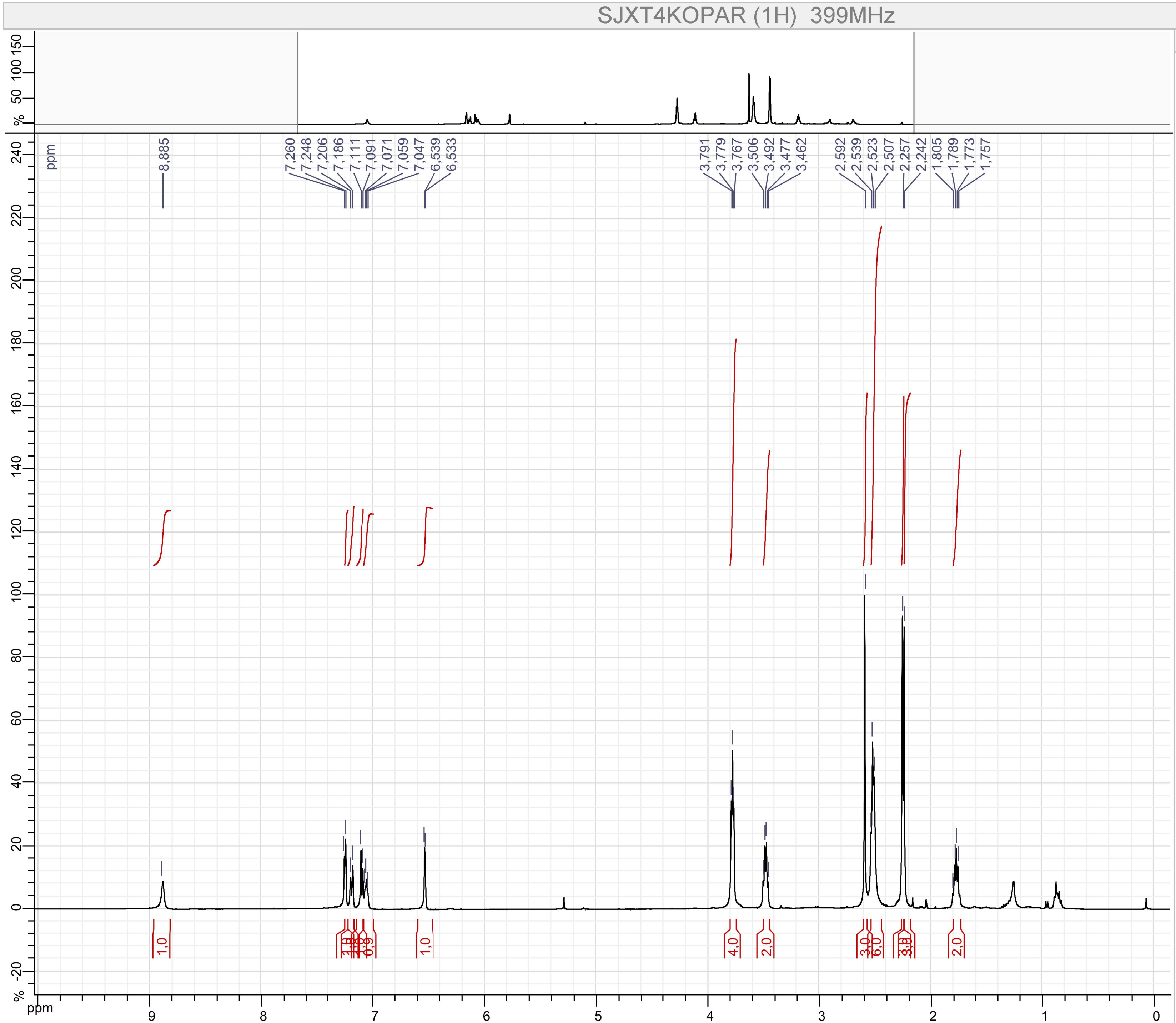


Parameters

SJXT4KOPAR (1H)

F1 :
- Size : 65536 points complex
- Spectral Width : 7494.2334 Hz
- Carrier Frequency : 399.7822 MHz
- Nucleus : 1H

Type : UNKNOWN
Spectro : JEOL 399 MHz
Probe : Unknown
Date : Fri Jul 12 22:58:34 EDT 2019
Temperature : UNKNOWN
Solvent :

Compound 37

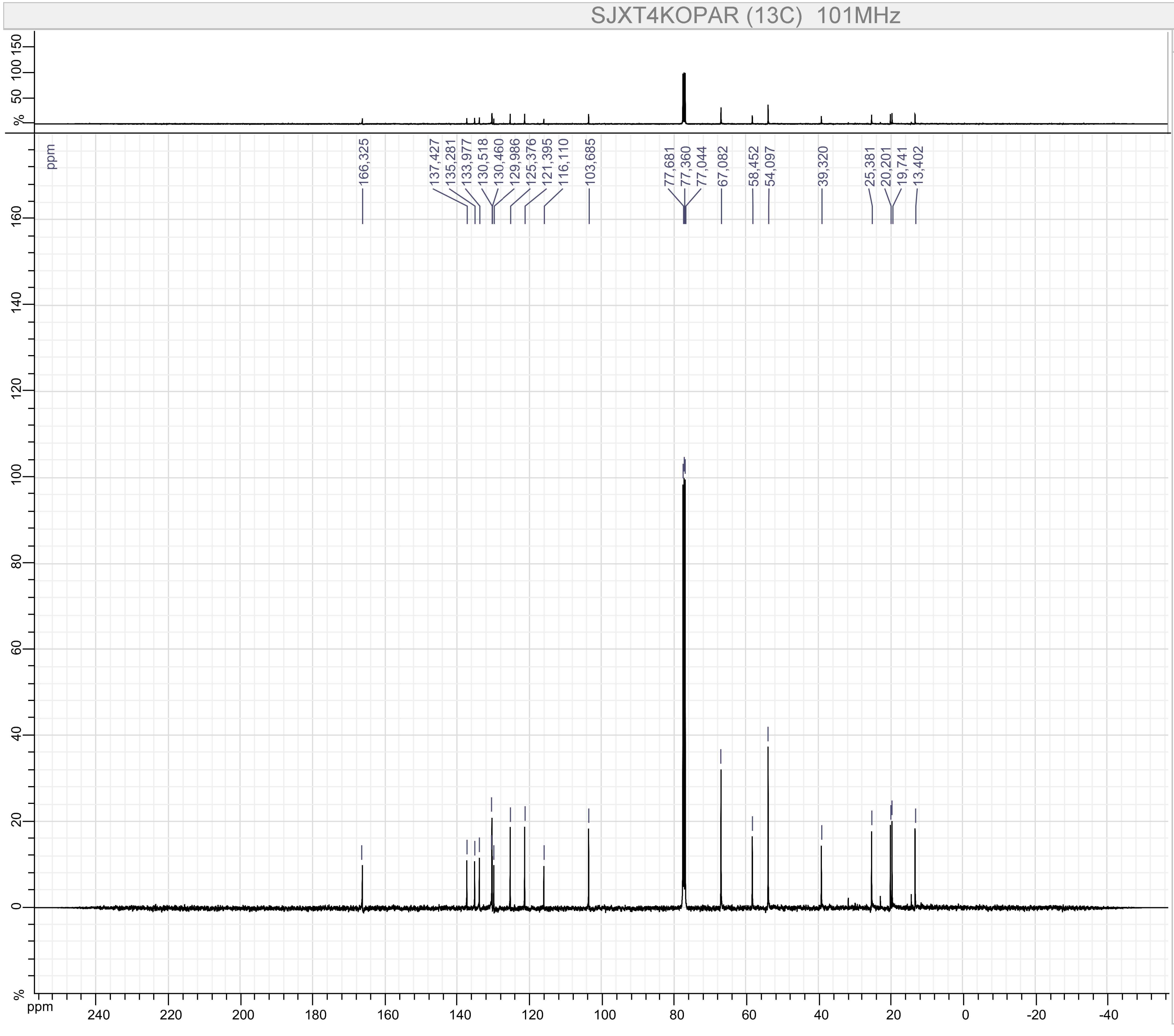


Parameters

SJXT4KOPAR (13C)

F1 :
- Size : 65536 points complex
- Spectral Width : 31566.62 Hz
- Carrier Frequency : 100.52531 MHz
- Nucleus : 13C

Type : UNKNOWN
Spectro : JEOL 101 MHz
Probe : Unknown
Date : Fri Jul 12 23:01:00 EDT 2019
Temperature : UNKNOWN
Solvent :

Compound 39

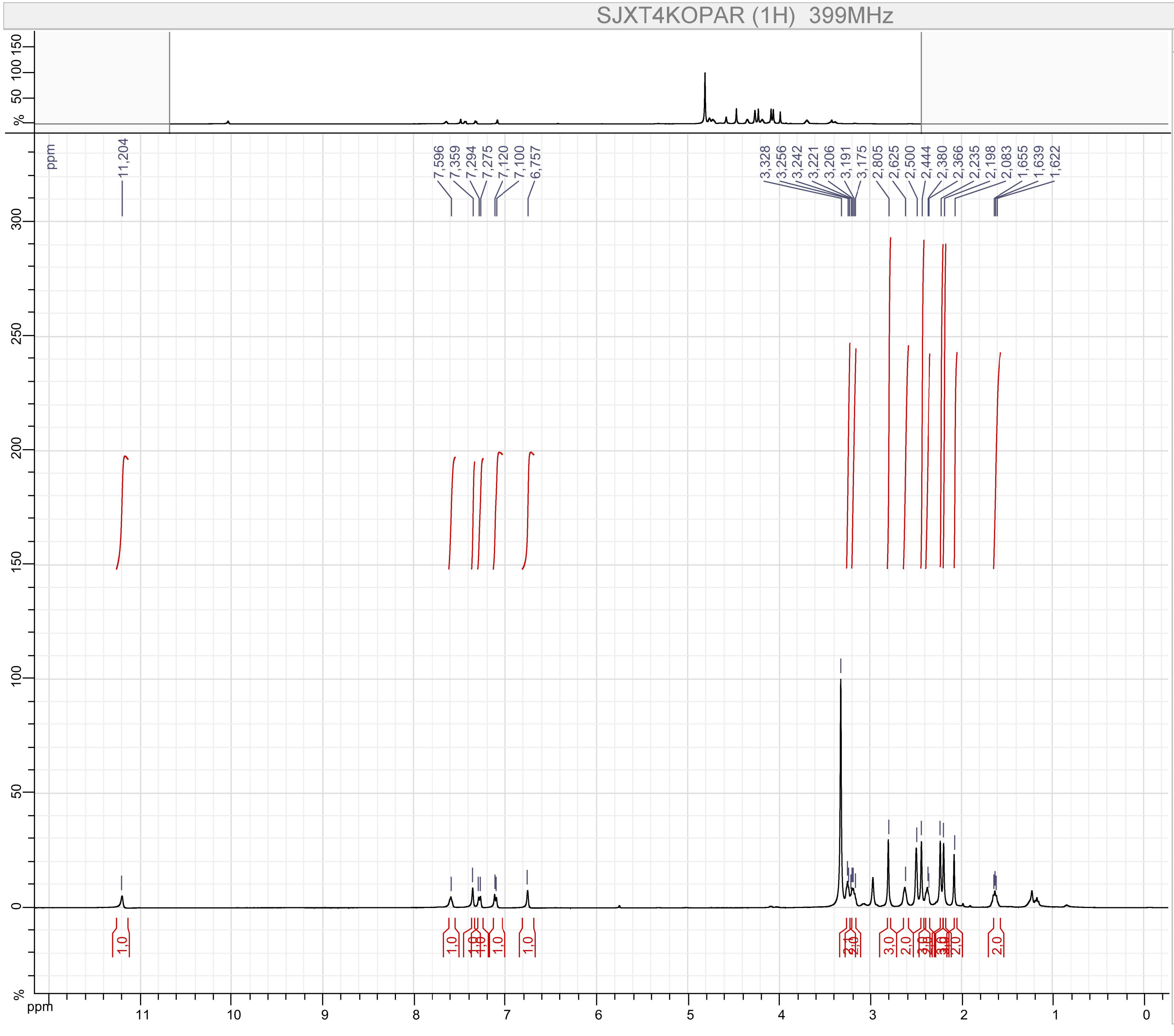


Parameters

SJXT4KOPAR (1H)

F1 :
- Size : 65536 points complex
- Spectral Width : 7494.2334 Hz
- Carrier Frequency : 399.7822 MHz
- Nucleus : 1H

Type : UNKNOWN
Spectro : JEOL 399 MHz
Probe : Unknown
Date : Mon Aug 26 19:24:22 EDT 2019
Temperature : UNKNOWN
Solvent :

Compound 39

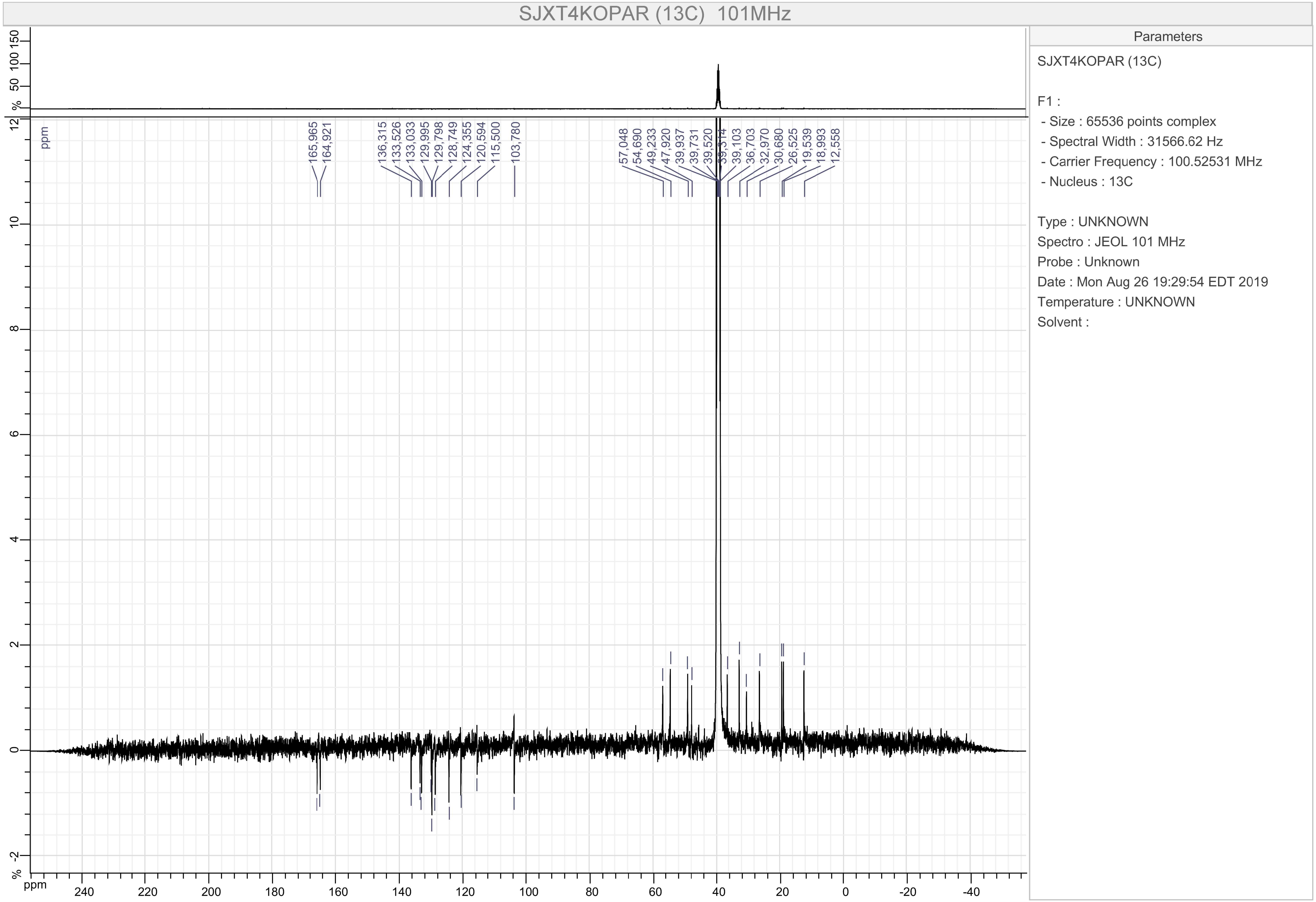
SJXT4KOPAR (13C) 101MHz
165,965
164,921
136,315
133,526
133,033
129,995
129,798
128,749
124,355
120,594
115,500
103,780
57,048
54,690
49,233
47,920
39,937
39,731
39,520
39,314
39,103
36,703
32,970
30,680
26,525
19,539
18,993
12,558
ppm
Parameters
SJXT4KOPAR (13C)
F1 :
- Size : 65536 points complex
- Spectral Width : 31566.62 Hz
- Carrier Frequency : 100.52531 MHz
- Nucleus : 13C
Type : UNKNOWN
Spectro : JEOL 101 MHz
Probe : Unknown
Date : Mon Aug 26 19:29:54 EDT 2019
Temperature : UNKNOWN
Solvent :

Compound 40

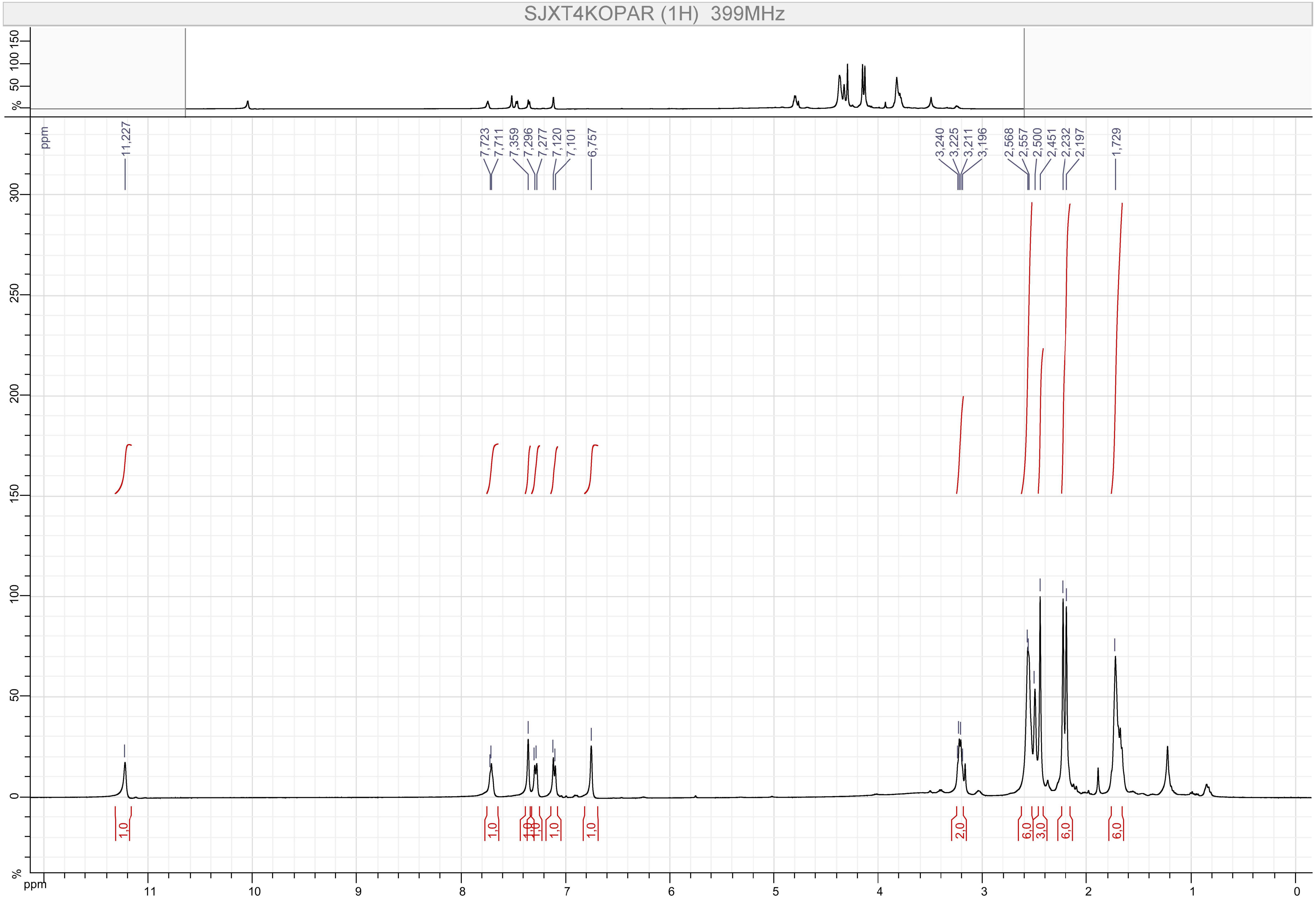
SJXT4KOPAR (1H) 399MHz
ppm
11,227
7,723
7,711
7,359
7,296
7,277
7,120
7,101
6,757
3,240
3,225
3,211
3,196
2,568
2,557
2,500
2,451
2,232
2,197
1,729
1,0
1,0
1,0
1,0
1,0
1,0
2,0
6,0
3,0
6,0
6,0
%
ppm

Compound 40

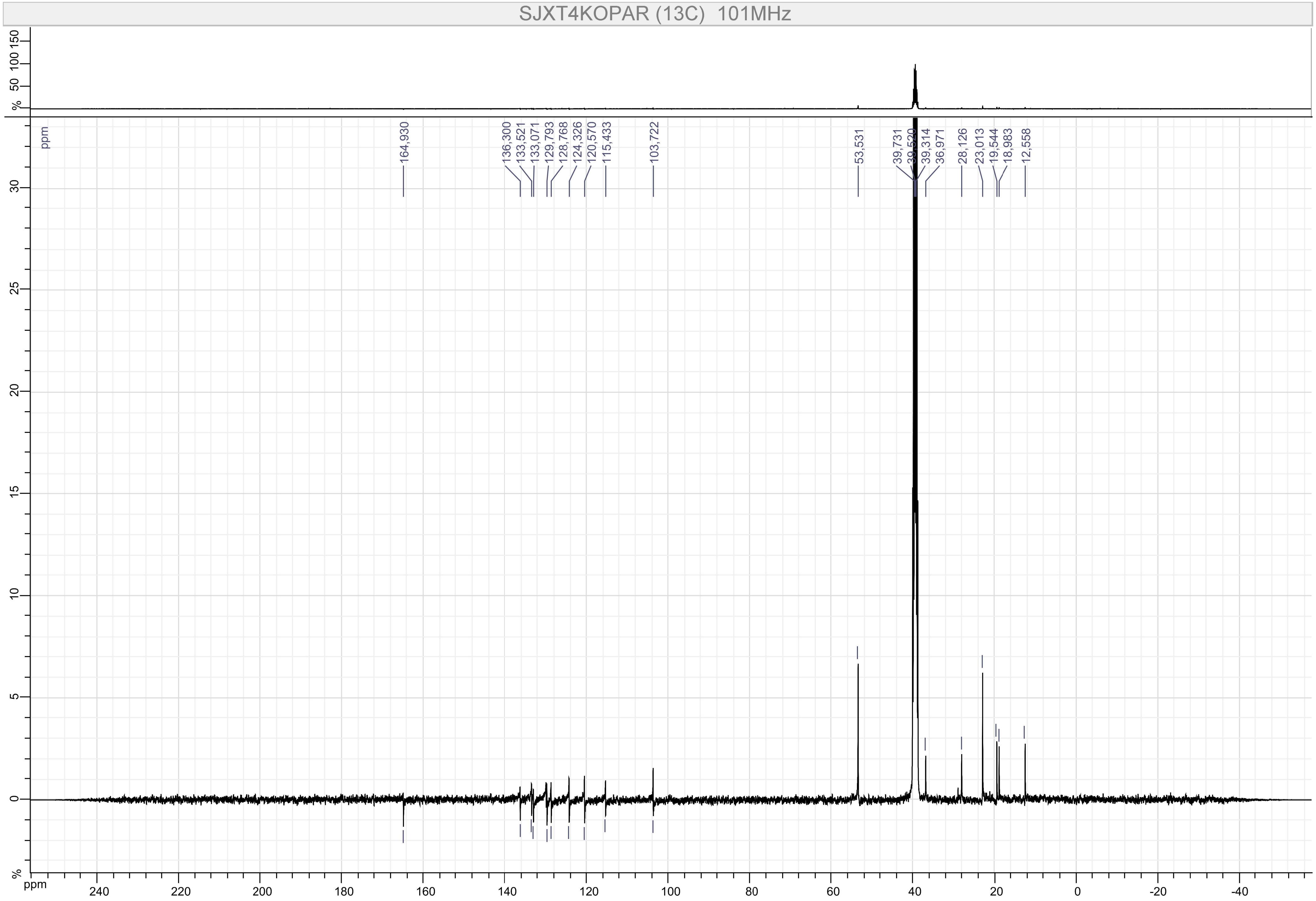
SJXT4KOPAR (13C) 101MHz
164,930
136,300
133,521
133,071
129,793
128,768
124,326
120,570
115,433
103,722
53,531
39,731
39,520
39,314
36,971
28,126
23,013
19,544
18,983
12,558
ppm
%

Compound 41

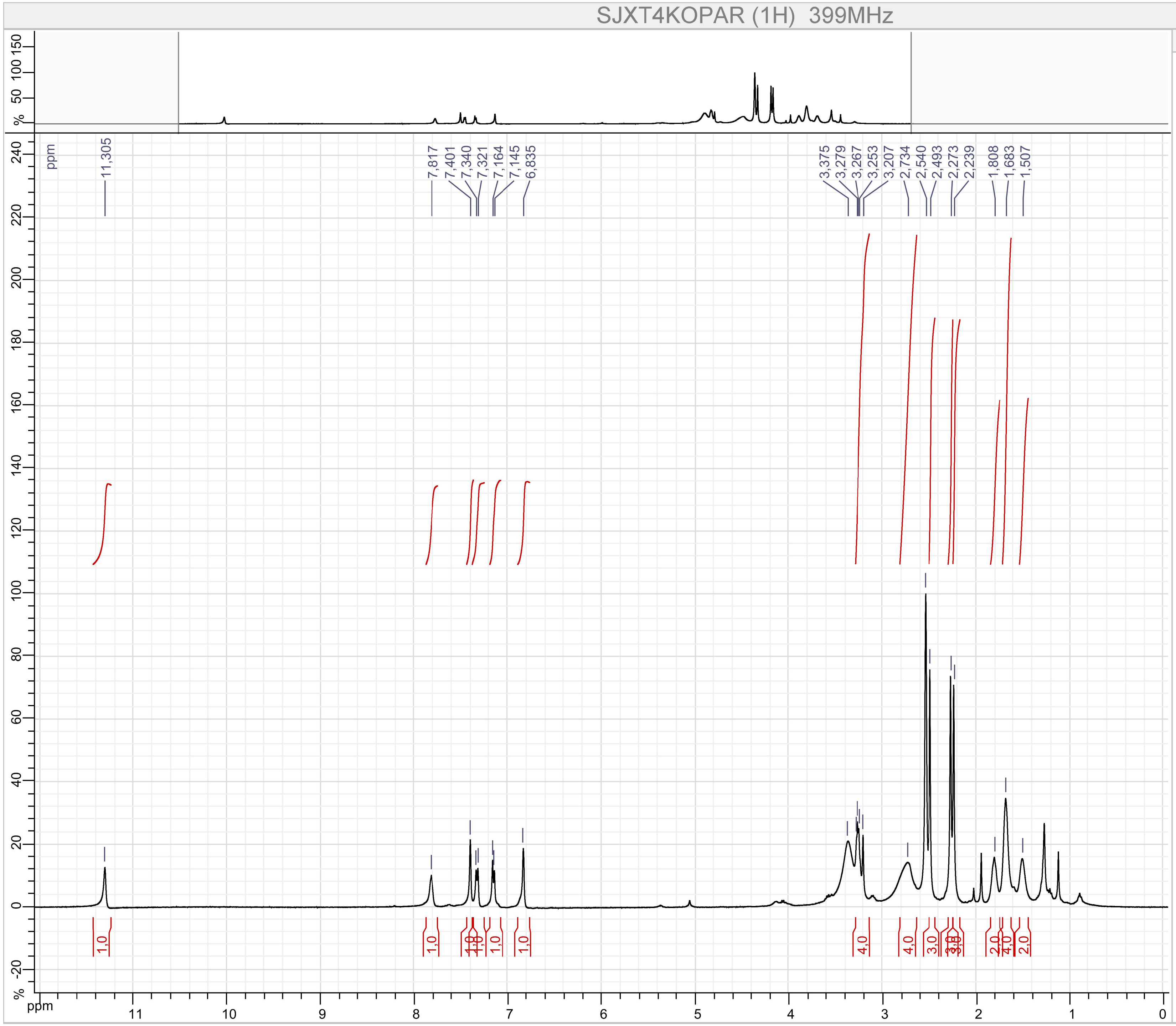


Parameters

SJXT4KOPAR (1H)

F1 :
- Size : 65536 points complex
- Spectral Width : 7494.2334 Hz
- Carrier Frequency : 399.7822 MHz
- Nucleus : 1H

Type : UNKNOWN
Spectro : JEOL 399 MHz
Probe : Unknown
Date : Thu Aug 16 18:25:03 EDT 2018
Temperature : UNKNOWN
Solvent :

Compound 41

AW-1002-600mhz.12.fid

$^{13}$C NMR (151 MHz, DMSO) δ 165.58, 136.54, 133.85, 133.61, 129.99, 129.06, 124.48, 120.75, 115.17, 103.93, 52.89, 36.25, 29.12, 25.12, 23.70, 21.20, 19.70, 19.14, 12.73.

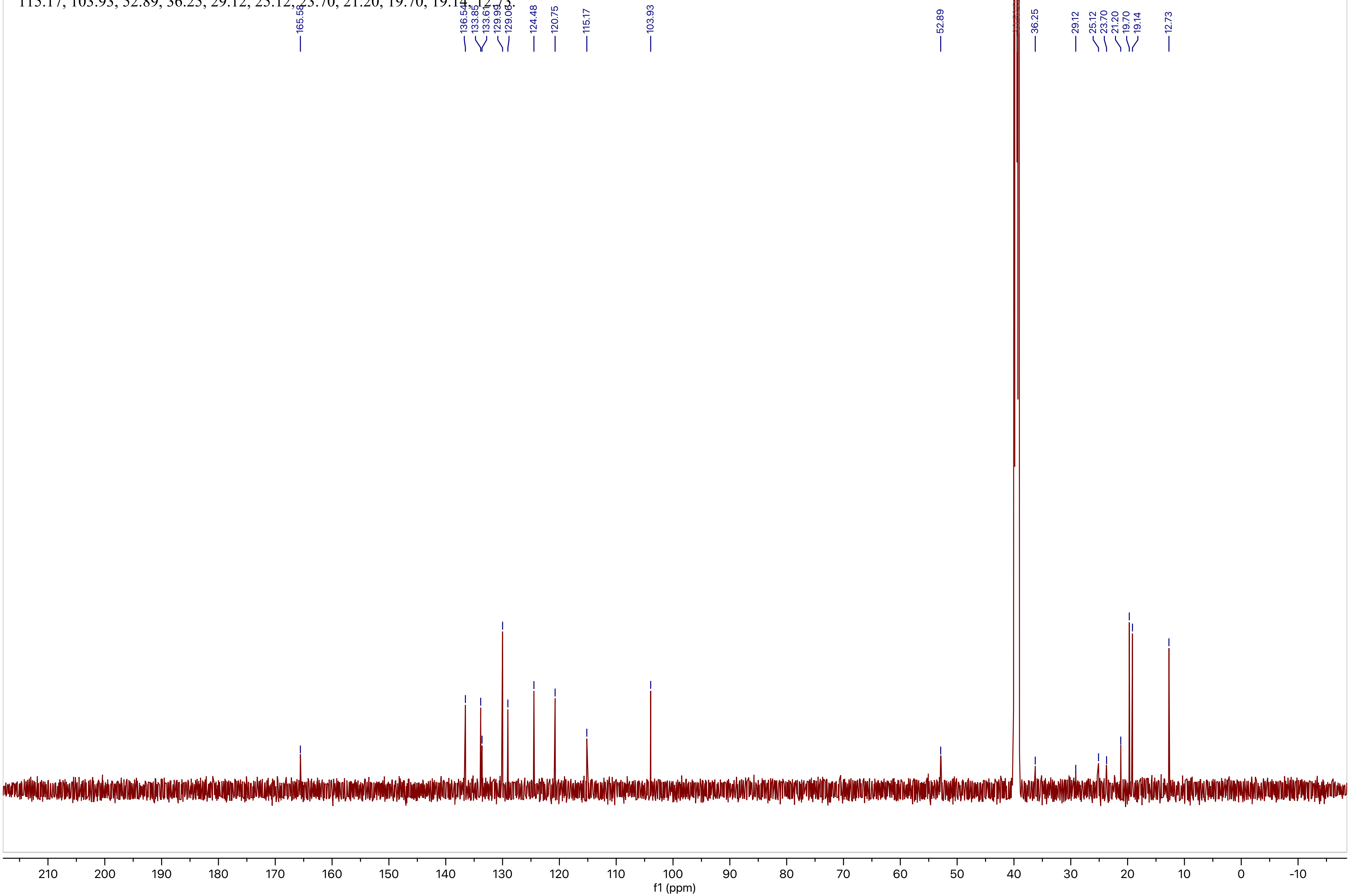

### Compound 42

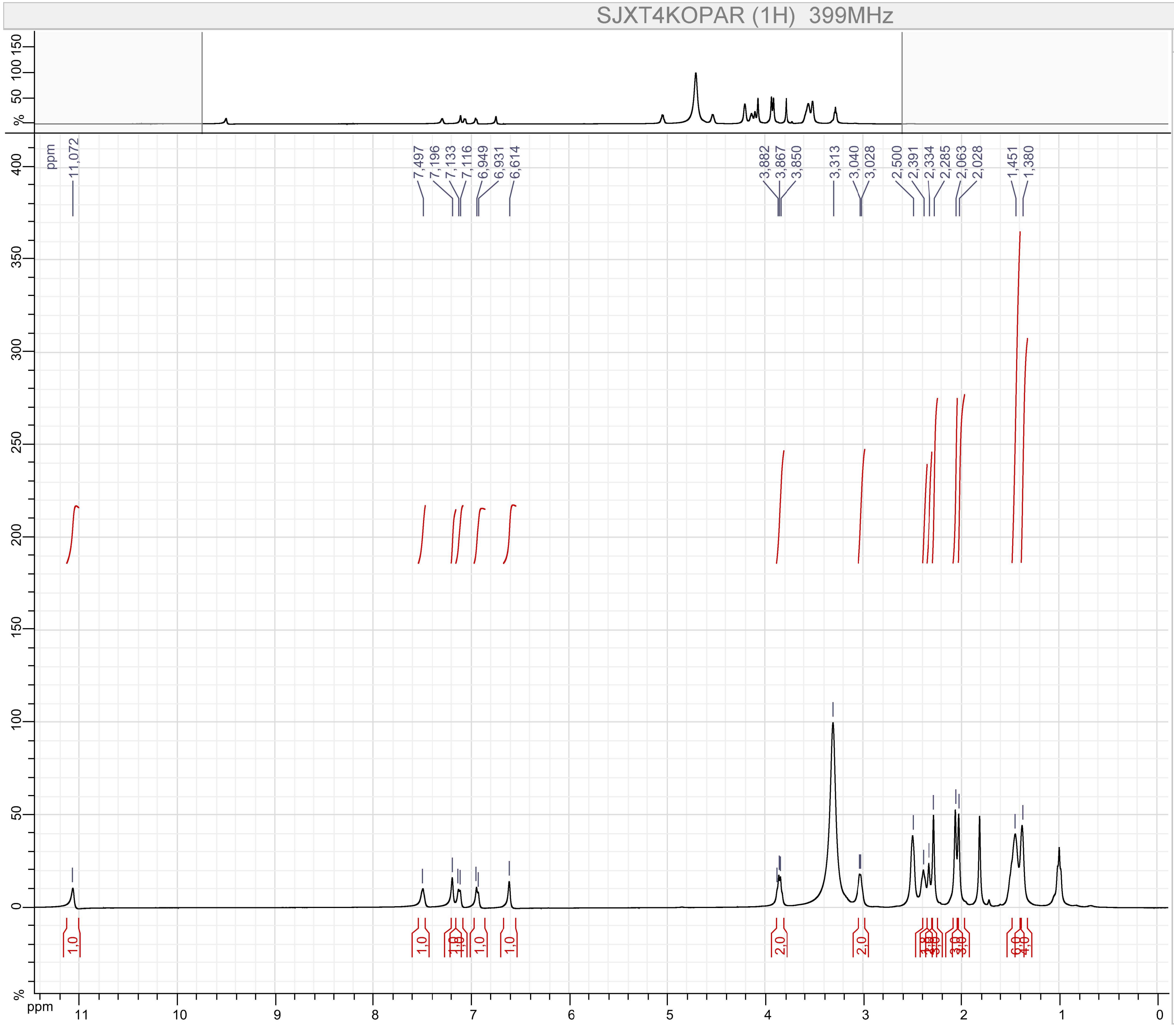


Parameters

SJXT4KOPAR (1H)

F1 :
- Size : 65536 points complex
- Spectral Width : 7494.2334 Hz
- Carrier Frequency : 399.7822 MHz
- Nucleus : 1H

Type : UNKNOWN
Spectro : JEOL 399 MHz
Probe : Unknown
Date : Tue Jul 02 22:12:10 EDT 2019
Temperature : UNKNOWN
Solvent :

Compound 42

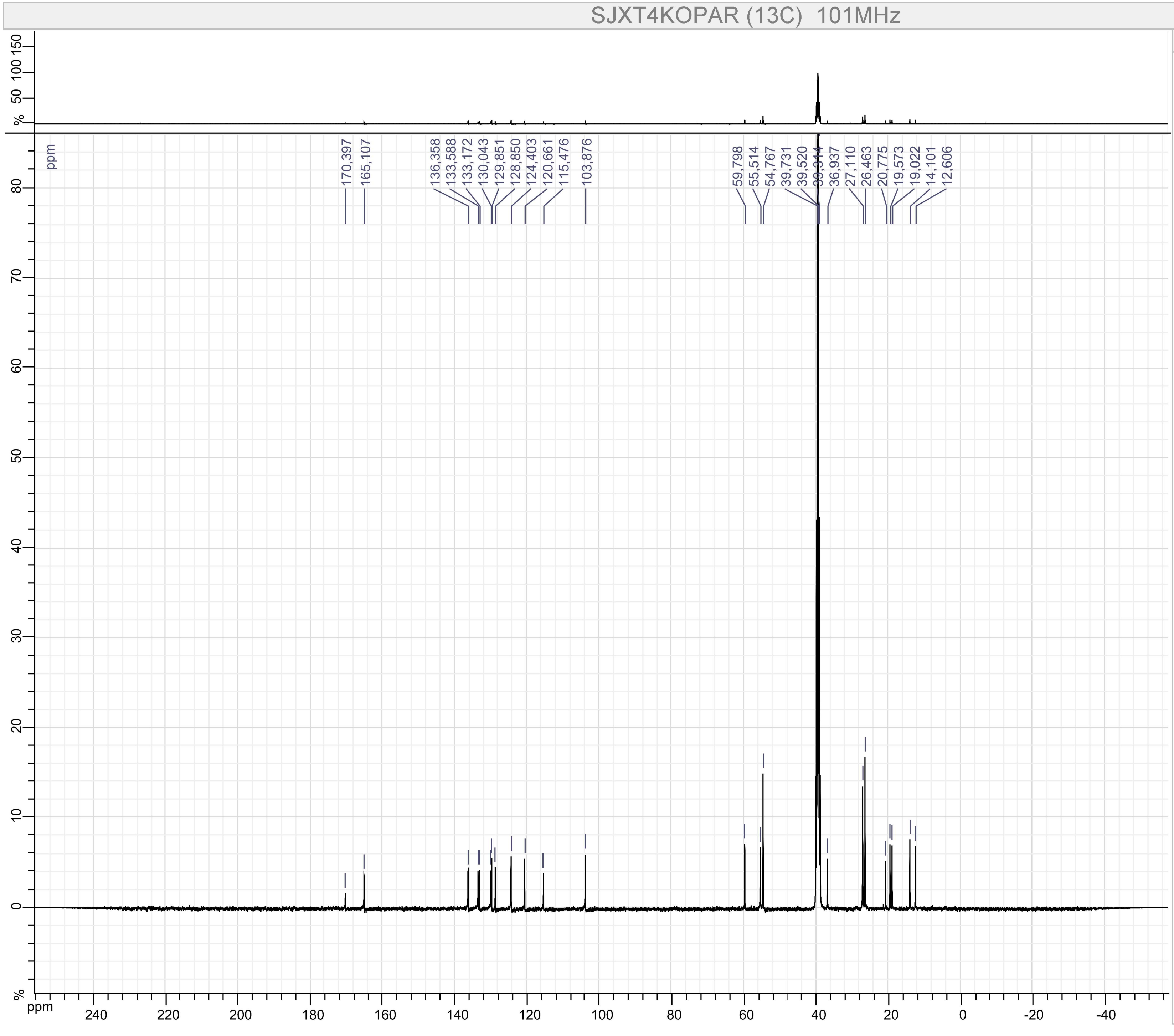


Parameters

SJXT4KOPAR (13C)

F1 :

- Size : 65536 points complex
- Spectral Width : 31566.62 Hz
- Carrier Frequency : 100.52531 MHz
- Nucleus : 13C

Type : UNKNOWN

Spectro : JEOL 101 MHz

Probe : Unknown

Date : Tue Jul 02 22:15:04 EDT 2019

Temperature : UNKNOWN

Solvent :

Compound 43

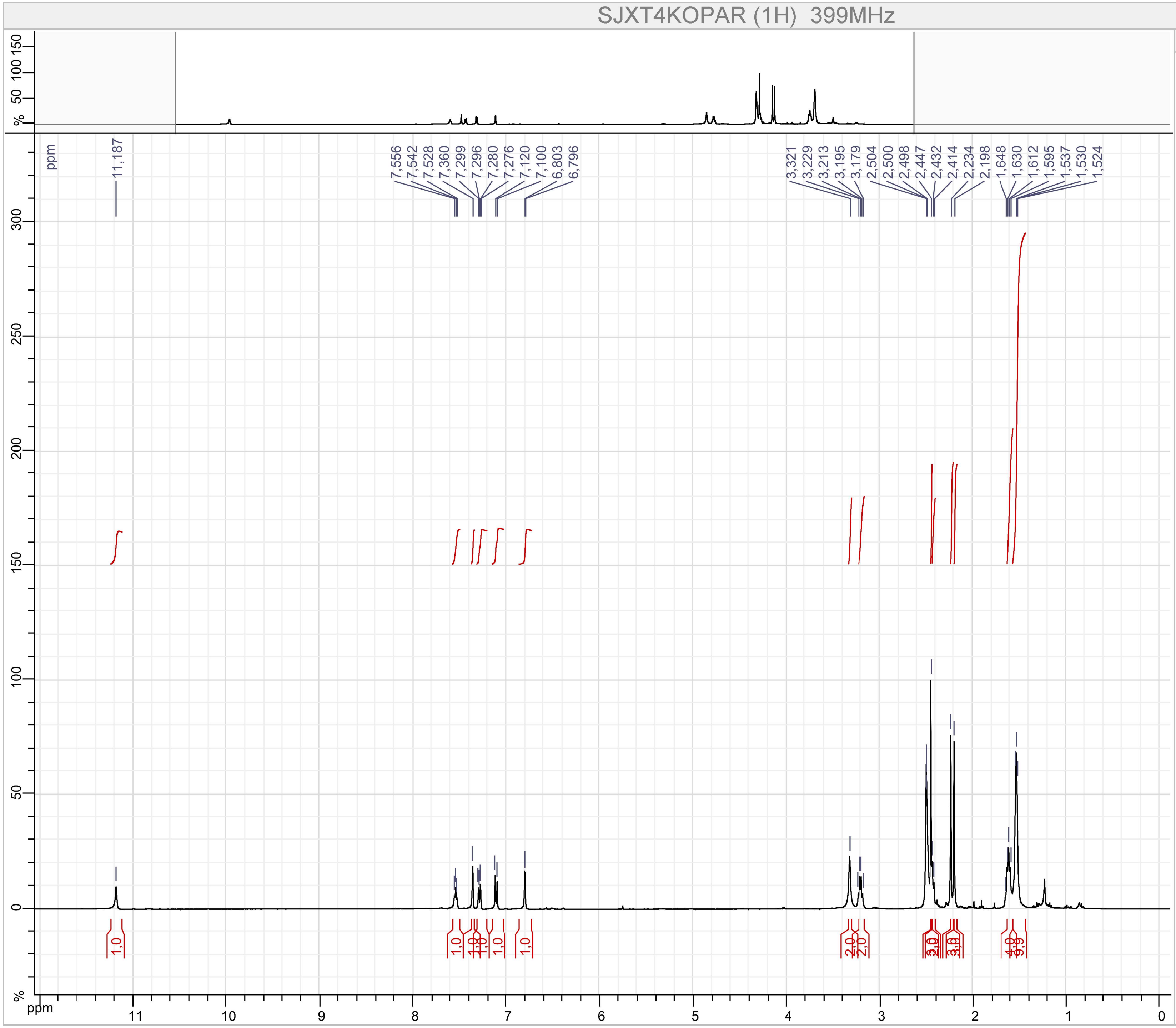


Parameters

SJXT4KOPAR (1H)

F1 :

- Size : 65536 points complex
- Spectral Width : 7494.2334 Hz
- Carrier Frequency : 399.7822 MHz
- Nucleus : 1H

Type : UNKNOWN

Spectro : JEOL 399 MHz

Probe : Unknown

Date : Sun Nov 25 21:14:57 EST 2018

Temperature : UNKNOWN

Solvent :

Compound 43

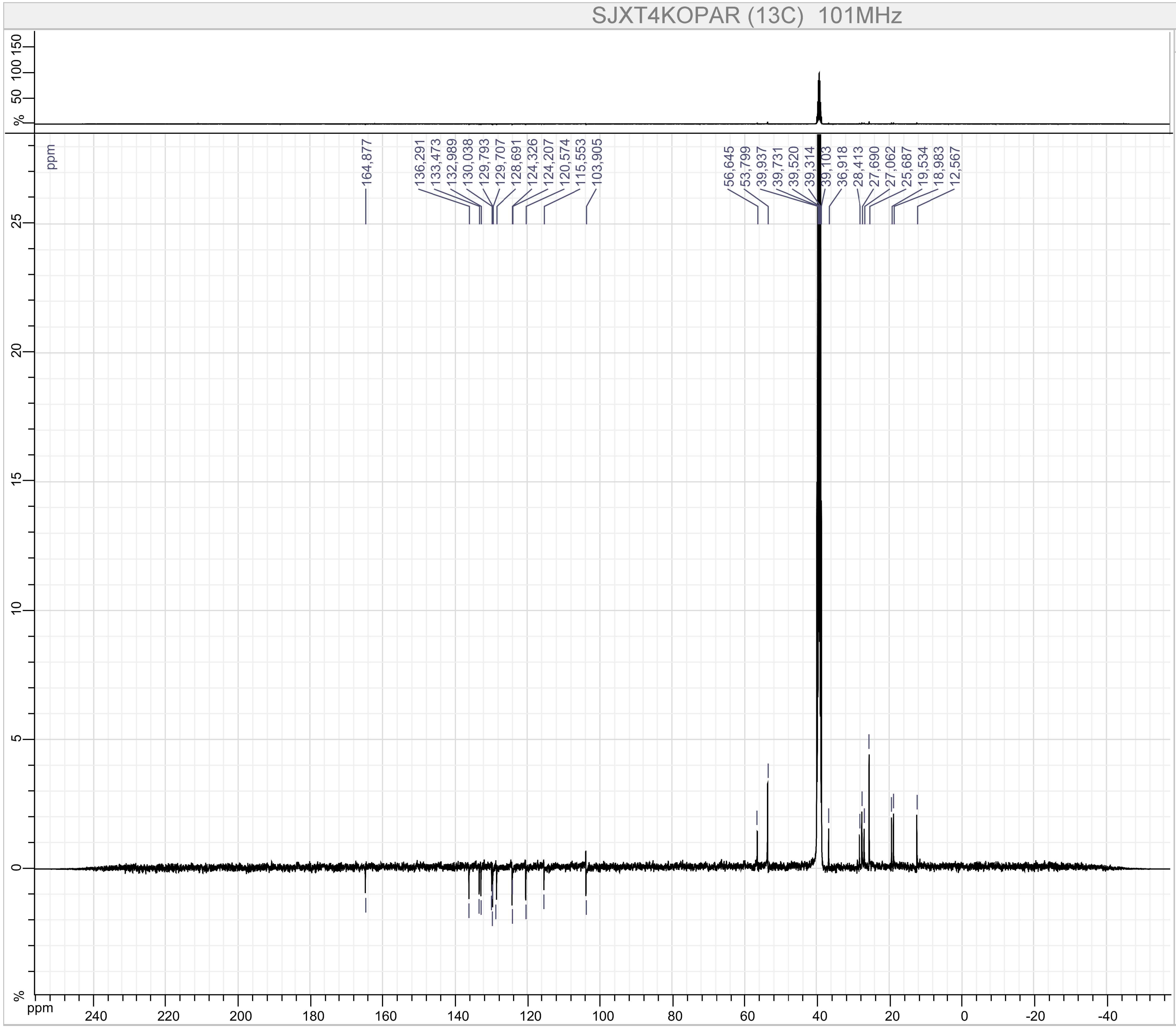


Parameters

SJXT4KOPAR (13C)

F1 :
- Size : 65536 points complex
- Spectral Width : 31566.62 Hz
- Carrier Frequency : 100.52531 MHz
- Nucleus : 13C

Type : UNKNOWN
Spectro : JEOL 101 MHz
Probe : Unknown
Date : Sun Nov 25 21:19:12 EST 2018
Temperature : UNKNOWN
Solvent :

Compound 44

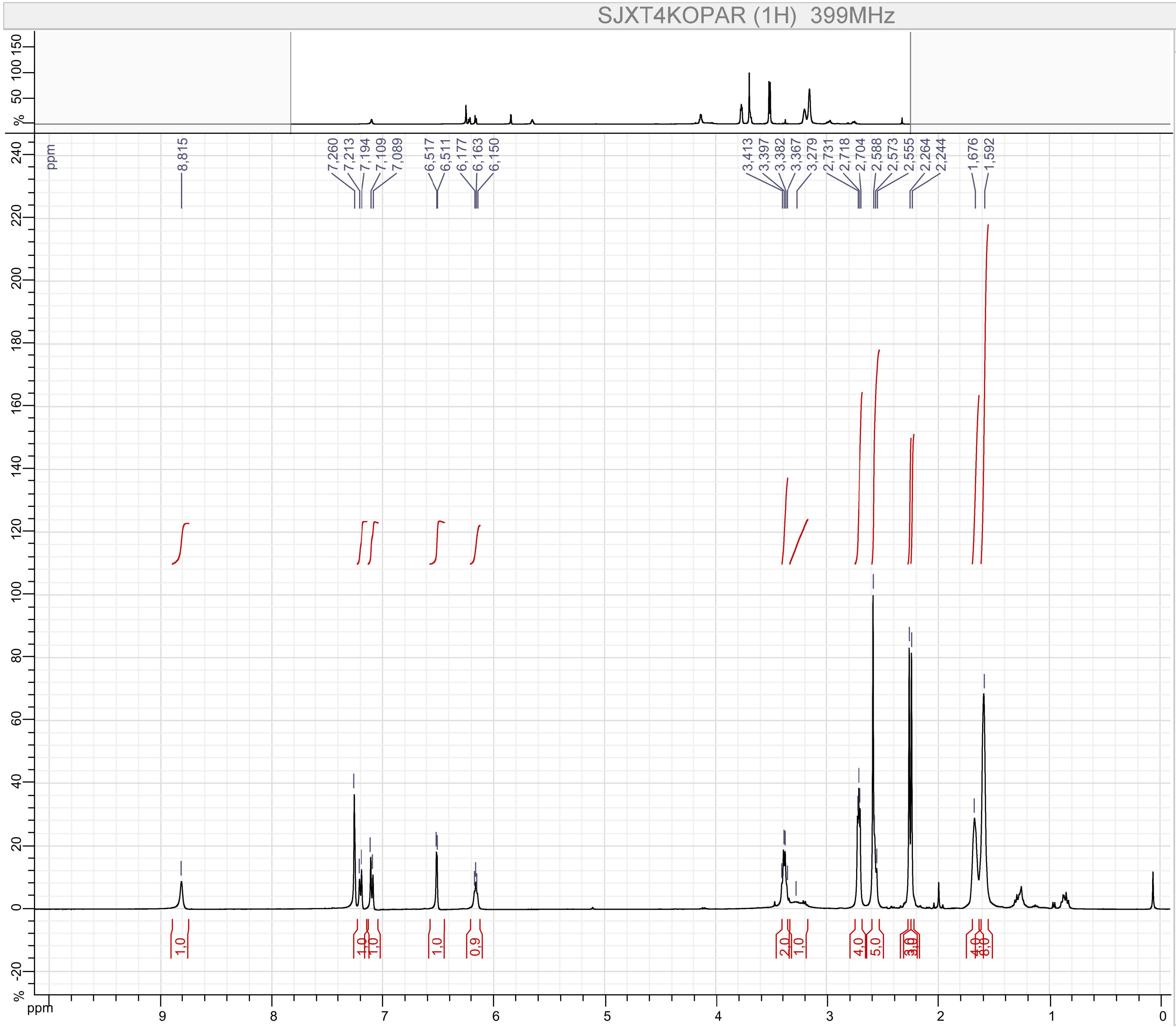


Parameters

SJXT4KOPAR (1H)

F1 :
- Size : 65536 points complex
- Spectral Width : 7494.2334 Hz
- Carrier Frequency : 399.7822 MHz
- Nucleus : 1H

Type : UNKNOWN
Spectro : JEOL 399 MHz
Probe : Unknown
Date : Tue Jul 02 22:40:56 EDT 2019
Temperature : UNKNOWN
Solvent :

Compound 44

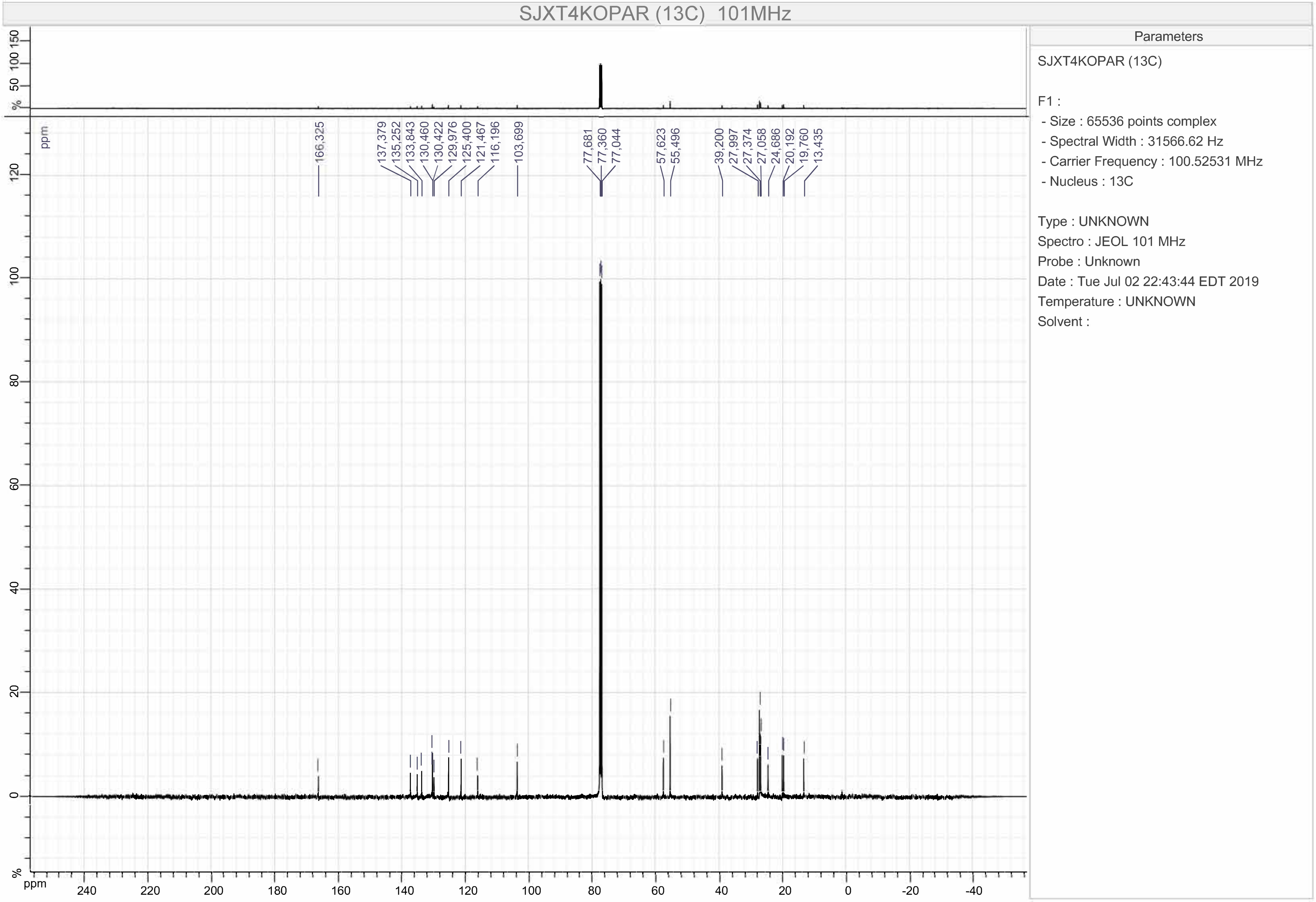

SJXT4KOPAR (13C) 101MHz
166,325
137,379
135,252
133,843
130,460
130,422
129,976
125,400
121,467
116,196
103,699
77,681
77,360
77,044
57,623
55,496
39,200
27,997
27,374
27,058
24,686
20,192
19,760
13,435
ppm
%
Parameters
SJXT4KOPAR (13C)
F1 :
- Size : 65536 points complex
- Spectral Width : 31566.62 Hz
- Carrier Frequency : 100.52531 MHz
- Nucleus : 13C
Type : UNKNOWN
Spectro : JEOL 101 MHz
Probe : Unknown
Date : Tue Jul 02 22:43:44 EDT 2019
Temperature : UNKNOWN
Solvent :

Compound 45

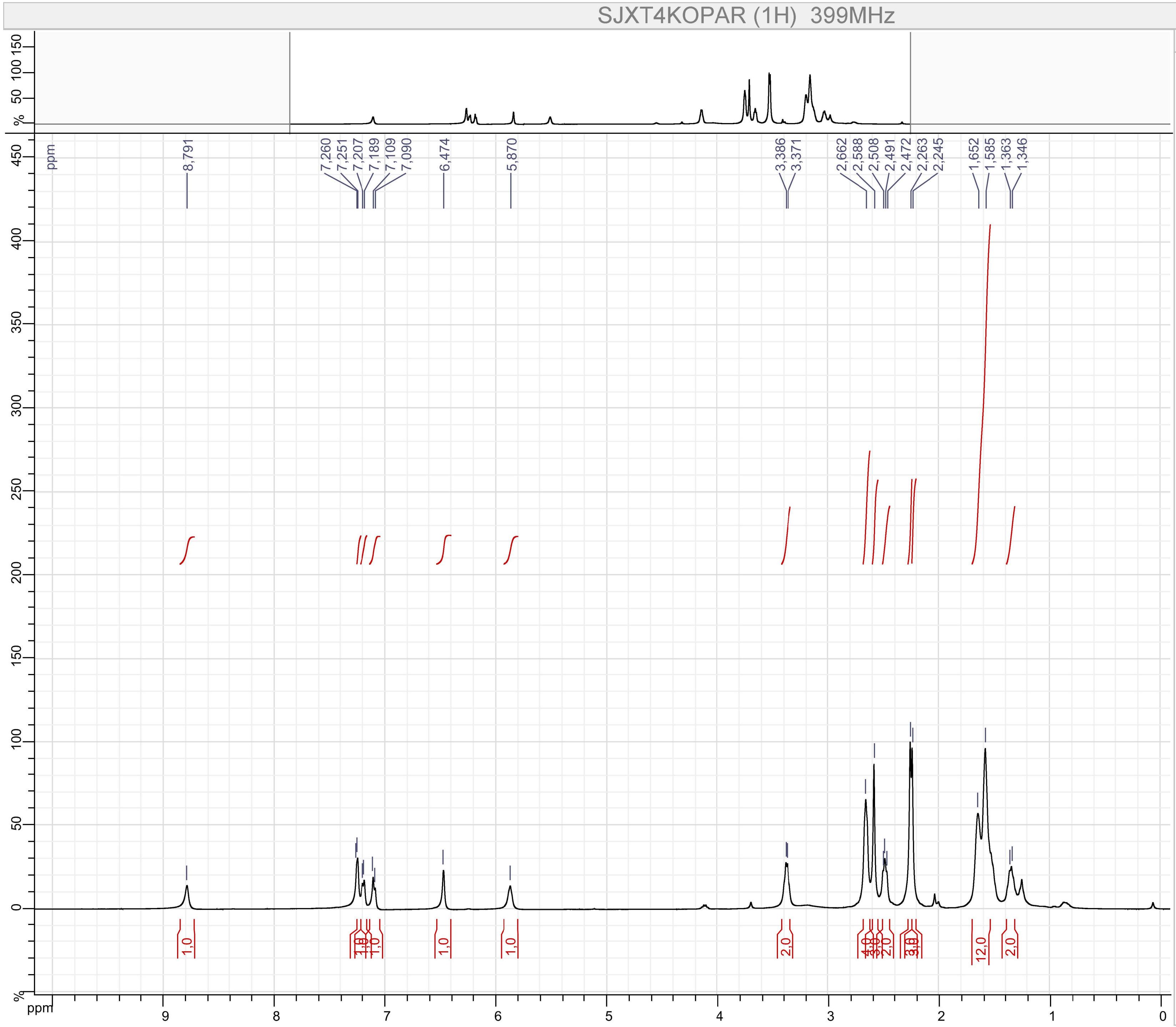


Parameters

SJXT4KOPAR (1H)

F1 :
- Size : 65536 points complex
- Spectral Width : 7494.2334 Hz
- Carrier Frequency : 399.7822 MHz
- Nucleus : 1H

Type : UNKNOWN
Spectro : JEOL 399 MHz
Probe : Unknown
Date : Fri Sep 06 22:00:34 EDT 2019
Temperature : UNKNOWN
Solvent :

Compound 45

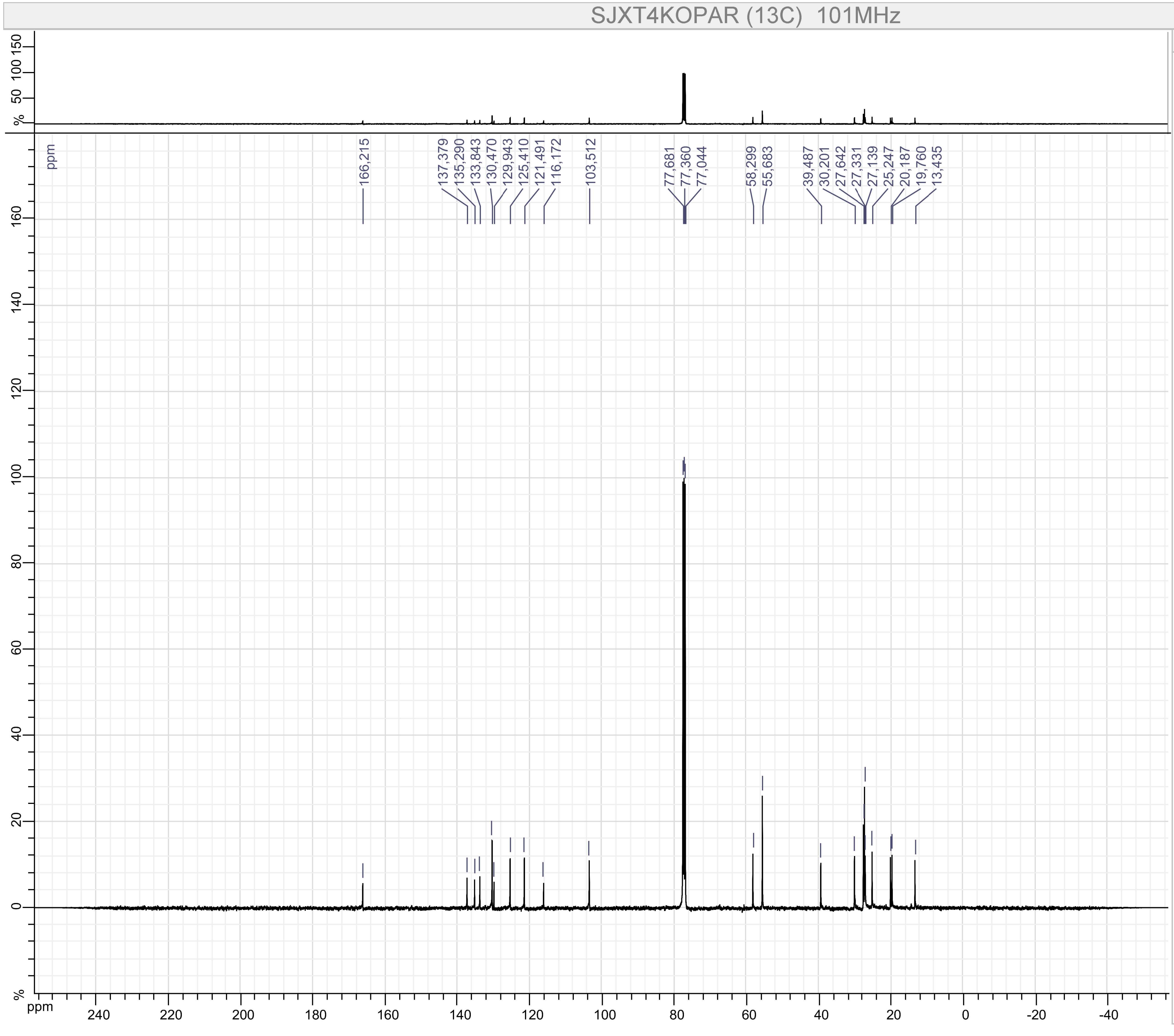


Parameters

SJXT4KOPAR (13C)

F1 :
- Size : 65536 points complex
- Spectral Width : 31566.62 Hz
- Carrier Frequency : 100.52531 MHz
- Nucleus : 13C

Type : UNKNOWN
Spectro : JEOL 101 MHz
Probe : Unknown
Date : Fri Sep 06 22:03:31 EDT 2019
Temperature : UNKNOWN
Solvent :
